# Centered cross-context perturbation transfer on the public 2025 Virtual Cell Challenge benchmark

**DOI:** 10.64898/2026.10.20.760514

**Authors:** Haoyang Luan, Letian Chen, Gufeng Yu, Yang Yang

**Author notes:** Correspondence: Yang Yang.

## Abstract

Predicting transcriptional responses across cellular contexts requires transferring both expression direction and differential-expression priority. We developed a compact numerical method that combines public perturbation summaries with a differential-expression kernel fitted to H1 CRISPRi training targets. The method centers an independent KOLF2.1J source on known training targets, transfers its response signs and magnitude ranks, and uses isotonic regression to render predicted expression means. We evaluated the frozen model retrospectively on the released public TEST100 of the 2025 Virtual Cell Challenge, using an external KOLF source released in 2026. The model achieved DES 0.371834, PDS 0.8614, and MAE 0.302206. With reconstructed historical anchors, its composite score was 0.341344, compared with the archived champion score of 0.340052, a difference of +0.001292. The comparison showed higher DES and lower PDS, with zero normalized MAE contributions for both methods. Source centering also improved development PDS across all three TRAIN folds and official VALIDATION. These results demonstrate a practical numerical approach to cross-context perturbation transfer and differential-expression ranking.

## Introduction

Perturbation prediction asks whether a measured response in one setting can inform an unmeasured response in another. The 2025 Virtual Cell Challenge formalized this problem using CRISPRi perturbations in H1 human embryonic stem cells. It released 150 TRAIN targets, 50 VALIDATION targets, and 100 TEST targets over 18,080 genes [1]. Public perturbation resources provide complementary evidence from other cell backgrounds [3-6]. Their numerical scales, common responses, and missing measurements differ.

The benchmark evaluates three properties. Differential Expression Score (DES) measures recovery of significant genes. Perturbation Discrimination Score (PDS) measures retrieval of the corresponding real perturbation. Mean Absolute Error (MAE) measures pseudobulk expression accuracy [2,8,9]. These objectives need not improve together. L1-based PDS is sensitive to response scaling [7]. The aggregate score also truncates negative contributions relative to baseline, so MAE can cease to influence ranking [8].

We separated response direction from DE priority. Public response summaries supply expression patterns, while a kernel fitted to H1 TRAIN labels predicts DE rankings. We then centered the KOLF source on known training targets before transferring its signs and magnitude ranks. This operation uses no query expression. An isotonic renderer combines the adapted directions with the learned DE order.

The resulting method is deterministic and numerical. It does not use a prediction agent, protein embeddings, or language-model embeddings. We report its development behavior, frozen public TEST evaluation, and exact comparison with the published champion aggregate. Our focus is the measured effect of source centering and rank-constrained rendering, rather than reconstruction of an unpublished competition model.

## Results

### Public TEST performance reflects a DES-PDS tradeoff

Frozen V5 achieved a reconstructed historical score of 0.341344449 on public TEST100. The archive reports 0.340052247 for BM_xTVC, model xTrimoSCPerturb, submission xTVCM_v01 [2]. The numerical difference was +0.001292202.

V5 increased raw DES by 0.023012925654259553 and decreased PDS by 0.0106 relative to the archive. Its MAE was lower, but both MAEs exceeded the fixed anchor and therefore contributed zero to the aggregate. The DES component increased by 0.0254300841, whereas the PDS component decreased by 0.0215534770. Their equally weighted combination explains the small overall difference.

**Figure 1.**
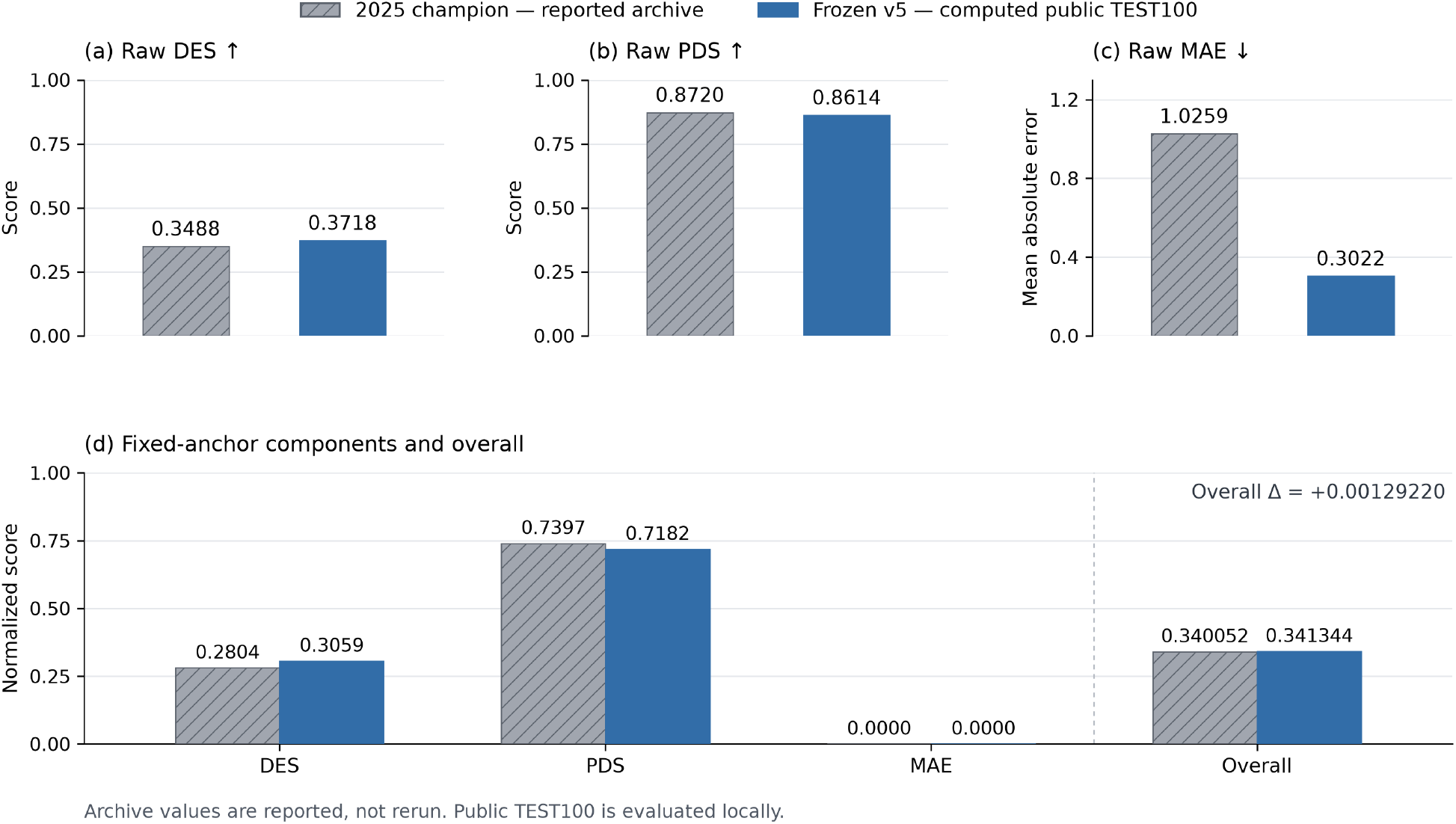
Archived and locally computed TEST performance. Panels show raw DES, PDS, and MAE, followed by the fixed-anchor normalized components and overall score. Grey hatched bars represent the archive-reported BM_xTVC/xTVCM_v01 values. Blue bars represent frozen V5 evaluated locally on public TEST100. Higher DES and PDS are favorable; lower MAE is favorable. Both normalized MAE contributions are zero. The comparison uses reconstructed historical anchors. No uncertainty bars are shown.

**Figure 2.**
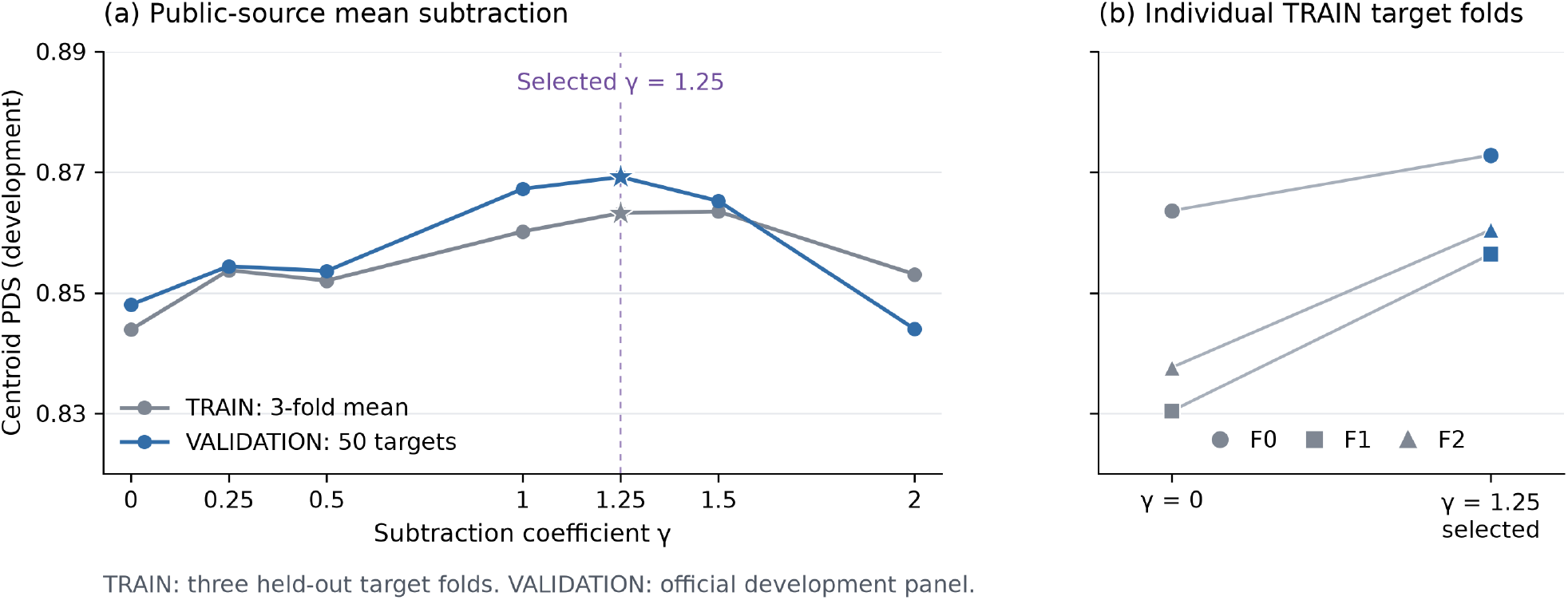
Development effect of KOLF source centering. The first panel plots mean TRAIN three-fold and official VALIDATION centroid PDS over coefficients 0, 0.25, 0.5, 1, 1.25, 1.5, and 2. The selected coefficient 1.25 is marked. The second panel shows the three actual TRAIN fold pairs from coefficient 0 to 1.25: F0, 0.8636 to 0.8728; F1, 0.8304 to 0.8564; F2, 0.8376 to 0.8604. These are development measurements, not confidence intervals. Centering affected source signs, magnitude ranks, and threshold eligibility together.

### Centering improved development discrimination

Subtracting the KOLF source mean estimated from available fit targets improved development PDS. Increasing the centering coefficient from 0 to 1.25 raised mean TRAIN three-fold PDS from 0.8438666667 to 0.8632. The corresponding fold changes were 0.8636 to 0.8728, 0.8304 to 0.8564, and 0.8376 to 0.8604. VALIDATION PDS increased from 0.848 to 0.8692.

Native VALIDATION DES changed from 0.3947980235 to 0.3944960522. MAE decreased from 0.2923363712 to 0.2846686323. The reconstructed development score increased from 0.3407206724 to 0.3549784275. A coefficient of 1 gave PDS 0.8672 and score 0.3536623563. The effect was not monotone. At coefficient 2, TRAIN PDS was 0.8530666667 and VALIDATION PDS was 0.844.

Raising only the uncentered source threshold to 0.12, 0.16, or 0.24 reduced the pooled development objective. Threshold adjustment alone therefore did not reproduce the centering gain. Centering changes signs, magnitude ranks, and threshold eligibility together. This comparison does not isolate a biological mechanism.

For selected V5, TRAIN three-fold MAE was 0.3872227989 and the DE-topN proxy was 0.3876358717. Native TRAIN DES was not computed. The selection objective was 0.3500948953 and used the top-N proxy. Native VALIDATION confirmation was a separate calculation.

### Sequential evaluation and computational cost

Four frozen versions were evaluated on public TEST100 (Table 2). Each prediction was fixed before its corresponding evaluation. These measurements document the sequential evaluation history rather than independent replications.

**Table 1.** Archived champion and locally evaluated V5. Values are rounded here. Champion values are archive-reported, whereas V5 values were computed on the released public TEST100 using the stated evaluator and reconstructed anchors.

| Quantity | Archived BM_xTVC | Frozen V5 |
| --- | --- | --- |
| Raw DES | 0.348821552 | 0.371834478 |
| Raw PDS | 0.872000000 | 0.861400000 |
| Raw MAE | 1.025874829 | 0.302205621 |
| Normalized DES | 0.280425142 | 0.305855227 |
| Normalized PDS | 0.739731598 | 0.718178121 |
| Normalized MAE | 0 | 0 |
| Overall | 0.340052247 | 0.341344449 |

**Table 2.** Recorded public TEST evaluations. All scores use the same fixed reconstructed anchors. Rows identify successive frozen versions, not independent replicates.

| Version | DES | PDS | MAE | Score |
| --- | --- | --- | --- | --- |
| v1 | 0.370076922852 | 0.8149 | 0.194522297233 | 0.309180186089 |
| v2 | 0.370815295399 | 0.8366 | 0.233661419787 | 0.324160037710 |
| v3 | 0.371847148058 | 0.8401 | 0.314435694180 | 0.326912353452 |
| v5 | 0.371834477549 | 0.8614 | 0.302205620930 | 0.341344449246 |

The coefficient-1 v4 prediction was generated but never TEST-scored. Coefficient 1.25 superseded it before centered-model TEST evaluation.

The final inference job completed in 11.72 s, including exact VALIDATION replay and TEST prediction generation. Native public TEST scoring took 158.669 s, including 66.174 s for the target Mann-Whitney U map. It used 15 spawned workers and a reused exact reference cache. Peak reported parent RSS was 14,684,912 KiB, not total process-group memory. These measurements exclude source downloads and the preceding development search.

## Methods

### Computational assistance

OpenAI Codex assisted code development, experiment orchestration, and analysis. ChatGPT, with GPT-6 displayed as the selected model, provided technical consultation and manuscript drafting. Numerical claims were checked against saved computation reports. Prompt and reply records were archived. The final frozen predictor is numerical and does not call a language model.

### Data, splits, and preprocessing

We used the released H1 CRISPRi TRAIN, VALIDATION, and TEST objects [1]. Development used three fixed target folds. Each F100 fit predicted its disjoint V50. The KOLF response center and H1 fitted response statistics used the corresponding fit targets. Fixed public Replogle scalers were retained. Final inference refitted on all 150 TRAIN targets. VALIDATION was repeatedly used for development and selection.

The retrospective TEST series contained four sequential frozen evaluations. The coordinator and evaluator saw earlier TEST aggregates. Model-development agents did not receive TEST expression or evaluation reports, and no TEST per-target analysis was used. Prediction freezing before each evaluation did not make the overall study a single untouched blind test.

TRAIN teacher profiles were computed by normalizing individual cells to 54,611, applying log1p, and averaging within targets. Teacher means were Float32. This is not log transformation of a pseudobulk count mean. The prediction reference comprised 38,176 TRAIN non-targeting cells. Let *c* be their mean and 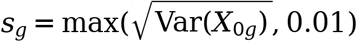.

Native real-object evaluation used the median positive library total of each original full object before log1p Float32 conversion. This value was 54,312 for VALIDATION and 54,125.5 for TEST. Predictions retained the TRAIN-derived scale. Real and predicted controls were handled separately.

Replogle inputs comprised K562 genome-wide, RPE1, and K562 essential GEM-Z summaries [3]. X-Atlas inputs were HCT116 and HEK293T mean-log1p CP10k differences [6]. These source units were kept separate. KOLF inputs came from the author viewer, not automatically from the journal analysis pipeline [4,5]. Its 10,167-target by 38,606-gene panel stores signed int8 values multiplied by 0.04. It lacks the original finite/missing mask. Quantized zeros were not treated as proof of original missingness. Identity matching remained literal and audited; TAZ was unresolved.

KOLF was available for 138 of 150 TRAIN targets and 91 of 100 TEST targets. Missing-source queries retained the older prediction branches. The current KOLF product is a 2026 release, so this study does not represent a method available by the original 2025 deadline.

### Numerical expression initializer

Let Π clip values to [0,14.999999046325684]. The base *B* used full GEM-Z feature axes of 8,248, 8,749, and 8,563 genes without PCA. Original own-source coordinates were retained. Fixed public centers and population standard deviations standardized observed features. Scaler genes required at least two observations and variance above 10^−24^. Missing numerical entries were zero after standardization, with row availability tracked separately.

For source *j*, center its *n*_*j*_ available fit feature rows to obtain *X*_*j*_. Let *Y*_*j*_ contain H1 teacher deltas from *c*, with mean 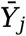. Define 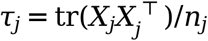.

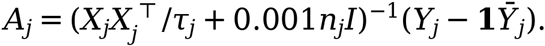

For a query feature vector *X*_*j*_ (*q*) and fit feature center 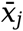,

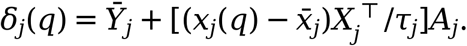

A source was enabled only when *n*_*j*_ ≥ and τ_*j*_*>*10^−24^. Each expert became max(*c* + *δ*_*j*_, 0) − *c* before averaging. Available experts received equal weight. With no available expert, the fallback was the all-fit mean response. Adding *c* and applying Π produced *B*. The solve and base storage used Float64. No residual-memory or agent correction entered *B*.

For the complete query prediction panel, let 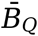 be the mean predicted vector. We formed

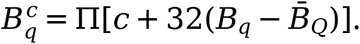

For the mean available GEM-Z response *Z* _*q*_,

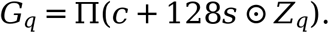

Missing and own-target coordinates in *G*_*q*_ retained *B*_*q*_. We then computed

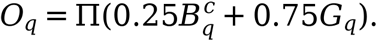

For the mean available X-Atlas delta 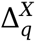,

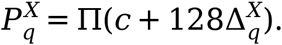

Missing and own-target coordinates in 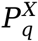 retained *O q*. The initializer used for V5 was

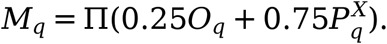

The *B*^*c*^,*G,O*, and *M* stages used Float32 encodings. Query-panel centering used predictions only.

### Centered source quantile transfer

For each gene, we averaged finite observed KOLF values across source-known fit targets to obtain 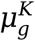. Coordinates with no observations used zero. The final centered source was

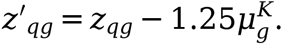

Availability masks were unchanged. Eligible coordinates required an available source value, *c*_*g*_ > 0, *M*_*qg*_ > 0, and exclusion of the query’s own gene.

For eligible genes, let 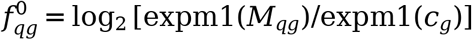. We ranked ~*z’_qg_*~ with average ties and divided by the eligible-gene count to obtain *r*_*qg*_. If *Q*_*q*_ is the linear empirical quantile function of eligible 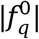, the transferred fold change was

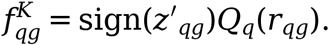

We decoded signed fold changes using

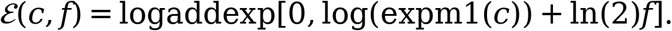

The substitution was applied only when ~*z*’_qg_~ ≥Float32(0.08). Weak or missing coordinates retained exact *M* before rendering. Ranks and quantiles were computed on the full eligible set before thresholding. This transfers relative magnitudes without equating KOLF z-scores with H1 log-expression differences.

### Independent DE kernels and isotonic rendering

Within each fit set, raw DE p values received a complete target-by-gene BH correction. Let *D* be the strict FDR <0.05mask and *f* its target mean.

For the original DE kernel, we aligned the three GEM-Z blocks to 18,080 genes, zero-filled missing coordinates, L2-normalized each block per target, and concatenated them. Their Gram matrix was divided by the mean diagonal of its fit block to obtain *K*. DE scores were

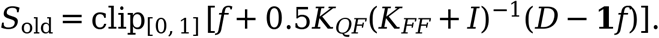

A separate KOLF kernel used uncentered aligned source vectors, missing entries zero, own-input coordinates zero, and per-target L2 normalization. It used only source-known fit rows, their label mean *fK*, and the same trace scaling and ridge equation. KOLF scores were encoded as Float32 before promotion to Float64.

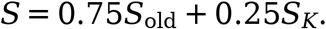

Queries without KOLF support retained *S*_old_. The expression-centering coefficient did not alter this kernel.

For each target, *u*_*qg*_ was the average rank of *S*_*qg*_ divided by 18,080. An increasing, unweighted `IsotonicRegression` fitted *u*_*q*_ to the actual absolute log2-FC of the adapted expression over genes with *c*_*g*_ > 0. It predicted all *u*_*q*_ with boundary clipping. If the fitted function is *h*_*q*_,

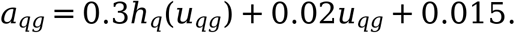

The final mean was

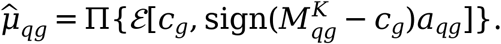

Here *M* ^*K*^ denotes the quantile-adapted expression. At *c*_*g*_ = 0, the decoded mean is zero. The literal own gene was set to zero when *c*_*g*_ > 0, excluding unresolved TAZ. This override does not directly affect PDS because that coordinate is excluded. Final means were Float32.

### Development selection and evaluation

We examined centering coefficients 0, 0.25, 0.5, 1, 1.25, 1.5, and 2. Development selection used

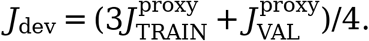

The DE-topN proxy ranked genes by actual absolute FC and used the real DEG count, without the complete native predicted-side significance calculation. It was recorded separately from DES. Native VALIDATION subsequently confirmed the selected recipe. Earlier development rows used their own specified mixing and curve settings.

One mean vector was repeated to each target’s independently public cell count and appended to the original TRAIN controls. The evaluated TEST representation comprised 132,670 query cells and 38,176 controls, totaling 170,846 cells. It represents repeated pseudobulk profiles, not sampled cellular heterogeneity.

Evaluation used the original `vcc` profile [8,9]. DE used two-sided nonexperimental Mann-Whitney U tests with tie correction, `is_log1p=True`, and `exp_post_agg=True`. Real and predicted sides each received global BH correction across their complete panel. TEST contained 1,808,000 hypotheses and VALIDATION 904,000.

Fold change was expml(mean log target)/expm1(mean log control). Both zero gave 1. Reference zero with positive target gave 20; target zero with positive reference gave 0.05. Other ratios were not clipped. DE numerics were converted to Float32. DES filtered strict FDR <0.05, sorted by absolute log2-FC, and divided overlap at the real significant-gene count by that count. A shorter predicted list did not change the denominator.

PDS used L1 distances between independently control-subtracted pseudobulk profiles. It excluded the query’s literal own-gene coordinate and ranked its paired real target among the complete real panel. PDS was*1-r/N* for zero-based rank *r*. MAE used all-gene pseudobulk means.

Exact constant-target MWU preserved installed typed arithmetic. Native DES and MAE/PDS functions calculated final metrics. Original-order Polars Float32 group means were computed in gene-column blocks. No full submission H5AD or private-runner bridge was used.

With anchors *b*_*D*_ = 0.09505113849605429, bp = 0.5082, and *b*_*M*_ *=* 0.0249575331993401,

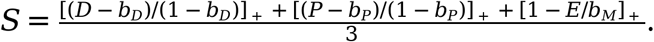

Here [*x*] + = max(0, *x*), and *D,P,E* are target-averaged DES, PDS, and MAE. Anchors were algebraically recovered from 100 archive rows, not obtained from an original baseline CSV. The MAE anchor was identified from one archived row with a nonzero MAE component.

## Discussion

Centering provides a small, effective adjustment to cross-context response transfer. It removes a recurring source component estimated from known fit targets before using query-specific signs and magnitude ranks. The selected coefficient exceeded one, so the operation was not simple mean subtraction. It could reverse weak responses and change which coordinates passed the strong-source threshold. The measured improvement supports this preprocessing choice for the studied data, not a mechanistic interpretation of the removed component.

The method also separates information sources by role. Direct public profiles provide directions and relative amplitudes. H1-supervised kernels provide DE priority. Isotonic rendering connects these quantities without a large learned expression decoder. Keeping the KOLF DE kernel uncentered distinguishes response adaptation from target-similarity estimation.

The benchmark result depends on its objective. L1 retrieval is scale-sensitive [7], and DES depends on both significance and effect ranking. Repeated mean profiles can score well without representing single-cell variability. V5’s lower MAE than the archive did not contribute to its score. Both normalized MAE terms were zero. Thus the numerical advantage reflects the DES-PDS balance, not uniform superiority across predictive properties.

External same-target responses make this cross-context transfer, not prediction of targets unseen in every dataset. Quantization and missingness in the derived KOLF viewer further constrain what can be inferred from individual source values. The initializer also depends on the requested query panel through its prediction mean. The KOLF center instead depends only on known fit targets. These two centering operations have different information boundaries.

This study compares the frozen predictor on released public TEST100 with reported archived benchmark values. Reconstructed historical anchors provide the fixed scoring scale. Together with the development analyses, the comparison characterizes the DES-PDS tradeoff of centered cross-context transfer and rank-constrained rendering under the specified public-data evaluation.

## Supporting information

Supplementary reproducibility code and frozen predictions

## Data and code availability

The accompanying 7.6 MB code archive contains the frozen inference and scoring code, version pins, the exact 100-target TEST prediction in NPZ format, the fitted source center, figures, and supporting evidence.

The compact archive omits large input matrices. These remain available from the original providers for the Virtual Cell Challenge [1], Replogle perturbation screens [3], KOLF2.1J [5], and X-Atlas/Orion [6].

## Reproducibility appendix

Inference used Python 3.11.10 and scikit-learn 1.9.1. Evaluation used cell-eval 0.6.6, pdex 0.1.27, NumPy 2.4.6, SciPy 1.17.1, Polars 2.0.0, and Scanpy 1.11.5. These versions identify the present computation, not the historical private environment.

Independent VALIDATION replay reproduced all 904,000 predicted Float32 mean entries exactly. Process/native raw-DE parity covered 904,000 VALIDATION rows. The prediction SHA256 is `cea4874ec557b56622d66e5a414ebe8ecb59fee078e5f613a3316cbc9fffb4df.` The TEST summary SHA256 is `b681574e7e6a484110597c5442cb8121abe58ae6cf344cfae00c7dbbb03f9362.` Full source, input-object, fitted-center, and runtime records are in `final_results_v5.json`.

