## Supplementary reproducibility code and frozen predictions for "Centered cross-context perturbation transfer on the public 2025 Virtual Cell Challenge benchmark": prepare_training.py

```
"""Stream training counts into a compact, explicitly normalized model dataset.

This representation is for target-level CV and numerical modeling. Its 10,000
count normalization is explicit and is not a historical leaderboard score.
The final evaluator's native preprocessing is recorded separately.
"""
from __future__ import annotations

import argparse
from datetime import datetime, timezone
import hashlib
import json
from pathlib import Path


def sha256(path: Path) -> str:
    digest = hashlib.sha256()
    with path.open("rb") as stream:
        while block := stream.read(8 * 1024 * 1024):
            digest.update(block)
    return digest.hexdigest()


def prepare(source: Path, output: Path, target_sum: float, chunk_rows: int) -> dict:
    import anndata as ad
    import numpy as np
    from scipy import sparse

    output.mkdir(parents=True, exist_ok=True)
    data = ad.read_h5ad(source, backed="r")
    try:
        if "target_gene" not in data.obs:
            raise ValueError(f"Missing target_gene. Actual obs columns: {list(data.obs)}")
        labels = data.obs["target_gene"].astype(str).to_numpy()
        groups, codes = np.unique(labels, return_inverse=True)
        if "non-targeting" not in groups:
            raise ValueError("Required control identity non-targeting is missing")
        expected = set(json.loads((output.parent.parent / "manifests" /
                                  "training_target_identities.json").read_text(
                                      encoding="utf-8-sig"))["target_genes"])
        actual = set(groups) - {"non-targeting"}
        if actual != expected:
            raise ValueError(f"Training target mismatch missing={expected-actual}, extra={actual-expected}")
        n_groups, n_genes = len(groups), data.n_vars
        gene_names = data.var_names.astype(str).to_numpy()
        if len(set(gene_names)) != n_genes:
            raise ValueError("Duplicate gene identities")
        count = np.bincount(codes, minlength=n_groups)
        sums = np.zeros((n_groups, n_genes), dtype=np.float64)
        squares = np.zeros_like(sums)
        raw_sums = np.zeros_like(sums)
        row_umi = np.empty(data.n_obs, dtype=np.float64)
        rng = np.random.default_rng(20251013)
        control_rows = np.flatnonzero(labels == "non-targeting")
        chosen_control = np.sort(rng.choice(control_rows,
                                           size=min(512, len(control_rows)), replace=False))
        controls = np.empty((len(chosen_control), n_genes), dtype=np.float32)
        control_lookup = {int(row): i for i, row in enumerate(chosen_control)}
        sample_min = float("inf")
        sample_max = 0.0
        for start in range(0, data.n_obs, chunk_rows):
            end = min(start + chunk_rows, data.n_obs)
            raw = data.X[start:end].copy()
            if sparse.issparse(raw):
                raw = raw.tocsr().astype(np.float32)
                payload = raw.data
                umi = np.asarray(raw.sum(axis=1)).ravel()
            else:
                raw = np.asarray(raw, dtype=np.float32)
                payload = raw
                umi = raw.sum(axis=1, dtype=np.float64)
            if not np.isfinite(payload).all() or np.any(payload < 0):
                raise ValueError("Counts must be finite and nonnegative")
            if np.any(np.abs(payload - np.rint(payload)) > 1e-4):
                raise ValueError("Input is not integer-like raw counts; inspect before preprocessing")
            if payload.size:
                sample_min = min(sample_min, float(payload.min()))
                sample_max = max(sample_max, float(payload.max()))
            row_umi[start:end] = umi
            design = sparse.csr_matrix((np.ones(end-start),
                                       (codes[start:end], np.arange(end-start))),
                                      shape=(n_groups, end-start))
            aggregate = design @ raw
            raw_sums += aggregate.toarray() if sparse.issparse(aggregate) else aggregate
            scale = target_sum / np.maximum(umi, 1.0)
            if sparse.issparse(raw):
                normal = raw.multiply(scale[:, None]).tocsr()
                normal.data = np.log1p(normal.data)
            else:
                normal = np.log1p(raw * scale[:, None])
            aggregate = design @ normal
            sums += aggregate.toarray() if sparse.issparse(aggregate) else aggregate
            square = normal.multiply(normal) if sparse.issparse(normal) else np.square(normal)
            aggregate = design @ square
            squares += aggregate.toarray() if sparse.issparse(aggregate) else aggregate
            for row in chosen_control[(chosen_control >= start) & (chosen_control < end)]:
                sample = normal[int(row)-start]
                controls[control_lookup[int(row)]] = sample.toarray().ravel() if sparse.issparse(sample) else sample
            print(f"training rows {end}/{data.n_obs}", flush=True)
        means = sums / count[:, None]
        variance = np.maximum(squares / count[:, None] - means**2, 0)
        np.savez_compressed(output / "training_pseudobulk.npz",
                            groups=groups.astype(str), genes=gene_names.astype(str),
                            means=means.astype(np.float32), variances=variance.astype(np.float32),
                            raw_means=(raw_sums / count[:, None]).astype(np.float32),
                            cell_counts=count, controls=controls,
                            control_row_ids=chosen_control, target_sum=np.array(target_sum))
        audit = {
            "created_utc": datetime.now(timezone.utc).isoformat(),
            "source_path": str(source), "source_sha256": sha256(source),
            "source_bytes": source.stat().st_size,
            "n_cells": data.n_obs, "n_genes": n_genes,
            "n_targets": len(actual), "obs_columns": list(data.obs),
            "var_columns": list(data.var), "layers": list(data.layers),
            "control_label": "non-targeting", "control_cells": len(control_rows),
            "normalization": {"input": "integer raw counts", "target_sum": target_sum,
                              "transform": "log1p", "scope": "fixed-scale model representation",
                              "official_score": False},
            "observed_count_min": sample_min, "observed_count_max": sample_max,
            "median_library_size": float(np.median(row_umi)),
            "control_reservoir_seed": 20251013,
            "target_counts": dict(zip(groups, count.astype(int).tolist())),
            "output_sha256": sha256(output / "training_pseudobulk.npz")
        }
        (output / "training_inventory.json").write_text(json.dumps(audit, indent=2)+"\n",
                                                       encoding="utf-8")
        return audit
    finally:
        data.file.close()


def main() -> None:
    parser = argparse.ArgumentParser()
    parser.add_argument("--root", type=Path, default=Path("/data1/luanhaoyang/virtualcell"))
    parser.add_argument("--target-sum", type=float, default=10000.0)
    parser.add_argument("--chunk-rows", type=int, default=1024)
    args = parser.parse_args()
    if args.target_sum <= 0 or args.chunk_rows < 1:
        parser.error("target-sum and chunk-rows must be positive")
    root = args.root.resolve()
    if root != Path("/data1/luanhaoyang/virtualcell"):
        raise ValueError("Use only the authorized remote project root")
    result = prepare(root / "data/train/adata_Training.h5ad",
                     root / "data/processed", args.target_sum, args.chunk_rows)
    print(json.dumps({k: result[k] for k in ("n_cells", "n_genes", "n_targets", "output_sha256")}))


if __name__ == "__main__":
    main()
```
