## Supplementary reproducibility code and frozen predictions for "Centered cross-context perturbation transfer on the public 2025 Virtual Cell Challenge benchmark": fast_gct_de_krr_rank_20261020.py

```
"""TRAIN-only target-fold KRR on existing cross-cell source effects and F100 D.

Each original F100 label mask D is fitted; its V50 labels are used only to
measure top-N overlap. This is a ranking proxy, never official DES.
"""
from __future__ import annotations

from datetime import datetime, timezone
import hashlib
import json
import os
from pathlib import Path
import sys
import time

import numpy as np

ROOT=Path('/data1/luanhaoyang/virtualcell')
BANK=ROOT/'data/processed/gct_TRAIN_raw_bank_v1/raw_bank.npz'
SCOPE=ROOT/'runs/training150_protein_prior_v2/prediction_scope_metadata.json'
OUT=ROOT/'runs/fast_gct_de_krr_rank_20261020'
ALPHAS=(.001,.01,.1,1.,10.)
BETAS=(.5,1.,2.)


def sha256(p):
    h=hashlib.sha256()
    with p.open('rb') as f:
        for block in iter(lambda:f.read(8*1024**2),b''):
            h.update(block)
    return h.hexdigest()


def build_source_gram(z,mask):
    blocks=[]
    for source in range(3):
        effect=np.where(mask[:,:,source],z[:,:,source],0.).astype(np.float32)
        norm=np.sqrt(np.sum(effect.astype(np.float64)**2,axis=1))
        effect/=np.maximum(norm[:,None],1e-12).astype(np.float32)
        blocks.append(effect)
    x=np.concatenate(blocks,axis=1)
    gram=(x@x.T).astype(np.float64)
    if not np.isfinite(gram).all() or np.max(np.abs(gram-gram.T))>1e-4:
        raise ValueError('Invalid source Gram')
    return (gram+gram.T)/2.


def main():
    if os.name!='posix' or not set(range(24,32))<=os.sched_getaffinity(0):
        raise RuntimeError('99.32 CPU24-31 only')
    os.sched_setaffinity(0,set(range(24,32)))
    sys.path.insert(0,str(ROOT/'ops'))
    from fast_gct_de_topn_proxy_20261020 import read_true_de,topn
    started=time.monotonic()
    with np.load(BANK,allow_pickle=False) as a:
        targets=tuple(map(str,a['targets']))
        genes=tuple(map(str,a['genes']))
        z=a['source_delta'].copy()
        mask=a['source_mask'].copy()
    if len(targets)!=150 or len(genes)!=18080 or z.shape!=(150,18080,3) or mask.shape!=z.shape:
        raise ValueError('Original source bank axes differ')
    gram=build_source_gram(z,mask)
    diag=np.diag(gram)
    zero_source_targets=int(np.sum(diag<=1e-12))
    cosine=gram/np.sqrt(np.maximum(diag[:,None],1e-12)*np.maximum(diag[None,:],1e-12))
    scope=json.loads(SCOPE.read_text(encoding='utf-8'))
    fold_data=[]
    records=[]
    for fold in scope['folds']:
        name=fold['fold_id'];query=tuple(fold['query_targets'])
        prior=ROOT/f'runs/training150_DE_macro_calibration_v1/priors/{name}_outer_fit/prior.npz'
        real=ROOT/f'runs/training150_DE_macro_outer_scoring_v1/panels/{name}/real_query_family_DE.parquet'
        with np.load(prior,allow_pickle=False) as a:
            fit=tuple(map(str,a['fit_targets']))
            if set(fit)!=set(targets)-set(query) or tuple(map(str,a['genes']))!=genes:
                raise ValueError('F100 prior axes differ')
            d=a['D'].astype(np.float64)
            f=a['f'].astype(np.float64)
        if d.shape!=(100,18080) or not np.allclose(f,d.mean(axis=0),atol=1e-12):
            raise ValueError('F100 D frequency differs')
        truth=read_true_de(real,query,genes)
        fi=np.array([targets.index(t) for t in fit])
        qi=np.array([targets.index(t) for t in query])
        fold_data.append({'name':name,'prior':prior,'real':real,'fit':fi,'query':qi,
                          'D':d,'freq':f,'truth':truth})
        baseline=topn(np.broadcast_to(f,truth.shape),truth)
        scores={}
        for kernel_name in ('cosine','trace_linear'):
            scale=1. if kernel_name=='cosine' else float(np.mean(diag[fi]))
            kfit=gram[np.ix_(fi,fi)]/scale if kernel_name=='trace_linear' else cosine[np.ix_(fi,fi)]
            kq=gram[np.ix_(qi,fi)]/scale if kernel_name=='trace_linear' else cosine[np.ix_(qi,fi)]
            residual=d-f[None,:]
            for alpha in ALPHAS:
                coef=np.linalg.solve(kfit+alpha*np.eye(len(fi)),residual)
                correction=kq@coef
                for beta in BETAS:
                    predicted=np.clip(f[None,:]+beta*correction,0.,1.)
                    key=f'{kernel_name}_a{alpha:g}_b{beta:g}'
                    score=topn(predicted,truth)
                    scores[key]=score
        record={'fold':name,'baseline_prior':baseline,'scores':scores,
                'gram_diagonal_mean_fit':float(np.mean(diag[fi])),
                'gram_offdiagonal_mean_fit':float(np.mean(gram[np.ix_(fi,fi)]-np.diag(diag[fi]))),
                'real_DE_sha256':sha256(real),'F100_prior_sha256':sha256(prior)}
        records.append(record)
        best=max(scores,key=lambda k:scores[k]['mean_overlap_nonempty'])
        print(json.dumps({'fold':name,'prior':baseline['mean_overlap_nonempty'],
                          'best':best,'best_topn':scores[best]['mean_overlap_nonempty']}),flush=True)
    if [r['fold'] for r in records]!=['F0','F1','F2']:
        raise ValueError('Original three folds differ')
    keys=list(records[0]['scores'])
    macro={k:float(np.mean([r['scores'][k]['mean_overlap_nonempty'] for r in records])) for k in keys}
    best=max(macro,key=macro.get)
    kernel_name,alpha,beta=best.split('_a')[0],float(best.split('_a')[1].split('_b')[0]),float(best.split('_b')[1])
    full=np.empty((150,18080),dtype=np.float32)
    for fold in fold_data:
        fi=fold['fit'];qi=fold['query'];f=fold['freq'];d=fold['D']
        scale=1. if kernel_name=='cosine' else float(np.mean(diag[fi]))
        kfit=gram[np.ix_(fi,fi)]/scale if kernel_name=='trace_linear' else cosine[np.ix_(fi,fi)]
        kq=gram[np.ix_(qi,fi)]/scale if kernel_name=='trace_linear' else cosine[np.ix_(qi,fi)]
        coef=np.linalg.solve(kfit+alpha*np.eye(len(fi)),d-f[None,:])
        full[qi]=np.clip(f[None,:]+beta*(kq@coef),0.,1.).astype(np.float32)
    OUT.mkdir(mode=0o700,parents=True,exist_ok=False)
    out=OUT/'best_global_fixed_tuple_rank_scores.npz'
    with out.open('xb') as f:
        np.savez_compressed(f,targets=np.asarray(targets),genes=np.asarray(genes),de_score=full)
    result={'state':'complete','utc':datetime.now(timezone.utc).isoformat(),
            'scope':'TRAIN original 3xF100/V50; F100 D fit only; V50 D top-N evaluation only',
            'official_DES':False,'validation_test_expression_read':False,
            'kernels':'per-target/per-source L2-normalized public GEM-Z effects; cosine or fit-trace-normalized linear',
            'zero_source_targets_fall_back_to_prior':zero_source_targets,
            'alpha_grid':ALPHAS,'beta_grid':BETAS,
            'baseline_prior_macro':float(np.mean([r['baseline_prior']['mean_overlap_nonempty'] for r in records])),
            'macro_scores':macro,'best_global_fixed_tuple':{'key':best,'topn_overlap':macro[best]},
            'best_score_npz':{'path':str(out),'sha256':sha256(out),'key':'de_score',
                              'dtype':str(full.dtype),'shape':list(full.shape)},
            'folds':records,'bank_sha256':sha256(BANK),'elapsed_seconds':time.monotonic()-started}
    with (OUT/'summary.json').open('x',encoding='utf-8') as f:
        json.dump(result,f,indent=2,allow_nan=False);f.write('\n')
    print(json.dumps({'state':'complete','baseline_prior_macro':result['baseline_prior_macro'],
                      'best_global_fixed_tuple':result['best_global_fixed_tuple'],
                      'best_score_npz':result['best_score_npz']}),flush=True)


if __name__=='__main__':main()
```
