## Supplementary reproducibility code and frozen predictions for "Centered cross-context perturbation transfer on the public 2025 Virtual Cell Challenge benchmark": run_numeric_kolf_centered_R1_v5_20261021.py

```
"""Exact selected VALIDATION replay, then prediction-only centered R1 v5 inference."""
from pathlib import Path
import hashlib,json,os,subprocess,time
ROOT=Path('/data1/luanhaoyang/virtualcell');PY=ROOT/'envs/vcc2025-py311/bin/python'
OUT=ROOT/'runs/numeric_kolf_centered_R1_v5_20261021'
os.sched_setaffinity(0,set(range(16,24)));OUT.mkdir(parents=True,exist_ok=False);started=time.monotonic()
def sha(path):return hashlib.sha256(Path(path).read_bytes()).hexdigest()
def run(args):subprocess.run([str(PY),'-B']+[str(x) for x in args],cwd=ROOT,check=True)
script=ROOT/'scripts/predict_numeric_kolf_centered_R1_v5.py'
expected=ROOT/'runs/fast_kolf_public_center_extend_20261021_v1/center_gamma1.25_validation.npz'
if sha(expected)!='419ca8f1f6de0897158b71bea5a10003626660a05ceebe0db63ce5c4cfdb5771':raise ValueError('Wrong selected VALIDATION replay input')
run([script,'--expression-predictions',ROOT/'runs/fast_xatlas_refine_20261020_v1/best_validation.npz',
     '--de-scores',ROOT/'runs/fast_validation_krr_rankshape_20261020_v1/validation_de_scores.npz',
     '--query-kolf-bank',ROOT/'data/processed/kolf2_1j_browser_selected_v1/source_bank.npz',
     '--replay',expected,'--replay-atol','0','--output',OUT/'validation_replay'])
validation=json.loads((OUT/'validation_replay/prediction.json').read_text())
if validation['replay']['changed_entries']!=0:raise RuntimeError('Selected VALIDATION replay changed; no TEST prediction')
run([script,'--features',ROOT/'runs/numeric_test_primary_20261021_v1/predictions.npz',
     '--query-kolf-bank',ROOT/'data/processed/kolf_TEST_public_20261021_v1/source_bank.npz',
     '--output',OUT/'test_predictions'])
receipt={'state':'complete','validation_replay':validation,
         'test_prediction':json.loads((OUT/'test_predictions/prediction.json').read_text()),
         'wrapper_sha256':sha(__file__),'query_expression_or_evaluation_report_read':False,
         'TEST_scoring_run':False,'elapsed_seconds':time.monotonic()-started}
(OUT/'receipt.json').write_text(json.dumps(receipt,indent=2)+'\n');print(json.dumps(receipt),flush=True)
```
