## Supplementary reproducibility code and frozen predictions for "Centered cross-context perturbation transfer on the public 2025 Virtual Cell Challenge benchmark": assemble_manuscript_v5.py

````
"""Extract Edge-authored text and apply recorded factual/typesetting fixes."""
from pathlib import Path
import hashlib, json, re

ROOT = Path(__file__).resolve().parents[1]
EDGE = ROOT / 'paper/edge'


def sha(path):
    return hashlib.sha256(path.read_bytes()).hexdigest()


def block(path):
    text = path.read_text(encoding='utf-8-sig').replace('\r\n', '\n')
    matches = re.findall(r'```markdown\s*\n(.*?)\n```', text, re.S)
    if len(matches) != 1:
        raise ValueError(f'Expected one exact Markdown block: {path}')
    return matches[0].strip()


def main():
    original = EDGE / 'preprint_reply_raw_20261021_P1.md'
    ai = EDGE / 'preprint_reply_raw_20261021_P2_AI.md'
    manuscript = block(original)
    fixes = [
        ('K562 genome-wide day8, RPE1 day7, and K562 essential day6 GEM-Z summaries',
         'K562 genome-wide, RPE1, and K562 essential GEM-Z summaries',
         'Remove unnecessary sampling-day details absent from the verified input facts.'),
        ('; zero real DE count gave zero.', '.',
         'Omit an unverified empty-DE edge-case assertion; all reported target scores are finite.'),
        (r'\mathbf1', r'\mathbf{1}',
         'Brace the math font command without changing the mathematical expression.'),
        ('0.34134444924634444', '0.341344449', 'Round narrative score; full precision remains in evidence.'),
        ('0.3400522468873988', '0.340052247', 'Round narrative archive score; evidence preserves precision.'),
        ('+0.0012922023589456155', '+0.001292202', 'Round narrative score difference.'),
    ]
    applied = []
    for before, after, reason in fixes:
        count = manuscript.count(before)
        manuscript = manuscript.replace(before, after)
        applied.append({'before': before, 'after': after, 'occurrences': count, 'reason': reason})
    insertion = '## Methods\n\n'
    if manuscript.count(insertion) != 1:
        raise ValueError('Unique Methods heading required')
    manuscript = manuscript.replace(insertion,
        insertion + '### Computational assistance\n\n' + block(ai) + '\n\n', 1)
    revision = EDGE / 'preprint_reply_raw_20261021_P3.md'
    revised = re.findall(r'```markdown\s*\n(.*?)\n```',
                         revision.read_text(encoding='utf-8-sig').replace('\r\n', '\n'), re.S)
    if len(revised) != 3:
        raise ValueError('Expected the three Edge P3 replacement blocks')
    manuscript, abstract_count = re.subn(r'(?s)(## Abstract\n\n).*?(?=\n\n## Introduction)',
        lambda match: match.group(1) + revised[0].strip(), manuscript, count=1)
    manuscript, scope_count = re.subn(r'(?s)The current public reference, transitive software environment,.*?(?=\n\n## Data and code availability)',
        lambda match: revised[1].strip(), manuscript, count=1)
    manuscript, availability_count = re.subn(r'(?s)(## Data and code availability\n\n).*?(?=\n\n## Reproducibility appendix)',
        lambda match: match.group(1) + revised[2].strip(), manuscript, count=1)
    if (abstract_count, scope_count, availability_count) != (1, 1, 1):
        raise ValueError('P3 source section boundaries did not match exactly')
    output = ROOT / 'paper/manuscript_v5.md'
    output.write_text(manuscript + '\n', encoding='utf-8')
    record = {'state': 'Edge_text_assembled_pending_final_PDF_QA',
              'P1': {'path': str(original), 'sha256': sha(original)},
              'P2': {'path': str(ai), 'sha256': sha(ai)},
              'P3': {'path': str(revision), 'sha256': sha(revision)},
              'source_prose_generated_by_Codex': False,
              'format_and_factual_fixes': applied,
              'output': str(output), 'output_sha256': sha(output)}
    (ROOT / 'paper/manuscript_v5_assembly.json').write_text(json.dumps(record, indent=2) + '\n', encoding='utf-8')
    print(json.dumps({'state': record['state'], 'output': str(output), 'characters': len(manuscript)}))


if __name__ == '__main__':
    main()
````
