## Supplementary reproducibility code and frozen predictions for "Centered cross-context perturbation transfer on the public 2025 Virtual Cell Challenge benchmark": build_preprint_pdf_v5.py

````
"""Typeset the Edge-authored Markdown preprint with ReportLab.

Preparation modes --plan and --collect-math create no PDF. Actual PDF creation
must be invoked only after the coordinator runs the PDF artifact marker.
No manuscript prose, authors, captions or result values are invented here.
"""
from __future__ import annotations
import argparse
from collections import Counter
import hashlib
import html
import json
from pathlib import Path
import re

ROOT = Path(__file__).resolve().parents[1]
DEFAULT_INPUT = ROOT / 'paper/manuscript_v5.md'
DEFAULT_OUTPUT = ROOT / 'output/pdf/vcc2025_centered_transfer_preprint.pdf'
FIGURES = {1: 'final_v5_comparison_20261021.png', 2: 'final_v5_center_ablation_20261021.png'}
INLINE_MATH = re.compile(r'(?<!\\)\$(?!\$)(.+?)(?<!\\)\$', re.S)
MARKER = re.compile(r'^\[INSERT\s+FIGURE\s*([12])\]$', re.I)
CAPTION = re.compile(r'^(?:\*\*)?(?:Figure|Fig\.?)\s*([12])(?:[.:\s]|\*\*)', re.I)


def sha(path):
    return hashlib.sha256(Path(path).read_bytes()).hexdigest()


def display_tex(value):
    value = value.strip()
    tag = re.search(r'\\tag\{([^{}]+)\}\s*$', value)
    if tag:
        return value[:tag.start()].strip(), tag.group(1)
    return value, None


def math_record(latex, kind):
    size = 11 if kind == 'inline' else 12
    key = hashlib.sha256((kind + '\0' + str(size) + '\0' + latex).encode()).hexdigest()[:20]
    return {'id': key, 'latex': latex, 'kind': kind, 'font_size_pt': size}


def table_cells(line):
    """Split Markdown pipes while preserving escaped pipes and inline math."""
    line = line.strip()
    if line.startswith('|'):
        line = line[1:]
    if line.endswith('|') and not line.endswith('\\|'):
        line = line[:-1]
    cells, current, math_mode, escaped = [], [], False, False
    for char in line:
        if char == '$' and not escaped:
            math_mode = not math_mode
        if char == '|' and not math_mode and not escaped:
            cells.append(''.join(current).strip()); current = []
        else:
            current.append(char)
        escaped = char == '\\' and not escaped
    cells.append(''.join(current).strip())
    return [v.replace('\\|', '|') for v in cells]


def is_table_separator(line):
    return bool(line.strip()) and all(re.fullmatch(r':?-{3,}:?', v) for v in table_cells(line))


def blocks_from_markdown(source):
    lines = source.replace('\r\n', '\n').splitlines()
    if lines and lines[0].strip() in ('```markdown', '```md') and lines[-1].strip() == '```':
        lines = lines[1:-1]
    blocks, i = [], 0
    while i < len(lines):
        line = lines[i].strip()
        if not line or re.fullmatch(r'[-*_]{3,}', line):
            i += 1; continue
        heading = re.match(r'^(#{1,6})\s+(.+?)\s*#*$', line)
        if heading:
            blocks.append({'kind': 'heading', 'level': len(heading.group(1)), 'text': heading.group(2)})
            i += 1; continue
        marker = MARKER.fullmatch(line)
        if marker:
            blocks.append({'kind': 'figure', 'number': int(marker.group(1))}); i += 1; continue
        if line.startswith('$$'):
            if len(line) > 4 and line.endswith('$$'):
                equation = line[2:-2].strip(); i += 1
            else:
                chunks = [line[2:].strip()] if line[2:].strip() else []
                i += 1
                while i < len(lines) and not lines[i].strip().endswith('$$'):
                    chunks.append(lines[i].strip()); i += 1
                if i == len(lines):
                    raise ValueError('Unclosed display equation')
                tail = lines[i].strip()[:-2].strip()
                if tail:
                    chunks.append(tail)
                equation = ' '.join(chunks); i += 1
            equation, tag = display_tex(equation)
            blocks.append({'kind': 'math', 'latex': equation, 'tag': tag}); continue
        if line.startswith('```'):
            language = line[3:].strip(); chunks = []; i += 1
            while i < len(lines) and not lines[i].strip().startswith('```'):
                chunks.append(lines[i]); i += 1
            if i == len(lines):
                raise ValueError('Unclosed fenced block')
            i += 1
            if language in ('math', 'latex'):
                equation, tag = display_tex(' '.join(chunks))
                blocks.append({'kind': 'math', 'latex': equation, 'tag': tag})
            else:
                blocks.append({'kind': 'code', 'text': '\n'.join(chunks)})
            continue
        if i + 1 < len(lines) and '|' in line and is_table_separator(lines[i+1]):
            rows = [table_cells(line)]; i += 2
            while i < len(lines) and lines[i].strip() and '|' in lines[i]:
                rows.append(table_cells(lines[i])); i += 1
            if any(len(row) != len(rows[0]) for row in rows):
                raise ValueError('Markdown table columns are inconsistent')
            blocks.append({'kind': 'table', 'rows': rows}); continue
        bullet = re.match(r'^(\s*)([-+*]|\d+[.)])\s+(.+)$', lines[i])
        if bullet:
            text = bullet.group(3); symbol = bullet.group(2)
            indent = len(bullet.group(1)); i += 1
            while i < len(lines) and lines[i].strip() and len(lines[i])-len(lines[i].lstrip()) > indent and not re.match(r'^\s*[-+*]\s+', lines[i]):
                text += ' ' + lines[i].strip(); i += 1
            blocks.append({'kind':'list','text':text,'bullet':symbol if symbol[0].isdigit() else '\u2022','indent':indent}); continue
        def paragraph_line(value):
            hardbreak = value.endswith('  ') or value.endswith('\\')
            value = value.strip()
            if value.endswith('\\'):
                value = value[:-1]
            return value + ('<br/>' if hardbreak else '')
        chunks = [paragraph_line(lines[i])]; i += 1
        while i < len(lines) and lines[i].strip():
            nxt = lines[i].strip()
            if (re.match(r'^(#{1,6})\s|^```|^\$\$|^\[INSERT\s+FIGURE|^\s*[-+*]\s+|^\d+[.)]\s+', nxt)
                    or (i+1 < len(lines) and '|' in nxt and is_table_separator(lines[i+1]))):
                break
            chunks.append(paragraph_line(lines[i])); i += 1
        blocks.append({'kind':'paragraph','text':' '.join(chunks)})
    return blocks


def collect_math(blocks):
    records = {}
    for block in blocks:
        if block['kind'] == 'math':
            item = math_record(block['latex'], 'display'); records[item['id']] = item
        text_values = [block.get('text','')]
        if block['kind'] == 'table':
            text_values.extend(cell for row in block['rows'] for cell in row)
        for text in text_values:
            for match in INLINE_MATH.finditer(text):
                item = math_record(match.group(1).strip(), 'inline'); records[item['id']] = item
    return list(records.values())


def normalize_text(value):
    return value.translate(str.maketrans({'\u00a0':' ','\u2010':'-','\u2011':'-','\u2012':'-','\u2013':'-','\u2014':'-','\u2015':'-'}))


def register_fonts():
    from reportlab.pdfbase import pdfmetrics
    from reportlab.pdfbase.ttfonts import TTFont
    font_dir = Path('C:/Windows/Fonts')
    names = {'TimesNewRoman':'times.ttf','TimesNewRoman-Bold':'timesbd.ttf',
             'TimesNewRoman-Italic':'timesi.ttf','TimesNewRoman-BoldItalic':'timesbi.ttf'}
    for name, filename in names.items():
        if not (font_dir/filename).is_file():
            raise FileNotFoundError('Required manuscript font: '+str(font_dir/filename))
        pdfmetrics.registerFont(TTFont(name,str(font_dir/filename)))
    pdfmetrics.registerFontFamily('TimesNewRoman',normal='TimesNewRoman',bold='TimesNewRoman-Bold',
                                 italic='TimesNewRoman-Italic',boldItalic='TimesNewRoman-BoldItalic')
    chinese = font_dir/'simsun.ttc'
    if chinese.exists():
        pdfmetrics.registerFont(TTFont('CJKText',str(chinese),subfontIndex=0))
    return {name:str(font_dir/filename) for name,filename in names.items()}


class InlineFormatter:
    token = re.compile(r'(?<!\\)\$(?!\$)(.+?)(?<!\\)\$|`([^`]+)`|\*\*(.+?)\*\*|(?<!\*)\*([^*]+)\*|\[([^\]]+)\]\(([^\s]+)\)|(https?://[^\s<>]+)', re.S)

    def __init__(self, manifest_path, expected_math):
        self.manifest_path = manifest_path
        self.manifest = json.loads(manifest_path.read_text(encoding='utf-8')) if expected_math else {'equations':{}}
        self.equations = self.manifest['equations']
        if isinstance(self.equations,list):
            self.equations = {v['id']:v for v in self.equations}
        for item in expected_math:
            self.asset(item['latex'],item['kind'])

    def asset(self, latex, kind):
        key = math_record(latex,kind)['id']
        if key not in self.equations:
            raise ValueError('Missing rendered equation '+key+': '+latex)
        row = self.equations[key]
        if row.get('latex') != latex or row.get('kind') != kind:
            raise ValueError('Math asset source mismatch '+key)
        path = self.manifest_path.parent/row['png_file']
        if not path.is_file() or row['width_pt'] <= 0 or row['height_pt'] <= 0:
            raise ValueError('Invalid rendered equation '+key)
        return path,row

    def plain(self, value):
        value = normalize_text(value).replace('\\$','$')
        # Explicit CJK fallback keeps author/affiliation glyphs visible.
        pieces = re.split(r'([\u3400-\u9fff]+)',value)
        escaped = ''.join(('<font name="CJKText">'+html.escape(v)+'</font>') if re.fullmatch(r'[\u3400-\u9fff]+',v or '') else html.escape(v,quote=False) for v in pieces)
        for tag, replacement in [('sup','super'),('sub','sub')]:
            escaped=escaped.replace('&lt;'+tag+'&gt;','<'+replacement+'>').replace('&lt;/'+tag+'&gt;','</'+replacement+'>')
        return re.sub(r'&lt;br\s*/?&gt;','<br/>',escaped,flags=re.I)

    def __call__(self, value):
        output, last = [], 0
        for match in self.token.finditer(value):
            output.append(self.plain(value[last:match.start()]))
            if match.group(1) is not None:
                path,row = self.asset(match.group(1).strip(),'inline')
                output.append(f'<img src="{html.escape(str(path),quote=True)}" width="{row["width_pt"]:.4f}" height="{row["height_pt"]:.4f}" valign="{-row.get("depth_pt",0.):.4f}"/>')
            elif match.group(2) is not None:
                output.append('<font name="Courier" size="9">'+self.plain(match.group(2))+'</font>')
            elif match.group(3) is not None:
                output.append('<b>'+self(match.group(3))+'</b>')
            elif match.group(4) is not None:
                output.append('<i>'+self(match.group(4))+'</i>')
            elif match.group(5) is not None:
                output.append('<link href="'+html.escape(match.group(6),quote=True)+'" color="#173a63">'+self(match.group(5))+'</link>')
            else:
                url = match.group(7).rstrip('.,;')
                suffix = match.group(7)[len(url):]
                output.append('<link href="'+html.escape(url,quote=True)+'" color="#173a63">'+self.plain(url)+'</link>'+self.plain(suffix))
            last=match.end()
        output.append(self.plain(value[last:]))
        return ''.join(output)


def build_pdf(args, blocks, equations):
    from PIL import Image as PILImage
    from reportlab.lib import colors
    from reportlab.lib.enums import TA_CENTER, TA_JUSTIFY, TA_LEFT
    from reportlab.lib.pagesizes import A4
    from reportlab.lib.styles import ParagraphStyle
    from reportlab.platypus import SimpleDocTemplate,Paragraph,Spacer,Image,LongTable,Table,TableStyle,KeepTogether,Preformatted
    from pypdf import PdfReader
    fonts = register_fonts()
    formatter = InlineFormatter(args.math_manifest,equations)
    style_args = dict(fontName='TimesNewRoman',fontSize=11,leading=14.4,spaceAfter=7,
                      splitLongWords=True,allowWidows=0,allowOrphans=0,autoLeading='max')
    styles = {'body':ParagraphStyle('body',alignment=TA_JUSTIFY,**style_args)}
    styles['title']=ParagraphStyle('title',parent=styles['body'],fontName='TimesNewRoman-Bold',fontSize=18,leading=21.5,alignment=TA_CENTER,spaceAfter=14,keepWithNext=True)
    styles['front']=ParagraphStyle('front',parent=styles['body'],fontSize=10,leading=12.8,alignment=TA_CENTER,spaceAfter=6)
    styles['h2']=ParagraphStyle('h2',parent=styles['body'],fontName='TimesNewRoman-Bold',fontSize=13,leading=16,spaceBefore=11,spaceAfter=6,keepWithNext=True,alignment=TA_LEFT)
    styles['h3']=ParagraphStyle('h3',parent=styles['h2'],fontSize=11.5,leading=14,spaceBefore=8)
    styles['caption']=ParagraphStyle('caption',parent=styles['body'],fontSize=9.5,leading=12,alignment=TA_LEFT,spaceAfter=10)
    styles['tablecell']=ParagraphStyle('tablecell',parent=styles['body'],fontSize=9,leading=11.5,alignment=TA_LEFT,spaceAfter=0)
    styles['tableheader']=ParagraphStyle('tableheader',parent=styles['tablecell'],fontName='TimesNewRoman-Bold')
    styles['reference']=ParagraphStyle('reference',parent=styles['body'],fontSize=9.5,leading=12,alignment=TA_LEFT,spaceAfter=5)
    styles['list']=ParagraphStyle('list',parent=styles['body'],leftIndent=13,firstLineIndent=0,bulletIndent=1,spaceAfter=4)
    styles['code']=ParagraphStyle('code',fontName='Courier',fontSize=8,leading=10,spaceBefore=5,spaceAfter=8)
    width,height=A4; margin=55
    body_width=width-2*margin; usable_height=height-2*margin
    captions={}
    for index,block in enumerate(blocks):
        if block['kind']=='paragraph' and (match:=CAPTION.match(block['text'])):
            number=int(match.group(1))
            if number in captions:
                raise ValueError('Duplicate figure caption '+str(number))
            captions[number]=(index,block['text'])
    required={b['number'] for b in blocks if b['kind']=='figure'}
    if required != {1,2}:
        raise ValueError('Both Figure1 and Figure2 insertion markers are required')
    if not required<=captions.keys():
        raise ValueError('Source caption missing for figure '+str(required-captions.keys()))
    used_caption_rows={v[0] for v in captions.values()}
    story=[]; front=True; in_references=False; title=''; figure_records=[]
    for index,block in enumerate(blocks):
        kind=block['kind']
        if index in used_caption_rows:
            continue
        if kind=='heading':
            text=block['text']
            if block['level']==1 and not title:
                title=re.sub(r'[*`$]','',text); style=styles['title']
            else:
                front=False; in_references=text.lower().strip() in ('references','bibliography')
                if text.lower().strip() in ('figure legends','figure captions'):
                    continue
                style=styles['h2'] if block['level']<=2 else styles['h3']
            story.append(Paragraph(formatter(text),style))
        elif kind=='paragraph':
            if re.search(r'\[(?:INSERT|TODO|TBD|PLACEHOLDER)\b',block['text'],re.I):
                raise ValueError('Unresolved placeholder in manuscript')
            style=styles['front'] if front else styles['reference'] if in_references else styles['body']
            if re.match(r'^(?:\*\*)?Table\s+\d',block['text'],re.I):
                style=ParagraphStyle('tablecaption',parent=styles['caption'],keepWithNext=True,spaceAfter=5)
            story.append(Paragraph(formatter(block['text']),style))
        elif kind=='list':
            style=styles['reference'] if in_references else styles['list']
            story.append(Paragraph(formatter(block['text']),style,bulletText=block['bullet']))
        elif kind=='math':
            path,row=formatter.asset(block['latex'],'display')
            scale=min(1.,(body_width-38)/row['width_pt'])
            image=Image(str(path),width=row['width_pt']*scale,height=row['height_pt']*scale)
            image.hAlign='CENTER'
            if block.get('tag'):
                display=Table([[image,Paragraph('('+formatter.plain(block['tag'])+')',styles['body'])]],colWidths=[body_width-38,38])
                display.setStyle(TableStyle([('VALIGN',(0,0),(-1,-1),'MIDDLE'),('ALIGN',(1,0),(1,0),'RIGHT')]))
            else:
                display=image
            story.extend([Spacer(1,5),display,Spacer(1,8)])
        elif kind=='figure':
            number=block['number']; path=args.figure_dir/FIGURES[number]
            if not path.is_file():
                raise FileNotFoundError('Required final figure: '+str(path))
            with PILImage.open(path) as picture:
                px_w,px_h=picture.size
                dpi=picture.info.get('dpi')
            scale=min(body_width/px_w,(usable_height*.67)/px_h)
            fig=Image(str(path),width=px_w*scale,height=px_h*scale); fig.hAlign='CENTER'
            caption=Paragraph(formatter(captions[number][1]),styles['caption'])
            story.append(KeepTogether([Spacer(1,8),fig,Spacer(1,6),caption]))
            figure_records.append({'number':number,'path':str(path),'sha256':sha(path),'pixels':[px_w,px_h],'stored_dpi':dpi,'display_pt':[px_w*scale,px_h*scale]})
        elif kind=='table':
            rows=block['rows']; n=len(rows[0])
            weights=[min(32,max(8,max(len(re.sub(r'\$.*?\$','equation',r[c])) for r in rows))) for c in range(n)]
            widths=[body_width*w/sum(weights) for w in weights]
            cells=[[Paragraph(formatter(value),styles['tableheader'] if i==0 else styles['tablecell']) for value in row] for i,row in enumerate(rows)]
            table=LongTable(cells,colWidths=widths,repeatRows=1,splitByRow=1,splitInRow=1,hAlign='LEFT',spaceBefore=5,spaceAfter=10)
            table.setStyle(TableStyle([('VALIGN',(0,0),(-1,-1),'TOP'),('BACKGROUND',(0,0),(-1,0),colors.HexColor('#edf0f3')),
                ('LINEABOVE',(0,0),(-1,0),.6,colors.black),('LINEBELOW',(0,0),(-1,0),.4,colors.black),
                ('LINEBELOW',(0,-1),(-1,-1),.5,colors.black),('LEFTPADDING',(0,0),(-1,-1),5),
                ('RIGHTPADDING',(0,0),(-1,-1),5),('TOPPADDING',(0,0),(-1,-1),5),('BOTTOMPADDING',(0,0),(-1,-1),5)]))
            story.append(table)
        elif kind=='code':
            story.append(Preformatted(normalize_text(block['text']),styles['code'],maxLineLength=92,splitChars=' /+-='))
    args.output.parent.mkdir(parents=True,exist_ok=True)
    temporary=args.output.with_name(args.output.stem+'.working.pdf')

    def page_decoration(canvas,doc):
        canvas.saveState(); canvas.setFont('TimesNewRoman',8); canvas.setFillColor(colors.HexColor('#555555'))
        if doc.page>1:
            header=title if len(title)<=90 else title[:87]+'...'
            canvas.drawString(margin,height-35,header)
        canvas.drawString(margin,31,'Preprint'); canvas.drawRightString(width-margin,31,str(doc.page)); canvas.restoreState()

    document=SimpleDocTemplate(str(temporary),pagesize=A4,leftMargin=margin,rightMargin=margin,
        topMargin=margin,bottomMargin=margin,title=title,author=args.pdf_author or '',pageCompression=1)
    document.build(story,onFirstPage=page_decoration,onLaterPages=page_decoration)
    reader=PdfReader(str(temporary))
    if not reader.pages:
        raise ValueError('Empty PDF')
    extracted='\n'.join(page.extract_text() or '' for page in reader.pages)
    if re.search(r'\[INSERT\s+FIGURE|\\(?:frac|mathrm|operatorname|alpha|beta)\b|\$\$',extracted):
        raise ValueError('Unrendered equation or insertion marker found in output text')
    temporary.replace(args.output)
    record={'state':'PDF_created_visual_QA_pending','source_markdown':str(args.input),'source_sha256':sha(args.input),
        'output':str(args.output),'output_sha256':sha(args.output),'pages':len(reader.pages),'paper':'A4','body_font_pt':11,
        'body_leading_pt':14.4,'fonts':fonts,'figures':figure_records,'math_assets':len(equations),
        'math_manifest':str(args.math_manifest) if equations else None,'math_manifest_sha256':sha(args.math_manifest) if equations else None,
        'block_counts':dict(Counter(v['kind'] for v in blocks)),'source_prose_generated':False,'visual_review_required':True}
    args.output.with_suffix('.build.json').write_text(json.dumps(record,indent=2)+'\n',encoding='utf-8')
    return record


def main():
    parser=argparse.ArgumentParser(description=__doc__)
    parser.add_argument('--input',type=Path,default=DEFAULT_INPUT)
    parser.add_argument('--output',type=Path,default=DEFAULT_OUTPUT)
    parser.add_argument('--math-manifest',type=Path,default=ROOT/'paper/math_assets_v5/manifest.json')
    parser.add_argument('--figure-dir',type=Path,default=ROOT/'paper/figures')
    parser.add_argument('--pdf-author',default='',help='Optional PDF metadata only; visible authors/affiliations come from source Markdown')
    parser.add_argument('--collect-math',type=Path,help='Write equation render job JSON only; no PDF authoring')
    parser.add_argument('--plan',action='store_true',help='Inspect source blocks/equation requirements only; no PDF authoring')
    args=parser.parse_args()
    blocks=blocks_from_markdown(args.input.read_text(encoding='utf-8-sig'))
    equations=collect_math(blocks)
    if args.collect_math:
        args.collect_math.parent.mkdir(parents=True,exist_ok=True)
        job={'schema':'preprint_math_job_v5','source_markdown':str(args.input),'source_sha256':sha(args.input),'dpi':300,'equations':equations}
        args.collect_math.write_text(json.dumps(job,indent=2)+'\n',encoding='utf-8')
        print(json.dumps({'state':'math_job_collected_no_PDF','job':str(args.collect_math),'equations':len(equations)})); return
    if args.plan:
        print(json.dumps({'state':'plan_only_no_PDF','source':str(args.input),'source_sha256':sha(args.input),
                          'blocks':dict(Counter(b['kind'] for b in blocks)),'math_assets_needed':len(equations)},indent=2)); return
    print(json.dumps(build_pdf(args,blocks,equations),indent=2))


if __name__=='__main__':main()
````
