## Supplementary reproducibility code and frozen predictions for "Centered cross-context perturbation transfer on the public 2025 Virtual Cell Challenge benchmark": preprint_prompt_20261021_P1.md

# Manuscript task P1, frozen numerical V5, 21 October 2026

Write a complete English research preprint from the verified facts below. This is the final centered R1 model, superseding earlier M1/v1 writing in this conversation. The model contains no prediction agent. The authors want a clear, useful paper about a compact numerical method and its measured benchmark result.

Deliver the entire manuscript as a single fenced `markdown` code block so it can be copied exactly. Include title, author block, abstract, Introduction, Results, Methods, Discussion, Data and code availability, two figure captions, tables and References. Aim for 2,500-3,500 words excluding references. Use a concise, accurate title about centered cross-context perturbation transfer on the public 2025 Virtual Cell Challenge benchmark. Do not put an unqualified competition-victory claim in the title.

Use short, natural academic sentences. Do not join clauses with em dashes, en dashes or double hyphens. Use ASCII hyphens for compound words. Minimize clause-joining colons and semicolons. State measured evidence directly and confidently. Keep the conditions needed to understand the comparison beside the claims. Avoid defensive boilerplate, repeated caveats and paragraphs about the writing process.

The main numerical result is 0.34134444924634444 versus the archived champion score 0.3400522468873988, a difference of +0.0012922023589456155. The abstract must identify this as a retrospective evaluation on the released public TEST100 with fixed reconstructed historical anchors. The original champion checkpoint and private scoring reference were not rerun. Describe these facts once where relevant. This is not a new official leaderboard entry or proof of statistical significance. Current external source products include a 2026 KOLF release, so do not describe the model as eligible for the original 2025 deadline.

Report the final raw DES/PDS/MAE and explain the actual tradeoff. DES is higher than the archive, PDS is lower, and both methods have a zero normalized MAE contribution. Higher benchmark score alone does not establish simulation of cellular heterogeneity or a biological mechanism.

Methods must be sufficiently explicit to implement the frozen model. Define TRAIN control means, the numerical initializer, source centering on known fit targets, the strong-source threshold, quantile transfer, the independent DE kernel, isotonic magnitude rendering, own-gene handling and public-count mean-vector replication. State the exact split design, repeated official VALIDATION development, four sequential frozen TEST evaluations and the separation of model developers from TEST expression/evaluation reports. Do not claim a single untouched blind test. Distinguish the TRAIN top-N proxy from native DES. Use the actual preprocessing and metric definitions in the evidence.

Keep hashes, paths, version inventory and implementation checks compact. They belong in a short reproducibility appendix or data/code section rather than filling the Results narrative. Do not invent a public code repository, deposited weights, unperformed tests, biological validation, error bars, confidence intervals, funding, author contributions or competing-interest declarations. Input source files and exact code are identified in the accompanying evidence archive.

Use standard Markdown headings and tables. Use standard LaTeX `$...$` inline math and one short equation per `$$...$$` display block. Avoid aligned/cases/array environments and long multi-line equations, for reliable downstream typesetting. Figures will be inserted by the local typesetter. Put a marker `[INSERT FIGURE 1]` and `[INSERT FIGURE 2]` where each belongs and write complete scientific captions. Figure 1 compares archive-reported champion and locally computed V5 raw DES/PDS/MAE plus the normalized components and overall. Figure 2 plots development PDS over the source-centering coefficient, with the selected coefficient 1.25 marked, and can show actual fold results if supplied. Do not describe plots that are not supplied.

Use numbered references with direct DOI or primary-source URLs. The supplied reference list is verified; do not invent extra bibliographic details. You may cite a software or dataset page directly. Paraphrase rather than quote. Keep descriptions of each external source brief and focus most text on this study's new methods and measurements.

Authors in this exact order:
Haoyang Luan (2), Letian Chen (1,2), Gufeng Yu (2), Yang Yang (2, corresponding).
1. Shanghai Innovation Institute, Shanghai, China.
2. AGI Institute, School of Computer Science, Shanghai Jiao Tong University, Shanghai 200240, China.
Other author.

Return manuscript prose directly, with no introductory chat or checking checklist. End outside the code block with the exact marker VCC-PREPRINT-V5-P1-END.

## Verified primary references

1. Arc Institute. Virtual Cell Challenge dataset documentation. https://github.com/ArcInstitute/arc-virtual-cell-atlas/blob/main/virtual-cell-challenge/README.md . H1 hESC CRISPRi benchmark, released TRAIN/VALIDATION/TEST.
2. Arc Institute. Virtual Cell Challenge 2025 Wrap-Up: Winners and Reflections. 6 December 2025. https://arcinstitute.org/news/virtual-cell-challenge-2025-wrap-up . Winner BM_xTVC, xTrimoSCPerturb. Archived numeric results at https://virtualcellchallenge.org/2025 .
3. Replogle JM et al. Mapping information-rich genotype-phenotype landscapes with genome-scale Perturb-seq. Cell (2022). https://doi.org/10.1016/j.cell.2022.05.013 . Author processed-data deposit https://plus.figshare.com/articles/dataset/20029387 .
4. Nourreddine S, Doctor Y, Dailamy A et al. A genome-scale CRISPRi perturbation atlas of human induced pluripotent stem cells. Nature Biotechnology (2026), published 1 July 2026. https://doi.org/10.1038/s41587-026-03199-w . Publisher DOI metadata independently checked through Crossref https://crossmark.crossref.org/dialog/?doi=10.1038%2Fs41587-026-03199-w .
5. Nourreddine, Doctor et al. KOLF2.1J Perturb-Seq Atlas author viewer and code. https://y-doctor.github.io/KOLF2.1J_Perturbation_Cell_Atlas/ ; fixed source https://github.com/y-doctor/KOLF2.1J_Perturbation_Cell_Atlas/tree/8f84531a784f24b14d7e2c9b80c375d7e78fd46c . Preprint predecessor https://doi.org/10.1101/2024.11.03.621734 . Describe the derived viewer panel separately from the journal experiment, not as identical processing.
6. Arc Institute. PIE X-atlas Orion derived public dataset. https://huggingface.co/datasets/arcinstitute/PIE_x_atlas_orion . Fixed revision ecb4491ea49fcf28df463fac1fc9ef00c0c3ac17. HCT116 and HEK293T delta profiles. Underlying study Huang AC et al., X-Atlas/Orion: Genome-wide Perturb-seq datasets via a scalable fix-cryopreserve platform for training dose-dependent biological foundation models, bioRxiv (2025), https://doi.org/10.1101/2025.06.11.659105 .
7. Liu Q, Zhang Q, Du J, Zhao S, Wang J. Effects of Distance Metrics and Scaling on the Perturbation Discrimination Score. arXiv (2025). https://doi.org/10.48550/arXiv.2511.16954 .
8. Arc Institute. cell-eval, v0.6.6. https://github.com/ArcInstitute/cell-eval/tree/v0.6.6 ; aggregate scoring https://github.com/ArcInstitute/cell-eval/blob/v0.6.6/src/cell_eval/_score.py .
9. Arc Institute. pdex, v0.1.27. https://github.com/ArcInstitute/pdex/blob/pdex-0.1.27/src/pdex/_single_cell.py .

## Verified technical evidence


# V5 technical facts for Edge — 2026-10-21

Purpose: evidence input only. Write the manuscript through the authorized Edge academic-writing workflow. Do not upgrade the comparison into a 2025 prize/private-leaderboard win. All numbers below are measured aggregates; full provenance is final_results_v5.json.

## Result and evaluation scope

- Final frozen public TEST100: DES=0.37183447754891852; PDS=0.86140000000000017; MAE=0.30220562092959879; fixed-anchor average=0.34134444924634444.
- Archive-reported BM_xTVC / xTVCM_v01 / xTrimoSCPerturb score=0.34005224688739882; local difference=+0.0012922023589456155. Archived raw DES=.34882155189465897, PDS=.872, MAE=1.0258748292922975. Champion checkpoint/predictions were not rerun.
- Primary scope: retrospective reanalysis of the released public 2025 benchmark, using a 2026 external source; no official private leaderboard/prize equivalence established.
- Native VALIDATION50: DES=0.39449605216692274, PDS=0.86919999999999997, MAE=0.28466863233537937, fixed-anchor development score=0.35497842748162273.
- TRAIN150 target-level 3fold (each fit100/query50): PDS=0.8632, MAE=0.38722279893791373, DE-topN proxy=0.38763587167635533. V5 native TRAIN DES not computed. Never call topN native DES.
- Selection: (3*TRAIN3fold topN-based joint+VALIDATION topN-based joint)/4=0.35009489530924903; later native VAL confirmation is separate. Official VAL was repeatedly used for development.
- TEST history (all same frozen anchors; four sequential freezes, not a single untouched blind test):
- v1: DES=0.370076922852; PDS=0.8149; MAE=0.194522297233; score=0.309180186089.
- v2: DES=0.370815295399; PDS=0.8366; MAE=0.233661419787; score=0.32416003771.
- v3: DES=0.371847148058; PDS=0.8401; MAE=0.31443569418; score=0.326912353452.
- v4 gamma1: prediction generated, never TEST-scored; superseded by gamma1.25 before centered-model TEST evaluation.
- v5: DES=0.371834477549; PDS=0.8614; MAE=0.30220562093; score=0.341344449246.
- Coordinator/evaluator saw earlier TEST aggregate outcomes. Model-development agents had no TEST expression/evaluation reports; no TEST per-target analysis was used. Disclose this retrospective sequential evaluation design.

## Model formulas transcribed from frozen source

- Numerical only; no internal prediction agent. Deterministic inference; target folds frozen; no sampling seed needed.
- Legacy multi-arm archive contributes B/targets/genes only; v5 reads original numerical expression_direction_source/de_score features, not prior agent arms. Public-center means use each F100 fit subset in CV; full TRAIN150 for final inference.
- c=TRAIN NTC log1p mean; s=control SD floor; C clips [0,nextafter(F32(15),0)]. B=frozen legacy numeric B array.
- Bcenter=C(c+32*(B-mean_query(B))).astype(F32).
- G=C(c+128*s*mean_available_three_GEMZ_source_z); missing/own coordinates fall back to B; G stored F32.
- O=C(.25*Bcenter+.75*G).astype(F32).
- X=C(c+128*mean_available_HCT116_HEK293T_delta); missing/own coordinates fall back to O.
- M=C(.25*O+.75*X).astype(F32); v5 consumes this expression feature, not the original power-curve output.
- old_DE_KRR: Concatenate three masked per-source L2-normalized response blocks; Gram/mean_fit_diagonal=K; S_old=clip(f+.5*K_qf*(K_ff+I)^(-1)*(D-f),0,1), F32; f=mean_fit(D).
- KOLF_public_center: mu_g=sum_observed_TRAIN150(z_tg)/n_observed; 0 if n=0. z_centered=z_query-1.25*mu. Source availability mask is unchanged; own input set0 for kernels; TAZ unresolved.
- KOLF_DE_KRR: Uncentered aligned KOLF vectors, masked0 and own input0, per-target L2; use source-known fit rows only. K trace-scaled. S_K=clip(f_K+.5*K_qf*(K_ff+I)^(-1)*(D_known-f_K),0,1), cast F32 then F64. S=.75*S_old+.25*S_K; missing-source queries fall back to old DE before mixing.
- alpha1_quantile_expression: Eligible=available & c>0 & M>0, excluding own coordinate. f0=sign(M-c)*abs(log2(expm1(M)/expm1(c))). r=average_rank(abs(z_centered))/n_eligible. Map to sign(z_centered)*linear_quantile(abs(f0_eligible),r); decode with logaddexp(0,log(expm1(c))+ln(2)*f). Apply only when abs(z_centered)>=float32(.08); weak/missing coordinates retain exact M before isotonic rendering.
- isotonic_rendering: u=average_rank(S)/18080. Per target fit increasing IsotonicRegression(u, absFC(adapted_expression,c)) on c>0; predict all u with boundary clipping. a=.3*projection+.02*u+.015. Final mean=C(logaddexp(0,log(expm1(c))+ln(2)*sign(adapted_expression-c)*a)).astype(F32). Literal own gene set0 when TRAIN c>0, excluding unresolved TAZ.
- zero_FC_rules: For actual_abs_log2_fc, bothzero ratio1; refzero/targetpositive ratio20; targetzero/refpositive ratio.05; otherwise expm1(target)/expm1(reference), without generic ratio clipping.
- KOLF expression center is fitted from observed finite published TRAIN source coordinates only. Current final fit uses138/150 source-known TRAIN targets; source-known TEST91/100. Missing queries retain old DE fallback. Quantized zeros do not reveal the original missing/finite mask.
- VALIDATION replay: all904000 F32 mean entries identical to the selected development candidate (0 changed, max difference0). TEST means were frozen before that version evaluation; source arrays remain remote.

## Metrics and preprocessing

- 18080 genes, TRAIN150/VAL50/TEST100 perturbation targets. TEST170846 cells, including38176 controls; query132670 cells. Predictions: one F32 mean vector per target, repeated at fixed independent public target cell counts; full TRAIN NTC38176 cells retained.
- Counts -> Scanpy normalize_total(target_sum=None, positive-library median of each original full object) -> log1p F32. TRAIN prediction reference54611; real VAL54312; real TEST54125.5. Real and predicted controls are independent.
- Native TEST bulk: original-order Polars Float32 group_by.mean, independent gene blocks. Native L1 PDS excludes literal own gene; MAE is native pseudobulk MAE.
- Native DES: two-sided nonexperimental pdex MWU, tie correction, is_log1p/exp_post_agg; global BH family100*18080=1808000 (VAL904000), FDR<.05; rank abs(log2 FC), overlap at actual DEG count (k=None). Exact constant-target MWU transports unchanged arithmetic. Process/full VALIDATION raw parity904000 rows passed.
- cell-eval0.6.6; pdex0.1.27; NumPy2.4.6; SciPy1.17.1; Polars2.0.0; Scanpy1.11.5. Inference Python3.11.10/scikit-learn1.9.1.
- Fixed reconstructed archive anchors: DES=.09505113849605429, PDS=.5082, MAE=.0249575331993401. Mean of max(0,(DES-b_DES)/(1-b_DES)), max(0,(PDS-b_PDS)/(1-b_PDS)), max(0,1-MAE/b_MAE).
- Anchors were algebraically recovered from100 archive rows, not an original baseline CSV; MAE anchor had one independently active row. See research/historical_reference_anchor_20261015.md.
- V5 normalized components: {"DES":0.3058552265515585,"PDS":0.7181781211874749,"MAE":0.0}; MAE component0 (also0 for archived champion).
- V5 scoring total158.669s; MWU66.174s; peak parent RSS14684912KiB. No full H5AD submission pipeline/private-reference bridge. No confidence interval or champion checkpoint replay. Never infer biological fidelity from score alone.

## Data/version identity and citations

- Official datasets: https://github.com/ArcInstitute/arc-virtual-cell-atlas/blob/main/virtual-cell-challenge/README.md ; champion archive https://virtualcellchallenge.org/2025 ; authorship https://arcinstitute.org/news/virtual-cell-challenge-2025-wrap-up .
- GCS H5AD generations TRAIN1765904883947296 / VAL1765905029341737 / TEST1765905295402700. TEST11950739168bytes, CRC32C JMP5RQ==; full SHA256 below. Validation raw SHA not newly asserted; all generations/CRC and measured artifacts are in machine JSON.
- KOLF journal: Nourreddine, Doctor, Dailamy et al., Nature Biotechnology, published2026-07-01, https://doi.org/10.1038/s41587-026-03199-w . Current author viewer commit8f84531a784f24b14d7e2c9b80c375d7e78fd46c, https://github.com/y-doctor/KOLF2.1J_Perturbation_Cell_Atlas/tree/8f84531a784f24b14d7e2c9b80c375d7e78fd46c .
- KOLF viewer: signed int8*.04, mean NTC z from CPM+log1p exporter;10167 targets x38606 genes. Current all-pert panel first reachable commit2026-04-23; Figshare release2026-05-18. Current panel availability before2025 competition cutoff unestablished. 2024 preprint/2025 repository creation do not prove it. Figshare https://doi.org/10.25452/figshare.plus.27261219.v1 .
- KOLF viewer is not automatically the journal processing recipe; no original finite/missing mask. MIT repository license and CC-BY4.0 Figshare data license recorded; binary-specific license not independently resolved.
- X-atlas contexts HCT116/HEK293T, https://huggingface.co/datasets/arcinstitute/PIE_x_atlas_orion , frozen revision ecb4491ea49fcf28df463fac1fc9ef00c0c3ac17. GEM-Z public K562 GWPS/RPE1/K562essential source blocks.

## Key SHA256 records

- v5_freeze: f7c008f4f55d4deef3f05df870e5d7274ec18662ae44bf21752c0b4c4635a8c9
- v5_prediction: cea4874ec557b56622d66e5a414ebe8ecb59fee078e5f613a3316cbc9fffb4df
- v5_inference_code: ece7309e1f17c1cdbe3ff74e74ba124bc409076b0e6b2bbdf1d6a5099647118f
- v5_score_config: 136f3db1b2679ae5d7dafdcfce3bd1ee286f6bdfd48a4f470e5a0b5f92549c26
- v5_TEST_summary: b681574e7e6a484110597c5442cb8121abe58ae6cf344cfae00c7dbbb03f9362
- TEST_H5AD: ba5ce66729e4c8e50b6f147c7d73d3f9f28f76c91fa43ff33e3f6bc2ed831f03
- TEST_real_DE: d01903bce4818bf8dd9b396258ba4891e37f877919f09fc82fefda8b78c925ff
- fitted_public_center: f7b084cf6117baea7a9c6a853bdae77a79588ce9b1d69659deeb1a523296bae7

All local source/report SHA256s, frozen input-array hashes, versions, native components, previous TEST aggregates and explicit limitations are in final_results_v5.json. No full per-target data in this packet.


# B initializer technical definition — v5

Role: the `B` field of the frozen **uncalibrated** multi-arm archive, used only as a numerical input to `predict_numeric_ranktransfer_v1.py`. The other archive arms are not B. Current v5 prediction features inherit this B through the original M expression initializer.

## Inputs and preprocessing

- Three separate original GEM-Z contexts, in order: `K562_gwps_normalized_bulk_01`, `rpe1_normalized_bulk_01`, `K562_essential_normalized_bulk_01`; feature dimensions 8248/8749/8563. All-public scaler genes, centers and population SDs are frozen; genes need >=2 observations and variance>1e-24. Features are `(source_value-public_center)/public_SD` on observed entries, with missing numerical entries0 and an explicit row-availability flag.
- Source rank128 basis objects provide the stored scaler only. **No PCA projection** is applied. B uses `original_features`, retaining original own-source coordinates; `selfmasked_features` belong to the separate retrieval branch.
- Exact/unique audited public identity alignment, no new alias expansion; TAZ remains unresolved. Query encoding does not refit the public scaler. Fitted ridge feature centers below use only available H1 fit-target rows.
- TRAIN teacher is `data/processed_ntc_scale_v1/training_pseudobulk.npz`, SHA256 `939ed693fd30c555794036ba2df930edfade95ebc5da53bf2304a9196281ac33`. Counts use target54611 then log1p; teacher means are F32. Set `c=NTC_mean.astype(F64)` and `Y_i=teacher_mean_i-c` in F64. Final fit uses all150 TRAIN targets and18080 output genes. CV B uses its current F100 fit rows.

## Exact source solve and projection order

For context j, let I_j contain its available fit rows and n_j=|I_j|. Enable the context only if n_j>=3 and the kernel trace scale exceeds1e-24.

```text
xbar_j = mean(x_j[I_j]); X_j = x_j[I_j] - xbar_j
tau_j = trace(X_j @ X_j.T) / n_j
ybar_j = mean(Y[I_j])
A_j = solve(X_j @ X_j.T / tau_j + .001*n_j*I, Y[I_j]-ybar_j)
delta_j(q) = ybar_j + ((x_j(q)-xbar_j) @ X_j.T / tau_j) @ A_j
p_j(q) = maximum(c+delta_j(q), 0) - c
delta(q) = equal mean of p_j(q) over enabled, query-available contexts
           or mean_all_fit(Y) when no context is available
B(q) = clip(c+delta(q), 0, nextafter(float32(15),0))
```

Alpha=.001 multiplies each context's available fit-row count in the trace-normalized kernel. Shrinkage=1; arithmetic/storage is F64. Each expert receives its lower projection **before** the context average; the upper CAP is applied after averaging. Response-centering ybar_j is added back.

## Branch identity and source code

- `src/virtualcell/final_training150_base_catalogue_v1.py`: `fit_base_states`, `predict_base_delta`, `pack_base`. This serializes the exact original `stage2_fit_dataset.source_prediction` + `residual_transfer_diagnostic.gemz_baseline` arithmetic.
- `src/virtualcell/dual_source_ridge.py`: `SourceScaler.encode`; `src/virtualcell/final150_public_query_features_v1.py`: `encode_public_source`, `source_views_from_output`.
- `scripts/prepare_training150_residual_v3.py` prepares source features. `scripts/fit_final_training150_base_catalogue_v1.py` fits and serializes the full150 numerical base.
- `src/virtualcell/residual_agent_final150_v1.py`: `Final150Context` sets `baseline_means=clip(c+predict_base_delta(...),0,CAP)`; `consume_four_arms` sets `uncalibrated['B']=baseline_means.copy()` separately from corrected/calibrated arms.
- B has **no** donor OOF residual correction, rank8 SVD program, GenePT/protein target kernel, retrieval weights, trust bound, agent decision or DE calibration. The word residual in its ridge means only centered supervised responses `Y-ybar`; it is not the later agent-residual branch.

Fitted B state: `/data1/luanhaoyang/virtualcell/runs/final_training150_base_catalogue_v1/final_base_full150_state.npz`, SHA256 `448f3d42b423398aab5d418ff673a1bbf8962d032f14a6da579fe73c468897fd`. Frozen source manifest SHA256 `5f72535fd14fe9750743ba45d95641b9bc6dd4023d116cbfc9ddf32d8bf7bc48`. Full prepared feature paths/hashes, source-code hashes and original configs are in `B_initializer_v5.json`. NumPy2.4.6 was pinned for the base fit. Arrays remain remote.

Documentation verification: existing code and small frozen metadata only; no new fit, prediction, experiment or array read.


# Supplementary measured facts for preprint P1

- Figure 1 assets: paper/figures/final_v5_comparison_20261021.png and .svg. Four panels. Raw DES, PDS and MAE, followed by fixed-anchor normalized DES/PDS/MAE and overall. Grey hatched archive-reported champion, blue locally computed frozen V5. No invented uncertainty estimates.
- Figure 2 assets are being produced from the seven measured center coefficients 0, .25, .5, 1, 1.25, 1.5, 2. Plot development centroid PDS for mean TRAIN three-fold queries and official VALIDATION; mark selected 1.25. Second panel uses three actual fold pairs.
- R1 gamma0 to selected gamma1.25 TRAIN mean PDS: .8438666666666667 to .8632; VALIDATION: .848 to .8692.
- Actual TRAIN fold PDS pairs: F0 .8636 to .8728; F1 .8304 to .8564; F2 .8376 to .8604. These are development-fold values, not confidence intervals.
- Native VALIDATION gamma0: DES .3947980234987502, PDS .848, MAE .2923363712016672, fixed-anchor score .34072067241135257.
- Native VALIDATION gamma1: DES .3946032706577774, PDS .8672, MAE .28613066620454874, fixed-anchor score .35366235627746057.
- Native VALIDATION gamma1.25: DES .39449605216692274, PDS .8692, MAE .28466863233537937, fixed-anchor score .3549784274816227.
- At center coefficient 2, development TRAIN PDS .8530666666666665 and VALIDATION .844. The effect is not monotone in the coefficient.
- Raising the original uncentered source threshold alone (.12/.16/.24) reduced the pooled development objective, rather than reproducing the centered-model gain. Mean centering changes magnitude ranks and threshold-based source selection together; this ablation does not isolate a biological mechanism.
- Final V5 independent inference job, including exact VALIDATION replay and TEST prediction generation, completed in 11.72 seconds. Native public TEST scoring took 158.669 seconds, including a 66.174-second target MWU map, using 15 spawned workers and a reused exact reference cache. Peak reported parent RSS is 14,684,912 KiB, not total cgroup peak. These measurements do not include downloading public source data or the historical model-development search.
