## Supplementary reproducibility code and frozen predictions for "Centered cross-context perturbation transfer on the public 2025 Virtual Cell Challenge benchmark": preprint_prompt_20261021_P2_AI.md

VCC-PREPRINT-V5-P2-AI

Write only one 40-80-word Methods paragraph disclosing the actual AI-assisted workflow for this manuscript. This is an addition to the P1 preprint, not a model redesign or a request to revise the entire paper.

Use these verified workflow facts only: OpenAI Codex assisted code development, experiment orchestration and analysis. ChatGPT, whose visible selected model was GPT-6, provided technical consultation and manuscript drafting. Numerical claims were checked against saved computation reports. Prompt and reply records were archived. The final frozen predictor is numerical and does not call a language model.

Do not claim human author review, author approval or other procedures that were not supplied. Do not describe an agent inside the predictor. Do not invent a model version, date, affiliation, data-use restriction or compliance assurance. Follow the same short, direct academic style and advisor rules as P1, with ASCII hyphens and no clause-linking em dashes.

Return only the paragraph in a single fenced markdown code block. Finish outside it with the exact marker VCC-PREPRINT-V5-P2-AI-END.
