## Supplementary reproducibility code and frozen predictions for "Centered cross-context perturbation transfer on the public 2025 Virtual Cell Challenge benchmark": preprint_prompt_20261021_P3.md

VCC-PREPRINT-V5-P3

Revise only the three items below from your P1 manuscript. Keep the title, authors and all other text unchanged. This is a targeted prose revision using verified project facts, not a request for further experiments or a full manuscript rewrite. Follow the same advisor rules: short direct academic prose, ASCII hyphens, no clause-linking em dashes, and few colons or semicolons. Use affirmative descriptions rather than a stack of defensive caveats. Do not invent claims, repository URLs, licenses or legal assurances.

1. Abstract. Write 150-180 words. Preserve the method and the retrospective released public 2025 benchmark setting with the external KOLF source released in 2026. State the verified raw metrics DES 0.371834, PDS 0.8614, MAE 0.302206 and composite score 0.341344. The archived champion score is 0.340052, with a difference of +0.001292. State the DES/PDS tradeoff without a significance claim. Keep detailed evaluation history in Methods rather than repeating it as a list of negations in the Abstract.

2. DiscussionScope. Replace only the final Discussion paragraph with about 50 words. Describe positively what the comparison covers: released public TEST data, reconstructed historical anchors, and a comparison with reported archived values. Do not claim equivalence to the original private scoring reference.

3. DataCodeAvailability. Replace the two availability paragraphs. The actual accompanying compact code archive is 7.6 MB. It contains frozen inference and scoring code, version pins, the exact 100-target prediction NPZ, the fitted source center, figures and evidence. Large input matrices are omitted and remain available from the original providers cited as references [1,3,5,6] in P1. Describe this actual supplement. Do not invent a repository URL or add unresolved license/legal boilerplate.

Return only three separate fenced markdown code blocks, labelled Abstract, DiscussionScope and DataCodeAvailability immediately before their respective blocks. End outside the code blocks with the exact marker VCC-PREPRINT-V5-P3-END.
