## Supplementary reproducibility code and frozen predictions for "Centered cross-context perturbation transfer on the public 2025 Virtual Cell Challenge benchmark": plot_center_ablation_v5_20261021.py

```
"""Public-center coefficient ablation using existing TRAIN/VALIDATION PDS only.

No predictions, fitting, TEST per-target data, confidence intervals or causal
interpretations are produced. Writes standalone SVG, 300dpi PNG and numbers.
"""
from pathlib import Path
import argparse,csv,hashlib,json,math
ROOT=Path(__file__).resolve().parents[1]
def sha(path):return hashlib.sha256(Path(path).read_bytes()).hexdigest()
def main():
    p=argparse.ArgumentParser(description=__doc__)
    p.add_argument('--base-report',type=Path,default=ROOT/'reports/fast_kolf_public_center_20261021_summary.json')
    p.add_argument('--extension-report',type=Path,default=ROOT/'reports/fast_kolf_public_center_extend_20261021_summary.json')
    p.add_argument('--output-prefix',type=Path,default=ROOT/'paper/figures/final_v5_center_ablation_20261021')
    args=p.parse_args();rows={};source={}
    for path in (args.base_report,args.extension_report):
        report=json.loads(path.read_text(encoding='utf-8-sig'))
        for row in report['all_rows']:
            gamma=float(row['gamma'])
            if gamma not in rows:rows[gamma]=row;source[gamma]=path
    gammas=(0.,.25,.5,1.,1.25,1.5,2.);selected_gamma=1.25
    if any(g not in rows for g in gammas):raise ValueError('Missing measured coefficient row')
    values=[]
    for gamma in gammas:
        row=rows[gamma];fold={r['fold']:float(r['pds']) for r in row['train_folds']}
        values.append({'gamma':gamma,'TRAIN_threefold_mean_PDS':float(row['train_pds']),
                       'VALIDATION50_PDS':float(row['validation_pds']),
                       'TRAIN_F0_PDS':fold['F0'],'TRAIN_F1_PDS':fold['F1'],'TRAIN_F2_PDS':fold['F2'],
                       'pooled_development_topN_joint_proxy':float(row['pooled_topN_joint_proxy']),
                       'selected_frozen_gamma':gamma==selected_gamma,
                       'source_report':str(source[gamma]),'source_report_sha256':sha(source[gamma])})
    import matplotlib
    matplotlib.use('Agg')
    import matplotlib.pyplot as plt
    from matplotlib.lines import Line2D
    import numpy as np
    plt.rcParams.update({'font.family':'DejaVu Sans','font.size':10,'axes.titlesize':11,
                         'axes.labelsize':10,'xtick.labelsize':9.5,'ytick.labelsize':9.5,
                         'legend.fontsize':9.5,'svg.fonttype':'none','axes.linewidth':.7,
                         'axes.spines.top':False,'axes.spines.right':False})
    train_color='#7F8793';val_color='#326DA8';grid='#E5E8EC';selected_color='#704E9D'
    fig,(ax,bx)=plt.subplots(1,2,figsize=(9.3,3.8),sharey=True,
                              gridspec_kw={'width_ratios':[1.8,1]})
    fig.subplots_adjust(left=.08,right=.985,bottom=.22,top=.86,wspace=.24)
    train=np.asarray([r['TRAIN_threefold_mean_PDS'] for r in values]);val=np.asarray([r['VALIDATION50_PDS'] for r in values])
    ax.plot(gammas,train,'o-',color=train_color,linewidth=1.6,markersize=5,label='TRAIN: 3-fold mean')
    ax.plot(gammas,val,'o-',color=val_color,linewidth=1.6,markersize=5,label='VALIDATION: 50 targets')
    selected=values[gammas.index(selected_gamma)];baseline=values[0]
    ax.axvline(selected_gamma,color=selected_color,linestyle=(0,(3,3)),linewidth=.9,alpha=.65,zorder=0)
    for key,color in [('TRAIN_threefold_mean_PDS',train_color),('VALIDATION50_PDS',val_color)]:
        ax.scatter([selected_gamma],[selected[key]],marker='*',s=145,c=color,edgecolors='white',linewidths=.65,zorder=5)
    ax.set_title('(a) Public-source mean subtraction',loc='left',pad=12)
    ax.set_xlabel('Subtraction coefficient γ');ax.set_ylabel('Centroid PDS (development)')
    ax.set_xticks(gammas,['0','0.25','0.5','1','1.25','1.5','2']);ax.set_xlim(-.07,2.07)
    ax.legend(loc='lower left',frameon=False,borderaxespad=.4,handlelength=2.)
    bx.set_title('(b) Individual TRAIN target folds',loc='left',pad=12)
    markers=('o','s','^');fold_records=[];handles=[]
    for name,marker in zip(('F0','F1','F2'),markers):
        old=baseline[f'TRAIN_{name}_PDS'];new=selected[f'TRAIN_{name}_PDS']
        bx.plot([0,1],[old,new],color='#A7AFBA',linewidth=1.25,zorder=1)
        bx.scatter([0],[old],marker=marker,s=48,color=train_color,edgecolor='white',linewidth=.55,zorder=3)
        bx.scatter([1],[new],marker=marker,s=48,color=val_color,edgecolor='white',linewidth=.55,zorder=3)
        handles.append(Line2D([],[],color='none',marker=marker,markersize=6,markerfacecolor=train_color,
                              markeredgecolor=train_color,label=name))
        fold_records.append({'fold':name,'gamma0_PDS':old,'selected_gamma1p25_PDS':new,'PDS_change':new-old})
    bx.set_xticks([0,1],['γ = 0','γ = 1.25\nselected']);bx.set_xlim(-.23,1.23)
    bx.legend(handles=handles,loc='lower center',frameon=False,ncol=3,columnspacing=.6,handletextpad=.3)
    for axis in (ax,bx):
        axis.set_ylim(.82,.89);axis.set_yticks([.83,.85,.87,.89])
        axis.grid(axis='y',color=grid,linewidth=.7,zorder=0);axis.set_axisbelow(True)
    ax.text(selected_gamma,.8865,'Selected γ = 1.25',ha='center',va='top',fontsize=9.5,color=selected_color,
            bbox={'facecolor':'white','edgecolor':'none','pad':2.})
    fig.text(.08,.04,'TRAIN: three held-out target folds. VALIDATION: official development panel.',
             ha='left',fontsize=9.5,color='#505864')
    prefix=args.output_prefix;prefix.parent.mkdir(parents=True,exist_ok=True);artifacts=[]
    for ext in ('svg','png'):
        path=prefix.with_suffix('.'+ext);fig.savefig(path,dpi=300,facecolor='white')
        artifacts.append({'path':str(path),'sha256':sha(path),'bytes':path.stat().st_size})
    plt.close(fig)
    csv_path=prefix.with_name(prefix.name+'_values.csv')
    with csv_path.open('w',newline='',encoding='utf-8') as f:
        w=csv.DictWriter(f,fieldnames=list(values[0]));w.writeheader();w.writerows(values)
    metadata={'figure':'Empirical public-center coefficient ablation','selected_gamma':selected_gamma,
              'metric':'Centroid development PDS; native DES and TEST metrics are not plotted',
              'TRAIN_scope':'Original3xF100/V50 target CV; equal-weight fold mean',
              'VALIDATION_scope':'Official50-target development panel',
              'source_reports':{str(p):sha(p) for p in (args.base_report,args.extension_report)},
              'code_sha256':sha(__file__),'matplotlib_version':matplotlib.__version__,
              'figsize_inches':[9.3,3.8],'png_dpi':300,'minimum_font_points':9.5,
              'all_points':values,'paired_folds':fold_records,
              'selected_minus_gamma0':{'TRAIN_PDS':selected['TRAIN_threefold_mean_PDS']-baseline['TRAIN_threefold_mean_PDS'],
                                      'VALIDATION_PDS':selected['VALIDATION50_PDS']-baseline['VALIDATION50_PDS']},
              'error_bars_or_confidence_intervals':False,'curve_interpolation':'Straight segments between measured coefficients',
              'biological_causality_claimed':False,'TEST_per_target_data_used':False,'large_arrays_read':False,
              'values_csv':str(csv_path),'values_csv_sha256':sha(csv_path),'artifacts':artifacts}
    path=prefix.with_name(prefix.name+'_metadata.json');path.write_text(json.dumps(metadata,indent=2)+'\n',encoding='utf-8')
    print(json.dumps({'state':'complete','artifacts':artifacts,'values':str(csv_path),'metadata':str(path)},indent=2))
if __name__=='__main__':main()
```
