## Supplementary reproducibility code and frozen predictions for "Centered cross-context perturbation transfer on the public 2025 Virtual Cell Challenge benchmark": plot_final_v5_comparison_20261021.py

```
"""Standalone raw-metric and fixed-anchor v5 comparison figure; SVG and PNG.

Consumes only compact verified evidence JSON. The archived champion is a
reported reference, not a rerun. This script performs no model computation.
"""
from pathlib import Path
import argparse,csv,hashlib,json,math

ROOT=Path(__file__).resolve().parents[1]
METRICS=('DES','PDS','MAE')

def sha(path):return hashlib.sha256(Path(path).read_bytes()).hexdigest()

def main():
    p=argparse.ArgumentParser(description=__doc__)
    p.add_argument('--input',type=Path,default=ROOT/'paper/evidence/final_results_v5.json')
    p.add_argument('--output-prefix',type=Path,default=ROOT/'paper/figures/final_v5_comparison_20261021')
    args=p.parse_args()
    data=json.loads(args.input.read_text(encoding='utf-8-sig'))
    test=data['TEST'];archive=data['archived_champion'];anchors=data['anchors']
    public_raw={k:float(test['raw'][k]) for k in METRICS}
    archive_raw={k:float(archive[k]) for k in METRICS}
    public_norm={k:float(test['normalized_components'][k]) for k in METRICS}
    archive_norm={k:max(0.,(archive_raw[k]-anchors[k])/(1.-anchors[k])) for k in ('DES','PDS')}
    archive_norm['MAE']=max(0.,1.-archive_raw['MAE']/anchors['MAE'])
    public_overall=float(test['historical_anchor_score'])
    archive_overall=float(archive['overall'])
    if not all(math.isfinite(v) for v in list(public_raw.values())+list(archive_raw.values())+
               list(public_norm.values())+list(archive_norm.values())+[public_overall,archive_overall]):
        raise ValueError('Complete finite measured/reported results required')
    import matplotlib
    matplotlib.use('Agg')
    import matplotlib.pyplot as plt
    from matplotlib.patches import Patch
    from matplotlib.ticker import MaxNLocator
    import numpy as np
    plt.rcParams.update({'font.family':'DejaVu Sans','font.size':10,'axes.titlesize':11,
                         'axes.labelsize':10,'xtick.labelsize':10,'ytick.labelsize':9.5,
                         'legend.fontsize':10,'svg.fonttype':'none','pdf.fonttype':42,
                         'axes.spines.top':False,'axes.spines.right':False,'axes.linewidth':.7,
                         'hatch.linewidth':.55,'savefig.facecolor':'white'})
    archive_color='#A1A6AF';model_color='#326DA8';edge='#5B626C';grid='#E5E8EC'
    figsize=(10.8,6.2)
    fig=plt.figure(figsize=figsize)
    gs=fig.add_gridspec(2,3,height_ratios=[1,1.06],left=.08,right=.985,bottom=.11,top=.875,
                        hspace=.52,wspace=.40)
    raw_axes=[]
    for col,key in enumerate(METRICS):
        ax=fig.add_subplot(gs[0,col]);raw_axes.append(ax)
        vals=[archive_raw[key],public_raw[key]]
        bars=ax.bar([0,1],vals,width=.55,color=[archive_color,model_color],
                    edgecolor=[edge,model_color],linewidth=.65,zorder=3)
        bars[0].set_hatch('///')
        ymax=1. if key!='MAE' else math.ceil(max(vals)*1.22*10)/10
        ax.set_ylim(0,ymax);ax.set_xlim(-.65,1.65);ax.set_xticks([])
        ax.set_title(f"({chr(97+col)}) Raw {key} {'↓' if key=='MAE' else '↑'}",loc='left',pad=11)
        ax.set_ylabel('Mean absolute error' if key=='MAE' else 'Score')
        ax.grid(axis='y',color=grid,linewidth=.7,zorder=0);ax.set_axisbelow(True)
        ax.yaxis.set_major_locator(MaxNLocator(nbins=4,min_n_ticks=4))
        for b,v in zip(bars,vals):
            ax.text(b.get_x()+b.get_width()/2,v+.024*ymax,f'{v:.4f}',ha='center',va='bottom',fontsize=10)
        ax.spines['bottom'].set_visible(False)
    ax=fig.add_subplot(gs[1,:]);width=.32;x=np.arange(4)
    av=[archive_norm[k] for k in METRICS]+[archive_overall]
    pv=[public_norm[k] for k in METRICS]+[public_overall]
    for dx,vals,color,hatch in [(-width/2,av,archive_color,'///'),(width/2,pv,model_color,None)]:
        bars=ax.bar(x+dx,vals,width,color=color,hatch=hatch,edgecolor=edge if hatch else color,
                    linewidth=.65,zorder=3)
        for j,(b,v) in enumerate(zip(bars,vals)):
            label=f'{v:.6f}' if j==3 else f'{v:.4f}'
            ax.text(b.get_x()+b.get_width()/2,v+.018,label,ha='center',va='bottom',fontsize=9.5)
    ax.set_title('(d) Fixed-anchor components and overall',loc='left',pad=11)
    ax.set_ylabel('Normalized score');ax.set_xticks(x,['DES','PDS','MAE','Overall'])
    ax.set_xlim(-.6,3.6);ax.set_ylim(0,1.0);ax.set_yticks(np.arange(0,1.01,.25))
    ax.axvline(2.5,color='#B7BCC4',linewidth=.8,linestyle=(0,(3,3)))
    ax.grid(axis='y',color=grid,linewidth=.7,zorder=0);ax.set_axisbelow(True)
    delta=public_overall-archive_overall
    ax.text(.995,.96,f'Overall Δ = {delta:+.8f}',transform=ax.transAxes,
            ha='right',va='top',fontsize=10,color='#2F3742')
    legend=[Patch(facecolor=archive_color,edgecolor=edge,hatch='///'),
            Patch(facecolor=model_color,edgecolor=model_color)]
    fig.legend(legend,['2025 champion — reported archive','Frozen v5 — computed public TEST100'],
               loc='upper center',bbox_to_anchor=(.52,.985),frameon=False,ncol=2,columnspacing=2.8)
    fig.text(.08,.025,'Archive values are reported, not rerun. Public TEST100 is evaluated locally.',
             fontsize=9.5,color='#505864',ha='left')
    prefix=args.output_prefix;prefix.parent.mkdir(parents=True,exist_ok=True);artifacts=[]
    for ext in ('svg','png'):
        path=prefix.with_suffix('.'+ext);fig.savefig(path,dpi=300,facecolor='white')
        artifacts.append({'path':str(path),'sha256':sha(path),'bytes':path.stat().st_size})
    plt.close(fig)
    values=[]
    for ident,label,raw,norm,score in [
        ('reported_archive','2025 champion reported archive',archive_raw,archive_norm,archive_overall),
        ('computed_public_TEST100','Frozen v5 computed public TEST100',public_raw,public_norm,public_overall)]:
        values.append({'series_id':ident,'series_label':label,**{f'raw_{k}':raw[k] for k in METRICS},
                       **{f'normalized_{k}':norm[k] for k in METRICS},'overall':score})
    csv_path=prefix.with_name(prefix.name+'_values.csv')
    with csv_path.open('w',newline='',encoding='utf-8') as f:
        w=csv.DictWriter(f,fieldnames=list(values[0]));w.writeheader();w.writerows(values)
    metadata={'figure':'Final fixed v5 aggregate comparison','input_json':str(args.input.resolve()),
              'input_sha256':sha(args.input),'code_sha256':sha(__file__),
              'matplotlib_version':matplotlib.__version__,'figsize_inches':list(figsize),
              'png_dpi':300,'minimum_font_points':9.5,'colors':{'reported_archive':archive_color,'v5':model_color},
              'numbers':values,'anchors':anchors,'overall_difference':delta,
              'normalization_formulas':{'DES':'max(0,(DES-a_DES)/(1-a_DES))',
                                        'PDS':'max(0,(PDS-a_PDS)/(1-a_PDS))',
                                        'MAE':'max(0,1-MAE/a_MAE)','overall':'mean of the three normalized components'},
              'archive_source':archive,'TEST_source_report':test.get('source_report'),
              'TEST_source_report_sha256':test.get('source_report_sha256'),
              'archive_rerun':False,'error_bars_included':False,'large_arrays_read':False,
              'official_private_leaderboard_equivalence':test.get('official_private_leaderboard_equivalence',False),
              'raw_y_limits':{'DES':[0,1],'PDS':[0,1],'MAE':list(raw_axes[2].get_ylim())},
              'normalized_and_overall_y_limits':[0,1],
              'values_csv':str(csv_path),'values_csv_sha256':sha(csv_path),'artifacts':artifacts}
    path=prefix.with_name(prefix.name+'_metadata.json')
    path.write_text(json.dumps(metadata,indent=2)+'\n',encoding='utf-8')
    print(json.dumps({'state':'complete','artifacts':artifacts,'values':str(csv_path),'metadata':str(path)},indent=2))

if __name__=='__main__':main()
```
