## Supplementary reproducibility code and frozen predictions for "Centered cross-context perturbation transfer on the public 2025 Virtual Cell Challenge benchmark": render_mathtext_assets_v5.py

```
"""Render a compact preprint_math_job_v5 JSON into transparent PNG math assets.

LaTeX fields contain one MathText equation without outer dollar delimiters.
Unsupported syntax is an error; no source-text fallback is drawn. No PDF is
created. Matplotlib/Pillow are imported only for an actual render invocation.
"""
from pathlib import Path
import argparse,hashlib,json,math,re,sys

def sha(path):return hashlib.sha256(Path(path).read_bytes()).hexdigest()

def mathtext_aliases(latex):
    """Equivalent TeX spellings only; original source remains in manifest."""
    rendered=re.sub(r'\\ge(?![A-Za-z])',r'\\geq',latex)
    rendered=re.sub(r'\\le(?![A-Za-z])',r'\\leq',rendered)
    rendered=re.sub(r'\\mathcal\s+([A-Za-z])',r'\\mathcal{\1}',rendered)
    return rendered

def render(job_path,out):
    job=json.loads(job_path.read_text(encoding='utf-8-sig'))
    if job.get('schema')!='preprint_math_job_v5':raise ValueError('Expected preprint_math_job_v5 schema')
    dpi=int(job.get('dpi',300))
    if dpi<=0:raise ValueError('Positive DPI required')
    equations=job.get('equations')
    if not isinstance(equations,list):raise ValueError('equations must be a list')
    import matplotlib
    matplotlib.use('Agg')
    from matplotlib.mathtext import MathTextParser
    from matplotlib.font_manager import FontProperties
    import numpy as np
    import PIL
    from PIL import Image
    matplotlib.rcParams['mathtext.fontset']='dejavuserif'
    parser=MathTextParser('agg');out.mkdir(parents=True,exist_ok=True);assets={};errors=[]
    for item in equations:
        ident=str(item['id']);kind=item['kind'];latex=item['latex']
        if not re.fullmatch(r'[A-Za-z0-9_-]{1,80}',ident) or ident in assets:raise ValueError('Invalid or duplicate equation ID')
        if kind not in ('inline','display') or not isinstance(latex,str) or not latex.strip():raise ValueError('Equation kind/latex required')
        if '$' in latex:raise ValueError('latex must not contain outer or embedded dollar delimiters: '+ident)
        size=float(item.get('font_size_pt',11. if kind=='inline' else 12.))
        if not math.isfinite(size) or size<=0:raise ValueError('Positive finite font size required')
        prop=FontProperties(family='DejaVu Serif',size=size)
        rendered_latex=mathtext_aliases(latex)
        try:parsed=parser.parse('$'+rendered_latex+'$',dpi=dpi,prop=prop)
        except Exception as exc:
            errors.append({'id':ident,'latex':latex,'kind':kind,'error':str(exc)})
            continue
        alpha=np.asarray(parsed.image,dtype=np.uint8)
        if alpha.ndim!=2 or not alpha.size:raise ValueError('Empty equation raster: '+ident)
        height,width=map(int,alpha.shape)
        rgba=np.zeros((height,width,4),np.uint8);rgba[:,:,3]=alpha
        name=ident+'.png';path=out/name;Image.fromarray(rgba).save(path,dpi=(dpi,dpi))
        depth_px=float(parsed.depth)
        assets[ident]={'latex':latex,'rendered_latex':rendered_latex,'syntax_alias_applied':latex!=rendered_latex,
                       'kind':kind,'png_file':name,'width_px':width,'height_px':height,
                       'width_pt':width*72./dpi,'height_pt':height*72./dpi,'depth_px':depth_px,
                       'depth_pt':depth_px*72./dpi,'inline_valign_pt':-depth_px*72./dpi,
                       'dpi':dpi,'font_size_pt':size,'font_family':'DejaVu Serif','mathtext_fontset':'dejavuserif',
                       'parser_width_px':float(parsed.width),'parser_height_px':float(parsed.height),
                       'parser_ox_px':float(parsed.ox),'parser_oy_px':float(parsed.oy),
                       'png_sha256':sha(path),'transparent_background':True}
    if errors:
        (out/'errors.json').write_text(json.dumps({'schema':'preprint_math_errors_v5','errors':errors},indent=2)+'\n',encoding='utf-8')
        raise ValueError('Unsupported MathText equations: '+json.dumps(errors))
    if (out/'errors.json').exists():(out/'errors.json').unlink()
    manifest={'schema':'preprint_math_assets_v5','source_job_path':str(job_path.resolve()),
              'source_job_sha256':sha(job_path),'renderer_code_sha256':sha(__file__),
              'dpi':dpi,'matplotlib_version':matplotlib.__version__,'pillow_version':PIL.__version__,
              'font_family':'DejaVu Serif','mathtext_fontset':'dejavuserif',
              'baseline_policy':'For ReportLab inline image use valign=-depth_pt; PNG dimensions include exact agg raster extent',
              'png_paths_relative_to':'manifest parent','equations':assets}
    path=out/'manifest.json';path.write_text(json.dumps(manifest,indent=2)+'\n',encoding='utf-8')
    return {'state':'complete','manifest':str(path),'equations':len(assets),'dpi':dpi}

if __name__=='__main__':
    p=argparse.ArgumentParser(description=__doc__)
    p.add_argument('--input',type=Path,required=True,help='preprint_math_job_v5 JSON')
    p.add_argument('--output-dir',type=Path,required=True)
    args=p.parse_args()
    try:print(json.dumps(render(args.input,args.output_dir)))
    except Exception as exc:
        print(json.dumps({'state':'failed','error_type':type(exc).__name__,'message':str(exc)}),file=sys.stderr)
        raise SystemExit(2)
```
