## Supplementary reproducibility code and frozen predictions for "Centered cross-context perturbation transfer on the public 2025 Virtual Cell Challenge benchmark": README.md

# Frozen numerical v5 reproducibility source package

This is the final compact source/evidence supplement, including the completed
Edge-authored manuscript Markdown. The final main manuscript PDF is provided
separately. No model fitting, prediction, evaluation, new numerical test or PDF
creation was run while assembling this package.

## Contents

- `outputs/predictions_v5.npz`: final 100 × 18,080 Float32 predicted means, with
  `targets`, `genes`, and `predicted_means` members.
- `outputs/fitted_public_center.npz`: TRAIN-fitted public KOLF mean and observed
  fit counts. It is shared by the validation replay and final prediction.
- `scripts/`, `ops/fast_gct_de_krr_rank_20261020.py`, and `src/virtualcell/`:
  the frozen numerical model, initializer, source/scaler math, and exact scorer.
- `runs/training_meanflat_DES_prepare_v1/scripts/`: the four exact mathematical
  helper modules actually imported by the final scorer. These are retained
  under their existing relative library path. Their historical runner entry
  points are not required; final scoring imports the math functions only.
- `configs/`: the final model freeze, actual scoring config, and source pins.
- `paper/evidence/`: compact results, initializer specification, author metadata.
- `paper/figures/`: SVG, 300 dpi PNG, complete plotting numbers and metadata.
- `paper/manuscript_v5.md`, its assembly receipt/source, and `paper/edge/`:
  final manuscript and the three writing prompts, replies and model observations.
- `paper/build_preprint_pdf_v5.py` and `paper/render_mathtext_assets_v5.py`:
  small standalone typesetting/equation-rendering sources. The separately
  provided main PDF and regenerable math PNG cache are not bundled.
- `provider_links.json`, `external_assets.json`, `runtime/pip_freeze.txt`,
  `runtime/retrieval_receipt.json`, `manifest.json`, and `filelist.csv`:
  public providers, omitted data dependencies, runtime and file identities.

Pure catalogue/GenePT helper functions remain in original numerical modules
for source fidelity. The selected model does not consume an internal prediction
agent, agent responses/caches, donor residuals, residual programs or GenePT.

## Exact included derived outputs

```text
predictions_v5.npz
  cea4874ec557b56622d66e5a414ebe8ecb59fee078e5f613a3316cbc9fffb4df
fitted_public_center.npz
  f7b084cf6117baea7a9c6a853bdae77a79588ce9b1d69659deeb1a523296bae7
```

The included prediction is the frozen output that was scored. The validation
replay changed 0 of 904,000 means and had maximum absolute difference 0.

## Runtime and file layout

The original compute host is 99.32 and project root is
`/data1/luanhaoyang/virtualcell`. The original interpreter is
`envs/vcc2025-py311/bin/python` (Python 3.11.10). Prediction used NumPy 2.4.6,
SciPy 1.17.1 and scikit-learn 1.9.1; evaluation used cell-eval 0.6.6,
pdex 0.1.27 and Polars 2.0.0. Figure production used Matplotlib 3.11.2.
`runtime/pip_freeze.txt` records the readable environment at packaging time,
including figure dependencies, without credentials or environment variables.

Commands below are Linux commands for the original project layout and
already prepared assets. They are documented commands, not executed packaging
steps. Use a fresh output directory. Source files preserve the original
relative paths and some frozen modules retain an absolute project ROOT.
Reproducing on another host requires recreating that layout or adapting a
separate working copy and recording the changed source hashes.

## Replot the included aggregates

From the unpacked bundle root, plotting needs only the included JSON/CSV
sources and a Matplotlib environment. Both scripts determine ROOT from their
own location. On the original server, use its existing interpreter:

```bash
/data1/luanhaoyang/virtualcell/envs/vcc2025-py311/bin/python -B paper/plot_final_v5_comparison_20261021.py
/data1/luanhaoyang/virtualcell/envs/vcc2025-py311/bin/python -B paper/plot_center_ablation_v5_20261021.py
```

## Rebuild the separately provided manuscript PDF

The final Markdown is included byte-for-byte with SHA256
`99469851f1ede6b9fa3198a2ee1966a0f7613e2d7eee14848ccb91506b04055a`.
The assembly receipt records the exact source replies and mechanical fixes;
its historical state describes the time of assembly. The archived model
metadata distinguishes the visibly selected GPT-6 model from Pro settings
observed earlier in the same session. Conversation URLs are provenance links,
not access tokens or a dependency for running the numerical model.

The pure renderers need Matplotlib and ReportLab. Generate the small equation
assets from Markdown before typesetting; the source package deliberately omits
the regenerable assets and the final PDF. From the unpacked bundle root:

```bash
python paper/build_preprint_pdf_v5.py --collect-math paper/math_job_v5.json
python paper/render_mathtext_assets_v5.py --input paper/math_job_v5.json --output-dir paper/math_assets_v5
python paper/build_preprint_pdf_v5.py --output output/pdf/vcc2025_centered_transfer_preprint.pdf
```

The PDF builder reads the manuscript author block and the two figure PNGs. The
writing records contain no server credential files or browser-control code.

## Repeat v5 inference on the prepared original server

Required but not included: TRAIN control/raw bank, TRAIN150 DE prior, aligned
TRAIN/query public KOLF banks, and the original numeric feature archive
containing `expression_direction_source` and `de_score`. Their exact paths and
hashes are in `external_assets.json` and the frozen prediction receipt.

```bash
cd /data1/luanhaoyang/virtualcell
envs/vcc2025-py311/bin/python -B scripts/predict_numeric_kolf_centered_R1_v5.py \
  --features runs/numeric_test_primary_20261021_v1/predictions.npz \
  --query-kolf-bank data/processed/kolf_TEST_public_20261021_v1/source_bank.npz \
  --output runs/reproduce_v5_prediction_NEW
```

The script fits the independent KOLF DE head to the source-known TRAIN150
subset, fits its public mean on TRAIN source rows, and reads only prediction
features for queries. It does not read query expression or evaluation reports.

## Recreate the preceding numeric features, if needed

The initializer consumes only the `B`, `targets`, and `genes` members of the
existing historical numerical archive. That whole archive is not distributed
because it also contains unused historical arms. B itself is defined fully in
`paper/evidence/B_initializer_v5.json` and its original pure functions are
`fit_base_states`, `predict_base_delta`, `pack_base`, and `unpack_base` in
`src/virtualcell/final_training150_base_catalogue_v1.py`.

The B fit state, original public scaler/basis features, TRAIN teacher, and
prepared query feature matrices remain external dependencies. No historical
approval/agent orchestration is needed to understand the B math. A clean-host
end-to-end B reconstruction still requires those pinned public/large assets;
this bundle does not claim a one-command reconstruction from missing data.

```bash
envs/vcc2025-py311/bin/python -B scripts/predict_numeric_ranktransfer_v1.py \
  --legacy-b runs/final150_agent_prediction_assembly_v1/vcc2025_test_v1/consume/all_four_external_uncalibrated_F64.npz \
  --xatlas-bank data/processed/arc_pie_xatlas_test_selected_v1/source_bank.npz \
  --gemz-bank data/processed/gct_TEST_public_source_bank_v1/source_bank.npz \
  --alpha 1 --beta .5 --eps .01 --power 2.5 --scale 1.5 \
  --output runs/reproduce_original_numeric_features_NEW
```

Only its `expression_direction_source` and `de_score` fields are inputs to
v5. The earlier initializer's power-curve predicted means are not used by v5.
Use `--gemz-bank` as shown; the optional old validation fallback import path
is intentionally outside this compact package. The included source preparation
modules require their pinned public objects/metadata. TRAIN preparation uses
normalization target 54611 explicitly; the historical helper default 10000 is
not the final model's training normalization.

## Re-evaluate the included frozen prediction

This requires the original full TRAIN non-targeting reference H5AD, native
real TEST cache, public counts and exact process-reference cache. None of
those expression matrices or large caches is included. The actual config
references the canonical frozen prediction path; the included output has the
same SHA256. On the original server, with those assets present:

```bash
envs/vcc2025-py311/bin/python -B scripts/score_frozen_meanflat_panel_process_v1.py \
  --process-reference-cache data/processed/mwu_constant_reference_process_20261021_v1 \
  --process-workers 15 \
  --config configs/numeric_test_final_metrics_20261021_v5.json \
  --outdir runs/reproduce_v5_final_metrics_NEW
```

The process wrapper changes transport only. It retains the exact constant-MWU
core, global BH family, native Polars bulk reduction, and official metric
functions. Cached reference manifests contain original absolute core paths;
retain those paths or rebuild the process cache with the same core on a clean
host. The direct final-v2 scorer can be used without the process wrapper if no
process cache is available, but still requires the real cache and full TRAIN
control H5AD.

If the native real cache is absent, its included preparer requires the large
official public count H5AD and a frozen prediction identity before evaluation:

```bash
envs/vcc2025-py311/bin/python -B scripts/compute_real_de_panel.py \
  --real-h5ad data/test/adata_Test.h5ad \
  --prediction-archive runs/numeric_kolf_centered_R1_v5_20261021/test_predictions/predictions.npz \
  --prediction-sha256 cea4874ec557b56622d66e5a414ebe8ecb59fee078e5f613a3316cbc9fffb4df \
  --input-space counts --workers 8 --batch-size 100 \
  --outdir runs/reproduce_public_TEST_real_cache_NEW
```

A copied scoring config must then point to the new cache and record its own
hash. This preparation materializes a large Float32 memmap and full-panel DE;
there is no standalone reevaluation without these real-data dependencies.

## Scope of the recorded comparison

The results are a retrospective local public-TEST fixed-anchor reconstruction.
The archive champion is a reported reference, not a checkpoint rerun. The full
H5AD submission pipeline/private leaderboard was not executed. Current KOLF
browser-release eligibility during 2025 is not established. Exact raw values,
normalization formulas, prior public-TEST evaluation history and source dates
are retained in the canonical evidence JSON.

No raw expression, large databases, agent caches, server access records,
credential helper files, API keys or authentication material are included.
