## Supplementary reproducibility code and frozen predictions for "Centered cross-context perturbation transfer on the public 2025 Virtual Cell Challenge benchmark": historical_reference_anchor_20261015.md

# 2025 VCC historical-reference anchor check (2026-10-15 UTC)

This is a read-only audit of Arc's public leaderboard, dataset listing, object metadata, and `cell-eval` source. No test H5AD bytes, expression values, or test-derived statistics were read. The check does **not** establish that the public test H5AD is identical to the private final-scoring reference.

## What is verified

| Question | Result and source |
| --- | --- |
| Historical score to exceed | The current [Arc 2025 archive](https://virtualcellchallenge.org/2025) serves BM_xTVC / `xTVCM_v01` at rank 1, overall `0.3400522468873988`, raw DES `0.34882155189465897`, PDS `0.8719999999999999`, MAE `1.0258748292922975`. The page's `self.__next_f.push` `finalEntries` was parsed fresh at `2026-10-15T04:15:41Z`; HTML SHA-256 `faf06af2010fd0e953aa1cb1c9debdc6b4e20f505d32c5d6f4dad5b08de21810`. [Arc's winner announcement](https://arcinstitute.org/news/virtual-cell-challenge-2025-wrap-up) binds BM_xTVC to the model name xTrimoSCPerturb. |
| Historical numeric implementation | Arc's [parity note](https://github.com/ArcInstitute/cell-eval2/blob/0ce25cf4c495e76e60dfca4ddc8f822507aa0e18/docs/parity/v1-parity-divergences.md) names `cell-eval 0.6.6` + `pdex 0.1.27` as its VCC `v0_6_6` reference and reports bit-identical leaderboard replay. Its reproduction environment is explicitly **not committed**. The public [0.6.6 scorer](https://github.com/ArcInstitute/cell-eval/blob/v0.6.6/src/cell_eval/_score.py) defines the clipped three-metric normalization. This identifies a reference *code path*, not the full historical runtime or real-side inputs. |
| Recoverable normalization constants | The local [100-row archive reconstruction](archived_baseline_inference.json) gives DES `0.09505113849605429`, PDS `0.5082`, MAE `0.0249575331993401`, reproducing all visible scores with maximum residual `5.55e-17`. They are inferred anchors, **not an official baseline aggregate CSV**. MAE has one independently active row. |
| Official public data lineage | The [Arc Atlas README](https://github.com/ArcInstitute/arc-virtual-cell-atlas/blob/main/virtual-cell-challenge/README.md) says its released train/validation/held-out test datasets were leveraged throughout the Challenge. [Arc's 2026 announcement](https://arcinstitute.org/news/virtual-cell-challenge-2026) calls the full 2025 data, including formerly held-out test perturbations, downloadable. These statements establish official dataset lineage, not value/control/preprocessing identity with the scoring server. |
| Current public test-object identity | Fresh [GCS metadata-only lookup](https://storage.googleapis.com/storage/v1/b/arc-institute-virtual-cell-atlas/o/virtual-cell-challenge%2F2025%2Ftest%2Fadata_Test.h5ad?fields=name%2Csize%2Cgeneration%2Cmetageneration%2Ccrc32c%2CtimeCreated%2Cupdated%2Cmetadata%2Cmd5Hash) reports generation `1765905295402700`, 11,950,739,168 bytes, CRC32C `JMP5RQ==`, created `2025-12-16T17:14:55.529Z`, metageneration 1, and no returned custom lineage metadata or MD5. A later bucket creation date does not prove reprocessing; copying an unchanged object also creates a new generation. |
| Scope of publicly listed artifacts | A fresh [versioned 2025-prefix inventory](https://storage.googleapis.com/storage/v1/b/arc-institute-virtual-cell-atlas/o?prefix=virtual-cell-challenge%2F2025%2F&versions=true&fields=items%28name%2Cgeneration%2Csize%29%2CnextPageToken) returned 49 items, no next page, and no baseline aggregate, real DE table, scorer manifest, or historical submission artifact. The current public [cell-eval2 complete tree](https://api.github.com/repos/ArcInstitute/cell-eval2/git/trees/0ce25cf4c495e76e60dfca4ddc8f822507aa0e18?recursive=1) has 230 entries (`truncated=false`) and no committed VCC replay recipe or baseline CSV. These are searches of stated public locations, not a proof of global nonexistence. |
| Does the archived leaderboard identify its scored reference? | Freshly parsed top-100 `finalEntries` have `panelId`, `partition`, and `anchorVersion` all null in all 100 visible rows. The public rows supply scores but no reference object generation/hash bridge. This is reproducible with the local [archive parser](infer_archived_baseline.py) against the [Arc 2025 archive](https://virtualcellchallenge.org/2025). |

## Remaining identity gap and safe checks

The official statement that Arc released the Challenge's test perturbations is strong provenance but does not expose a hash or logical mapping to the private reference scored in November 2025. The recovered anchors likewise cannot identify a high-dimensional matrix. The original final-test baseline aggregate CSV, private reference/control manifest, production real DE cache and its clip settings, complete transitive runtime lock, and an authenticated 2025 prediction with archived raw metrics remain missing from the inspected public sources. The [0.6.6 baseline CLI](https://github.com/ArcInstitute/cell-eval/blob/v0.6.6/src/cell_eval/_cli/_baseline.py) precomputes DE with `clip_value=2**20`, while ordinary [pdex 0.1.27](https://github.com/ArcInstitute/pdex/blob/pdex-0.1.27/src/pdex/_single_cell.py) recomputation defaults to 20; an unspecified historical cached-DE path therefore matters to DES.

Before the final freeze, permitted read-only checks are generation-specific API/HEAD metadata, the small public target-count CSV, exact source-tree identities, and train-only evaluator sensitivity tests. After all model and agent predictions are frozen and test expression is authorized for scoring, a scoring-only worker can build the public-panel baseline and compare **all three** raw aggregates to the recovered anchors under preregistered scorer/cache branches. A match supports numerical continuity but alone does not prove full reference identity; a logged original Arc reference manifest or replay of authentic archived predictions against the public panel would be stronger. Until then report a comparison as provisional historical replay, not an official competition win or a reproduction of the champion model.
