## Supplementary reproducibility code and frozen predictions for "Centered cross-context perturbation transfer on the public 2025 Virtual Cell Challenge benchmark": mwu_constant_reference.py

```
"""Experimental constant-target MWU specialization, never the default scorer.

This reproduces pinned SciPy 1.17.1's per-gene float32/float64 rank and tie
reductions, including underflow. A float64 analytical p value is not substituted
for the official float32 p value. Only genuinely constant targets are accepted.
"""
from __future__ import annotations

import hashlib
import importlib.metadata
import json
from pathlib import Path
import time

import numpy as np

VERSION = "constant_MWU_scipy1171_dtype_v1"


class ConstantReference:
    def __init__(self,control,genes):
        if importlib.metadata.version("scipy") != "1.17.1":
            raise ValueError("Constant specialization requires audited SciPy1.17.1")
        if importlib.metadata.version("pdex") != "0.1.27":
            raise ValueError("Constant specialization requires audited pdex0.1.27")
        control=np.asarray(control)
        if control.ndim != 2 or min(control.shape) < 1 or control.dtype not in (np.dtype("float32"),np.dtype("float64")):
            raise ValueError("Reference must be a nonempty float32/float64 log1p matrix")
        if not np.isfinite(control).all() or np.any(control < 0) or np.any(control >= 15):
            raise ValueError("Reference values must be finite nonnegative log1p below15")
        genes=list(map(str,genes))
        if len(genes)!=control.shape[1] or len(set(genes))!=len(genes):
            raise ValueError("Reference gene identities must be unique and aligned")
        started=time.monotonic()
        self.genes,self.n,self.dtype=genes,len(control),control.dtype
        self.reference=control
        self.sorted=np.sort(control.T,axis=1)
        self.starts=[]
        self.counts=[]
        self.values=[]
        self.reference_means=[]
        import pdex._single_cell as official
        if official.use_experimental:
            raise ValueError("Experimental pdex implementation is incompatible")
        for j,row in enumerate(self.sorted):
            starts=np.flatnonzero(np.r_[True,row[1:]!=row[:-1]])
            counts=np.diff(np.r_[starts,self.n])
            self.starts.append(starts)
            self.counts.append(counts)
            self.values.append(row[starts])
            self.reference_means.append(official._sample_mean(np.ascontiguousarray(control[:,j]),is_log1p=True,exp_post_agg=True))
        self.build_elapsed_seconds=time.monotonic()-started
        self.reference_sha256=hashlib.sha256(np.ascontiguousarray(control).tobytes()).hexdigest()

    def gene(self,gene_index,mean,n_cells):
        """Return official-typed U/p/means/FC for one constant target column."""
        from scipy import special
        from scipy.stats import mannwhitneyu
        import scipy.stats._mannwhitneyu as mwu
        import pdex._single_cell as official
        j=int(gene_index)
        n=int(n_cells)
        if n<1 or n!=n_cells:
            raise ValueError("Positive integer target count required")
        mu=self.dtype.type(mean)
        if not np.isfinite(mu) or not 0<=mu<15:
            raise ValueError("Constant target mean outside registered log domain")
        less=int(np.searchsorted(self.sorted[j],mu,side="left"))
        equal=int(np.searchsorted(self.sorted[j],mu,side="right"))-less
        # Recreate SciPy's rank-data tie layout, not a compact tie-count sum:
        # float32 pairwise reduction depends on the intervening zero positions.
        t=np.zeros(n+self.n,dtype=self.dtype)
        starts=self.starts[j]+(self.values[j]>mu)*n
        t[starts]=self.counts[j]
        t[less]=equal+n
        rank=self.dtype.type(less+(equal+n+1)/2)
        ranks=np.full(n,rank,dtype=self.dtype)
        R1=np.sum(ranks,axis=-1)
        U1=R1-n*(n+1)/2
        U2=n*self.n-U1
        method=mwu._mwu_choose_method(n,self.n,np.any(t>1))
        if method=="exact":
            # n=1 and a completely untied combined sample: preserve the actual
            # SciPy exact distribution rather than assuming asymptotic behavior.
            actual=mannwhitneyu(np.full(n,mu,dtype=self.dtype),np.ascontiguousarray(self.reference[:,j]),use_continuity=True)
            U1,p=actual.statistic,actual.pvalue
        else:
            U=np.maximum(U1,U2)
            z=mwu._get_mwu_z(U,n,self.n,t,continuity=True,xp=np)
            p=special.ndtr(-z)
            p*=2
            p=np.clip(p,0.,1.)
        target_mean=official._sample_mean(np.full(n,mu,dtype=self.dtype),is_log1p=True,exp_post_agg=True)
        reference_mean=self.reference_means[j]
        return {"target_mean":target_mean,"reference_mean":reference_mean,
                "percent_change":official._percent_change(target_mean,reference_mean),
                "fold_change":official._fold_change(target_mean,reference_mean,clip_value=20.),
                "p_value":p,"statistic":U1}, {"method":method,"control_less_count":less,
                "control_equal_count":equal,"analytic_float64_U":n*(less+.5*equal),
                "actual_U_minus_analytic":float(U1)-n*(less+.5*equal)}

    def raw_frame(self,means,n_cells,target):
        import pandas as pd
        import polars as pl
        means=np.asarray(means)
        if means.shape!=(len(self.genes),) or means.dtype!=self.dtype:
            raise ValueError("Constant encoded means must match reference gene axis and dtype")
        started=time.monotonic()
        rows,methods,errors=[],{},[]
        for j,gene in enumerate(self.genes):
            values,diag=self.gene(j,means[j],n_cells)
            rows.append(dict(target=str(target),reference="non-targeting",feature=gene,**values))
            methods[diag["method"]]=methods.get(diag["method"],0)+1
            errors.append(abs(diag["actual_U_minus_analytic"]))
        # Exactly the official pdex pandas -> Polars construction, retaining
        # the same scalar types and their mixed-column dtype promotion.
        frame=pl.DataFrame(pd.DataFrame(rows))
        return frame,{"version":VERSION,"dtype":str(self.dtype),"reference_sha256":self.reference_sha256,
              "reference_cells":self.n,"target_cells":int(n_cells),"gene_count":len(self.genes),
              "method_counts":methods,"maximum_U_difference_from_float64_formula":max(errors),
              "elapsed_seconds":time.monotonic()-started,"official_backend_default_changed":False,
              "official_score":False,"benchmark_comparable":False}
```
