## Supplementary reproducibility code and frozen predictions for "Centered cross-context perturbation transfer on the public 2025 Virtual Cell Challenge benchmark": mwu_constant_reference_pdex_v2.py

```
"""Frozen v1 SciPy specialization plus the actual pdex NaN-p postprocess.

The v1 module is retained unchanged. SciPy1.17.1 float32 all-tied variance can
round negative for some legitimate cell counts. pdex replaces the resulting
NaN p with one after dataframe construction; this wrapper does exactly that.
"""
from __future__ import annotations

import polars as pl
from mwu_constant_reference import ConstantReference as ScipyConstantReference

VERSION="constant_MWU_pdex0127_scipy1171_dtype_v2"


class ConstantReference(ScipyConstantReference):
    def raw_frame(self,means,n_cells,target):
        frame,diagnostics=super().raw_frame(means,n_cells,target)
        replacements=int(frame.select((pl.col("p_value").is_nan()|pl.col("p_value").is_null()).sum()).item())
        frame=frame.with_columns(pl.col("p_value").fill_nan(1.).fill_null(1.))
        diagnostics.update(version=VERSION,frozen_SciPy_core_version=diagnostics["version"],
             NaN_p_values_replaced_by_one=replacements,
             NaN_p_postprocessing="official pdex pandas p_value.fillna(1.0)")
        return frame,diagnostics
```
