## Supplementary reproducibility code and frozen predictions for "Centered cross-context perturbation transfer on the public 2025 Virtual Cell Challenge benchmark": score_training_fold.py

```
"""Full-panel training-only VCC diagnostics with a prediction-before-truth gate.

The official pdex backend is the default and sole implemented backend. Its raw
per-target tests may be reused across panels; adjusted p values never are.
Draft protocols permit metadata preparation, not real expression evaluation.
"""
from __future__ import annotations

import argparse
from datetime import datetime, timezone
import gc
import hashlib
import importlib.metadata
import json
import os
from pathlib import Path
import sys
import time

ROOT = Path("/data1/luanhaoyang/virtualcell")
CONTROL = "non-targeting"
METRICS = {"des": "overlap_at_N", "pds": "discrimination_score_l1", "mae": "mae"}
RAW_COLUMNS = ["target", "reference", "feature", "target_mean", "reference_mean",
               "percent_change", "fold_change", "p_value", "statistic"]


def now():
    return datetime.now(timezone.utc).isoformat()


def sha(path):
    digest = hashlib.sha256()
    with Path(path).open("rb") as stream:
        while block := stream.read(8 * 1024 * 1024):
            digest.update(block)
    return digest.hexdigest()


def canonical(value):
    return json.dumps(value, sort_keys=True, separators=(",", ":"), ensure_ascii=False, allow_nan=False)


def logical_sha(value):
    return hashlib.sha256(canonical(value).encode()).hexdigest()


def matrix_sha(matrix, block_rows=256):
    """Canonical C-order float32 values, independent of H5AD storage encoding."""
    import numpy as np
    from scipy import sparse
    digest = hashlib.sha256()
    for start in range(0, matrix.shape[0], block_rows):
        block = matrix[start:start+block_rows]
        if sparse.issparse(block):
            block = block.toarray()
        block = np.ascontiguousarray(block, dtype=np.float32)
        digest.update(block.tobytes(order="C"))
    return digest.hexdigest()


def selected_matrix_sha(matrix, indices, block_rows=256):
    import numpy as np
    from scipy import sparse
    digest = hashlib.sha256()
    for start in range(0,len(indices),block_rows):
        block = matrix[indices[start:start+block_rows], :]
        if sparse.issparse(block):
            block = block.toarray()
        digest.update(np.ascontiguousarray(block,dtype=np.float32).tobytes(order="C"))
    return digest.hexdigest()


def read_json(path):
    return json.loads(Path(path).read_text(encoding="utf-8-sig"))


def write_json(path, data):
    path = Path(path)
    path.parent.mkdir(exist_ok=True, parents=True)
    with path.open("x", encoding="utf-8") as stream:
        json.dump(data, stream, indent=2, sort_keys=True, allow_nan=False)


def protocol(path, execution=False):
    result = read_json(path)
    ev = result["evaluator"]
    required = {"cell_eval": "0.6.6", "pdex": "0.1.27", "profile": "vcc",
                "pert_col": "target_gene", "control_pert": CONTROL, "de_method": "wilcoxon",
                "allow_discrete": False, "USE_EXPERIMENTAL": "0", "backend": "official_pdex_0.1.27"}
    if any(ev.get(k) != v for k, v in required.items()) or ev["pdex_kwargs"] != {
            "tie_correct": True, "is_log1p": True, "exp_post_agg": True, "clip_value": 20.0}:
        raise ValueError("The official pinned training evaluator configuration changed")
    if result["normalization"]["target_sum"] != 54611.0 or result["normalization"]["additional_prediction_normalization"]:
        raise ValueError("Full-reference NTC-L protocol requires per-cell L54611 and no second prediction normalization")
    if any(result.get(k) for k in ("official_score", "benchmark_comparable", "champion_claim_supported", "validation_test_expression_access")):
        raise ValueError("This runner is only a training diagnostic")
    if Path(result["source"]["path"]) != ROOT / "data/train/adata_Training.h5ad":
        raise ValueError("Only the registered official training object is supported")
    if result["control_reference"]["policy"] != "all_training_NTC_in_original_source_row_order" or result["control_reference"]["expected_cells"] != 38176:
        raise ValueError("A subset cannot silently replace the full training NTC reference")
    if execution and (result["status"] != "frozen_by_root" or not result["real_DE_execution_authorized"] or result["renderer_selection"] is None):
        raise ValueError("Real scoring awaits a separately frozen root protocol and renderer")
    return result


def h5_column(group, key):
    """Read observation/variable metadata only; callers never inspect X here."""
    import h5py
    import numpy as np
    node = group[key]
    if isinstance(node, h5py.Dataset):
        values = node.asstr()[:] if node.dtype.kind in "OSU" else node[:]
    elif isinstance(node, h5py.Group) and "codes" in node and "categories" in node:
        categories = h5_column(node, "categories")
        codes = np.asarray(node["codes"][:], dtype=np.int64)
        if np.any(codes < 0) or np.any(codes >= len(categories)):
            raise ValueError("Metadata has missing or invalid categorical codes")
        values = np.asarray(categories)[codes]
    else:
        raise ValueError(f"Unsupported H5AD metadata column {key}")
    return np.asarray(values)


def h5_index(group):
    key = group.attrs.get("_index", "_index")
    key = key.decode() if isinstance(key, bytes) else str(key)
    return h5_column(group, key).astype(str)


def prepare_metadata(protocol_path, query_path, output):
    """Legitimate identities/counts/row IDs only, before prediction rendering."""
    import h5py
    import numpy as np
    cfg = protocol(protocol_path)
    query = read_json(query_path)
    targets, fit = list(map(str, query["query_targets"])), list(map(str, query["fit_targets"]))
    if not targets or len(set(targets)) != len(targets) or len(set(fit)) != len(fit) or set(targets) & set(fit) or CONTROL in set(targets) | set(fit):
        raise ValueError("Query/fit target identities must be unique, nonempty and disjoint")
    source = cfg["source"]
    inventory_path = Path(source["inventory_path"])
    if sha(inventory_path) != source["inventory_sha256"]:
        raise ValueError("Teacher metadata inventory changed")
    inventory = read_json(inventory_path)
    if inventory["source_sha256"] != source["sha256"] or inventory["normalization"]["target_sum"] != cfg["normalization"]["target_sum"]:
        raise ValueError("Inventory source or normalization differs")
    evidence_path = Path(cfg["normalization"]["scale_evidence_path"])
    if sha(evidence_path) != cfg["normalization"]["scale_evidence_sha256"]:
        raise ValueError("Training-control scale evidence changed")
    evidence = read_json(evidence_path)
    if evidence["NTC_raw_library"]["median"] != cfg["normalization"]["target_sum"]:
        raise ValueError("NTC-only median does not match the requested scale")
    with h5py.File(source["path"], "r") as handle:
        labels = h5_column(handle["obs"], "target_gene").astype(str)
        cell_ids, genes = h5_index(handle["obs"]), h5_index(handle["var"])
        # Deliberately no X, layers, numeric expression slices, or query stats.
    if len(labels) != source["expected_cells"] or len(genes) != source["expected_genes"] or len(set(genes)) != len(genes) or len(set(cell_ids)) != len(cell_ids):
        raise ValueError("Training metadata axes differ from the registered identity")
    all_targets = set(labels) - {CONTROL}
    if len(all_targets) != source["expected_targets"] or set(targets) | set(fit) != all_targets:
        raise ValueError("Query/fit split must cover exactly the official 150 training targets")
    control_rows = np.flatnonzero(labels == CONTROL).astype(int).tolist()
    if len(control_rows) != cfg["control_reference"]["expected_cells"]:
        raise ValueError("Full NTC reference count differs")
    target_rows = {target: np.flatnonzero(labels == target).astype(int).tolist() for target in targets}
    counts = {target: len(rows) for target, rows in target_rows.items()}
    if any(counts[t] != inventory["target_counts"][t] or counts[t] <= 0 for t in targets):
        raise ValueError("Actual query target counts disagree with the inventory")
    result = {"schema_version": 1, "panel_id": query["panel_id"], "registered_at_utc": now(),
              "source": source, "normalization": cfg["normalization"], "query_targets": targets, "fit_targets": fit,
              "target_counts": counts, "target_source_rows": target_rows,
              "target_source_rows_sha256": {t: logical_sha(rows) for t, rows in target_rows.items()},
              "control_source_rows": control_rows, "control_source_rows_sha256": logical_sha(control_rows),
              "control_cell_ids": cell_ids[control_rows].tolist(), "control_cell_ids_sha256": logical_sha(cell_ids[control_rows].tolist()),
              "genes": genes.tolist(), "gene_order_sha256": logical_sha(genes.tolist()),
              "protocol_sha256_at_metadata_preparation": sha(protocol_path), "query_manifest_sha256": sha(query_path),
              "query_X_values_accessed": False, "contains_expression_or_outcome_statistics": False,
              "official_score": False}
    write_json(output, result)
    return {"metadata": str(output), "sha256": sha(output), "panel_id": result["panel_id"], "target_counts": counts,
            "control_source_rows_sha256": result["control_source_rows_sha256"], "control_cell_ids_sha256": result["control_cell_ids_sha256"],
            "query_X_values_accessed": False}


def parse_records(items):
    records = []
    for item in items:
        role, value = item.split("=", 1)
        path = Path(value).resolve(strict=True)
        if not role or not path.is_file() or role in {r["role"] for r in records}:
            raise ValueError("Unique role=regular_file records are required")
        records.append({"role": role, "path": str(path), "sha256": sha(path)})
    return records


def check_prediction_metadata(adata, panel):
    import numpy as np
    if not adata.obs_names.is_unique or adata.var_names.astype(str).tolist() != panel["genes"] or "target_gene" not in adata.obs:
        raise ValueError("Prediction gene/target identities differ from the registered panel")
    labels = adata.obs.target_gene.astype(str).to_numpy()
    counts = {str(t): int(n) for t, n in adata.obs.target_gene.astype(str).value_counts().items() if n > 0}
    expected = dict(panel["target_counts"], **{CONTROL: len(panel["control_source_rows"])})
    if counts != expected:
        raise ValueError("Prediction counts must equal actual registered query counts and full NTC count")
    controls = np.flatnonzero(labels == CONTROL)
    if adata.obs_names.astype(str).to_numpy()[controls].tolist() != panel["control_cell_ids"]:
        raise ValueError("Prediction controls must carry actual source cell IDs in registered order")
    if "source_row_id" not in adata.obs or adata.obs.iloc[controls].source_row_id.astype(int).tolist() != panel["control_source_rows"]:
        raise ValueError("Prediction control source-row identities/order differ")
    return labels, controls


def commit(protocol_path, panel_path, predictions, artifacts, output):
    """Hash and validate every prediction arm before any query truth is opened."""
    import anndata as ad
    cfg, panel = protocol(protocol_path, execution=True), read_json(panel_path)
    if panel["source"] != cfg["source"] or panel["normalization"] != cfg["normalization"] or panel["query_X_values_accessed"]:
        raise ValueError("Panel provenance differs from the frozen protocol")
    for key in ("control_source_rows_sha256", "control_cell_ids_sha256"):
        declared = cfg["control_reference"][key]
        if declared is None or declared != panel[key]:
            raise ValueError(f"Root protocol must freeze the actual {key}")
    arms, artifact_records = parse_records(predictions), parse_records(artifacts)
    if not arms or set(cfg["required_artifact_roles"]) != {r["role"] for r in artifact_records}:
        raise ValueError("Every required model/cache/renderer/settings/knowledge artifact must be frozen")
    if any(Path(record["path"]) == Path(cfg["source"]["path"]) for record in [*arms,*artifact_records]):
        raise ValueError("Raw query truth cannot be supplied as a pre-truth prediction/model artifact")
    control_hashes = []
    for record in arms:
        data = ad.read_h5ad(record["path"], backed="r")
        try:
            _, indices = check_prediction_metadata(data, panel)
            digest = selected_matrix_sha(data.X,indices)
            record["control_values_sha256"] = digest
            control_hashes.append(digest)
        finally:
            data.file.close()
    if len(set(control_hashes)) != 1 or cfg["control_reference"]["normalized_control_values_sha256"] != control_hashes[0]:
        raise ValueError("Every arm must use the same actually registered full NTC values")
    manifest = {"schema_version": 1, "committed_at_utc": now(), "scorer_sha256": sha(Path(__file__)),
                "protocol": {"path": str(Path(protocol_path).resolve()), "sha256": sha(protocol_path)},
                "panel": {"path": str(Path(panel_path).resolve()), "sha256": sha(panel_path)},
                "predictions": arms, "artifacts": artifact_records,
                "query_truth_not_yet_opened_by_this_scorer": True, "official_score": False}
    write_json(output, manifest)
    return manifest


def verify_commit(path):
    manifest = read_json(path)
    if manifest.get("schema_version") != 1 or not manifest.get("query_truth_not_yet_opened_by_this_scorer") or manifest["scorer_sha256"] != sha(Path(__file__)):
        raise ValueError("Prediction commit or scorer identity is invalid")
    timestamp = datetime.fromisoformat(manifest["committed_at_utc"])
    if timestamp.tzinfo is None or timestamp > datetime.now(timezone.utc):
        raise ValueError("Prediction commit has an invalid timestamp")
    for record in [manifest["protocol"], manifest["panel"], *manifest["predictions"], *manifest["artifacts"]]:
        if sha(record["path"]) != record["sha256"]:
            raise ValueError(f"Artifact changed after prediction commit: {record['path']}")
    cfg = protocol(manifest["protocol"]["path"], execution=True)
    if {r["role"] for r in manifest["artifacts"]} != set(cfg["required_artifact_roles"]) or not manifest["predictions"]:
        raise ValueError("Committed model/cache/renderer/settings/knowledge artifacts are incomplete")
    if len(set(r["role"] for r in manifest["predictions"])) != len(manifest["predictions"]):
        raise ValueError("Prediction arm names must be unique")
    return manifest


def normalize_teacher_counts(raw, target_sum):
    """Match pinned prepare_training.py arithmetic, then encode cell float32."""
    import numpy as np
    from scipy import sparse
    if sparse.issparse(raw):
        raw = raw.tocsr().astype(np.float32)
        payload, library = raw.data, np.asarray(raw.sum(axis=1)).ravel()
    else:
        raw = np.asarray(raw, dtype=np.float32)
        payload, library = raw, raw.sum(axis=1, dtype=np.float64)
    if not np.isfinite(payload).all() or np.any(payload < 0) or np.any(np.abs(payload-np.rint(payload)) > 1e-4):
        raise ValueError("Query input must be finite nonnegative integer raw counts")
    scale = float(target_sum) / np.maximum(library, 1.0)
    if sparse.issparse(raw):
        normal = raw.multiply(scale[:, None]).tocsr()
        normal.data = np.log1p(normal.data)
        normal = normal.toarray()
    else:
        normal = np.log1p(raw * scale[:, None])
    return np.asarray(normal, dtype=np.float32)


def check_log_matrix(matrix, chunk_rows=256):
    import numpy as np
    from scipy import sparse
    found_fractional = False
    for start in range(0, matrix.shape[0], chunk_rows):
        block = matrix[start:start+chunk_rows]
        values = block.data if sparse.issparse(block) else np.asarray(block)
        if not np.isfinite(values).all() or np.any(values < 0) or np.any(values >= 15):
            raise ValueError("Scoring input must be finite nonnegative log1p float32 values below 15")
        found_fractional |= bool(np.any(values != np.floor(values)))
    if not found_fractional:
        raise ValueError("Entire input is integer-valued and ambiguous to the official conversion heuristic")


def runtime():
    os.environ["USE_EXPERIMENTAL"] = "0"
    os.environ["CUDA_VISIBLE_DEVICES"] = ""
    if "pdex._single_cell" in sys.modules and sys.modules["pdex._single_cell"].use_experimental:
        raise ValueError("pdex was previously imported in experimental mode")
    versions = {k: importlib.metadata.version(k) for k in ["cell-eval", "pdex", "scipy", "numpy", "polars"]}
    if versions["cell-eval"] != "0.6.6" or versions["pdex"] != "0.1.27":
        raise ValueError("Pinned evaluator versions required")
    import pdex._single_cell as backend
    if backend.use_experimental:
        raise ValueError("Official nonexperimental pdex backend required")
    return dict(versions=versions, pdex_backend_sha256=sha(backend.__file__), scorer_sha256=sha(Path(__file__)))


def with_global_bh(raw_frames, targets, genes):
    """Never accept an already adjusted per-target cache as a panel FDR."""
    import numpy as np
    import polars as pl
    from scipy.stats import false_discovery_control
    if len(raw_frames) != len(targets):
        raise ValueError("Every query target requires one full-gene raw table")
    for frame, target in zip(raw_frames, targets):
        if set(frame.columns) != set(RAW_COLUMNS) or frame.height != len(genes):
            raise ValueError("Raw cache must contain exactly original raw fields and the entire gene panel")
        if frame["feature"].to_list() != genes or frame["target"].unique().to_list() != [target] or frame["reference"].unique().to_list() != [CONTROL]:
            raise ValueError("Raw DE cache target/reference/gene identity mismatch")
    combined = pl.concat(raw_frames, how="vertical")
    pvalues = combined["p_value"].to_numpy()
    if not np.isfinite(pvalues).all() or np.any(pvalues < 0) or np.any(pvalues > 1):
        raise ValueError("Raw DE has invalid p values")
    return combined.with_columns(pl.Series("fdr", false_discovery_control(pvalues, method="bh")))


def raw_de(target_matrix, control_matrix, target, genes, settings, identity, cache):
    import anndata as ad
    import numpy as np
    import pandas as pd
    import polars as pl
    import pdex._single_cell as backend
    key = logical_sha(identity)
    directory = Path(cache) / key
    if directory.exists():
        manifest = read_json(directory / "manifest.json")
        if manifest["identity"] != identity or sha(directory / "raw.parquet") != manifest["raw_sha256"]:
            raise ValueError("Existing raw-DE cache failed identity/hash validation")
        frame = pl.read_parquet(directory / "raw.parquet")
        if "fdr" in frame.columns:
            raise ValueError("Raw-DE cache may not retain cross-panel adjusted p values")
        return frame, dict(key=key, hit=True, manifest_sha256=sha(directory / "manifest.json"))
    directory.mkdir(exist_ok=False, parents=True)
    started = time.monotonic()
    matrix = np.concatenate([control_matrix, target_matrix]).astype(np.float32, copy=False)
    data = ad.AnnData(matrix, obs=pd.DataFrame({"target_gene": [CONTROL]*len(control_matrix)+[target]*len(target_matrix)}),
                      var=pd.DataFrame(index=genes))
    frame = backend.parallel_differential_expression(data, reference=CONTROL, groupby_key="target_gene",
            num_workers=settings["num_workers"], batch_size=settings["batch_size"], metric="wilcoxon", as_polars=True,
            **settings["pdex_kwargs"]).select(RAW_COLUMNS)
    frame.write_parquet(directory / "raw.parquet")
    manifest = {"identity": identity, "raw_sha256": sha(directory / "raw.parquet"), "created_at_utc": now(),
                "elapsed_seconds": time.monotonic()-started, "rows": frame.height, "fdr_cached": False,
                "backend": "official_pdex_0.1.27", "official_score": False}
    write_json(directory / "manifest.json", manifest)
    return frame, dict(key=key, hit=False, manifest_sha256=sha(directory / "manifest.json")),


def resource_estimate(panel, cfg):
    cells = len(panel["control_source_rows"]) + sum(panel["target_counts"].values())
    dense = cells * len(panel["genes"]) * 4
    largest_pair = (len(panel["control_source_rows"])+max(panel["target_counts"].values()))*len(panel["genes"])*4
    # Conservative budget: real/pred arrays, Polars column copies and temporaries,
    # plus one target/control concat and official pdex shared-memory copy.
    ram = 4*dense + 2*largest_pair + 4*1024**3
    return {"panel_cells_per_object": cells, "dense_float32_bytes_per_object": dense,
            "largest_pdex_shared_memory_bytes": largest_pair, "conservative_RAM_bytes": ram,
            "full_BH_family": len(panel["query_targets"])*len(panel["genes"]),
            "CPU_workers": cfg["evaluator"]["num_workers"], "GPU_use": False,
            "disk_minimum_bytes_excluding_input_predictions": dense+2*largest_pair,
            "estimate_is_not_measured_peak": True}


def evaluate(commit_path, cache, output):
    """Only this entry opens real expression, strictly after all-arm commit."""
    committed = verify_commit(commit_path)
    cfg = protocol(committed["protocol"]["path"], execution=True)
    panel = read_json(committed["panel"]["path"])
    output = Path(output)
    if output.exists():
        raise FileExistsError("Use a fresh output directory; never overwrite fold evaluation")
    output.mkdir(parents=True)
    if os.name != "posix" or not output.resolve().is_relative_to(ROOT / "runs") or not Path(cache).resolve().is_relative_to(ROOT):
        raise ValueError("Real training scoring runs only in the authorized remote project")
    estimate = resource_estimate(panel, cfg)
    if estimate["conservative_RAM_bytes"] > cfg["resources"]["maximum_memory_gib"]*1024**3:
        raise MemoryError("Panel exceeds the frozen conservative memory budget")
    memory = {line.split(":")[0]: int(line.split()[1])*1024 for line in Path("/proc/meminfo").read_text().splitlines() if line.startswith(("MemAvailable:", "MemTotal:"))}
    if estimate["conservative_RAM_bytes"] > memory["MemAvailable"]*(1-cfg["resources"]["minimum_free_memory_fraction"]):
        raise MemoryError("Available memory does not leave the required reserve for other users")
    shm = os.statvfs("/dev/shm")
    if estimate["largest_pdex_shared_memory_bytes"] > shm.f_bavail*shm.f_frsize*.8:
        raise MemoryError("Official pdex shared-memory allocation exceeds safe available space")
    write_json(output / "before_query_truth.json", {"at_utc": now(), "prediction_commit_sha256": sha(commit_path),
                "all_prediction_hashes_verified": True, "resource_estimate": estimate,
                "scorer_sha256": sha(Path(__file__)), "ground_truth_not_yet_opened": True, "official_score": False})
    versions = runtime()
    # The complete source byte-hash is now permitted: prediction commit already
    # passed. Metadata preparation above never hashes or inspects query X.
    if sha(cfg["source"]["path"]) != cfg["source"]["sha256"]:
        raise ValueError("Raw training source byte identity changed")
    import anndata as ad
    import numpy as np
    import pandas as pd
    from cell_eval import MetricsEvaluator
    genes, targets = panel["genes"], panel["query_targets"]
    rows = panel["control_source_rows"] + [r for t in targets for r in panel["target_source_rows"][t]]
    labels = [CONTROL]*len(panel["control_source_rows"]) + [t for t in targets for _ in panel["target_source_rows"][t]]
    truth_path = output / "real_normalized_cells.npy"
    matrix = np.lib.format.open_memmap(truth_path, mode="w+", dtype=np.float32, shape=(len(rows),len(genes)))
    source = ad.read_h5ad(cfg["source"]["path"], backed="r")
    opened_at = now()
    try:
        if source.var_names.astype(str).tolist() != genes:
            raise ValueError("Raw training gene order changed")
        actual_labels = source.obs.target_gene.astype(str).to_numpy()
        if actual_labels[rows].tolist() != labels or source.obs_names.astype(str).to_numpy()[panel["control_source_rows"]].tolist() != panel["control_cell_ids"]:
            raise ValueError("Raw training target/control row identities changed")
        chunk = cfg["resources"]["normalization_chunk_rows"]
        for start in range(0,len(rows),chunk):
            matrix[start:start+chunk] = normalize_teacher_counts(source.X[rows[start:start+chunk], :], cfg["normalization"]["target_sum"])
        matrix.flush()
    finally:
        source.file.close()
    check_log_matrix(matrix)
    control = matrix[:len(panel["control_source_rows"])]
    control_hash = matrix_sha(control)
    if control_hash != cfg["control_reference"]["normalized_control_values_sha256"]:
        raise ValueError("Actual raw-normalized full NTC reference differs from frozen prediction controls")
    obs = pd.DataFrame({"target_gene": labels}, index=[f"TRAIN_REAL_{r}" for r in rows])
    real = ad.AnnData(matrix, obs=obs, var=pd.DataFrame(index=genes))
    common = {"runtime": versions, "normalization": cfg["normalization"], "gene_order_sha256": panel["gene_order_sha256"],
              "control_source_rows_sha256": panel["control_source_rows_sha256"], "control_values_sha256": control_hash}
    raw_real, real_cache = [], {}
    start = len(control)
    for target in targets:
        size = panel["target_counts"][target]
        target_matrix = matrix[start:start+size]
        identity = dict(common, kind="real", target=target, count=size, source_sha256=cfg["source"]["sha256"],
                        target_row_ids_sha256=panel["target_source_rows_sha256"][target], target_values_sha256=matrix_sha(target_matrix))
        frame, receipt = raw_de(target_matrix, control, target, genes, cfg["evaluator"], identity, cache)
        raw_real.append(frame)
        real_cache[target] = receipt
        start += size
    de_real = with_global_bh(raw_real, targets, genes)
    de_real.write_parquet(output / "real_full_panel_de.parquet")
    de_real.write_csv(output / "real_full_panel_de.csv")
    summary = {"panel_id": panel["panel_id"], "query_truth_opened_at_utc": opened_at, "query_targets": targets,
               "actual_target_counts": panel["target_counts"], "control_values_sha256": control_hash, "control_count": len(control),
               "normalization": cfg["normalization"], "runtime": versions, "real_raw_cache": real_cache,
               "full_BH_family": len(targets)*len(genes), "resource_estimate": estimate, "arms": {},
               "prediction_commit_sha256": sha(commit_path), "official_score": False,
               "benchmark_comparable": False, "champion_claim_supported": False, "validation_test_expression_accessed": False}
    for record in committed["predictions"]:
        # Whole files are immutable and were hashed before any query access.
        arm = record["role"]
        arm_output = output / arm
        arm_output.mkdir()
        predicted = ad.read_h5ad(record["path"])
        pred_labels, pred_control_rows = check_prediction_metadata(predicted, panel)
        check_log_matrix(predicted.X)
        if selected_matrix_sha(predicted.X,pred_control_rows) != control_hash:
            raise ValueError("Prediction reference differs from actual full NTC values")
        raw_frames, receipts = [], {}
        for target in targets:
            x = predicted.X[np.flatnonzero(pred_labels == target)]
            if hasattr(x,"toarray"):
                x = x.toarray()
            identity = dict(common, kind="pred", target=target, count=len(x), target_values_sha256=matrix_sha(x))
            frame, receipt = raw_de(x, control, target, genes, cfg["evaluator"], identity, cache)
            raw_frames.append(frame)
            receipts[target] = receipt
            del x
        de_pred = with_global_bh(raw_frames, targets, genes)
        de_pred.write_parquet(arm_output / "pred_full_panel_de.parquet")
        de_pred.write_csv(arm_output / "pred_full_panel_de.csv")
        evaluator = MetricsEvaluator(adata_pred=predicted, adata_real=real, de_pred=de_pred, de_real=de_real,
                control_pert=CONTROL, pert_col="target_gene", de_method="wilcoxon", allow_discrete=False,
                num_threads=cfg["evaluator"]["num_workers"], batch_size=cfg["evaluator"]["batch_size"],
                outdir=str(arm_output/"vcc"), pdex_kwargs=cfg["evaluator"]["pdex_kwargs"].copy())
        result, aggregate = evaluator.compute(profile="vcc", break_on_error=True)
        if set(result.columns) != {"perturbation", *METRICS.values()} or set(result["perturbation"].to_list()) != set(targets):
            raise ValueError("Official metrics did not cover the exact entire query panel")
        mean = aggregate.filter(aggregate["statistic"] == "mean")
        raw_metrics = {k: float(mean[v][0]) for k,v in METRICS.items()}
        if not all(np.isfinite(value) for value in raw_metrics.values()):
            raise ArithmeticError("VCC diagnostics include nonfinite raw metrics")
        anchors = cfg["historical_anchors"]
        components = {k: max(0., 1-raw_metrics[k]/anchors[k] if k == "mae" else (raw_metrics[k]-anchors[k])/(1-anchors[k])) for k in METRICS}
        summary["arms"][arm] = {"prediction_sha256": record["sha256"], "raw_metrics": raw_metrics,
             "historical_anchor_normalization_diagnostic": {"components": components, "mean_fraction": sum(components.values())/3,
                   "anchor_source_type": anchors["source_type"], "historical_comparison_supported": False},
             "raw_cache": receipts, "pred_FDR_discoveries": int((de_pred["fdr"] < .05).sum()),
             "real_FDR_discoveries": int((de_real["fdr"] < .05).sum())}
        del evaluator, predicted, de_pred, raw_frames
        gc.collect()
    summary["completed_at_utc"] = now()
    summary["actual_truth_values_file_sha256"] = sha(truth_path)
    write_json(output / "summary.json", summary)
    hashes = {str(p.relative_to(output)): sha(p) for p in sorted(output.rglob("*")) if p.is_file()}
    write_json(output / "completion.json", dict(exit_code=0, completed_at_utc=now(), output_hashes=hashes,
          cache_root=str(cache), prediction_commit_sha256=sha(commit_path), official_score=False))
    return summary


def main():
    parser = argparse.ArgumentParser(description=__doc__)
    subs = parser.add_subparsers(dest="command", required=True)
    meta = subs.add_parser("metadata")
    for name in ("protocol", "queries", "output"):
        meta.add_argument(f"--{name}", type=Path, required=True)
    freeze = subs.add_parser("commit")
    for name in ("protocol", "panel", "output"):
        freeze.add_argument(f"--{name}", type=Path, required=True)
    freeze.add_argument("--prediction", action="append", required=True)
    freeze.add_argument("--artifact", action="append", required=True)
    run = subs.add_parser("run")
    for name in ("commit", "cache", "outdir"):
        run.add_argument(f"--{name}", type=Path, required=True)
    args = parser.parse_args()
    if args.command == "metadata":
        result = prepare_metadata(args.protocol,args.queries,args.output)
    elif args.command == "commit":
        result = commit(args.protocol,args.panel,args.prediction,args.artifact,args.output)
    else:
        result = evaluate(args.commit,args.cache,args.outdir)
    print(json.dumps(result,indent=2,allow_nan=False))


if __name__ == "__main__":
    main()
```
