## Supplementary reproducibility code and frozen predictions for "Centered cross-context perturbation transfer on the public 2025 Virtual Cell Challenge benchmark": score_training_meanflat_DES.py

```
"""Six fixed post-screen TRAINING DES diagnostics; no validation/test reads.

Preparation materializes and commits every native-count H5AD before this
runner reads query raw-stat tables. The specialized backend is restricted to
constant float32 targets and does not change the official default scorer.
"""
from __future__ import annotations

import argparse
from concurrent.futures import ThreadPoolExecutor
import gc
import hashlib
import importlib.metadata
import inspect
import json
import os
from pathlib import Path
import sys
import time

sys.path.insert(0, str(Path(__file__).resolve().parent))
import score_training_fold as score

ROOT = score.ROOT
CAP = 14.999999046325684
ARMS = ["candidate_02_learned_only", "candidate_02_genuine_sol_mix25",
        "candidate_02_static_retrieval_mix25", "candidate_02_shuffled_mix25",
        "candidate_02_random_mix25", "GEMZ3_source_mean"]
GEMZ_EXPERTS = ["K562_gwps_normalized_bulk_01", "rpe1_normalized_bulk_01",
                "K562_essential_normalized_bulk_01"]
FLAGS = dict(official_score=False, benchmark_comparable=False,
             champion_claim_supported=False, unbiased_new_outer_result=False,
             validation_test_expression_accessed=False, DES_feedback_to_agent=False)


def record(path, role=None):
    path = Path(path).resolve(strict=True)
    return dict(path=str(path), sha256=score.sha(path), **({"role": role} if role else {}))


def verify_record(item):
    path = Path(item["path"]).resolve(strict=True)
    if not path.is_relative_to(ROOT) or score.sha(path) != item["sha256"]:
        raise ValueError("Frozen project artifact changed or escaped remote root")
    return path


def configuration(path, execution=False):
    cfg = score.read_json(path)
    if cfg["arms"] != ARMS or cfg["family_size"] != 34*18080 or cfg["fit_targets"] != 66 or cfg["query_targets"] != 34:
        raise ValueError("The root-selected six-arm inner66/query34 scope changed")
    if cfg["normalization_target_sum"] != 54611. or cfg["control_cells"] != 38176 or cfg["cap"] != CAP:
        raise ValueError("Normalization/reference/encoding contract changed")
    if cfg["renderer"] != "native_actual_count_mean_flat_float32" or cfg["additional_normalization"]:
        raise ValueError("Only the frozen constant-target renderer is supported")
    if any(cfg.get(k) is not v for k, v in FLAGS.items()) or cfg["whole_train_BH"] or cfg["candidate_selection_uses_DES"]:
        raise ValueError("Only post-screen training diagnostics without feedback are supported")
    if cfg["backend"] != "constant_MWU_pdex0127_scipy1171_dtype_v2" or cfg["workers"] not in range(1,5):
        raise ValueError("Audited constant backend and at most four target workers required")
    if execution and (cfg["status"] != "frozen_by_root" or cfg["real_DE_execution_authorized"] is not True):
        raise ValueError("Real six-arm execution awaits root approval of code and QA")
    return cfg


def query_panel(scope, panels):
    """Reuse only already registered observation identities and cell counts."""
    from read_fit_raw_DE import fit_scope
    fit, query = fit_scope(scope, panels[scope["outer_fold"]])
    base = panels[scope["outer_fold"]]
    for panel in panels:
        for key in ["genes", "source", "normalization", "control_source_rows", "control_cell_ids"]:
            if panel[key] != base[key] or panel["query_X_values_accessed"]:
                raise ValueError("Original metadata panels disagree or contain query expressions")
    rows = {t: v for p in panels for t, v in p["target_source_rows"].items()}
    counts = {t: v for p in panels for t, v in p["target_counts"].items()}
    if len(rows) != 150 or set(query)&set(scope["excluded_outer_query_targets"]):
        raise ValueError("Registered training target coverage or outer exclusion differs")
    result = dict(base, panel_id="outer0_inner0_postscreen_query34", fit_targets=fit,
                  query_targets=query, target_counts={t: counts[t] for t in query},
                  target_source_rows={t: rows[t] for t in query},
                  target_source_rows_sha256={t: score.logical_sha(rows[t]) for t in query})
    if len(fit) != 66 or len(query) != 34 or len(result["genes"]) != 18080 or min(result["target_counts"].values()) <= 8:
        raise ValueError("Only the actual large-count registered training panel is supported")
    return result


def gemz3_mean(metadata, control, common, deltas, available):
    """Exactly the already frozen diagnostic's three-source available mean."""
    import numpy as np
    source_ids = [metadata["expert_ids"].index(k) for k in GEMZ_EXPERTS]
    source, mask = deltas[:, source_ids, :], available[:, source_ids, :]
    count = mask.sum(axis=1)
    delta = np.divide(np.where(mask, source, 0).sum(axis=1), count,
                      out=np.broadcast_to(common, count.shape).copy(), where=count > 0)
    return np.maximum(control + delta, 0)


def resource_preflight(cfg, panel):
    """Conservative shared-CDF budget; no dense query truth is allocated."""
    control_bytes = 38176*18080*4
    estimated = control_bytes*8 + 8*1024**3
    mem = {line.split(":")[0]: int(line.split()[1])*1024 for line in Path("/proc/meminfo").read_text().splitlines()
           if line.startswith(("MemAvailable:", "MemTotal:"))}
    if estimated > cfg["maximum_memory_gib"]*1024**3 or estimated > mem["MemAvailable"]*.8:
        raise MemoryError("Shared reference preparation cannot preserve the memory reserve")
    disk = os.statvfs(ROOT)
    dense = (38176+sum(panel["target_counts"].values()))*18080*4
    required_disk = dense + 6*(control_bytes + 1024**3) + 2*1024**3
    if disk.f_bavail*disk.f_frsize < required_disk:
        raise OSError("Insufficient space for all six retained H5ADs and one writer scratch")
    return dict(memory=mem, conservative_peak_RAM_bytes=estimated,
                prediction_dense_scratch_bytes=dense, minimum_disk_bytes=required_disk,
                CPU_affinity=sorted(os.sched_getaffinity(0)), workers=cfg["workers"], GPU_use=False,
                estimate_is_not_measured_peak=True)


def pinned_runtime():
    runtime = score.runtime()
    expected = {"cell-eval": "0.6.6", "pdex": "0.1.27", "scipy": "1.17.1", "numpy": "2.4.6", "polars": "2.0.0"}
    if runtime["versions"] != expected:
        raise ValueError("The audited raw-stat and metric runtime changed")
    from cell_eval.metrics._de import de_overlap_metric
    from cell_eval._types._de import initialize_de_comparison
    import scipy.stats._mannwhitneyu as mwu
    return dict(runtime=runtime, direct_DES_metric_source_sha256=score.sha(inspect.getfile(de_overlap_metric)),
                DE_comparison_source_sha256=score.sha(inspect.getfile(initialize_de_comparison)),
                scipy_MWU_source_sha256=score.sha(mwu.__file__))


def load_inputs(cfg):
    paths = {r["role"]: verify_record(r) for r in cfg["inputs"]}
    panels = [score.read_json(verify_record(r)) for r in cfg["metadata_panels"]]
    scope = score.read_json(paths["scope"])
    panel = query_panel(scope, panels)
    semantic = score.read_json(paths["semantic_commit"])
    if semantic["prediction_sha256"] != score.sha(paths["semantic_predictions"]) or semantic["query_truth_opened"] or semantic["validation_test_expression_opened"]:
        raise ValueError("Original semantic predictions were not frozen before query truth")
    if not set(ARMS[:-1]).issubset(semantic["prediction_arms"]) or semantic["cache_sha256"] != score.sha(paths["prediction_agent_cache"]):
        raise ValueError("Frozen selected arms or genuine prediction-agent cache differ")
    return paths, panel, semantic


def prepare(protocol_path, run_dir):
    """All actual prediction H5ADs are frozen before any query cache read."""
    import anndata as ad
    import numpy as np
    from write_training_fold_flat import write_one
    cfg = configuration(protocol_path, execution=True)
    run_dir = run_dir.resolve()
    if run_dir != Path(cfg["run_dir"]).resolve() or not run_dir.is_relative_to(ROOT/"runs") or run_dir.exists():
        raise ValueError("Fresh explicit registered run directory required")
    os.sched_setaffinity(0, cfg["CPU_affinity"])
    paths, panel, semantic = load_inputs(cfg)
    runtime = pinned_runtime()
    estimate = resource_preflight(cfg, panel)
    control_manifest = score.read_json(paths["control_manifest"])
    if control_manifest["state"] != "complete" or control_manifest["query_expression_read"] or control_manifest["target_sum"] != 54611.:
        raise ValueError("The real full-training NTC provenance is invalid")
    if control_manifest["normalized_control_values_sha256"] != cfg["control_values_sha256"]:
        raise ValueError("Full-cell reference values differ from registration")
    with np.load(paths["semantic_predictions"], allow_pickle=False) as source:
        if source["targets"].astype(str).tolist() != panel["query_targets"] or source["genes"].astype(str).tolist() != panel["genes"]:
            raise ValueError("Semantic prediction axes differ from the exact inner panel")
        means = {arm: source[arm].copy() for arm in ARMS[:-1]}
    with np.load(paths["stage2_dataset"], allow_pickle=False) as source:
        metadata = json.loads(str(source["metadata"]))
        if metadata["fit_targets"] != panel["fit_targets"] or metadata["query_targets"] != panel["query_targets"] or metadata["gene_names"] != panel["genes"]:
            raise ValueError("GEMZ derivation inputs differ from frozen fit/query/gene axes")
        # These four arrays are prediction inputs. Do not load fit labels,
        # responses, current-fit FDR, raw p values or query truth arrays.
        means[ARMS[-1]] = gemz3_mean(metadata, source["control_mean"], source["common_delta"],
                                   source["query_expert_deltas"], source["query_expert_available"])
    run_dir.mkdir(parents=True)
    score.write_json(run_dir/"panel.json", panel)
    score.write_json(run_dir/"before_prediction_encoding.json", dict(at_utc=score.now(),
        protocol_sha256=score.sha(protocol_path), runtime=runtime, resources=estimate,
        original_semantic_commit_sha256=score.sha(paths["semantic_commit"]),
        post_screen_selection=True, this_runner_query_cache_opened=False, **FLAGS))
    encoded = {arm: np.clip(means[arm], 0., CAP).astype(np.float32) for arm in ARMS}
    np.savez_compressed(run_dir/"encoded_means.npz", targets=np.asarray(panel["query_targets"]),
                        genes=np.asarray(panel["genes"]), **encoded)
    predictions = []
    controls = ad.read_h5ad(paths["controls"], backed="r")
    try:
        if controls.shape != (38176,18080) or controls.var_names.astype(str).tolist() != panel["genes"] or controls.obs_names.astype(str).tolist() != panel["control_cell_ids"]:
            raise ValueError("Full reference axes or actual original cell identities changed")
        if controls.obs.source_row_id.astype(int).tolist() != panel["control_source_rows"] or score.matrix_sha(controls.X) != cfg["control_values_sha256"]:
            raise ValueError("Full reference rows/order/values changed")
        for arm in ARMS:
            print(f"prepare actual native-count H5AD {arm}", flush=True)
            receipt = write_one(means[arm], panel["genes"], panel, controls,
                                run_dir/f"{arm}.h5ad", run_dir/f".{arm}.scratch.npy")
            if receipt["control_values_sha256"] != cfg["control_values_sha256"]:
                raise ValueError("Materialized prediction control values changed")
            predictions.append(dict(receipt, role=arm))
    finally:
        controls.file.close()
    commit = dict(state="all_six_materialized_predictions_committed", at_utc=score.now(),
                  protocol=record(protocol_path), panel=record(run_dir/"panel.json"),
                  encoded_means=record(run_dir/"encoded_means.npz"), predictions=predictions,
                  inputs=cfg["inputs"], metadata_panels=cfg["metadata_panels"], runtime=runtime,
                  runner_sha256=score.sha(__file__), this_runner_query_cache_opened=False,
                  previous_query_MAE_PDS_used_for_arm_screening=True, **FLAGS)
    score.write_json(run_dir/"all_prediction_commit.json", commit)
    return commit


def official_DES(real, pred, targets):
    """Invoke the actual registered overlap-at-N function, including FC ranks."""
    import numpy as np
    from cell_eval._types._de import initialize_de_comparison
    from cell_eval.metrics._de import de_overlap_metric
    from cell_eval._types._de import DESortBy
    values = de_overlap_metric(initialize_de_comparison(real, pred), k=None, metric="overlap",
                               fdr_threshold=.05, sort_by=DESortBy.ABS_FOLD_CHANGE)
    if set(values) != set(targets) or not all(np.isfinite(v) for v in values.values()):
        raise ArithmeticError("Official DES did not return the exact finite query target set")
    return {t: float(values[t]) for t in targets}


def read_real_raw(cfg, panel):
    """Strict registered target cache resolver; numerical reads only query34."""
    import numpy as np
    import polars as pl
    ledger_record = next(r for r in cfg["inputs"] if r["role"] == "real_cache_ledger")
    ledger = score.read_json(verify_record(ledger_record))
    common = ledger["common_identity"]
    if ledger["scope"] != "registered_official_training_real_raw_cache" or ledger["NaN_p_postprocessed"] is not True or common["runtime"] != score.runtime():
        raise ValueError("Original official raw cache producer/runtime differs")
    expected_common = dict(normalization=panel["normalization"], gene_order_sha256=panel["gene_order_sha256"],
                           control_source_rows_sha256=panel["control_source_rows_sha256"], control_values_sha256=cfg["control_values_sha256"])
    if any(common[k] != v for k,v in expected_common.items()):
        raise ValueError("Original raw cache control/value/gene/normalization identity differs")
    frames, records = [], []
    cache = Path(cfg["raw_cache_root"]).resolve()
    for target in panel["query_targets"]:
        matches = [r for r in ledger["records"] if r["target"] == target]
        if len(matches) != 1:
            raise ValueError("Missing or colliding registered original target cache")
        r = matches[0]
        folder = (cache/r["cache_key"]).resolve()
        if not folder.is_relative_to(cache) or score.sha(folder/"manifest.json") != r["manifest_sha256"]:
            raise ValueError("Raw cache key escaped root or original manifest changed")
        m = score.read_json(folder/"manifest.json")
        identity = m["identity"]
        if identity != r["identity"] or score.logical_sha(identity) != r["cache_key"] or any(identity[k] != v for k,v in common.items()):
            raise ValueError("Original raw cache exact value identity changed")
        if identity["kind"] != "real" or identity["target"] != target or identity["count"] != panel["target_counts"][target] or identity["source_sha256"] != panel["source"]["sha256"] or identity["target_row_ids_sha256"] != panel["target_source_rows_sha256"][target]:
            raise ValueError("Raw cache has wrong training target/count/source-row identity")
        if identity["target_values_sha256"] != r["target_values_sha256"] or m["fdr_cached"] or m["rows"] != 18080 or m["backend"] != "official_pdex_0.1.27":
            raise ValueError("Incomplete, wrong-value or adjusted cache rejected")
        if score.sha(folder/"raw.parquet") != r["raw_sha256"] or r["raw_sha256"] != m["raw_sha256"]:
            raise ValueError("Original cached numerical bytes changed")
        frame = pl.read_parquet(folder/"raw.parquet")
        if {k:str(v) for k,v in frame.schema.items()} != r["raw_schema"]:
            raise ValueError("Original raw numerical dtypes changed")
        p = frame["p_value"].to_numpy()
        if p.dtype != np.dtype("float32") or not np.isfinite(p).all() or np.any(p<0) or np.any(p>1):
            raise ValueError("Wrong raw-p dtype or official postprocessed range")
        frames.append(frame)
        records.append(r)
    # with_global_bh additionally checks every raw field, exact original order,
    # target/reference/gene coverage and strict vertical dtype compatibility.
    return score.with_global_bh(frames, panel["query_targets"], panel["genes"]), records


def validate_materialized(path, encoded, panel, control_sha):
    import anndata as ad
    import numpy as np
    data = ad.read_h5ad(path, backed="r")
    try:
        labels, controls = score.check_prediction_metadata(data, panel)
        if score.selected_matrix_sha(data.X, controls) != control_sha:
            raise ValueError("Actual committed full reference changed")
        for target, mean in zip(panel["query_targets"], encoded):
            rows = np.flatnonzero(labels == target)
            for start in range(0,len(rows),256):
                block = np.asarray(data.X[rows[start:start+256],:])
                if block.dtype != np.dtype("float32") or not np.array_equal(block,np.broadcast_to(mean,block.shape)):
                    raise ValueError("Committed H5AD does not contain exactly constant encoded means")
    finally:
        data.file.close()


def parallel_raw(reference, means, counts, targets, workers):
    """Ordered map over shared read-only CDF; only asymptotic actual counts."""
    if min(counts) <= 8:
        raise ValueError("Exact-method small counts require the official backend")
    jobs = list(zip(means,counts,targets))
    def work(item):
        mean,count,target = item
        return reference.raw_frame(mean,count,target)
    started = time.monotonic()
    with ThreadPoolExecutor(max_workers=workers) as pool:
        results = list(pool.map(work,jobs))
    return [v[0] for v in results], [v[1] for v in results], time.monotonic()-started


def evaluate(commit_path, output):
    import anndata as ad
    import numpy as np
    import polars as pl
    from mwu_constant_reference_pdex_v2 import ConstantReference
    committed = score.read_json(commit_path)
    if committed["state"] != "all_six_materialized_predictions_committed" or committed["this_runner_query_cache_opened"] or committed["runner_sha256"] != score.sha(__file__):
        raise ValueError("Invalid or changed six-arm prediction commit")
    protocol_path = verify_record(committed["protocol"])
    cfg = configuration(protocol_path, execution=True)
    if Path(commit_path).resolve().parent != Path(cfg["run_dir"]).resolve() or output.resolve() != Path(cfg["run_dir"]).resolve()/"evaluation" or output.exists():
        raise ValueError("Explicit immutable committed run and fresh evaluation output required")
    os.sched_setaffinity(0, cfg["CPU_affinity"])
    for r in [committed["panel"],committed["encoded_means"],*committed["inputs"],*committed["metadata_panels"],*committed["predictions"]]:
        verify_record(r)
    if committed["runtime"] != pinned_runtime() or [r["role"] for r in committed["predictions"]] != ARMS:
        raise ValueError("Frozen metric/kernel runtime or all-arm set changed")
    panel = score.read_json(committed["panel"]["path"])
    resources = resource_preflight(cfg, panel)
    with np.load(committed["encoded_means"]["path"],allow_pickle=False) as saved:
        if saved["targets"].astype(str).tolist()!=panel["query_targets"] or saved["genes"].astype(str).tolist()!=panel["genes"]:
            raise ValueError("Committed encoded prediction axes changed")
        means = {arm:saved[arm].copy() for arm in ARMS}
    for r in committed["predictions"]:
        validate_materialized(r["path"],means[r["role"]],panel,cfg["control_values_sha256"])
    output.mkdir()
    score.write_json(output/"before_query_raw_cache.json",dict(at_utc=score.now(),
        all_prediction_commit_sha256=score.sha(commit_path), all_prediction_bytes_and_values_verified=True,
        query_targets=panel["query_targets"], fit_targets=panel["fit_targets"], resources=resources,
        previous_query_MAE_PDS_used_for_arm_screening=True, no_raw_expression_will_be_read=True, **FLAGS))
    real, real_records = read_real_raw(cfg,panel)
    real.write_parquet(output/"real_full_query34_de.parquet")
    control_path = next(r["path"] for r in cfg["inputs"] if r["role"] == "controls")
    control = ad.read_h5ad(control_path)
    if control.X.dtype != np.dtype("float32") or score.matrix_sha(control.X) != cfg["control_values_sha256"]:
        raise ValueError("The shared CDF actual decoded control identity changed")
    reference = ConstantReference(control.X, panel["genes"])
    targets = panel["query_targets"]
    counts = [panel["target_counts"][t] for t in targets]
    summary = dict(state="six_fixed_postscreen_training_DES_complete", completed_at_utc=None,
                   all_prediction_commit_sha256=score.sha(commit_path), phase="postscreen TRAINING inner-panel diagnostic",
                   previous_query_MAE_PDS_used_for_arm_screening=True, learned_only_is_full_agent_free=False,
                   actual_target_counts=panel["target_counts"], query_targets=targets,
                   real_raw_cache_receipts=real_records, full_BH_family=34*18080,
                   reference_cells=38176, normalization_target_sum=54611., native_dtype="float32",
                   reference_prepare_seconds=reference.build_elapsed_seconds, resources=resources,
                   official_preprocessing_pipeline_executed=False, official_default_backend_changed=False,
                   official_DES_function="cell_eval.metrics._de.de_overlap_metric(k=None,metric=overlap,fdr=.05,ABS_FOLD_CHANGE)",
                   arms={}, **FLAGS)
    for arm in ARMS:
        arm_out = output/arm
        arm_out.mkdir()
        print(f"raw-stat target map {arm}: {len(targets)} targets, workers={cfg['workers']}",flush=True)
        frames, diagnostics, elapsed = parallel_raw(reference,means[arm],counts,targets,cfg["workers"])
        raw = pl.concat(frames,how="vertical")
        raw.write_parquet(arm_out/"raw_full_panel_without_FDR.parquet")
        pred = score.with_global_bh(frames,targets,panel["genes"])
        pred.write_parquet(arm_out/"pred_full_query34_de.parquet")
        values = official_DES(real,pred,targets)
        pl.DataFrame({"perturbation":targets,"overlap_at_N":[values[t] for t in targets]}).write_csv(arm_out/"DES_per_target.csv")
        score.write_json(arm_out/"raw_backend_diagnostics.json",dict(targets=targets,diagnostics=diagnostics,
             elapsed_wall_seconds=elapsed, worker_count=cfg["workers"], **FLAGS))
        summary["arms"][arm] = dict(DES=float(np.mean(list(values.values()))),
            per_target_DES=values, raw_map_wall_seconds=elapsed,
            cumulative_target_seconds=sum(d["elapsed_seconds"] for d in diagnostics),
            real_FDR_discoveries=int((real["fdr"]<.05).sum()), predicted_FDR_discoveries=int((pred["fdr"]<.05).sum()))
        print(json.dumps({"arm":arm,**summary["arms"][arm]},sort_keys=True),flush=True)
        del frames,raw,pred
        gc.collect()
    summary["completed_at_utc"] = score.now()
    score.write_json(output/"summary.json",summary)
    hashes = {str(p.relative_to(output)):score.sha(p) for p in sorted(output.rglob("*")) if p.is_file()}
    score.write_json(output/"completion.json",dict(exit_code=0,at_utc=score.now(),output_hashes=hashes,
        all_prediction_commit_sha256=score.sha(commit_path),runner_sha256=score.sha(__file__),**FLAGS))
    return summary


def main():
    parser = argparse.ArgumentParser(description=__doc__)
    sub = parser.add_subparsers(dest="command",required=True)
    prep = sub.add_parser("prepare")
    prep.add_argument("--protocol",type=Path,required=True)
    prep.add_argument("--run-dir",type=Path,required=True)
    run = sub.add_parser("run")
    run.add_argument("--commit",type=Path,required=True)
    run.add_argument("--outdir",type=Path,required=True)
    args = parser.parse_args()
    if os.name != "posix":
        raise ValueError("Real preparation/evaluation is remote-only")
    result = prepare(args.protocol,args.run_dir) if args.command == "prepare" else evaluate(args.commit,args.outdir)
    print(json.dumps(result,sort_keys=True,allow_nan=False),flush=True)


if __name__ == "__main__":
    main()
```
