## Supplementary reproducibility code and frozen predictions for "Centered cross-context perturbation transfer on the public 2025 Virtual Cell Challenge benchmark": compute_real_de_panel.py

```
"""Compute and cache one native real-DE panel after prediction freezing.

Preparation only: no TEST input is opened unless this script is explicitly run.
Native Scanpy count normalization uses the complete object's positive-library
median. A remote Float32 memmap bounds memory; a single pdex call performs the
full target-by-gene BH family. Reuse the resulting real_de.csv for every model.
"""
from __future__ import annotations

import argparse
import hashlib
import importlib.metadata
import json
import os
from pathlib import Path
import resource
import time


def sha(path):
    h = hashlib.sha256()
    with Path(path).open('rb') as stream:
        for block in iter(lambda: stream.read(8 * 1024**2), b''):
            h.update(block)
    return h.hexdigest()


def write(path, value):
    Path(path).write_text(json.dumps(value, indent=2, allow_nan=False) + '\n', encoding='utf-8')


def main():
    parser = argparse.ArgumentParser(description=__doc__)
    parser.add_argument('--real-h5ad', type=Path, required=True)
    parser.add_argument('--prediction-archive', type=Path, required=True)
    parser.add_argument('--prediction-sha256', required=True)
    parser.add_argument('--outdir', type=Path, required=True)
    parser.add_argument('--workers', type=int, default=8)
    parser.add_argument('--batch-size', type=int, default=100)
    parser.add_argument('--chunk-rows', type=int, default=512)
    parser.add_argument('--pert-col', default='target_gene')
    parser.add_argument('--control', default='non-targeting')
    parser.add_argument('--input-space', choices=('counts', 'normlog'), default='counts')
    args = parser.parse_args()
    if os.name != 'posix':
        raise RuntimeError('Run on the authorized numerical server')
    if sha(args.prediction_archive) != args.prediction_sha256:
        raise ValueError('Frozen prediction archive hash differs')
    os.environ.update(USE_EXPERIMENTAL='0', CUDA_VISIBLE_DEVICES='', OMP_NUM_THREADS='1',
                      OPENBLAS_NUM_THREADS='1', MKL_NUM_THREADS='1', POLARS_MAX_THREADS='4')
    args.outdir.mkdir(parents=True, exist_ok=False)
    started = time.monotonic()
    import anndata as ad
    import numpy as np
    import pandas as pd
    import scanpy as sc
    from scipy import sparse
    import pdex._single_cell as pdex
    versions = {p: importlib.metadata.version(p) for p in ('cell-eval', 'pdex', 'scanpy', 'scipy', 'numpy', 'polars')}
    if versions['cell-eval'] != '0.6.6' or versions['pdex'] != '0.1.27' or pdex.use_experimental:
        raise ValueError('Pinned official nonexperimental evaluator required')
    with np.load(args.prediction_archive, allow_pickle=False) as archive:
        expected_targets = archive['targets'].astype(str).tolist()
        expected_genes = archive['genes'].astype(str).tolist()
    # All real expression/statistics reads start here, after the supplied frozen hash.
    real_sha = sha(args.real_h5ad)
    data = ad.read_h5ad(args.real_h5ad, backed='r')
    try:
        labels = data.obs[args.pert_col].astype(str).to_numpy()
        genes = data.var_names.astype(str).tolist()
        targets = sorted(set(labels) - {args.control})
        counts = {t: int(np.sum(labels == t)) for t in sorted(set(labels))}
        if set(targets) != set(expected_targets) or genes != expected_genes or args.control not in counts:
            raise ValueError('Frozen prediction and real panel axes differ')
        n, g = data.shape
        median = None
        if args.input_space == 'counts':
            libraries = np.empty(n, dtype=np.float64)
            for start in range(0, n, args.chunk_rows):
                block = data.X[start:start + args.chunk_rows]
                payload = block.data if sparse.issparse(block) else np.asarray(block)
                if not np.isfinite(payload).all() or np.any(payload < 0) or np.any(np.abs(payload - np.rint(payload)) > 1e-4):
                    raise ValueError('Declared count input must contain finite nonnegative integers')
                libraries[start:start + block.shape[0]] = np.asarray(block.sum(axis=1)).ravel()
            median = float(np.median(libraries[libraries > 0]))
        matrix_path = args.outdir / 'real_native_normlog_F32.npy'
        matrix = np.lib.format.open_memmap(matrix_path, mode='w+', dtype=np.float32, shape=(n, g))
        for start in range(0, n, args.chunk_rows):
            block = data.X[start:start + args.chunk_rows]
            if args.input_space == 'counts':
                part = ad.AnnData(X=block)
                sc.pp.normalize_total(part, target_sum=median, inplace=True)
                sc.pp.log1p(part)
                block = part.X
            if sparse.issparse(block):
                block = block.toarray()
            block = np.asarray(block, dtype=np.float32)
            if not np.isfinite(block).all() or np.any(block < 0):
                raise ValueError('Normalized expression is not finite/nonnegative')
            matrix[start:start + block.shape[0]] = block
            if start % (args.chunk_rows * 40) == 0:
                print(json.dumps({'state': 'normalizing', 'rows_done': start + block.shape[0], 'rows_total': n}), flush=True)
        matrix.flush()
        np.savez_compressed(args.outdir / 'panel_axes.npz', labels=labels.astype(str),
                            genes=np.asarray(genes), targets=np.asarray(targets),
                            counts=np.asarray([counts[t] for t in targets], dtype=np.int64))
    finally:
        data.file.close()
    normal_seconds = time.monotonic() - started
    dense_bytes = n * g * 4
    write(args.outdir / 'inputs.json', {'real_h5ad': str(args.real_h5ad), 'real_sha256': real_sha,
          'prediction_archive': str(args.prediction_archive), 'prediction_sha256': args.prediction_sha256,
          'source_sha256': sha(__file__), 'versions': versions, 'input_space': args.input_space,
          'native_positive_library_median': median, 'shape': [n, g], 'target_counts': counts,
          'targets': targets, 'genes_sha256': hashlib.sha256('\n'.join(genes).encode()).hexdigest(),
          'BH_family_size': len(targets) * g, 'workers': args.workers, 'batch_size': args.batch_size,
          'normalized_matrix_bytes': dense_bytes, 'pdex_shared_memory_bytes': dense_bytes,
          'panel_axes_sha256': sha(args.outdir / 'panel_axes.npz'),
          'normalization_seconds': normal_seconds})
    # One whole panel: pdex skips reference-vs-itself rows before global BH.
    normalized = ad.AnnData(X=matrix, obs=pd.DataFrame({args.pert_col: labels}), var=pd.DataFrame(index=genes))
    de_started = time.monotonic()
    frame = pdex.parallel_differential_expression(normalized, reference=args.control,
            groupby_key=args.pert_col, num_workers=args.workers, batch_size=args.batch_size,
            metric='wilcoxon', tie_correct=True, is_log1p=True, exp_post_agg=True,
            clip_value=20, as_polars=True)
    if frame.height != len(targets) * g or set(frame['target'].unique().to_list()) != set(targets):
        raise ValueError('Native real DE target/gene coverage differs')
    frame.write_csv(args.outdir / 'real_de.csv')
    frame.write_parquet(args.outdir / 'real_de.parquet')
    summary = {'state': 'complete', 'de_seconds': time.monotonic() - de_started,
               'elapsed_seconds': time.monotonic() - started, 'rows': frame.height,
               'real_de_csv_sha256': sha(args.outdir / 'real_de.csv'),
               'real_de_parquet_sha256': sha(args.outdir / 'real_de.parquet'),
               'peak_owner_RSS_KiB': resource.getrusage(resource.RUSAGE_SELF).ru_maxrss,
               'peak_child_RSS_KiB': resource.getrusage(resource.RUSAGE_CHILDREN).ru_maxrss,
               'prediction_sha256_before_first_real_read': args.prediction_sha256,
               'reuse': 'Pass real_de.csv as MetricsEvaluator de_real; do not recompute per prediction'}
    write(args.outdir / 'summary.json', summary)
    write(args.outdir / 'completion.json', {'exit_code': 0, 'summary_sha256': sha(args.outdir / 'summary.json')})
    print(json.dumps(summary), flush=True)


if __name__ == '__main__':
    main()
```
