## Supplementary reproducibility code and frozen predictions for "Centered cross-context perturbation transfer on the public 2025 Virtual Cell Challenge benchmark": mwu_constant_reference_process_v1.py

```
"""Spawned-process sharing for the unchanged exact constant-MWU v2 core.

Only the transport and reference storage differ. All gene/raw-frame arithmetic
is delegated unchanged to mwu_constant_reference_pdex_v2.ConstantReference.
"""
from __future__ import annotations
from concurrent.futures import ProcessPoolExecutor
import hashlib,json,multiprocessing as mp,os,pickle,time
from pathlib import Path

_REFERENCE=None

def _sha(path):
    h=hashlib.sha256()
    with Path(path).open('rb') as stream:
        for block in iter(lambda:stream.read(8*1024**2),b''):h.update(block)
    return h.hexdigest()

def write_reference_cache(reference,path):
    """Persist exact sorted values and tie metadata once; no numeric conversion."""
    import numpy as np
    path=Path(path);path.mkdir(parents=True,exist_ok=False);started=time.monotonic()
    sorted_values=np.lib.format.open_memmap(path/'sorted.npy',mode='w+',dtype=reference.dtype,shape=reference.sorted.shape)
    for start in range(0,len(reference.genes),256):sorted_values[start:start+256]=reference.sorted[start:start+256]
    sorted_values.flush();del sorted_values
    offsets=np.r_[0,np.cumsum([len(v) for v in reference.starts],dtype=np.int64)]
    np.save(path/'offsets.npy',offsets)
    for name in ('starts','counts','values'):
        np.save(path/(name+'.npy'),np.concatenate(getattr(reference,name)))
    # Preserve the original list's scalar classes as well as their values.
    with (path/'reference_means.pkl').open('wb') as stream:pickle.dump(reference.reference_means,stream,protocol=5)
    import mwu_constant_reference,mwu_constant_reference_pdex_v2
    manifest=dict(format='exact_constant_MWU_process_memmap_v1',genes=reference.genes,reference_cells=reference.n,dtype=str(reference.dtype),sorted_shape=list(reference.sorted.shape),
                  reference_sha256=reference.reference_sha256,reference_prepare_seconds=reference.build_elapsed_seconds,cache_write_seconds=time.monotonic()-started,
                  unchanged_core_sha256={str(Path(p).resolve()):_sha(p) for p in (mwu_constant_reference.__file__,mwu_constant_reference_pdex_v2.__file__)})
    (path/'manifest.json').write_text(json.dumps(manifest,indent=2));return manifest

def load_reference_cache(path):
    """Rehydrate exact class fields; reference=None is valid only for n>8."""
    import numpy as np
    from mwu_constant_reference_pdex_v2 import ConstantReference
    path=Path(path);manifest=json.loads((path/'manifest.json').read_text())
    if manifest['format']!='exact_constant_MWU_process_memmap_v1':raise ValueError('Wrong cache format')
    for p,digest in manifest['unchanged_core_sha256'].items():
        if _sha(p)!=digest:raise ValueError('Exact core changed since cache build')
    ref=ConstantReference.__new__(ConstantReference)
    ref.genes=manifest['genes'];ref.n=manifest['reference_cells'];ref.dtype=np.dtype(manifest['dtype']);ref.reference=None
    ref.sorted=np.load(path/'sorted.npy',mmap_mode='r',allow_pickle=False)
    offsets=np.load(path/'offsets.npy',mmap_mode='r',allow_pickle=False)
    for name in ('starts','counts','values'):
        flat=np.load(path/(name+'.npy'),mmap_mode='r',allow_pickle=False)
        setattr(ref,name,[flat[offsets[i]:offsets[i+1]] for i in range(len(ref.genes))])
    with (path/'reference_means.pkl').open('rb') as stream:ref.reference_means=pickle.load(stream)
    ref.reference_sha256=manifest['reference_sha256'];ref.build_elapsed_seconds=0.
    if not ref.sorted.flags.c_contiguous or ref.sorted.flags.writeable or ref.sorted.shape!=tuple(manifest['sorted_shape']):raise ValueError('Read-only contiguous sorted mmap required')
    return ref

def _initialize(cache,cpus):
    global _REFERENCE
    identity=mp.current_process()._identity
    if cpus:os.sched_setaffinity(0,{cpus[(identity[-1]-1)%len(cpus)]})
    _REFERENCE=load_reference_cache(cache)

def _work(job):
    means,count,target=job
    if count<=8:raise ValueError('Small-count exact branch requires original control')
    return _REFERENCE.raw_frame(means,count,target)

def parallel_raw(cache,means,counts,targets,workers=8,cpus=None):
    """Ordered spawn map; never fork an initialized Polars threadpool."""
    if len(means)!=len(counts) or len(counts)!=len(targets) or min(counts)<=8:raise ValueError('Aligned asymptotic-count target jobs required')
    started=time.monotonic()
    with ProcessPoolExecutor(max_workers=workers,mp_context=mp.get_context('spawn'),initializer=_initialize,initargs=(str(cache),list(cpus or []))) as pool:
        results=list(pool.map(_work,zip(means,counts,targets),chunksize=1))
    return [v[0] for v in results],[v[1] for v in results],time.monotonic()-started
```
