## Supplementary reproducibility code and frozen predictions for "Centered cross-context perturbation transfer on the public 2025 Virtual Cell Challenge benchmark": predict_numeric_kolf_centered_R1_v5.py

```
"""Centered R1 v5 inference: fit-public gamma1.25, quantile alpha1, DE-kernel beta.25.

Uses TRAIN150 DE labels/control, public KOLF responses, and old numerical
prediction features. Query expression and evaluation reports are not inputs.
"""
from __future__ import annotations
import argparse,json,os,sys,time
from pathlib import Path
for k in ('OMP_NUM_THREADS','OPENBLAS_NUM_THREADS','MKL_NUM_THREADS'):os.environ[k]='4'
import numpy as np
import scipy,sklearn
from scipy.stats import rankdata
from sklearn.isotonic import IsotonicRegression
from predict_numeric_kolf_isotonic_v2 import ROOT,CAP,sha,aligned,actual_abs_log2_fc

PARAMETERS={'expression_quantile_alpha':1.,'KOLF_DE_score_mix_beta':.25,
            'KOLF_expression_public_center_gamma':1.25,
            'KOLF_center_mean':'per gene, observed finite public TRAIN150 source only; zero if no observed fit coordinate',
            'KOLF_DE_kernel_ridge_alpha':1.,'KOLF_DE_kernel_residual_beta':.5,
            'isotonic_scale':.3,'strict_rank_coefficient':.02,'magnitude_floor':.015,
            'source_direction_threshold_quantized_units':2,'source_quantization_step':.04,
            'source_direction_threshold_decoded_float32':float(np.float32(.08)),
            'source_absZ_rank':'average rank / eligible gene count',
            'old_M_absFC_quantile_method':'linear','own_input_excluded':True,
            'own_gene_zero':True,'unresolved_TAZ_missing':True}

def get_bank(path,targets,genes):
    with np.load(path,allow_pickle=False) as a:
        z=aligned(a,targets,genes,'source_delta')[:,:,0].astype(float)
        mask=aligned(a,targets,genes,'source_mask')[:,:,0].astype(bool)
    return z,mask

def fit_public_center(train_z,train_mask):
    valid=train_mask&np.isfinite(train_z);counts=valid.sum(axis=0)
    mean=np.divide(np.where(valid,train_z,0.).sum(axis=0),counts,
                   out=np.zeros(train_z.shape[1]),where=counts>0)
    return mean,counts,{'public_fit_targets':train_z.shape[0],
                        'known_public_fit_targets':int(np.any(valid,axis=1).sum()),
                        'genes_with_observed_fit_mean':int((counts>0).sum()),
                        'mean_semantics':'Observed finite published decoded coordinates; no original finite/missing mask recovered',
                        'direction_threshold_policy':'Apply abs(centered response)>=float32(.08); source coverage mask unchanged',
                        'DE_kernel_centered':False}

def kolf_de_score(train_z,train_mask,query_z,query_mask,train,targets,genes,fit,d,old):
    """Same fit-known subset, L2+fit-trace solve and F32 cache as R1 screen."""
    z=np.concatenate([train_z,query_z]);mask=np.concatenate([train_mask,query_mask])
    all_targets=tuple(train)+tuple(targets);gi={g:i for i,g in enumerate(genes)}
    x=np.where(mask,z,0.)
    for i,t in enumerate(all_targets):
        if t in gi:x[i,gi[t]]=0.
    norm=np.linalg.norm(x,axis=1);known=norm>1e-12
    x=np.divide(x,norm[:,None],out=np.zeros_like(x),where=known[:,None])
    gram=x@x.T;diag=np.diag(gram);train_index={t:i for i,t in enumerate(train)}
    fi=np.asarray([train_index[t] for t in fit]);selection=known[fi]
    fi=fi[selection];d=d[selection];frequency=d.mean(axis=0)
    scale=float(diag[fi].mean());coef=np.linalg.solve(gram[np.ix_(fi,fi)]/scale+np.eye(len(fi)),d-frequency)
    qi=np.arange(len(train),len(all_targets));qknown=known[qi]
    # The development R1 16-arm screen consumed a float32 saved KOLF score.
    new=old.astype(float).copy()
    new[qknown]=np.clip(frequency+.5*(gram[np.ix_(qi[qknown],fi)]/scale@coef),0.,1.)
    new=new.astype(np.float32).astype(float)
    mixed=.75*old.astype(float)+.25*new
    return mixed,{'fit_total':len(fit),'source_known_fit':len(fi),
                  'source_missing_fit_excluded':[t for t,k in zip(fit,selection) if not k],
                  'source_known_queries':int(qknown.sum()),
                  'source_missing_queries_old_DE_fallback':[t for t,k in zip(targets,qknown) if not k],
                  'fit_trace_scale':scale,'KOLF_score_intermediate_dtype':'float32',
                  'kernel':'Aligned public KOLF vectors; own input0; masked0; per-target L2; known-fit trace mean',
                  'intercept':'DE mask mean over source-known TRAIN150 fitting subset'}

def render(expression,score,c,z,mask,targets,genes):
    expression=expression.astype(float);c=c.astype(float);gi={g:i for i,g in enumerate(genes)}
    eligible=mask&(c[None,:]>0)&(expression>0)
    for i,t in enumerate(targets):
        if t in gi:eligible[i,gi[t]]=False
    strong=eligible&(abs(z)>=float(np.float32(.08)))
    f0=np.sign(expression-c)*actual_abs_log2_fc(expression,c);mapped=f0.copy()
    for i in range(len(targets)):
        ok=eligible[i]
        if ok.any():
            r=rankdata(abs(z[i,ok]),method='average')/int(ok.sum())
            mapped[i,ok]=np.sign(z[i,ok])*np.quantile(abs(f0[i,ok]),r,method='linear')
    logc=np.log(np.expm1(c),where=c>0,out=np.full_like(c,-np.inf))
    # alpha1 still follows the same arithmetic as the fixed 16-arm screen.
    f=(1.-1.)*f0+1.*mapped
    decoded=np.clip(np.logaddexp(0.,logc+np.log(2.)*f),0.,CAP)
    adapted=expression.copy();adapted[strong]=decoded[strong]
    desired=actual_abs_log2_fc(adapted,c);sign=np.sign(adapted-c)
    u=np.stack([rankdata(row,method='average')/len(row) for row in score])
    projection=np.empty_like(desired)
    for i in range(len(targets)):
        model=IsotonicRegression(increasing=True,out_of_bounds='clip').fit(u[i,c>0],desired[i,c>0])
        projection[i]=model.predict(u[i])
    magnitude=.3*projection+.02*u+.015
    means=np.clip(np.logaddexp(0.,logc+np.log(2.)*sign*magnitude),0.,CAP).astype(np.float32)
    own=[]
    for i,t in enumerate(targets):
        if t!='TAZ' and t in gi and c[gi[t]]>0:means[i,gi[t]]=0.;own.append(t)
    if not np.isfinite(means).all():raise ValueError('Nonfinite prediction')
    return means,{'quantile_eligible_coordinates':int(eligible.sum()),
                  'quantile_strong_source_coordinates':int(strong.sum()),'own_zero_applied':len(own),
                  'weak_or_missing_expression_policy':'Exact old M coordinate retained before isotonic rendering'}

def infer(args):
    started=time.monotonic();os.sched_setaffinity(0,set(range(16,24)))
    with np.load(args.train_bank,allow_pickle=False) as a:
        train=tuple(map(str,a['targets']));genes=tuple(map(str,a['genes']));c=a['control_mean'].astype(float)
    inputs=[args.train_bank,args.train_kolf_bank,args.query_kolf_bank,args.train_DE_prior]
    if args.features:
        if args.expression_predictions or args.de_scores:raise ValueError('Choose combined or separate prediction features')
        with np.load(args.features,allow_pickle=False) as a:
            targets=tuple(map(str,a['targets']));expression=aligned(a,targets,genes,'expression_direction_source')
            old=aligned(a,targets,genes,'de_score').astype(float)
        inputs.append(args.features)
        allowed={str(args.features):['targets','genes','expression_direction_source','de_score']}
    else:
        if not args.expression_predictions or not args.de_scores:raise ValueError('Expression and DE prediction inputs required')
        with np.load(args.expression_predictions,allow_pickle=False) as a:
            targets=tuple(map(str,a['targets']));expression=aligned(a,targets,genes,'predicted_means')
        with np.load(args.de_scores,allow_pickle=False) as a:old=aligned(a,targets,genes,'de_score').astype(float)
        inputs.extend([args.expression_predictions,args.de_scores])
        allowed={str(args.expression_predictions):['targets','genes','predicted_means'],str(args.de_scores):['targets','genes','de_score']}
    if len(targets) not in (50,100) or len(set(targets))!=len(targets) or set(train)&set(targets):
        raise ValueError('Expected independent50/100 target prediction panel')
    train_z,train_mask=get_bank(args.train_kolf_bank,train,genes)
    z,mask=get_bank(args.query_kolf_bank,targets,genes)
    with np.load(args.train_DE_prior,allow_pickle=False) as a:
        fit=tuple(map(str,a['fit_targets']));d=a['D'].astype(float)
        if set(fit)!=set(train) or tuple(map(str,a['genes']))!=genes:raise ValueError('TRAIN150 DE label axes')
    score,kernel=kolf_de_score(train_z,train_mask,z,mask,train,targets,genes,fit,d,old)
    public_mean,public_counts,centering=fit_public_center(train_z,train_mask)
    centered_z=z-1.25*public_mean[None,:]
    means,rendering=render(expression,score,c,centered_z,mask,targets,genes)
    centering['published_zero_became_threshold_strong']=int((mask&(z==0)&(abs(centered_z)>=float(np.float32(.08)))).sum())
    replay=None
    if args.replay:
        with np.load(args.replay,allow_pickle=False) as a:expected=aligned(a,targets,genes,'predicted_means')
        delta=abs(means-expected)
        replay={'expected_path':str(args.replay),'expected_sha256':sha(args.replay),
                'changed_entries':int(np.sum(means!=expected)),
                'max_absolute_mean_difference':float(delta.max()),'allowed_absolute_tolerance':args.replay_atol}
        if replay['max_absolute_mean_difference']>args.replay_atol:raise ValueError('Replay mismatch '+json.dumps(replay))
    args.output.mkdir(parents=True,exist_ok=False);path=args.output/'predictions.npz'
    np.savez_compressed(path,targets=np.asarray(targets),genes=np.asarray(genes),predicted_means=means)
    center_path=args.output/'fitted_public_center.npz'
    np.savez_compressed(center_path,genes=np.asarray(genes),mean=public_mean,observed_fit_counts=public_counts)
    centering.update(fitted_center_path=str(center_path),fitted_center_sha256=sha(center_path))
    helper=Path(__file__).with_name('predict_numeric_kolf_isotonic_v2.py')
    receipt={'state':'complete','recipe':'Selected_centered_R1_gamma1p25_quantile_alpha1_DEbeta025_v5','parameters':PARAMETERS,
             'prediction_path':str(path),'prediction_sha256':sha(path),'code_path':str(Path(__file__).resolve()),
             'code_sha256':sha(__file__),'pure_helper_code_sha256':{str(helper):sha(helper)},
             'input_sha256':{str(p):sha(p) for p in dict.fromkeys(inputs)},'prediction_feature_members_read':allowed,
             'query_targets':len(targets),'genes':len(genes),'kernel_fit':kernel,'renderer':rendering,'public_centering':centering,
             'source_units':'Published author mean NTC z, signed int8*.04; no original missing/finite mask',
             'query_expression_read':False,'query_evaluation_report_read':False,'prediction_agent_used':False,
             'runtime_versions':{'python':sys.version.split()[0],'numpy':np.__version__,'scipy':scipy.__version__,'scikit_learn':sklearn.__version__},
             'replay':replay,'elapsed_seconds':time.monotonic()-started}
    (args.output/'prediction.json').write_text(json.dumps(receipt,indent=2)+'\n')
    print(json.dumps(receipt),flush=True)

if __name__=='__main__':
    p=argparse.ArgumentParser(description=__doc__)
    p.add_argument('--features',type=Path)
    p.add_argument('--expression-predictions',type=Path)
    p.add_argument('--de-scores',type=Path)
    p.add_argument('--train-bank',type=Path,default=ROOT/'data/processed/gct_TRAIN_raw_bank_v1/raw_bank.npz')
    p.add_argument('--train-kolf-bank',type=Path,default=ROOT/'data/processed/kolf2_1j_browser_selected_v1/source_bank.npz')
    p.add_argument('--train-DE-prior',dest='train_DE_prior',type=Path,default=ROOT/'runs/training150_DE_macro_full150_final_prior_v1/priors/TRAIN150_final_fit/prior.npz')
    p.add_argument('--query-kolf-bank',type=Path,required=True)
    p.add_argument('--output',type=Path,required=True)
    p.add_argument('--replay',type=Path)
    p.add_argument('--replay-atol',type=float,default=0.,help='Exact selected development replay required before prediction')
    infer(p.parse_args())
```
