## Supplementary reproducibility code and frozen predictions for "Centered cross-context perturbation transfer on the public 2025 Virtual Cell Challenge benchmark": predict_numeric_kolf_isotonic_v2.py

```
"""Fixed numerical KOLF direction + isotonic magnitude inference.

Inputs are existing numerical predictions, TRAIN control statistics and public
KOLF2.1J responses. No query expression or evaluation report is an input.
"""
from __future__ import annotations
import argparse
from collections import Counter
import hashlib,json,os,re,time
from pathlib import Path
for key in ('OMP_NUM_THREADS','OPENBLAS_NUM_THREADS','MKL_NUM_THREADS'):os.environ[key]='4'
import numpy as np
import scipy,sklearn,sys
from scipy.stats import rankdata
from sklearn.isotonic import IsotonicRegression

ROOT=Path('/data1/luanhaoyang/virtualcell')
CAP=float(np.nextafter(np.float32(15),np.float32(0)))
PARAMETERS={'KOLF_gain':16.,'KOLF_direction_weight':.25,'KOLF_DE_rank_weight':0.,
            'isotonic_scale':.3,'strict_rank_coefficient':.02,'magnitude_floor':.015,
            'own_gene_zero':True,'unresolved_TAZ_missing':True}

def sha(path):
    h=hashlib.sha256()
    with Path(path).open('rb') as f:
        for b in iter(lambda:f.read(8*1024**2),b''):h.update(b)
    return h.hexdigest()

def aligned(archive,targets,genes,key):
    ids=tuple(map(str,archive['targets']))
    if len(set(ids))!=len(ids) or tuple(map(str,archive['genes']))!=tuple(genes):raise ValueError('Prediction axis')
    return archive[key][[ids.index(t) for t in targets]].copy()

def gene_columns(genes,source_genes,identity_path):
    """Same literal-first, unique stable-ID fallback as development bank."""
    value=json.loads(Path(identity_path).read_text())
    pairs=[(r['symbol'],r['stable_gene_id']) for r in value['output_gene_metadata']
           if r.get('stable_mapping_available') and r.get('stable_gene_id')]
    by_symbol={};by_id={}
    for symbol,ensg in pairs:
        if not re.fullmatch(r'ENSG\d{11}',ensg):raise ValueError('Nonliteral stable ID')
        by_symbol.setdefault(symbol,set()).add(ensg);by_id.setdefault(ensg,set()).add(symbol)
    mapping={s:next(iter(ids)) for s,ids in by_symbol.items()
             if len(ids)==1 and len(by_id[next(iter(ids))])==1}
    count=Counter(source_genes);index={g:i for i,g in enumerate(source_genes) if count[g]==1}
    cols=[];methods=[]
    for g in genes:
        if g=='TAZ':col,method=-1,'unresolved_TAZ'
        elif g in index:col,method=index[g],'literal_symbol'
        elif mapping.get(g) in index:col,method=index[mapping[g]],'unique_stable_id'
        else:col,method=-1,'missing_or_ambiguous'
        cols.append(col);methods.append(method)
    return np.asarray(cols),methods

def actual_abs_log2_fc(pred,control):
    """The pdex zero branches used by the measured development combination."""
    t=np.expm1(pred.astype(np.float64));r=np.expm1(control.astype(np.float64))[None,:]
    with np.errstate(divide='ignore',invalid='ignore',over='ignore'):
        ratio=np.divide(t,r,out=np.ones_like(t),where=r>0)
    ratio=np.where((r==0)&(t>0),20.,ratio)
    ratio=np.where((r>0)&(t==0),.05,ratio)
    return np.abs(np.log2(ratio))

def predict(expression,de_score,c,sd,z,mask,targets,genes):
    expression=expression.astype(float);de_score=de_score.astype(float)
    c=c.astype(float);sd=sd.astype(float);z=z.astype(float);mask=mask.astype(bool)
    shape=(len(targets),len(genes))
    if any(a.shape!=shape for a in (expression,de_score,z,mask)):raise ValueError('Feature geometry')
    u=np.stack([rankdata(x,method='average')/len(x) for x in de_score])
    desired=actual_abs_log2_fc(expression,c);projection=np.empty_like(desired)
    for i in range(len(targets)):
        iso=IsotonicRegression(increasing=True,out_of_bounds='clip').fit(u[i,c>0],desired[i,c>0])
        projection[i]=iso.predict(u[i])
    magnitude=.3*projection+.02*u+.015
    transferred=np.clip(c+16*sd*z,0.,CAP);transferred[~mask]=expression[~mask]
    sign=np.sign(.75*expression+.25*transferred-c)
    fallback=(z==0)|~mask;sign[fallback]=np.sign(expression-c)[fallback]
    logc=np.log(np.expm1(c),where=c>0,out=np.full_like(c,-np.inf))
    means=np.clip(np.logaddexp(0.,logc+np.log(2.)*sign*magnitude),0.,CAP).astype(np.float32)
    gi={g:i for i,g in enumerate(genes)};own=[]
    for i,t in enumerate(targets):
        if t!='TAZ' and t in gi and c[gi[t]]>0:means[i,gi[t]]=0.;own.append(t)
    if not np.isfinite(means).all():raise ValueError('Nonfinite prediction')
    return means,len(own)

def infer(args):
    started=time.monotonic();os.sched_setaffinity(0,set(range(32,40)))
    inputs=[args.train_bank,args.kolf_bank]
    with np.load(args.train_bank,allow_pickle=False) as a:
        genes=tuple(map(str,a['genes']));c=a['control_mean'].astype(float);sd=a['control_sd_floor'].astype(float)
    if args.features:
        if args.expression_predictions or args.de_scores:raise ValueError('Choose combined features or two feature files')
        with np.load(args.features,allow_pickle=False) as a:
            targets=tuple(map(str,a['targets']))
            expression=aligned(a,targets,genes,'expression_direction_source')
            de_score=aligned(a,targets,genes,'de_score')
        inputs.append(args.features)
        allowed_members={str(args.features):['targets','genes','expression_direction_source','de_score']}
    else:
        if not args.expression_predictions or not args.de_scores:raise ValueError('Two prediction features required')
        with np.load(args.expression_predictions,allow_pickle=False) as a:
            targets=tuple(map(str,a['targets']));expression=aligned(a,targets,genes,'predicted_means')
        with np.load(args.de_scores,allow_pickle=False) as a:de_score=aligned(a,targets,genes,'de_score')
        inputs += [args.expression_predictions,args.de_scores]
        allowed_members={str(args.expression_predictions):['targets','genes','predicted_means'],
                         str(args.de_scores):['targets','genes','de_score']}
    if len(set(targets))!=len(targets) or len(targets) not in (50,100):raise ValueError('Expected independent50/100 target panel')
    with np.load(args.kolf_bank,allow_pickle=False) as a:
        z=aligned(a,targets,genes,'source_delta')[:,:,0]
        mask=aligned(a,targets,genes,'source_mask')[:,:,0]
    means,own_count=predict(expression,de_score,c,sd,z,mask,targets,genes)
    replay=None
    if args.replay:
        with np.load(args.replay,allow_pickle=False) as a:expected=aligned(a,targets,genes,'predicted_means')
        replay={'expected_path':str(args.replay),'expected_sha256':sha(args.replay),
                'changed_entries':int(np.sum(means!=expected)),
                'max_absolute_mean_difference':float(np.max(np.abs(means-expected)))}
        if replay['changed_entries']:raise ValueError('Exact development replay differs: '+json.dumps(replay))
    out=args.output;out.mkdir(parents=True,exist_ok=False);path=out/'predictions.npz'
    np.savez_compressed(path,targets=np.asarray(targets),genes=np.asarray(genes),predicted_means=means)
    receipt={'state':'complete','recipe':'numeric_kolf_isotonic_v2','parameters':PARAMETERS,
             'prediction_path':str(path),'prediction_sha256':sha(path),'code_path':str(Path(__file__).resolve()),
             'code_sha256':sha(__file__),'input_sha256':{str(p):sha(p) for p in inputs},
             'prediction_feature_members_read':allowed_members,
             'query_targets':len(targets),'genes':len(genes),'own_zero_applied':own_count,
             'KOLF_available_targets':int(mask.any(axis=1).sum()),'KOLF_output_genes_observed':int(mask.any(axis=0).sum()),
             'fallback':'Missing public response or observed quantized zero retains old numerical M direction',
             'magnitude':'Per-query isotonic(absFC of old M) on fixed KRR percentile; fit only TRAIN control>0 genes',
             'query_expression_read':False,'query_evaluation_report_read':False,'prediction_agent_used':False,
             'runtime_versions':{'python':sys.version.split()[0],'numpy':np.__version__,
                                 'scipy':scipy.__version__,'scikit_learn':sklearn.__version__},
             'replay':replay,'elapsed_seconds':time.monotonic()-started}
    (out/'prediction.json').write_text(json.dumps(receipt,indent=2)+'\n')
    print(json.dumps(receipt),flush=True)

if __name__=='__main__':
    p=argparse.ArgumentParser(description=__doc__)
    p.add_argument('--features',type=Path)
    p.add_argument('--expression-predictions',type=Path)
    p.add_argument('--de-scores',type=Path)
    p.add_argument('--train-bank',type=Path,default=ROOT/'data/processed/gct_TRAIN_raw_bank_v1/raw_bank.npz')
    p.add_argument('--kolf-bank',type=Path,required=True)
    p.add_argument('--output',type=Path,required=True)
    p.add_argument('--replay',type=Path)
    infer(p.parse_args())
```
