## Supplementary reproducibility code and frozen predictions for "Centered cross-context perturbation transfer on the public 2025 Virtual Cell Challenge benchmark": predict_numeric_ranktransfer_v1.py

```
"""Numeric VCC inference: public response transfer, TRAIN DE KRR and rank curve.

Reads only TRAIN assets, public external responses, and the B/targets/genes
members of the legacy numerical prediction archive. No query expression input.
"""
from __future__ import annotations
import argparse
import hashlib
import json
import os
from pathlib import Path
import sys
import time

for key in ('OMP_NUM_THREADS','OPENBLAS_NUM_THREADS','MKL_NUM_THREADS'):
    os.environ[key]='4'
import numpy as np
from scipy.stats import rankdata

ROOT=Path('/data1/luanhaoyang/virtualcell')
CAP=float(np.nextafter(np.float32(15),np.float32(0)))
TRAIN=ROOT/'data/processed/gct_TRAIN_raw_bank_v1/raw_bank.npz'
PRIOR=ROOT/'runs/training150_DE_macro_full150_final_prior_v1/priors/TRAIN150_final_fit/prior.npz'


def sha(path):
    h=hashlib.sha256()
    with Path(path).open('rb') as f:
        for chunk in iter(lambda:f.read(8*1024**2),b''):h.update(chunk)
    return h.hexdigest()


def reorder(archive,targets,genes,key):
    at=list(map(str,archive['targets']))
    if list(map(str,archive['genes']))!=genes or len(set(at))!=len(at):
        raise ValueError('Source gene or target identities differ')
    return archive[key][[at.index(t) for t in targets]].copy()


def infer(args):
    started=time.monotonic()
    if not ROOT.is_dir():raise RuntimeError('Numerical arrays stay on99.32 /data1')
    sys.path[:0]=[str(ROOT/'ops'),str(ROOT/'scripts')]
    from fast_gct_de_krr_rank_20261020 import build_source_gram
    out=args.output.resolve();out.mkdir(parents=True,exist_ok=False)
    with np.load(TRAIN,allow_pickle=False) as a:
        train=list(map(str,a['targets']));genes=list(map(str,a['genes']))
        c=a['control_mean'].astype(float);sd=a['control_sd_floor'].astype(float)
        tz=a['source_delta'].copy();tm=a['source_mask'].copy()
    with np.load(args.legacy_b,allow_pickle=False) as a:
        # np.load is lazy: none of the historical prediction-agent arms is read.
        targets=list(map(str,a['targets']))
        b=reorder(a,targets,genes,'B').astype(float)
    if set(targets)&set(train) or len(targets) not in (50,100) or b.shape!=(len(targets),18080):
        raise ValueError('Expected independent official50/100 query identities')
    if args.gemz_bank:
        with np.load(args.gemz_bank,allow_pickle=False) as a:
            z=reorder(a,targets,genes,'source_delta').astype(float)
            mask=reorder(a,targets,genes,'source_mask').astype(bool)
            valid_source_orders=(['K562_gwps','rpe1','K562essential'],
                                 ['K562_gwps_normalized_bulk_01','rpe1_normalized_bulk_01',
                                  'K562_essential_normalized_bulk_01'])
            if 'source_ids' in a and list(map(str,a['source_ids'])) not in valid_source_orders:
                raise ValueError('Public GEM-Z source ordering differs')
    else:
        from fast_validation_rawsource_ensemble_20261020 import public_deltas
        cfg=json.loads((ROOT/'configs/gct_TRAIN_raw_bank_v1_frozen.json').read_text())
        z,mask,_=public_deltas(targets,genes,cfg)
    if z.shape!=(len(targets),18080,3) or mask.shape!=z.shape:
        raise ValueError('Public three-source shape differs')
    observed=mask.sum(axis=2)>0
    delta=np.divide(np.where(mask,z,0).sum(axis=2),mask.sum(axis=2),out=np.zeros(b.shape),where=observed)
    for i,t in enumerate(targets):
        if t in genes:observed[i,genes.index(t)]=False
    # Preserve the F32 intermediate encodings used by development candidates.
    shared=b.mean(axis=0)
    centered=np.clip(c+32.*(b-shared),0,CAP).astype(np.float32)
    raw=np.where(observed,np.clip(c+128.*sd*delta,0,CAP),b).astype(np.float32)
    old=np.clip(.25*centered.astype(float)+.75*raw.astype(float),0,CAP).astype(np.float32)
    with np.load(args.xatlas_bank,allow_pickle=False) as a:
        xt=list(map(str,a['targets']))
        if list(map(str,a['genes']))!=genes:raise ValueError('X-atlas gene identities differ')
        xi=[xt.index(t) for t in targets]
        xd=a['source_delta'][:,xi].astype(float);xm=a['source_mask'][:,xi].astype(bool)
    if xd.shape!=(2,len(targets),18080) or xm.shape!=xd.shape:
        raise ValueError('Expected context-first HCT116/HEK293T responses')
    known=xm.sum(axis=0)>0
    xdelta=np.divide(np.where(xm,xd,0).sum(axis=0),xm.sum(axis=0),out=np.zeros(b.shape),where=known)
    for i,t in enumerate(targets):
        if t in genes:known[i,genes.index(t)]=False
    xraw=np.where(known,np.clip(c+128.*xdelta,0,CAP),old)
    expression=np.clip(.25*old.astype(float)+.75*xraw,0,CAP).astype(np.float32)
    with np.load(PRIOR,allow_pickle=False) as a:
        fit=list(map(str,a['fit_targets']));d=a['D'].astype(float);frequency=a['f'].astype(float)
        if set(fit)!=set(train) or list(map(str,a['genes']))!=genes:raise ValueError('TRAIN DE prior identities differ')
    gram=build_source_gram(np.concatenate([tz,z]),np.concatenate([tm,mask]))
    fi=np.asarray([train.index(t) for t in fit]);qi=np.arange(len(train),len(train)+len(targets))
    k=gram/np.mean(np.diag(gram)[fi])
    de_score=np.clip(frequency+args.beta*k[np.ix_(qi,fi)]@np.linalg.solve(k[np.ix_(fi,fi)]+args.alpha*np.eye(len(fi)),d-frequency),0,1).astype(np.float32)
    u=np.stack([rankdata(row,method='average')/len(row) for row in de_score])
    sign=np.sign(expression.astype(float)-c)
    logc=np.log(np.expm1(c),where=c>0,out=np.full_like(c,-np.inf))
    fc=sign*args.scale*(args.eps+u**args.power)
    means=np.clip(np.logaddexp(0,logc+np.log(2)*fc),0,CAP).astype(np.float32)
    if not np.isfinite(means).all():raise ValueError('Nonfinite predictions')
    path=out/'predictions.npz'
    np.savez_compressed(path,targets=np.asarray(targets),genes=np.asarray(genes),predicted_means=means,de_score=de_score,expression_direction_source=expression)
    replay=None
    if args.replay:
        with np.load(args.replay,allow_pickle=False) as a:
            expected=reorder(a,targets,genes,'predicted_means')
        replay=dict(max_absolute_mean_difference=float(np.max(np.abs(means-expected))),
                    changed_entries=int(np.sum(means!=expected)),expected_sha256=sha(args.replay))
        if replay['max_absolute_mean_difference']>1e-6:
            raise ValueError('Prediction-only development replay differs: '+str(replay))
    inputs=[TRAIN,PRIOR,args.legacy_b,args.xatlas_bank]+([args.gemz_bank] if args.gemz_bank else [])
    report=dict(state='complete',prediction_path=str(path),prediction_sha256=sha(path),
                source_sha256=sha(__file__),input_sha256={str(p):sha(p) for p in inputs},
                alpha=args.alpha,beta=args.beta,eps=args.eps,scale=args.scale,power=args.power,
                source_gains=dict(center32=True,GEMZ=128,Xatlas=128),
                mixing=dict(Bcenter=.25,GEMZ=.75,previous=.25,Xatlas=.75),
                query_targets=len(targets),genes=len(genes),query_expression_read=False,
                prediction_agent_used=False,replay=replay,elapsed_seconds=time.monotonic()-started)
    (out/'prediction.json').write_text(json.dumps(report,indent=2),encoding='utf-8')
    print(json.dumps(report),flush=True)


if __name__=='__main__':
    p=argparse.ArgumentParser(description=__doc__)
    p.add_argument('--legacy-b',type=Path,required=True)
    p.add_argument('--xatlas-bank',type=Path,required=True)
    p.add_argument('--gemz-bank',type=Path)
    p.add_argument('--output',type=Path,required=True)
    p.add_argument('--replay',type=Path)
    p.add_argument('--alpha',type=float,default=1.)
    p.add_argument('--beta',type=float,default=.5)
    p.add_argument('--eps',type=float,default=.01)
    p.add_argument('--scale',type=float,default=1.)
    p.add_argument('--power',type=float,default=2.)
    infer(p.parse_args())
```
