## Supplementary reproducibility code and frozen predictions for "Centered cross-context perturbation transfer on the public 2025 Virtual Cell Challenge benchmark": prepare_kolf_public_query_bank_v2.py

```
"""Select public author KOLF rows using existing numerical target identities.

Uses already cached publisher chunks on /data1; no query expression input.
The same unique literal/stable output-gene join as development is retained.
"""
import argparse
from collections import Counter
import json,os,time
from pathlib import Path
import numpy as np
from predict_numeric_kolf_isotonic_v2 import ROOT,sha,gene_columns
COMMIT='8f84531a784f24b14d7e2c9b80c375d7e78fd46c'
def main(args):
    started=time.monotonic();os.sched_setaffinity(0,set(range(32,40)))
    with np.load(args.prediction_identities,allow_pickle=False) as a:
        targets=tuple(map(str,a['targets']));genes=tuple(map(str,a['genes']))
    if len(set(targets))!=len(targets) or len(set(genes))!=len(genes):raise ValueError('Duplicate output identities')
    source=ROOT/f'data/external/kolf2_1j_viewer2026/{COMMIT}'
    meta_path=source/'meta.json';meta=json.loads(meta_path.read_text())
    identity=ROOT/'data/processed/protein_identity_metadata_v1/identity_manifest.json'
    cols,methods=gene_columns(genes,meta['genes'],identity);valid=cols>=0
    target_index={t:i for i,t in enumerate(meta['perts'])}
    n=len(targets);z=np.zeros((n,len(genes),1),np.float32);mask=np.zeros_like(z,bool)
    source_rows=np.full((n,1),-1,np.int32);receipts=[];payload=0
    for i,t in enumerate(targets):
        if t not in target_index or t=='TAZ':continue
        r=target_index[t];chunk,within=divmod(r,meta['chunk_size']);path=source/f'chunk_{chunk}.bin'
        with path.open('rb') as f:
            f.seek(within*meta['row_bytes']);raw=f.read(meta['row_bytes'])
        if len(raw)!=meta['row_bytes']:raise ValueError('Short public source row')
        q=np.frombuffer(raw,dtype=np.int8)
        # Keep F32 multiplication order exactly equal to the development bank.
        z[i,valid,0]=q[cols[valid]].astype(np.float32)*float(meta['scale'])
        mask[i,valid,0]=True;source_rows[i,0]=r;payload+=len(raw)
        receipts.append({'target':t,'author_row':r,'chunk':str(path),'offset':within*meta['row_bytes'],
                         'bytes':len(raw),'selected_row_sha256':__import__('hashlib').sha256(raw).hexdigest()})
    out=args.output;out.mkdir(parents=True,exist_ok=False);path=out/'source_bank.npz'
    np.savez_compressed(path,targets=np.asarray(targets),genes=np.asarray(genes),contexts=np.asarray(['KOLF2.1J_iPSC']),
                        source_delta=z,source_mask=mask,source_target_available=mask.any(axis=1),source_row_index=source_rows)
    used_chunks=sorted({r['chunk'] for r in receipts})
    report={'state':'complete','namespace':'kolf_TEST_public_20261021_v1','author_commit':COMMIT,
            'source_units':'published signed int8*.04 NTC-referenced CPM+log1p z-score',
            'quantization_step':float(meta['scale']),'original_missing_finite_mask_available':False,
            'zero_policy':'Observed quantized code, uncertain tiny effect; inference retains old M direction',
            'targets':n,'genes':len(genes),'available_targets':int(mask.any(axis=1).sum()),
            'available_output_genes':int(valid.sum()),'alignment_method_counts':dict(Counter(methods)),
            'missing_targets':[t for i,t in enumerate(targets) if not mask[i].any()],
            'source_bank_path':str(path),'source_bank_sha256':sha(path),'code_sha256':sha(__file__),
            'mapping_code_sha256':sha(Path(__file__).with_name('predict_numeric_kolf_isotonic_v2.py')),
            'identity_manifest_sha256':sha(identity),'author_meta_sha256':sha(meta_path),
            'prediction_identity_path':str(args.prediction_identities),'prediction_identity_file_sha256':sha(args.prediction_identities),
            'prediction_identity_members_read':['targets','genes'],'source_rows':receipts,
            'public_chunk_sha256':{p:sha(p) for p in used_chunks},
            'cached_public_bytes_read':payload,'elapsed_seconds':time.monotonic()-started,
            'query_expression_read':False,'query_evaluation_report_read':False}
    (out/'source_bank.json').write_text(json.dumps(report,indent=2)+'\n')
    print(json.dumps({k:v for k,v in report.items() if k!='source_rows'}),flush=True)
if __name__=='__main__':
    p=argparse.ArgumentParser(description=__doc__)
    p.add_argument('--prediction-identities',type=Path,required=True)
    p.add_argument('--output',type=Path,required=True)
    main(p.parse_args())
```
