## Supplementary reproducibility code and frozen predictions for "Centered cross-context perturbation transfer on the public 2025 Virtual Cell Challenge benchmark": prepare_pie_source_rows.py

```
"""Select public PIE source rows, fetch bounded NPY ranges, and align identities.

No H1 expression/outcomes, fitting, evaluation or TEST identities are inputs.
`plan` is offline. `fetch` performs public HTTPS only when explicitly invoked.
`align` also supports the already-downloaded Replogle/Nadig directory.
"""
from __future__ import annotations

import argparse
import ast
from concurrent.futures import ThreadPoolExecutor
import csv
import hashlib
from http.client import IncompleteRead
import json
import os
from pathlib import Path
import re
import struct
import time
from urllib.parse import urlencode, urlparse
from urllib.request import Request, urlopen
import uuid

import numpy as np


def sha(path):
    h = hashlib.sha256()
    with Path(path).open('rb') as stream:
        for block in iter(lambda: stream.read(4 * 1024 * 1024), b''):
            h.update(block)
    return h.hexdigest()


def read_json(path):
    return json.loads(Path(path).read_text(encoding='utf-8-sig'))


def write_json(path, value):
    Path(path).write_text(json.dumps(value, indent=2) + '\n', encoding='utf-8')


def identities(paths):
    values = []
    for path in paths:
        path = Path(path)
        # Explicitly refuse the VCC TEST panel, including identity-only input.
        if 'test' in path.as_posix().lower():
            raise ValueError('Only TRAIN/VALIDATION identity files are permitted')
        if path.suffix == '.json':
            values.extend(read_json(path)['target_genes'])
        else:
            with path.open(encoding='utf-8-sig', newline='') as stream:
                values.extend(row['target_gene'] for row in csv.DictReader(stream))
    if any(not isinstance(x, str) or not x for x in values):
        raise ValueError('Invalid literal target identity')
    return list(dict.fromkeys(values))


def validate_meta(meta):
    if meta['format_version'] != 1 or meta['pert_kind'] != 'gene':
        raise ValueError('Unsupported publisher metadata')
    if len(meta['genes']) != meta['num_genes'] or len(set(meta['genes'])) != meta['num_genes']:
        raise ValueError('Gene axis must be unique')
    for key, number in [('context_to_id', 'num_contexts'), ('pert_to_id', 'num_perts')]:
        if sorted(meta[key].values()) != list(range(meta[number])):
            raise ValueError('Publisher vocabulary must be bijective and contiguous')
    if any('h1' == context.lower() for context in meta['context_to_id']):
        raise ValueError('H1 is not an external source context')


def load_metadata(directory):
    directory = Path(directory)
    meta = read_json(directory / 'meta.json')
    validate_meta(meta)
    # When fetched by the small metadata helper, verify every supplied receipt.
    manifest_path = directory / 'manifest.json'
    if manifest_path.exists():
        manifest = read_json(manifest_path)
        for item in manifest['files']:
            path = directory / item['name']
            if path.stat().st_size != item['bytes'] or sha(path) != item['sha256']:
                raise ValueError('Metadata receipt changed')
    arrays = {}
    for name in ['ctx_ids', 'pert_ids', 'ctrl_means']:
        path = directory / (name + '.npy')
        expected = meta['array_sha256'].get(path.name, meta['array_sha256'].get(name))
        if expected is None or sha(path) != expected:
            raise ValueError('Publisher small array SHA differs')
        arrays[name] = np.load(path, allow_pickle=False)
    if arrays['ctx_ids'].shape != (meta['num_rows'],) or arrays['pert_ids'].shape != (meta['num_rows'],):
        raise ValueError('Row-index shapes differ')
    if arrays['ctrl_means'].shape != (meta['num_contexts'], meta['num_genes']):
        raise ValueError('Control shape differs')
    return meta, arrays


def select_rows(meta, ctx, pert, targets):
    reverse = {value: key for key, value in meta['pert_to_id'].items()}
    wanted = set(targets) - {'TAZ'}  # no silent WWTR1/TAFAZZIN interpretation
    if ctx.dtype.kind not in 'iu' or pert.dtype.kind not in 'iu':
        raise ValueError('Source row IDs must be integer arrays')
    if np.any((ctx < 0) | (ctx >= meta['num_contexts'])) or np.any((pert < 0) | (pert >= meta['num_perts'])):
        raise ValueError('Source row IDs are out of range')
    rows = np.asarray([i for i, p in enumerate(pert) if reverse[int(p)] in wanted], dtype=np.int64)
    pairs = [(int(ctx[i]), int(pert[i])) for i in rows]
    if len(set(pairs)) != len(pairs):
        raise ValueError('Duplicate source context-target pair; no implicit aggregation')
    return rows


def npy_header(raw, total):
    if raw[:6] != b'\x93NUMPY' or len(raw) < 12:
        raise ValueError('Not a complete NPY header')
    version = tuple(raw[6:8])
    if version == (1, 0):
        offset = 10 + struct.unpack('<H', raw[8:10])[0]
        prefix = 10
    elif version in [(2, 0), (3, 0)]:
        offset = 12 + struct.unpack('<I', raw[8:12])[0]
        prefix = 12
    else:
        raise ValueError('Unsupported NPY version')
    if offset > len(raw) or offset > 4096:
        raise ValueError('NPY header exceeds bounded prefix')
    header = ast.literal_eval(raw[prefix:offset].decode('latin1' if version != (3, 0) else 'utf-8').strip())
    dtype, shape = np.dtype(header['descr']), tuple(header['shape'])
    if header['fortran_order'] or len(shape) != 2 or any(type(x) is not int or x < 1 for x in shape):
        raise ValueError('Only nonempty C-order row matrices are supported')
    if dtype.hasobject or dtype.kind not in 'fb' or dtype.itemsize not in [1, 4, 8]:
        raise ValueError('Only numeric floating / boolean arrays are supported')
    if offset + shape[0] * shape[1] * dtype.itemsize != total:
        raise ValueError('NPY shape/offset does not match pinned byte size')
    return offset, shape, dtype


def validate_range(status, headers, url, original_url, start, end, total):
    headers = {key.lower(): value for key, value in headers.items()}
    if status != 206 or headers.get('content-range') != f'bytes {start}-{end}/{total}':
        raise ValueError('Server did not return the exact pinned byte range')
    if headers.get('content-encoding', 'identity').lower() != 'identity':
        raise ValueError('Range is HTTP encoded')
    if 'content-length' in headers and int(headers['content-length']) != end - start + 1:
        raise ValueError('Range length differs')
    parsed = urlparse(url)
    host = parsed.hostname or ''
    if parsed.scheme != 'https' or not (host == 'huggingface.co' or host.endswith('.hf.co')):
        raise ValueError('Range redirect is outside publisher HTTPS hosts')
    if host == 'huggingface.co' and parsed.path != urlparse(original_url).path:
        raise ValueError('Publisher repository/revision changed')


def get_range(url, start, end, total, cache_dir=None):
    size = end - start + 1
    if not 0 <= start <= end < total or size > 4 * 1024 * 1024:
        raise ValueError('Range is outside the pinned object or 4MiB limit')
    identity = dict(url=url, start=start, end=end, total=total, bytes=size)
    cache = receipt = None
    if cache_dir is not None:
        cache_dir = Path(cache_dir)
        cache_dir.mkdir(parents=True, exist_ok=True)
        key = hashlib.sha256(json.dumps(identity, sort_keys=True).encode()).hexdigest()
        cache = cache_dir / (key + '.bin')
        receipt = cache.with_suffix('.json')
        if cache.exists() and receipt.exists():
            saved = read_json(receipt)
            if (any(saved.get(k) != v for k, v in identity.items()) or cache.stat().st_size != size
                    or sha(cache) != saved.get('sha256')):
                raise ValueError('Completed range cache SHA/identity differs')
            return cache.read_bytes()
    for attempt in range(1, 5):
        # Distinct URL per range/attempt prevents a stale HF redirect/range cache.
        query = urlencode(dict(download='1', pie_range=f'{start}-{end}', pie_nonce=uuid.uuid4().hex))
        request = Request(url + ('&' if '?' in url else '?') + query,
                          headers={'Range': f'bytes={start}-{end}', 'Accept-Encoding': 'identity',
                                   'Cache-Control': 'no-cache'})
        try:
            with urlopen(request, timeout=120) as response:
                validate_range(response.status, dict(response.headers), response.url, url, start, end, total)
                body = bytearray()
                # A read(n) may return fewer than n bytes without EOF. Keep reading,
                # including one extra byte to reject an overlong response body.
                while len(body) <= size:
                    block = response.read(min(65536, size + 1 - len(body)))
                    if not block:
                        break
                    body.extend(block)
            if len(body) != size:
                raise ValueError(f'Range body length differs: expected {size}, received {len(body)}')
            body = bytes(body)
            if cache is not None:
                temporary = cache.with_name(cache.name + '.part-' + uuid.uuid4().hex)
                try:
                    temporary.write_bytes(body)
                    os.replace(temporary, cache)
                finally:
                    temporary.unlink(missing_ok=True)
                temporary = receipt.with_name(receipt.name + '.part-' + uuid.uuid4().hex)
                try:
                    write_json(temporary, dict(**identity, sha256=hashlib.sha256(body).hexdigest()))
                    os.replace(temporary, receipt)
                finally:
                    temporary.unlink(missing_ok=True)
            return body
        except (OSError, ValueError, IncompleteRead) as error:
            if attempt == 4:
                raise RuntimeError(f'Range {start}-{end} failed after four bounded GET attempts: {type(error).__name__}: {error}') from error
            time.sleep(2 ** (attempt - 1))
    raise AssertionError('Unreachable range retry state')


def row_spans(rows, width, offset, max_bytes=4 * 1024 * 1024):
    spans = []
    for row in rows:
        row = int(row)
        if spans and row == spans[-1][1] + 1 and (row - spans[-1][0] + 1) * width <= max_bytes:
            spans[-1][1] = row
        else:
            spans.append([row, row])
    return [dict(first=a, last=b, start=offset + a * width, end=offset + (b + 1) * width - 1) for a, b in spans]


def fetch_selected(directory, inventory_path, targets, output, names, max_bytes, workers):
    directory, output = Path(directory), Path(output)
    if output.exists() or output.with_suffix('.json').exists():
        raise FileExistsError('Preserve existing selected source output')
    meta, small = load_metadata(directory)
    manifest = read_json(directory / 'manifest.json')
    revision, repo = manifest['revision'], manifest['repo']
    if not re.fullmatch('[0-9a-f]{40}', revision) or repo != 'arcinstitute/PIE_x_atlas_orion':
        raise ValueError('Expected immutable X-atlas publisher revision')
    inventory = {item['path'].split('/')[-1]: item for item in read_json(inventory_path)}
    rows = select_rows(meta, small['ctx_ids'], small['pert_ids'], targets)
    sizes = {'delta_p': 4, 'tested': 1, 'fdr': 4}
    budget = len(rows) * meta['num_genes'] * sum(sizes[name] for name in names) + 4096 * len(names)
    if budget > max_bytes:
        raise ValueError(f'Selected download exceeds cap: {budget} > {max_bytes}')
    arrays = dict(genes=np.asarray(meta['genes']), row_indices=rows,
                  ctx_ids=small['ctx_ids'][rows], pert_ids=small['pert_ids'][rows],
                  ctrl_means=small['ctrl_means'], metadata_json=np.asarray(json.dumps(meta)))
    receipts = []
    for name in names:
        entry = inventory[name + '.npy']
        expected = meta['array_sha256'].get(name + '.npy', meta['array_sha256'].get(name))
        if entry['lfs']['oid'] != expected:
            raise ValueError('Inventory and metadata publisher SHA disagree')
        total = entry['size']
        url = f'https://huggingface.co/datasets/{repo}/resolve/{revision}/preprocessed/{name}.npy'
        cache_dir = output.parent / 'range_cache'
        prefix = get_range(url, 0, min(4095, total - 1), total, cache_dir)
        offset, shape, dtype = npy_header(prefix, total)
        if shape != (meta['num_rows'], meta['num_genes']) or dtype.itemsize != sizes[name] or dtype.kind != ('b' if name == 'tested' else 'f'):
            raise ValueError('Actual publisher header differs from matrix metadata')
        spans = row_spans(rows, shape[1] * dtype.itemsize, offset)
        result = np.empty((len(rows), shape[1]), dtype=dtype)
        positions = {int(row): i for i, row in enumerate(rows)}
        def fetch(span):
            body = get_range(url, span['start'], span['end'], total, cache_dir)
            data = np.frombuffer(body, dtype=dtype).reshape(-1, shape[1])
            for i, row in enumerate(range(span['first'], span['last'] + 1)):
                result[positions[row]] = data[i]
            return dict(**span, sha256=hashlib.sha256(body).hexdigest())
        with ThreadPoolExecutor(max_workers=workers) as pool:
            ranges = list(pool.map(fetch, spans))
        arrays[name] = result
        receipts.append(dict(name=name, publisher_full_file_sha256=expected, full_file_sha256_verified=False,
                             header_prefix_sha256=hashlib.sha256(prefix).hexdigest(), header_offset=offset,
                             total_bytes=total, ranges=ranges))
    if not np.isfinite(arrays['delta_p'][arrays['tested']]).all():
        raise ValueError('Tested source delta is not finite')
    output.parent.mkdir(parents=True, exist_ok=True)
    with output.open('xb') as stream:
        np.savez_compressed(stream, **arrays)
    report = dict(state='public_source_rows_selected', rows=len(rows), genes=meta['num_genes'], repo=repo,
                  revision=revision, target_identity_count=len(targets), TAZ_excluded=True,
                  metadata_sha256=sha(directory / 'meta.json'), output_sha256=sha(output),
                  transfer_bytes=sum(sum(x['end']-x['start']+1 for x in r['ranges']) + min(4096,r['total_bytes']) for r in receipts),
                  transfer_bytes_definition='unique selected ranges; excludes retries and may reuse SHA-verified local cache',
                  range_cache=str(output.parent / 'range_cache'),
                  code_sha256=sha(__file__), arrays=receipts, H1_expression_read=False)
    write_json(output.with_suffix('.json'), report)
    return report


def mapping_from_manifest(path):
    value = read_json(path)
    if 'output_gene_metadata' in value:
        pairs = [(r['symbol'], r['stable_gene_id']) for r in value['output_gene_metadata']
                 if r.get('stable_mapping_available') and r.get('stable_gene_id')]
    else:
        pairs = list(value.get('symbol_to_ensembl', value).items())
    by_symbol, by_id = {}, {}
    for symbol, ensg in pairs:
        if not isinstance(symbol, str) or not isinstance(ensg, str) or not re.fullmatch(r'ENSG\d{11}', ensg):
            raise ValueError('Identity map must contain literal stable ENSG IDs')
        by_symbol.setdefault(symbol, set()).add(ensg)
        by_id.setdefault(ensg, set()).add(symbol)
    return {s: next(iter(ids)) for s, ids in by_symbol.items()
            if len(ids) == 1 and len(by_id[next(iter(ids))]) == 1}


def align_rows(meta, rows, ctx, pert, delta, tested, control, targets, genes, gene_map=None, fdr=None):
    validate_meta(meta)
    expected = (len(rows), meta['num_genes'])
    if delta.shape != expected or tested.shape != expected or tested.dtype.kind != 'b':
        raise ValueError('Selected delta/tested geometry or mask type differs')
    if ctx.shape != (len(rows),) or pert.shape != (len(rows),) or control.shape != (meta['num_contexts'], meta['num_genes']):
        raise ValueError('Selected identity/control geometry differs')
    if fdr is not None and fdr.shape != expected:
        raise ValueError('Selected FDR geometry differs')
    if not np.isfinite(delta[tested]).all() or not np.isfinite(control).all():
        raise ValueError('Observed source values are not finite')
    if len(set(targets)) != len(targets) or len(set(genes)) != len(genes):
        raise ValueError('Output identity axes must be unique')
    contexts = [name for name, _ in sorted(meta['context_to_id'].items(), key=lambda item: item[1])]
    source_index = {name: i for i, name in enumerate(meta['genes'])}
    columns = []
    for gene in genes:
        literal = gene_map.get(gene) if gene_map and gene_map.get(gene) in source_index else gene
        columns.append(source_index.get(literal, -1) if gene != 'TAZ' else -1)
    columns = np.asarray(columns)
    valid = columns >= 0
    shape = (len(targets), len(genes), len(contexts))
    out = np.full(shape, np.nan, dtype=np.float32)
    mask = np.zeros(shape, dtype=bool)
    ctrl = np.full((len(genes), len(contexts)), np.nan, dtype=np.float32)
    for context in range(len(contexts)):
        ctrl[valid, context] = control[context, columns[valid]]
    row_ids = np.full((len(targets), len(contexts)), -1, dtype=np.int64)
    target_index = {t: i for i, t in enumerate(targets) if t != 'TAZ'}
    reverse = {value: key for key, value in meta['pert_to_id'].items()}
    out_fdr = np.full(shape, np.nan, dtype=np.float32) if fdr is not None else None
    for i, row in enumerate(rows):
        target = reverse[int(pert[i])]
        if target not in target_index:
            continue
        ti, ci = target_index[target], int(ctx[i])
        if row_ids[ti, ci] != -1:
            raise ValueError('Duplicate context-target row; no implicit averaging')
        row_ids[ti, ci] = row
        observed = valid.copy()
        observed[valid] &= tested[i, columns[valid]] & np.isfinite(delta[i, columns[valid]])
        out[ti, observed, ci] = delta[i, columns[observed]]
        mask[ti, observed, ci] = True
        if fdr is not None:
            out_fdr[ti, observed, ci] = fdr[i, columns[observed]]
    result = dict(targets=np.asarray(targets), genes=np.asarray(genes), contexts=np.asarray(contexts),
                  source_delta=out, source_mask=mask, source_target_available=mask.any(axis=1),
                  source_row_index=row_ids, control_means=ctrl, control_mask=np.isfinite(ctrl))
    if out_fdr is not None:
        result['source_fdr'] = out_fdr
    return result


def main():
    parser = argparse.ArgumentParser(description=__doc__)
    parser.add_argument('action', choices=['plan', 'fetch', 'align'])
    parser.add_argument('--metadata-dir', type=Path, required=True)
    parser.add_argument('--targets', type=Path, nargs='+', required=True)
    parser.add_argument('--output', type=Path, required=True)
    parser.add_argument('--inventory', type=Path)
    parser.add_argument('--arrays', nargs='+', default=['delta_p', 'tested', 'fdr'], choices=['delta_p', 'tested', 'fdr'])
    parser.add_argument('--max-bytes', type=int, default=64 * 1024 * 1024)
    parser.add_argument('--workers', type=int, default=6)
    parser.add_argument('--selected', type=Path)
    parser.add_argument('--genes', type=Path)
    parser.add_argument('--gene-map', type=Path)
    parser.add_argument('--context-first', action='store_true', help='Save matrices as context,target,gene')
    args = parser.parse_args()
    if not 1 <= args.workers <= 16:
        parser.error('workers must be 1..16')
    targets = identities(args.targets)
    meta, small = load_metadata(args.metadata_dir)
    rows = select_rows(meta, small['ctx_ids'], small['pert_ids'], targets)
    if args.action == 'plan':
        report = dict(rows=len(rows), row_indices=rows.tolist(), source_genes=meta['num_genes'],
                      metadata_sha256=sha(args.metadata_dir / 'meta.json'),
                      selected_targets_by_context={name: sorted({t for t, p in meta['pert_to_id'].items()
                          if np.any((small['ctx_ids'][rows] == ci) & (small['pert_ids'][rows] == p))})
                          for name, ci in meta['context_to_id'].items()},
                      delta_tested_payload_bytes=len(rows) * meta['num_genes'] * 5,
                      delta_tested_fdr_payload_bytes=len(rows) * meta['num_genes'] * 9,
                      TAZ_excluded=True, H1_expression_read=False)
        args.output.parent.mkdir(parents=True, exist_ok=True)
        write_json(args.output, report)
    elif args.action == 'fetch':
        if not args.inventory or 'delta_p' not in args.arrays or 'tested' not in args.arrays:
            parser.error('fetch requires inventory and both delta_p/tested')
        report = fetch_selected(args.metadata_dir, args.inventory, targets, args.output, args.arrays, args.max_bytes, args.workers)
    else:
        if not args.genes:
            parser.error('align requires --genes identity CSV')
        if args.output.exists():
            raise FileExistsError(args.output)
        with args.genes.open(encoding='utf-8-sig', newline='') as stream:
            genes = [row[0] for row in csv.reader(stream) if row]
        mapping = mapping_from_manifest(args.gene_map) if args.gene_map else None
        if all(re.fullmatch(r'ENSG\d{11}', g) for g in meta['genes']) and not mapping:
            parser.error('ENSG source axis requires an explicit identity-only --gene-map')
        if args.selected:
            receipt = read_json(args.selected.with_suffix('.json'))
            if receipt['output_sha256'] != sha(args.selected) or receipt['metadata_sha256'] != sha(args.metadata_dir / 'meta.json'):
                raise ValueError('Selected source receipt mismatch')
            with np.load(args.selected, allow_pickle=False) as data:
                if data['row_indices'].tolist() != rows.tolist() or tuple(data['genes']) != tuple(meta['genes']):
                    raise ValueError('Selected source rows or genes differ from requested identities')
                values = align_rows(meta, rows, data['ctx_ids'], data['pert_ids'], data['delta_p'], data['tested'],
                                    small['ctrl_means'], targets, genes, mapping, data['fdr'] if 'fdr' in data else None)
            source_provenance = dict(selected_sha256=sha(args.selected), selected_receipt_sha256=sha(args.selected.with_suffix('.json')))
        else:
            # Reuse existing 181MB original source without a new download.
            loaded, source_provenance = {}, {}
            for name in ['delta_p', 'tested']:
                path = args.metadata_dir / (name + '.npy')
                expected = meta['array_sha256'].get(path.name, meta['array_sha256'].get(name))
                if sha(path) != expected:
                    raise ValueError('Original source SHA differs')
                loaded[name] = np.load(path, mmap_mode='r', allow_pickle=False)
                source_provenance[name + '_sha256'] = expected
            values = align_rows(meta, rows, small['ctx_ids'][rows], small['pert_ids'][rows], loaded['delta_p'][rows],
                                loaded['tested'][rows], small['ctrl_means'], targets, genes, mapping)
        available_by_context = values['source_target_available'].sum(axis=0).tolist()
        observed_by_context = values['source_mask'].sum(axis=(0, 1)).tolist()
        if args.context_first:
            for key in ['source_delta', 'source_mask', 'source_fdr']:
                if key in values:
                    values[key] = values[key].transpose(2, 0, 1)
            for key in ['source_target_available', 'source_row_index', 'control_means', 'control_mask']:
                values[key] = values[key].T
        args.output.parent.mkdir(parents=True, exist_ok=True)
        with args.output.open('xb') as stream:
            np.savez_compressed(stream, **values)
        report = dict(state='public_source_aligned', output_sha256=sha(args.output), source=source_provenance,
                      metadata_sha256=sha(args.metadata_dir / 'meta.json'), code_sha256=sha(__file__),
                      genes_sha256=sha(args.genes), mapping_sha256=sha(args.gene_map) if args.gene_map else None,
                      targets=len(targets), genes=len(genes), contexts=values['contexts'].tolist(),
                      source_delta_shape=list(values['source_delta'].shape),
                      layout='context,target,gene' if args.context_first else 'target,gene,context',
                      available_targets_by_context=available_by_context,
                      observed_gene_pairs_by_context=observed_by_context,
                      missing_policy='NaN plus false mask; no aliases or source-scale conversion', H1_expression_read=False)
        write_json(args.output.with_suffix('.json'), report)
    print(json.dumps({k: v for k, v in report.items() if k not in ['row_indices', 'selected_targets_by_context', 'arrays']}))


if __name__ == '__main__':
    main()
```
