## Supplementary reproducibility code and frozen predictions for "Centered cross-context perturbation transfer on the public 2025 Virtual Cell Challenge benchmark": prepare_TRAIN_gct_raw_bank_v1.py

```
"""Prepare pinned TRAIN/public inputs for GCT; no fit, score or external query.

Actual arrays stay on 99.32. This bank contains all TRAIN outcomes; subsequent
scope controllers must physically slice fit rows before calling any estimator.
"""
from __future__ import annotations

import argparse
from datetime import datetime, timezone
import gc
import hashlib
import json
import os
from pathlib import Path
import resource
import shutil
import sys
import time
import traceback

ROOT = Path('/data1/luanhaoyang/virtualcell')
OUTPUT = 'data/processed/gct_TRAIN_raw_bank_v1'
SOURCE_IDS = ('K562_gwps_normalized_bulk_01', 'rpe1_normalized_bulk_01',
              'K562_essential_normalized_bulk_01')


def sha(path):
    h = hashlib.sha256()
    with path.open('rb') as stream:
        while block := stream.read(8 * 1024**2):
            h.update(block)
    return h.hexdigest()


def write(path, value):
    with path.open('x', encoding='utf-8') as stream:
        json.dump(value, stream, indent=2, allow_nan=False)
        stream.write('\n')


def now():
    return datetime.now(timezone.utc).isoformat()


def handle():
    pid = os.getpid()
    tail = Path(f'/proc/{pid}/stat').read_text().rsplit(')', 1)[1].split()
    return dict(pid=pid, start_ticks=int(tail[19]), uid=os.getuid())


def local_path(relative):
    path = (ROOT / relative).resolve()
    if not path.is_relative_to(ROOT) or path == ROOT:
        raise ValueError('Path outside authorized project')
    return path


def run(cfg, config_sha):
    if os.name != 'posix' or Path(__file__).resolve() != ROOT/'scripts'/Path(__file__).name:
        raise ValueError('Actual data preparation is remote only')
    if cfg['schema'] != 'GCT_TRAIN_raw_bank_v1' or cfg['output'] != OUTPUT:
        raise ValueError('Unexpected configuration')
    cpus = {32, 33, 34, 35}
    if not cpus <= os.sched_getaffinity(0):
        raise ValueError('Required CPU affinity unavailable')
    mem = dict(line.split(':', 1) for line in Path('/proc/meminfo').read_text().splitlines())
    if int(mem['MemAvailable'].split()[0])*1024 < 100*1024**3:
        raise ValueError('Less than100GiB available; leave current evaluation alone')
    if shutil.disk_usage(ROOT).free < 128*1024**3:
        raise ValueError('Insufficient free disk')
    os.sched_setaffinity(0, cpus)
    resource.setrlimit(resource.RLIMIT_AS, (32*1024**3,)*2)
    for name in ('OPENBLAS_NUM_THREADS', 'OMP_NUM_THREADS', 'MKL_NUM_THREADS',
                 'NUMEXPR_NUM_THREADS', 'POLARS_MAX_THREADS'):
        os.environ[name] = '4'
    os.environ['CUDA_VISIBLE_DEVICES'] = ''
    # All bytes are checked before importing project modules or opening arrays.
    for relative, expected in cfg['pins'].items():
        if sha(local_path(relative)) != expected:
            raise ValueError('Input/code SHA mismatch: '+relative)
    import numpy as np
    sys.path.insert(0, str(ROOT/'src'))
    from virtualcell.transfer import read_source, gene_mapping, axis_hash, Z_SCALE
    from virtualcell.public_source_basis import resolve_targets
    output = local_path(OUTPUT)
    output.mkdir(exist_ok=False)
    write(output/'started.json', dict(UTC=now(), process=handle(), config_sha256=config_sha,
        CPU_affinity=sorted(cpus), RLIMIT_AS_bytes=32*1024**3,
        role='TRAIN_public_raw_preparation_no_model_or_score'))
    start = time.monotonic()
    try:
        with np.load(local_path(cfg['teacher']), allow_pickle=False) as data:
            groups = tuple(map(str, data['groups']))
            genes = tuple(map(str, data['genes']))
            means = data['means'].copy()
            variance = data['variances'].copy()
            counts = data['cell_counts'].copy()
            if (means.shape != (151, 18080) or variance.shape != means.shape or
                    len(set(groups)) != 151 or len(set(genes)) != 18080 or
                    groups.count('non-targeting') != 1 or float(data['target_sum']) != 54611):
                raise ValueError('Teacher shape/axis/normalization mismatch')
        cidx = groups.index('non-targeting')
        if int(counts[cidx]) != 38176 or not np.isfinite(means).all() or not np.isfinite(variance).all() or np.any(variance < 0):
            raise ValueError('Invalid full-NTC statistics')
        targets = tuple(t for t in groups if t != 'non-targeting')
        declared = json.loads(local_path(cfg['training_ids']).read_text(encoding='utf-8-sig'))['target_genes']
        if len(targets) != 150 or set(targets) != set(declared):
            raise ValueError('Not the fixed TRAIN150 identities')
        rows = [groups.index(t) for t in targets]
        target_means = means[rows].astype(np.float32)
        C = means[cidx].astype(np.float64)
        d = np.maximum(np.sqrt(variance[cidx].astype(np.float64)), .01)
        own = np.array([genes.index(t) if t in genes and t != 'TAZ' else -1 for t in targets], dtype=np.int64)
        Z = np.zeros((150, 18080, 3), dtype=np.float32)
        M = np.zeros(Z.shape, dtype=bool)
        audits = []
        for s, item in enumerate(cfg['sources']):
            source = read_source(local_path(item['path']))
            if source.source_id != SOURCE_IDS[s] or source.scale != Z_SCALE:
                raise ValueError('Source identity/units mismatch')
            mapping = resolve_targets(source, targets)
            known = [mapping[t] for t in targets if mapping[t] is not None]
            if len(known) != len(set(known)):
                raise ValueError('Duplicate canonical perturbation identity')
            encoded = tuple(mapping[t] or '__missing__'+t for t in targets)
            original, observed = source.aligned(encoded, source.genes)
            columns, alignment = gene_mapping(source.genes, source.metadata['source_expression_symbols'], genes)
            if 'TAZ' in genes:
                columns[genes.index('TAZ')] = -1
            matched = columns >= 0
            ev = original[:, columns[matched]]
            mask = observed[:, columns[matched]]
            if not np.isfinite(ev[mask]).all():
                raise ValueError('Nonfinite measured source value')
            Z[:, matched, s] = np.where(mask, ev, 0).astype(np.float32)
            M[:, matched, s] = mask
            audits.append(dict(source_id=source.source_id, target_mapping=mapping,
                alignment=alignment, forced_TAZ_output_unknown=True,
                measured_coordinates=int(mask.sum()), measured_zero_coordinates=int(np.sum(mask & (ev == 0))),
                source_context=source.metadata['source_context'],
                units='canonical population-weighted GEM-Z target minus population-weighted core control',
                source_clipping_applied=False))
            print(json.dumps(dict(state='source_aligned', source_id=source.source_id,
                                  measured_coordinates=int(mask.sum()))), flush=True)
            del source, original, observed, ev, mask
            gc.collect()
        with np.load(local_path(cfg['target_ada']), allow_pickle=False) as data:
            stored_targets = tuple(map(str, data['targets']))
            if len(stored_targets) != 150 or set(stored_targets) != set(targets):
                raise ValueError('Target ADA axis mismatch')
            order = [stored_targets.index(t) for t in targets]
            te = data['values'][order].astype(np.float64)
            ta = data['available'][order]
        with np.load(local_path(cfg['output_ada']), allow_pickle=False) as data:
            if tuple(map(str, data['genes'])) != genes or str(data['role']) != 'output_gene_prior':
                raise ValueError('Output ADA axis/role mismatch')
            ge = data['values'].astype(np.float64)
            ga = data['available']
        if te.shape != (150, 1536) or ge.shape != (18080, 1536) or ta.dtype.kind != 'b' or ga.dtype.kind != 'b':
            raise ValueError('ADA dimensions/masks mismatch')
        if not np.isfinite(te[ta]).all() or not np.isfinite(ge[ga]).all():
            raise ValueError('Nonfinite observed ADA vector')
        archive = output/'raw_bank.npz'
        np.savez(archive, targets=np.array(targets), genes=np.array(genes), source_ids=np.array(SOURCE_IDS),
                 source_delta=Z, source_mask=M, target_means=target_means, control_mean=C,
                 control_sd_floor=d, target_cell_counts=counts[rows], own_coordinate=own,
                 target_ada=te, target_ada_available=ta, output_ada=ge, output_ada_available=ga)
        # Verify array serialization and masks on actual saved bytes before commit.
        with np.load(archive, allow_pickle=False) as saved:
            for name, expected in (('source_delta', Z), ('source_mask', M), ('target_means', target_means),
                                   ('control_mean', C), ('control_sd_floor', d)):
                if not np.array_equal(saved[name], expected):
                    raise ValueError('Serialization mismatch: '+name)
        write(output/'manifest.json', dict(schema='GCT_TRAIN_raw_bank_v1', UTC=now(),
            config_sha256=config_sha, inputs_and_code_sha256=cfg['pins'],
            bank=dict(path=str(archive.relative_to(ROOT)), sha256=sha(archive), bytes=archive.stat().st_size),
            targets=list(targets), genes_axis_sha256=axis_hash(genes), source_ids=list(SOURCE_IDS),
            sources=audits, n_targets=150, n_genes=18080,
            target_ADA_available=int(ta.sum()), output_ADA_available=int(ga.sum()),
            control_definition='teacher full38176 NTC mean; sqrt population variance floored .01; float32 archived statistics promoted float64',
            normalization='fixed54611 per-cell log1p before averaging; no new normalization',
            source_missing_storage_zero=True, source_missing_is_observed_zero=False,
            own_coordinates_preserved_in_bank_but_excluded_by_model=True,
            PCA_or_supervised_transform_fitted=False, model_fitted=False, scoring_run=False,
            validation_test_accessed=False, TRAIN_outcomes_present=True,
            future_controller_must_physically_slice_fit_Y=True))
        write(output/'completion.json', dict(UTC=now(), exit_code=0, process=handle(),
            elapsed_seconds=time.monotonic()-start, manifest_sha256=sha(output/'manifest.json')))
        print(json.dumps(dict(state='TRAIN_raw_bank_committed', manifest_sha256=sha(output/'manifest.json'))), flush=True)
    except BaseException as exc:
        write(output/'failure.json', dict(UTC=now(), process=handle(), type=type(exc).__name__, message=str(exc)))
        traceback.print_exc()
        raise


if __name__ == '__main__':
    parser = argparse.ArgumentParser()
    parser.add_argument('--config', required=True, type=Path)
    parser.add_argument('--config-sha256', required=True)
    args = parser.parse_args()
    if sha(args.config) != args.config_sha256:
        raise ValueError('Configuration byte SHA mismatch')
    run(json.loads(args.config.read_text(encoding='utf-8-sig')), args.config_sha256)
```
