## Supplementary reproducibility code and frozen predictions for "Centered cross-context perturbation transfer on the public 2025 Virtual Cell Challenge benchmark": score_frozen_meanflat_panel_final_v2.py

```
"""Score frozen constant-target predictions against an independently built real cache.

Exact constant-MWU backend and native DES are reused. Native Polars group-by
means run in gene-column blocks to bound memory without changing the per-column
operation or complete row order. MAE/PDS call the installed official functions.
"""
from __future__ import annotations

import argparse
import csv
import gc
import hashlib
import importlib.metadata
import json
import os
from pathlib import Path
import resource
import sys
import time

ROOT = Path('/data1/luanhaoyang/virtualcell')
LIB = ROOT / 'runs/training_meanflat_DES_prepare_v1/scripts'
DEFAULT_CONTROL = ROOT / 'data/processed_ntc_scale_v1/full_training_NTC_L54611.h5ad'
ANCHORS = {'DES': .09505113849605429, 'PDS': .5082, 'MAE': .0249575331993401}


def sha(path):
    h = hashlib.sha256()
    with Path(path).open('rb') as stream:
        for block in iter(lambda: stream.read(8 * 1024**2), b''):
            h.update(block)
    return h.hexdigest()


def write(path, value):
    Path(path).write_text(json.dumps(value, indent=2, allow_nan=False) + '\n', encoding='utf-8')


def native_bulk_blocks(matrix, labels, block_genes=256):
    import numpy as np
    import polars as pl
    expected = np.unique(labels)
    parts = []
    for start in range(0, matrix.shape[1], block_genes):
        frame = pl.DataFrame(matrix[:, start:start + block_genes]).with_columns(groupby_key=labels)
        grouped = frame.group_by('groupby_key').mean().sort('groupby_key')
        if not np.array_equal(grouped['groupby_key'].to_numpy(), expected):
            raise ValueError('Native bulk group identity differs')
        parts.append(grouped.drop('groupby_key').to_numpy())
    return expected, np.concatenate(parts, axis=1)


def predicted_bulk(means, targets, counts, control, block_genes=256):
    import numpy as np
    # Match the meanflat renderer: original TRAIN control rows followed by each
    # target in frozen archive order, with its independently public cell count.
    labels = np.concatenate([np.repeat('non-targeting', len(control)),
                             np.repeat(targets, counts)])
    expected = np.unique(labels)
    parts = []
    for start in range(0, means.shape[1], block_genes):
        stop = min(start + block_genes, means.shape[1])
        block = np.empty((len(labels), stop - start), dtype=np.float32)
        block[:len(control)] = control[:, start:stop]
        offset = len(control)
        for mean, count in zip(means, counts):
            block[offset:offset + count] = mean[start:stop]
            offset += count
        keys, bulk = native_bulk_blocks(block, labels, stop - start)
        if not np.array_equal(keys, expected):
            raise ValueError('Predicted native bulk keys differ')
        parts.append(bulk)
    return expected, np.concatenate(parts, axis=1)


def official_bulk_metrics(real_keys, real_values, pred_keys, pred_values, genes):
    import anndata as ad
    import numpy as np
    import pandas as pd
    from cell_eval._types._anndata import PerturbationAnndataPair as Pair
    from cell_eval.metrics import _anndata as metrics
    if not np.array_equal(real_keys, pred_keys):
        raise ValueError('Real/pred native bulk keys differ')
    obs = pd.DataFrame({'target_gene': real_keys})
    var = pd.DataFrame(index=genes)
    pair = Pair(real=ad.AnnData(np.zeros_like(real_values), obs=obs.copy(), var=var.copy()),
                pred=ad.AnnData(np.zeros_like(pred_values), obs=obs.copy(), var=var.copy()),
                pert_col='target_gene', control_pert='non-targeting')
    pair.bulk_real['_default'] = (real_keys, real_values)
    pair.bulk_pred['_default'] = (pred_keys, pred_values)
    return {'MAE': metrics.mae(pair, embed_key=None),
            'PDS': metrics.discrimination_score(pair, metric='l1', embed_key=None, exclude_target_gene=True)}


def prepare_predictions(cfg, output, public_counts):
    """Independent prediction cache: no real file/cache is opened here."""
    import anndata as ad
    import numpy as np
    sys.path.insert(0, str(LIB))
    from mwu_constant_reference_pdex_v2 import ConstantReference
    import score_training_meanflat_DES as helper
    import score_training_fold as score
    started = time.monotonic()
    write(output / 'started.json', {'pid': os.getpid(), 'prediction_only': True,
          'test_truth_read': False, 'config': cfg, 'source_sha256': sha(__file__)})
    control = ad.read_h5ad(cfg.get('control_h5ad', str(DEFAULT_CONTROL)))
    genes = control.var_names.astype(str).tolist()
    if control.X.dtype != np.dtype('float32') or control.shape != (38176, len(genes)):
        raise ValueError('Original full TRAIN prediction NTC differs')
    reference = ConstantReference(control.X, genes)
    reference.sorted = np.ascontiguousarray(reference.sorted)
    print(json.dumps({'state': 'prediction_reference_ready', 'seconds': time.monotonic() - started}), flush=True)
    results = []
    for candidate in cfg['candidates']:
        t0 = time.monotonic()
        with np.load(candidate['path'], allow_pickle=False) as z:
            targets = z['targets'].astype(str).tolist()
            means = z[candidate.get('key', 'predicted_means')].copy()
            if z['genes'].astype(str).tolist() != genes:
                raise ValueError('Frozen prediction gene axis differs')
        counts = [public_counts[t] for t in targets]
        frames, diagnostics, raw_seconds = helper.parallel_raw(reference, means, counts, targets, cfg.get('workers', 4))
        predicted_de = score.with_global_bh(frames, targets, genes)
        keys, values = predicted_bulk(means, targets, counts, control.X, cfg.get('bulk_block_genes', 256))
        name = candidate['id']
        de_path = output / (name + '_pred_DE.parquet')
        bulk_path = output / (name + '_native_bulk.npz')
        predicted_de.write_parquet(de_path)
        np.savez_compressed(bulk_path, keys=keys, values=values, genes=np.asarray(genes))
        record = {'candidate': candidate, 'prediction_only': True, 'test_truth_read': False,
                  'targets': targets, 'counts_from_public_metadata': counts,
                  'counts_sha256': cfg['counts_sha256'], 'prediction_control_cells': len(control),
                  'reference_values_sha256': reference.reference_sha256,
                  'pred_DE_sha256': sha(de_path), 'native_bulk_sha256': sha(bulk_path),
                  'raw_map_seconds': raw_seconds, 'elapsed_seconds': time.monotonic() - t0,
                  'BH_family_size': len(targets) * len(genes), 'diagnostics': diagnostics}
        write(output / (name + '_prediction_cache.json'), record)
        results.append(record)
        print(json.dumps({'state': 'prediction_cache_complete', 'candidate': name,
                          'seconds': record['elapsed_seconds']}), flush=True)
        del frames, predicted_de
        gc.collect()
    summary = {'state': 'complete', 'prediction_only': True, 'test_truth_read': False,
               'candidates': results, 'elapsed_seconds': time.monotonic() - started}
    write(output / 'summary.json', summary)
    write(output / 'completion.json', {'exit_code': 0, 'summary_sha256': sha(output / 'summary.json')})


def main():
    parser = argparse.ArgumentParser(description=__doc__)
    parser.add_argument('--config', type=Path, required=True)
    parser.add_argument('--outdir', type=Path, required=True)
    parser.add_argument('--prediction-only', action='store_true')
    args = parser.parse_args()
    cfg = json.loads(args.config.read_text())
    os.sched_setaffinity(0, set(cfg.get('cpus', range(32, 40))))
    os.environ.update(USE_EXPERIMENTAL='0', CUDA_VISIBLE_DEVICES='', OMP_NUM_THREADS='1',
                      OPENBLAS_NUM_THREADS='1', MKL_NUM_THREADS='1', POLARS_MAX_THREADS='4')
    for candidate in cfg['candidates']:
        if sha(candidate['path']) != candidate['sha256']:
            raise ValueError('Frozen candidate hash differs before real cache access')
    if sha(cfg['counts_path']) != cfg['counts_sha256']:
        raise ValueError('Public frozen cell-count metadata hash differs')
    with Path(cfg['counts_path']).open(newline='') as stream:
        public_counts = {r['target_gene']: int(r['n_cells']) for r in csv.DictReader(stream)}
    args.outdir.mkdir(parents=True, exist_ok=False)
    started = time.monotonic()
    import anndata as ad
    import numpy as np
    import polars as pl
    sys.path.insert(0, str(LIB))
    from mwu_constant_reference_pdex_v2 import ConstantReference
    import score_training_meanflat_DES as helper
    import score_training_fold as score
    versions = {p: importlib.metadata.version(p) for p in ('cell-eval', 'pdex', 'scipy', 'numpy', 'polars')}
    if versions != {'cell-eval': '0.6.6', 'pdex': '0.1.27', 'scipy': '1.17.1', 'numpy': '2.4.6', 'polars': '2.0.0'}:
        raise ValueError('Pinned exact parity runtime differs')
    if args.prediction_only:
        prepare_predictions(cfg, args.outdir, public_counts)
        return
    cache = Path(cfg['real_cache_dir'])
    with np.load(cache / 'panel_axes.npz', allow_pickle=False) as z:
        labels, genes, real_targets = z['labels'].astype(str), z['genes'].astype(str).tolist(), z['targets'].astype(str).tolist()
    matrix = np.load(cache / 'real_native_normlog_F32.npy', mmap_mode='r', allow_pickle=False)
    if matrix.dtype != np.dtype('float32') or matrix.shape != (len(labels), len(genes)):
        raise ValueError('Real normalized memmap axes differ')
    block_genes = cfg.get('bulk_block_genes', 256)
    bulk_path = cache / 'real_native_bulk.npz'
    if bulk_path.exists():
        with np.load(bulk_path, allow_pickle=False) as z:
            real_keys, real_values = z['keys'].astype(str), z['values'].copy()
            if z['genes'].astype(str).tolist() != genes:
                raise ValueError('Cached native real bulk gene axis differs')
    else:
        real_keys, real_values = native_bulk_blocks(matrix, labels, block_genes)
        np.savez_compressed(bulk_path, keys=real_keys, values=real_values, genes=np.asarray(genes))
    del matrix
    gc.collect()
    real_de_path = cache / 'real_de.parquet'
    real_de = pl.read_parquet(real_de_path) if real_de_path.exists() else pl.read_csv(cache / 'real_de.csv')
    prediction_cache = Path(cfg['prediction_cache_dir']) if cfg.get('prediction_cache_dir') else None
    reference = None
    if prediction_cache is None:
        control = ad.read_h5ad(cfg.get('control_h5ad', str(DEFAULT_CONTROL)))
        if control.X.dtype != np.dtype('float32') or control.shape != (38176, len(genes)) or control.var_names.tolist() != genes:
            raise ValueError('Prediction full original TRAIN NTC differs')
        reference = ConstantReference(control.X, genes)
        reference.sorted = np.ascontiguousarray(reference.sorted)
    write(args.outdir / 'inputs.json', {'config': cfg, 'versions': versions, 'source_sha256': sha(__file__),
          'real_DE_sha256': sha(real_de_path if real_de_path.exists() else cache / 'real_de.csv'),
          'panel_axes_sha256': sha(cache / 'panel_axes.npz'), 'native_real_bulk_sha256': sha(bulk_path),
          'reference_values_sha256': reference.reference_sha256 if reference is not None else 'from_prediction_cache',
          'bulk_method': 'native Polars group_by.mean, complete original row order, Float32, independent gene blocks',
          'BH_family_size': len(real_targets) * len(genes), 'real_expression_refit': False})
    results = []
    for candidate in cfg['candidates']:
        t0 = time.monotonic()
        with np.load(candidate['path'], allow_pickle=False) as z:
            targets = z['targets'].astype(str).tolist()
            if set(targets) != set(real_targets) or z['genes'].astype(str).tolist() != genes:
                raise ValueError('Frozen prediction/query axes differ')
            means = z[candidate.get('key', 'predicted_means')].copy()
        counts = [public_counts[t] for t in targets]
        if means.dtype != np.dtype('float32') or means.shape != (len(targets), len(genes)):
            raise ValueError('Frozen Float32 mean array required')
        name = candidate['id']
        frames = None
        if prediction_cache is not None:
            receipt = json.loads((prediction_cache / (name + '_prediction_cache.json')).read_text())
            de_path = prediction_cache / (name + '_pred_DE.parquet')
            bulk_path = prediction_cache / (name + '_native_bulk.npz')
            if (receipt['candidate'] != candidate or receipt['counts_sha256'] != cfg['counts_sha256']
                    or sha(de_path) != receipt['pred_DE_sha256'] or sha(bulk_path) != receipt['native_bulk_sha256']):
                raise ValueError('Frozen prediction cache identity differs')
            pred_de = pl.read_parquet(de_path)
            with np.load(bulk_path, allow_pickle=False) as z:
                pred_keys, pred_values = z['keys'].astype(str), z['values'].copy()
                if z['genes'].astype(str).tolist() != genes:
                    raise ValueError('Cached prediction bulk gene axis differs')
            diagnostics, seconds = receipt['diagnostics'], receipt['raw_map_seconds']
        else:
            frames, diagnostics, seconds = helper.parallel_raw(reference, means, counts, targets, cfg.get('workers', 4))
            pred_de = score.with_global_bh(frames, targets, genes)
            pred_keys, pred_values = predicted_bulk(means, targets, counts, control.X, block_genes)
        des = helper.official_DES(real_de, pred_de, targets)
        bulk = official_bulk_metrics(real_keys, real_values, pred_keys, pred_values, genes)
        per_target = {'DES': des, **bulk}
        raw = {metric: float(np.mean(list(values.values()))) for metric, values in per_target.items()}
        from cell_eval._score import score_agg_metrics
        columns = {'DES': 'overlap_at_N', 'PDS': 'discrimination_score_l1', 'MAE': 'mae'}
        aggregate = pl.DataFrame({'statistic': ['mean'], **{columns[k]: [raw[k]] for k in ANCHORS}})
        anchors = pl.DataFrame({'statistic': ['mean'], **{columns[k]: [v] for k, v in ANCHORS.items()}})
        native_score = score_agg_metrics(results_user=aggregate, results_base=anchors)
        overall = float(native_score.filter(pl.col('metric') == 'avg_score')['from_baseline'][0])
        pred_de.write_parquet(args.outdir / (name + '_pred_DE.parquet'))
        native_score.write_csv(args.outdir / (name + '_anchor_score.csv'))
        aggregate.write_csv(args.outdir / (name + '_agg_results.csv'))
        result = {'candidate': candidate, 'raw': raw, 'per_target': per_target,
                  'historical_anchor_score': overall, 'historical_anchors': ANCHORS,
                  'native_metric_functions': True, 'exact_constant_MWU_backend': True,
                  'bulk_reduction': 'native Polars group_by.mean in gene blocks',
                  'raw_map_seconds': seconds, 'elapsed_seconds': time.monotonic() - t0,
                  'diagnostics': diagnostics, 'full_H5AD_submission_pipeline': False,
                  'official_private_leaderboard_equivalence': False}
        write(args.outdir / (name + '.json'), result)
        results.append(result)
        print(json.dumps({'state': 'candidate_complete', 'candidate': name, 'raw': raw, 'historical_anchor_score': overall}), flush=True)
        del frames, pred_de
        gc.collect()
    summary = {'state': 'complete', 'candidates': results, 'elapsed_seconds': time.monotonic() - started,
               'peak_RSS_KiB': resource.getrusage(resource.RUSAGE_SELF).ru_maxrss}
    write(args.outdir / 'summary.json', summary)
    write(args.outdir / 'completion.json', {'exit_code': 0, 'summary_sha256': sha(args.outdir / 'summary.json')})


if __name__ == '__main__':
    main()
```
