## Supplementary reproducibility code and frozen predictions for "Centered cross-context perturbation transfer on the public 2025 Virtual Cell Challenge benchmark": score_frozen_meanflat_panel_process_v1.py

```
"""Opt in to spawn-based exact MWU transport around the unchanged v2 scorer.

All original scorer arguments are forwarded. Additional arguments:
  --process-reference-cache PATH [--process-workers 8]
The original v2 scorer still builds/checks its reference and runs its unchanged
BH, native bulk reduction, and official metrics. Only its target map is swapped.
"""
from __future__ import annotations
import argparse,hashlib,json,os,sys
from pathlib import Path
ROOT=Path(__file__).resolve().parents[1]
LIB=ROOT/'runs/training_meanflat_DES_prepare_v1/scripts'

def main():
    parser=argparse.ArgumentParser(add_help=False)
    parser.add_argument('--process-reference-cache',type=Path,required=True)
    parser.add_argument('--process-workers',type=int)
    args,remainder=parser.parse_known_args()
    manifest_path=args.process_reference_cache/'manifest.json'
    manifest=json.loads(manifest_path.read_text())
    sys.path[:0]=[str(ROOT/'scripts'),str(LIB)]
    import mwu_constant_reference_process_v1 as process_backend
    import score_training_meanflat_DES as helper
    import score_frozen_meanflat_panel_final_v2 as original
    output=Path(remainder[remainder.index('--outdir')+1])
    calls=[]
    def exact_process_map(reference,means,counts,targets,workers):
        if reference.reference_sha256!=manifest['reference_sha256'] or reference.genes!=manifest['genes'] or reference.n!=manifest['reference_cells'] or str(reference.dtype)!=manifest['dtype']:
            raise ValueError('Process cache differs from the v2 scorer actual reference')
        cpus=sorted(os.sched_getaffinity(0));actual_workers=args.process_workers or workers
        # Spawn workers inherit single-thread library settings. The already
        # initialized parent Polars pool and original bulk kernels stay intact.
        keys=('OMP_NUM_THREADS','OPENBLAS_NUM_THREADS','MKL_NUM_THREADS','POLARS_MAX_THREADS')
        previous={key:os.environ.get(key) for key in keys};os.environ.update({key:'1' for key in keys})
        try:frames,diagnostics,seconds=process_backend.parallel_raw(args.process_reference_cache,means,counts,targets,actual_workers,cpus)
        finally:
            for key,value in previous.items():
                if value is None:os.environ.pop(key,None)
                else:os.environ[key]=value
        calls.append(dict(workers=actual_workers,cpus=cpus,targets=len(targets),seconds=seconds))
        (output/'process_backend.json').write_text(json.dumps(dict(backend='spawn_exact_constant_MWU_v1',reference_cache=str(args.process_reference_cache),
            cache_manifest_sha256=hashlib.sha256(manifest_path.read_bytes()).hexdigest(),
            backend_sha256=hashlib.sha256(Path(process_backend.__file__).read_bytes()).hexdigest(),
            wrapper_sha256=hashlib.sha256(Path(__file__).read_bytes()).hexdigest(),calls=calls,math_changed=False),indent=2))
        return frames,diagnostics,seconds
    helper.parallel_raw=exact_process_map
    sys.argv=[str(Path(original.__file__)),*remainder]
    original.main()
if __name__=='__main__':main()
```
