## Supplementary reproducibility code and frozen predictions for "Centered cross-context perturbation transfer on the public 2025 Virtual Cell Challenge benchmark": dual_source_ridge.py

```
"""No-PCA, full-source masked linear dual ridge; no data acquisition or CV.

Fit-only statistics are the default. Frozen public source centers/spreads may
be provided explicitly, without using their PCA components. Every supervised
matrix is reduced to explicit H1 fit rows before values are inspected.
"""
from __future__ import annotations
from dataclasses import dataclass
import numpy as np

from .transfer import _ids, axis_hash
from .public_source_basis import resolve_targets, _matrix_hash


@dataclass
class SourceScaler:
    source_id: str
    scale: str
    genes: tuple
    center: np.ndarray
    spread: np.ndarray
    metadata: dict

    @classmethod
    def fit_only(cls,source,fit_source_ids):
        ids=_ids(fit_source_ids,"source scaler fit targets"); genes=tuple(sorted(source.genes))
        x,mask=source.aligned(ids,genes)
        count=mask.sum(axis=0)
        center=np.divide(np.where(mask,x,0).sum(axis=0),count,out=np.zeros(len(genes)),where=count>0)
        centered=np.where(mask,x-center,0)
        variance=np.divide((centered**2).sum(axis=0),count,out=np.zeros(len(genes)),where=count>0)
        keep=(count>=2)&(variance>1e-24)
        return cls(source.source_id,source.scale,tuple(g for g,ok in zip(genes,keep) if ok),center[keep],np.sqrt(variance[keep]),
                   dict(scope="h1_fit_only",fit_source_ids=list(ids),source_file_sha256=source.metadata.get("file_sha256"),
                        input_gene_axis_sha256=axis_hash(source.genes),H1_response_used=False,PCA_used=False))

    @classmethod
    def from_public_basis(cls,basis):
        if basis.metadata["scope"]!="all_public" or basis.metadata["H1_response_used"] is not False or basis.metadata["excluded_source_target_ids"]:
            raise ValueError("Only explicit main-track frozen public source scalers are reusable")
        return cls(basis.source_id,basis.source_scale,basis.genes,basis.center.copy(),basis.scale.copy(),
                   dict(scope="frozen_public",public_basis_input_sha256=basis.metadata["basis_input_sha256"],
                        public_basis_target_ids=list(basis.basis_target_ids),
                        source_file_sha256=basis.metadata["source_file_sha256"],input_gene_axis_sha256=basis.metadata["input_source_gene_axis_sha256"],
                        H1_response_used=False,PCA_used=False,public_components_ignored=True))

    def encode(self,source,ids):
        if source.source_id!=self.source_id or source.scale!=self.scale:
            raise ValueError("Scaler source identity/scale differs")
        if self.center.shape!=(len(self.genes),) or self.spread.shape!=self.center.shape or not np.isfinite(self.center).all() or not np.isfinite(self.spread).all() or np.any(self.spread<=0):
            raise ValueError("Invalid source scaler arrays")
        x,mask=source.aligned(ids,self.genes)
        # Center-only imputation defines the numerical feature map. It is not
        # an observed zero delta and never makes an absent source row available.
        design=np.where(mask,(x-self.center)/self.spread,0)
        return design,mask.any(axis=1)


class DualSourceRidgeExpert:
    def __init__(self,*,alpha=1.,shrinkage=.5,scaling_scope="h1_fit_only",track="main_external"):
        if not np.isfinite(alpha) or alpha<=0 or not np.isfinite(shrinkage) or not 0<=shrinkage<=1:
            raise ValueError("Finite positive alpha and legal shrinkage required")
        if scaling_scope not in ("h1_fit_only","frozen_public") or track not in ("main_external","strict_all_context_unseen"):
            raise ValueError("Declare scaler scope and availability track")
        if track=="strict_all_context_unseen" and scaling_scope=="frozen_public":
            raise ValueError("Main all-public scalers cannot be reused for strict unseen")
        self.alpha=alpha; self.shrinkage=shrinkage; self.scaling_scope=scaling_scope; self.track=track

    def fit(self,sources,response_target_ids,response_genes,response_deltas,*,fit_target_ids,
            authoritative_target_ids=None,excluded_target_ids=(),scalers=None):
        ids=_ids(response_target_ids,"response targets"); self.genes=_ids(response_genes,"response genes")
        self.fit_target_ids=_ids(fit_target_ids,"fit targets"); self.excluded=_ids(excluded_target_ids,"excluded targets")
        values=np.asarray(response_deltas)
        if values.shape!=(len(ids),len(self.genes)) or len(self.fit_target_ids)<2 or not set(self.fit_target_ids)<=set(ids):
            raise ValueError("Explicit response and fit axes differ")
        if set(self.fit_target_ids)&set(self.excluded):
            raise ValueError("Fit/excluded targets overlap")
        self.authoritative=authoritative_target_ids or {}
        if self.track=="strict_all_context_unseen" and (not self.excluded or ("TAZ" in self.excluded and not self.authoritative.get("TAZ"))):
            raise ValueError("Strict track requires frozen and resolved excluded targets")
        lookup={target:i for i,target in enumerate(ids)}
        y=np.asarray(values[[lookup[target] for target in self.fit_target_ids]],dtype=float).copy()
        if not np.isfinite(y).all():
            raise ValueError("Fit responses must be finite")
        self.common_response=y.mean(axis=0); self.models={}
        for source in sources:
            if source.source_id in self.models:
                raise ValueError("Duplicate source view ID")
            context=source.metadata.get("source_context",{})
            if not context.get("cell_line") or "time_days" not in context:
                raise ValueError("Declare experimental background to group correlated RAW/Z views")
            background=(str(context["cell_line"]),str(context["time_days"]))
            mapped=resolve_targets(source,self.fit_target_ids+self.excluded,self.authoritative)
            known=[mapped[t] for t in self.fit_target_ids if mapped[t] is not None]
            if len(known)!=len(set(known)):
                raise ValueError("Fit targets resolve to a duplicate source gene")
            if self.track=="strict_all_context_unseen" and any(mapped[t] is not None and mapped[t] in set(known) for t in self.excluded):
                raise ValueError("Strict fit/excluded targets resolve to the same source gene")
            if self.scaling_scope=="h1_fit_only":
                if scalers is not None:
                    raise ValueError("Fit-only scaling cannot silently take public scalers")
                scaler=SourceScaler.fit_only(source,tuple(sorted(known)))
            else:
                if scalers is None or source.source_id not in scalers:
                    raise ValueError("Explicit frozen public scaler required")
                scaler=scalers[source.source_id]
                if scaler.metadata.get("scope")!="frozen_public" or scaler.metadata.get("H1_response_used") is not False or scaler.metadata.get("PCA_used") is not False:
                    raise ValueError("Unreviewed frozen scaler scope")
                if scaler.metadata["source_file_sha256"]!=source.metadata.get("file_sha256") or scaler.metadata["input_gene_axis_sha256"]!=axis_hash(source.genes):
                    raise ValueError("Frozen scaler source provenance differs")
                scaler_ids=tuple(scaler.metadata["public_basis_target_ids"]); scaler_genes=tuple(sorted(source.genes))
                sx,sm=source.aligned(scaler_ids,scaler_genes)
                if _matrix_hash(scaler_ids,scaler_genes,sx,sm)!=scaler.metadata["public_basis_input_sha256"]:
                    raise ValueError("Frozen scaler numeric source inputs changed")
                del sx,sm
            encoded_ids=tuple(mapped[t] or "__missing__"+t for t in self.fit_target_ids)
            design,available=scaler.encode(source,encoded_ids)
            model=dict(enabled=False,scaler=scaler,mapping=mapped,background=background,scale=source.scale)
            self.models[source.source_id]=model
            if available.sum()<3:
                continue
            x,ys=design[available],y[available]
            feature_center=x.mean(axis=0); x=x-feature_center
            gram=x@x.T
            trace_scale=float(np.trace(gram)/len(x))
            if not np.isfinite(trace_scale) or trace_scale<=1e-24:
                continue
            kernel=gram/trace_scale
            ycenter=ys.mean(axis=0)
            dual=np.linalg.solve(kernel+self.alpha*len(x)*np.eye(len(x)),ys-ycenter)
            model.update(enabled=True,fit_design=x,feature_center=feature_center,trace_scale=trace_scale,
                         dual_coefficients=dual,offset=ycenter-self.common_response,
                         fit_available_targets=int(available.sum()),normalized_kernel_mean_diagonal=float(np.trace(kernel)/len(x)))
        return self

    def predict(self,sources,*,query_ids,genes=None,return_diagnostics=False):
        queries=_ids(query_ids,"query targets")
        if set(queries)&set(self.fit_target_ids):
            raise ValueError("Fit/query targets overlap")
        if self.track=="strict_all_context_unseen" and not set(queries)<=set(self.excluded):
            raise ValueError("Strict query was not frozen as excluded")
        sources_by_id={s.source_id:s for s in sources}
        if len(sources_by_id)!=len(sources):
            raise ValueError("Duplicate source view ID")
        backgrounds={}; view_counts=np.zeros(len(queries),dtype=int)
        for source_id,model in self.models.items():
            source=sources_by_id.get(source_id)
            if source is None:
                continue
            if source.scale!=model["scale"]:
                raise ValueError("Source scale changed after fitting")
            context=source.metadata.get("source_context",{})
            if (str(context.get("cell_line")),str(context.get("time_days")))!=model["background"]:
                raise ValueError("Experimental background changed after fitting")
            if not model["enabled"] or self.track=="strict_all_context_unseen":
                continue
            mapped=resolve_targets(source,queries,self.authoritative)
            fit_genes={model["mapping"][t] for t in self.fit_target_ids if model["mapping"][t] is not None}
            if any(mapped[t] is not None and mapped[t] in fit_genes for t in queries):
                raise ValueError("Query resolves to a fit gene")
            encoded=tuple(mapped[t] or "__missing__"+t for t in queries)
            x,available=model["scaler"].encode(source,encoded)
            cross=(x-model["feature_center"])@model["fit_design"].T/model["trace_scale"]
            delta=cross@model["dual_coefficients"]+model["offset"]
            total,count=backgrounds.setdefault(model["background"],(np.zeros((len(queries),len(self.genes))),np.zeros(len(queries),dtype=int)))
            total+=np.where(available[:,None],delta,0); count+=available; view_counts+=available
        total=np.zeros((len(queries),len(self.genes))); counts=np.zeros(len(queries),dtype=int)
        for values,available_views in backgrounds.values():
            present=available_views>0
            total+=np.divide(values,available_views[:,None],out=np.zeros_like(values),where=present[:,None]); counts+=present
        prediction=self.common_response+self.shrinkage*np.divide(total,counts[:,None],out=np.zeros_like(total),where=counts[:,None]>0)
        if genes is not None:
            genes=_ids(genes,"prediction genes"); lookup={g:i for i,g in enumerate(self.genes)}
            if not set(genes)<=set(lookup): raise ValueError("Unfitted response gene")
            prediction=prediction[:,[lookup[g] for g in genes]]
        report=dict(expert="full_source_linear_dual_ridge",PCA_used=False,scaling_scope=self.scaling_scope,track=self.track,
                    fit_target_ids=list(self.fit_target_ids),query_ids=list(queries),alpha=self.alpha,shrinkage=self.shrinkage,
                    kernel_normalization="fit mean diagonal = 1; solve K + alpha*n_available_fit I",
                    available_views_per_query=view_counts.tolist(),experimental_backgrounds_per_query=counts.tolist(),
                    RAW_GEMZ_independent_evidence=False,background_aggregation="equal available views within background, then equal available backgrounds",
                    source_fits={key:{"enabled":model["enabled"],"background":list(model["background"]),"scaler_scope":model["scaler"].metadata["scope"],
                                      "supported_genes":len(model["scaler"].genes),"kernel_trace_scale":model.get("trace_scale"),
                                      "fit_available_targets":model.get("fit_available_targets")} for key,model in self.models.items()},
                    fallback="current fit-target mean H1 delta",H1_query_truth_accepted=False,agent_used=False,official_score=False)
        return (prediction,report) if return_diagnostics else prediction
```
