## Supplementary reproducibility code and frozen predictions for "Centered cross-context perturbation transfer on the public 2025 Virtual Cell Challenge benchmark": final_training150_base_catalogue_v1.py

```
"""Final TRAIN-only base fit state and original-B OOF residual catalogue.

No outcome-file reader, query selection, metric, actor or launcher.
"""
from __future__ import annotations
import numpy as np
from .stage2_fit_dataset import SOURCE_ALPHA,SOURCE_SHRINKAGE,array_hash,ids,logical_hash
from .residual_transfer_diagnostic import SOURCE_IDS,CAP,fit_programs

def fit_base_states(*,fit_target_ids,fit_deltas,genes,control_mean,baseline_views):
    """Serialize precisely the unchanged source_prediction solve, without PCA."""
    targets=ids(fit_target_ids,'final fit targets');genes=ids(genes,'output genes')
    y=np.array(fit_deltas,dtype=np.float64,copy=True);control=np.array(control_mean,dtype=np.float64,copy=True)
    if y.shape!=(len(targets),len(genes)) or control.shape!=(len(genes),) or not np.isfinite(y).all() or not np.isfinite(control).all():
        raise ValueError('Finite physical final-fit target/gene/control arrays required')
    if SOURCE_ALPHA!=.001 or SOURCE_SHRINKAGE!=1. or tuple(v.expert_id for v in baseline_views)!=SOURCE_IDS or len({v.background for v in baseline_views})!=3:
        raise ValueError('Exactly the existing three independent original GEMZ contexts and fixed ridge settings')
    common=y.mean(axis=0);states=[]
    scope=logical_hash(dict(fit_ids=list(targets),genes=list(genes),fit_delta_sha256=array_hash(y),control_sha256=array_hash(control)))
    for view in baseline_views:
        x,known=view.slice(targets);design=np.empty((0,x.shape[1]),np.float64)
        center=np.zeros(x.shape[1],np.float64);response_center=common.copy();dual=np.empty((0,len(genes)),np.float64);trace=0.;enabled=False
        if known.sum()>=3:
            design=x[known];center=design.mean(axis=0);design=design-center;gram=design@design.T
            trace=float(np.trace(gram)/len(design))
            if np.isfinite(trace) and trace>1e-24:
                response=y[known];response_center=response.mean(axis=0)
                dual=np.linalg.solve(gram/trace+SOURCE_ALPHA*len(design)*np.eye(len(design)),response-response_center)
                enabled=True
            else:design=np.empty((0,x.shape[1]),np.float64)
        states.append(dict(source_id=view.expert_id,background=view.background,enabled=enabled,fit_available=known.copy(),
            feature_center=center,fit_design=design,response_center=response_center,dual_coefficients=dual,trace_scale=trace,
            public_feature_declaration=view.declaration,fit_scope_hash=scope,alpha=.001,shrinkage=1.,PCA_used=False))
    return dict(fit_targets=targets,genes=genes,control_mean=control,common_delta=common,states=states,
        fit_scope_hash=scope,fit_response_sha256=array_hash(y),model_family='original frozen-public fullfeature noPCA GEMZ3')

def predict_base_delta(*,base,query_ids,query_views):
    """Exact expert lower projection, available-context mean, final CAP order."""
    query=ids(query_ids,'external query identities')
    if set(query)&set(base['fit_targets']):raise ValueError('Final model is an inference asset for disjoint external targets')
    if tuple(v.expert_id for v in query_views)!=SOURCE_IDS:raise ValueError('Same source order required')
    genes=base['genes'];control=base['control_mean'];common=base['common_delta']
    total=np.zeros((len(query),len(genes)));count=np.zeros_like(total,dtype=np.int64)
    for state,view in zip(base['states'],query_views,strict=True):
        xq,available=view.slice(query)
        if xq.shape[1]!=len(state['feature_center']) or tuple(view.background)!=tuple(state['background']):
            raise ValueError('Frozen source coordinate/background identity changed')
        prediction=np.broadcast_to(common,total.shape).copy();valid=np.zeros(len(query),bool)
        if state['enabled']:
            prediction[available]=state['response_center']+((xq[available]-state['feature_center'])@state['fit_design'].T/state['trace_scale'])@state['dual_coefficients']
            valid=available.copy()
        mask=np.broadcast_to(valid[:,None],total.shape)
        projected=np.maximum(control+np.where(mask,prediction,common),0)-control
        total+=np.where(mask,projected,0);count+=mask
    delta=np.divide(total,count,out=np.broadcast_to(common,total.shape).copy(),where=count>0)
    return np.clip(control+delta,0,CAP)-control

def catalogue_from_original_B(*,fit_target_ids,genes,fit_deltas,control_mean,original_B_means):
    targets=ids(fit_target_ids,'all final catalogue donor identities');genes=ids(genes,'output genes')
    y=np.asarray(fit_deltas,dtype=np.float64);control=np.asarray(control_mean,dtype=np.float64);B=np.asarray(original_B_means)
    if B.dtype!=np.float64 or y.shape!=(len(targets),len(genes)) or B.shape!=y.shape or control.shape!=(len(genes),):
        raise ValueError('Original full F64 B means and TRAIN-only response/control axes required')
    if y.shape!=(len(targets),len(genes)) or B.shape!=y.shape or not np.isfinite(y).all() or not np.isfinite(B).all() or np.any((B<0)|(B>CAP)):
        raise ValueError('Finite original uncalibrated OOF B required')
    donor_oof=B-control
    residual=y-donor_oof
    programs=fit_programs(residual,targets,genes,rank=8,completion_iterations=2)
    trust_bound=.5*float(np.quantile(np.abs(programs['completion']).sum(axis=1),.9))
    return dict(fit_targets=targets,genes=genes,donor_oof=donor_oof,residual_values=residual,programs=programs,trust_bound_L1=trust_bound,
        original_B_means_sha256=array_hash(B),donor_oof_sha256=array_hash(donor_oof),residual_sha256=array_hash(residual),
        definition='donor_oof=original committed uncalibrated B means minus frozen C; residual=TRAIN delta minus donor_oof',
        oof_source='original disjoint F100-to-V50 B prediction covering all150 once',
        in_sample_residuals_used=False,genuine_or_calibrated_residuals_used=False,residual_center_added_back=False,
        actual_rank=programs['actual_rank'],basis_fit_target_count=len(targets),final_inference_asset_only=True)

def pack_base(base):
    arrays=dict(fit_targets=np.asarray(base['fit_targets']),genes=np.asarray(base['genes']),control_mean=base['control_mean'],common_delta=base['common_delta'])
    states=[]
    for i,state in enumerate(base['states']):
        fields=('fit_available','feature_center','fit_design','response_center','dual_coefficients')
        for field in fields:arrays[f'context{i}_{field}']=state[field]
        states.append({k:v for k,v in state.items() if k not in fields})
    meta=dict(model_family=base['model_family'],fit_scope_hash=base['fit_scope_hash'],fit_response_sha256=base['fit_response_sha256'],
        states=states,array_hashes={k:array_hash(v) for k,v in arrays.items()},control_projection_order='each expert lower projection; equal available contexts; final CAP clip')
    return arrays,meta

def unpack_base(arrays,metadata):
    if set(arrays)!=set(metadata['array_hashes']) or any(array_hash(v)!=metadata['array_hashes'][k] for k,v in arrays.items()):
        raise ValueError('Immutable serialized base field/hash identity required')
    states=[]
    for i,state in enumerate(metadata['states']):
        out=dict(state)
        for key in ('fit_available','feature_center','fit_design','response_center','dual_coefficients'):out[key]=arrays[f'context{i}_{key}'].copy()
        states.append(out)
    return dict(fit_targets=tuple(map(str,arrays['fit_targets'])),genes=tuple(map(str,arrays['genes'])),control_mean=arrays['control_mean'].copy(),
        common_delta=arrays['common_delta'].copy(),states=states,fit_scope_hash=metadata['fit_scope_hash'],
        fit_response_sha256=metadata['fit_response_sha256'],model_family=metadata['model_family'])
```
