## Supplementary reproducibility code and frozen predictions for "Centered cross-context perturbation transfer on the public 2025 Virtual Cell Challenge benchmark": genept_target_kernel.py

```
"""Identity-feature target kernel expert, independent of archive preparation.

This module fits arrays only. It has no outcome-file reader, selector, metric,
launcher or provider call. Callers must physically slice the current fit labels.
"""
from __future__ import annotations

from dataclasses import dataclass
import hashlib
import json
from pathlib import Path

import numpy as np


def array_hash(value):
    value = np.ascontiguousarray(value)
    digest = hashlib.sha256(str(value.dtype).encode()+str(value.shape).encode())
    digest.update(value.tobytes())
    return digest.hexdigest()


def file_hash(path):
    digest = hashlib.sha256()
    with Path(path).open('rb') as stream:
        for block in iter(lambda: stream.read(1 << 20), b''):
            digest.update(block)
    return digest.hexdigest()


def ids(values, name):
    values = tuple(values)
    if not values or any(type(v) is not str or not v for v in values) or len(set(values)) != len(values):
        raise ValueError('Require nonempty unique literal '+name)
    return values


@dataclass(frozen=True)
class GenePTTargetFeatures:
    representation_id: str
    targets: tuple[str, ...]
    values: np.ndarray
    available: np.ndarray
    metadata: dict

    def __post_init__(self):
        target_ids = ids(self.targets, 'feature target identities')
        values = np.array(self.values, dtype=np.float64, copy=True)
        mask = np.asarray(self.available)
        if (values.ndim != 2 or values.shape[0] != len(target_ids) or values.shape[1] < 1 or
                mask.dtype.kind != 'b' or mask.shape != (len(target_ids),) or
                not np.isfinite(values[mask]).all()):
            raise ValueError('Require aligned finite observed embedding vectors and explicit bool mask')
        # Unavailable placeholders are never read into model statistics.
        mask = mask.copy()
        values.setflags(write=False)
        mask.setflags(write=False)
        object.__setattr__(self, 'targets', target_ids)
        object.__setattr__(self, 'values', values)
        object.__setattr__(self, 'available', mask)
        object.__setattr__(self, 'metadata', dict(self.metadata))

    def slice(self, target_ids):
        target_ids = ids(target_ids, 'requested target identities')
        lookup = {target: row for row, target in enumerate(self.targets)}
        if any(target not in lookup for target in target_ids):
            raise ValueError('Requested target absent from the frozen explicit-mask feature axis')
        rows = [lookup[target] for target in target_ids]
        return self.values[rows].copy(), self.available[rows].copy()


def load_genept_target_features(root, manifest_path, *, expected_manifest_sha256, representation_id):
    """Load the frozen metadata-only NPZ; no aliases and no expression file."""
    root = Path(root).resolve()
    manifest_path = Path(manifest_path).resolve()
    if not manifest_path.is_relative_to(root) or file_hash(manifest_path) != expected_manifest_sha256:
        raise ValueError('Feature manifest location or pinned SHA differs')
    manifest = json.loads(manifest_path.read_text(encoding='utf-8-sig'))
    if (manifest['state'] != 'identity_features_prepared' or manifest['n_targets'] != 150 or
            manifest['H1_expression_file_opened'] is not False or manifest['performance_CV_started'] is not False or
            manifest['alias_expansion_applied'] is not False or manifest['representations_independent'] is not True):
        raise ValueError('Require independently prepared TRAINING150 identity features')
    representation = manifest['representations'][representation_id]
    path = (root/representation['output_file']).resolve()
    if not path.is_relative_to(root) or file_hash(path) != representation['output_sha256']:
        raise ValueError('Representation NPZ location or pinned SHA differs')
    with np.load(path, allow_pickle=False) as saved:
        if set(saved.files) != {'targets', 'values', 'available', 'representation_id'}:
            raise ValueError('Unexpected feature NPZ schema')
        if saved['representation_id'].ndim != 0 or saved['representation_id'].item() != representation_id:
            raise ValueError('Representation identity differs')
        targets = tuple(saved['targets'].tolist())
        if saved['targets'].ndim != 1 or saved['targets'].dtype.kind != 'U':
            raise ValueError('Require a literal Unicode target axis')
        block = GenePTTargetFeatures(representation_id, targets, saved['values'], saved['available'],
            dict(feature_manifest_sha256=expected_manifest_sha256,
                 feature_npz_sha256=representation['output_sha256'], source=manifest['source'],
                 identity_sha256=manifest['inputs']['identities_sha256'],
                 basis_scope='frozen external pretrained dictionary', alias_expansion_applied=False,
                 H1_expression_opened=False, source_alignment=representation['aligned_features']))
    if (len(block.targets) != 150 or block.values.shape[1] != representation['dimension'] or
            int(block.available.sum()) != representation['n_available']):
        raise ValueError('Feature dimensions or availability differ from manifest')
    return block


@dataclass(frozen=True)
class GenePTKernelPrediction:
    deltas: np.ndarray
    available: np.ndarray
    metadata: dict


class GenePTTargetKernelExpert:
    """Current-fit-only scaler/PCA, trace-normalized dual ridge and common fallback.

    alpha multiplies n_available_fit in the normalized kernel. A caller may
    independently register PCA rank/alpha/shrinkage; this class selects nothing.
    Shrinkage pulls the known-target response towards the all-fit common delta.
    """
    def __init__(self, *, alpha, shrinkage=1., pca_rank=None, standardize=True):
        if (not np.isfinite(alpha) or alpha <= 0 or not np.isfinite(shrinkage) or
                not 0 <= shrinkage <= 1 or type(standardize) is not bool or
                (pca_rank is not None and (type(pca_rank) is not int or pca_rank < 1))):
            raise ValueError('Require positive alpha, bounded shrinkage and declared rank/scaling')
        self.alpha, self.shrinkage = float(alpha), float(shrinkage)
        self.pca_rank, self.standardize = pca_rank, standardize
        self.fitted = False

    def fit(self, *, fit_target_ids, fit_deltas, genes, features):
        fit_ids = ids(fit_target_ids, 'fit target identities')
        genes = ids(genes, 'response gene identities')
        response = np.array(fit_deltas, dtype=np.float64, copy=True)
        if (not isinstance(features, GenePTTargetFeatures) or response.shape != (len(fit_ids), len(genes)) or
                not np.isfinite(response).all()):
            raise ValueError('Require physically sliced finite current-fit response matrix')
        x, known = features.slice(fit_ids)
        self.features, self.fit_ids, self.genes = features, fit_ids, genes
        self.common = response.mean(axis=0)
        self.enabled = False
        self.metadata = dict(representation_id=features.representation_id,
            fit_target_ids=list(fit_ids), available_fit_target_ids=[t for t, k in zip(fit_ids, known) if k],
            n_fit=len(fit_ids), n_available_fit=int(known.sum()), feature_metadata=features.metadata,
            fit_response_sha256=array_hash(response), fit_feature_sha256=array_hash(x[known]),
            alpha=self.alpha, shrinkage=self.shrinkage, requested_pca_rank=self.pca_rank,
            current_fit_only_scaler=True, current_fit_only_PCA=True, standardize=self.standardize,
            query_truth_accepted=False, unknown_fallback='all current fit-target common delta',
            kernel_convention='K / (trace(K) / n_available_fit); regularizer alpha*n_available_fit')
        if known.sum() >= 2:
            observed = x[known]
            self.center = observed.mean(axis=0)
            spread = observed.std(axis=0)
            self.divisor = np.where(spread > 1e-12, spread, 1.) if self.standardize else np.ones(x.shape[1])
            design = (observed-self.center)/self.divisor
            self.components = None
            if self.pca_rank is not None:
                eigenvalues, eigenvectors = np.linalg.eigh(design@design.T)
                order = np.argsort(eigenvalues)[::-1]
                eigenvalues, eigenvectors = eigenvalues[order], eigenvectors[:, order]
                keep = np.flatnonzero(eigenvalues > max(1e-12, float(eigenvalues[0])*1e-12))[:self.pca_rank]
                self.components = (design.T@eigenvectors[:, keep]/np.sqrt(eigenvalues[keep])).T
                design = design@self.components.T
            gram = design@design.T
            trace_scale = float(np.trace(gram)/len(design))
            if np.isfinite(trace_scale) and trace_scale > 1e-24:
                self.design, self.trace_scale = design, trace_scale
                self.ycenter = response[known].mean(axis=0)
                self.coefficients = np.linalg.solve(gram/trace_scale+self.alpha*len(design)*np.eye(len(design)),
                                                    response[known]-self.ycenter)
                self.enabled = True
                self.metadata.update(kernel_trace_scale=trace_scale,
                    normalized_kernel_mean_diagonal=float(np.trace(gram/trace_scale)/len(design)),
                    raw_materialized_regularizer=self.alpha*len(design)*trace_scale,
                    normalized_regularizer=self.alpha*len(design), effective_rank=int(np.linalg.matrix_rank(design)),
                    fit_center_sha256=array_hash(self.center), fit_spread_sha256=array_hash(self.divisor),
                    PCA_components_sha256=None if self.components is None else array_hash(self.components))
        self.metadata.update(enabled=self.enabled)
        self.fitted = True
        return self

    def predict(self, *, query_target_ids):
        if not self.fitted:
            raise RuntimeError('Fit expert on current-fit responses first')
        query_ids = ids(query_target_ids, 'query target identities')
        if set(query_ids)&set(self.fit_ids):
            raise ValueError('Query target identities overlap current fit')
        x, known = self.features.slice(query_ids)
        available = known & self.enabled
        prediction = np.broadcast_to(self.common, (len(query_ids), len(self.genes))).copy()
        if available.any():
            query = (x[available]-self.center)/self.divisor
            if self.components is not None:
                query = query@self.components.T
            predicted_known = self.ycenter+(query@self.design.T/self.trace_scale)@self.coefficients
            prediction[available] = self.common+self.shrinkage*(predicted_known-self.common)
        if not np.isfinite(prediction).all():
            raise ValueError('Prediction became nonfinite')
        return GenePTKernelPrediction(prediction, available,
            dict(self.metadata, query_target_ids=list(query_ids),
                 available_query_target_ids=[t for t, k in zip(query_ids, available) if k],
                 fallback_query_target_ids=[t for t, k in zip(query_ids, available) if not k],
                 query_feature_sha256=array_hash(x[known]), prediction_sha256=array_hash(prediction),
                 query_truth_accessed=False))
```
