## Supplementary reproducibility code and frozen predictions for "Centered cross-context perturbation transfer on the public 2025 Virtual Cell Challenge benchmark": public_source_basis.py

```
"""External-only source pretraining and explicit H1 fit-target calibration.

Independent of frozen E1 v1. All public source targets may define a basis in
the main track; no H1 response enters pretraining. RAW log1p population profiles
and signed GEM-Z profiles remain separate blocks. Missing is never zero effect.
"""
from __future__ import annotations

from dataclasses import dataclass
from pathlib import Path
from collections import Counter, defaultdict
from dataclasses import asdict
import json
import hashlib
import re

import numpy as np

from .transfer import (ExternalSource, RAW_SCALE, Z_SCALE, _index, _column,
                       _booleans, _ids, parse_construct, axis_hash, file_hash,
                       gene_mapping)


POLICY = "canonical source ENSG targets including explicitly ENSG-bound RefSeq NM constructs; finite-positive population constructs; per-gene finite control difference before weighted construct aggregation; RAW log1p profile or signed GEM-Z identity; missing NaN plus mask"


def parse_public_construct(label,row):
    record=parse_construct(label,row)
    if record is not None:
        return record
    match=re.fullmatch(r"\d+_(.+)_(NM_\d+(?:\.\d+)?(?:,NM_\d+(?:\.\d+)?)*)_(ENSG\d+|nan)",label)
    if not match:
        return None
    return dict(row=int(row),symbol=match[1],target_unit=match[2],target_unit_type="refseq_transcript",
                promoter=None,ensembl_id=match[3] if match[3]!="nan" else None)


def _header(handle, obj):
    import h5py
    labels, genes = _index(handle["obs"]), _index(handle["var"])
    symbols = tuple(map(str, _column(handle["var"], "gene_name")))
    audit = obj["metadata"]
    for key, values in (("target_axis_sha256", labels), ("gene_axis_sha256", genes), ("gene_symbol_axis_sha256", symbols)):
        if axis_hash(values) != audit[key]:
            raise ValueError(f"Frozen source axis changed: {key}")
    matrix = handle["X"]
    if not isinstance(matrix, h5py.Dataset) or matrix.dtype.kind != "f" or matrix.shape != tuple(audit["shape"]):
        raise ValueError("Require audited dense floating point population profiles")
    controls = _booleans(_column(handle["obs"], "core_control"))
    if int(controls.sum()) != audit["core_control_rows"]:
        raise ValueError("Core-control identity differs")
    counts = np.asarray(_column(handle["obs"], "num_cells_filtered"), dtype=float)
    if counts.shape != (len(labels),):
        raise ValueError("Population count axis differs")
    eligible = np.isfinite(counts) & (counts > 0)
    parsed, excluded = {}, []
    for row, label in enumerate(labels):
        if controls[row]:
            continue
        record = parse_public_construct(label, row)
        if record is None:
            excluded.append(dict(row=row, label=label, reason="noncore_non_targeting_control" if label.endswith("_non-targeting_non-targeting_non-targeting") else "unparsed_noncontrol_construct"))
        elif not record["ensembl_id"] or not re.fullmatch(r"ENSG\d+", record["ensembl_id"]):
            excluded.append(dict(**record, reason="missing_canonical_target_ensembl"))
        elif not eligible[row]:
            excluded.append(dict(**record, reason="invalid_population_cell_count"))
        else:
            parsed[row] = record
    if not np.any(controls & eligible):
        raise ValueError("No finite-positive population core control")
    # Every legacy H1 matched construct must retain exactly its audited identity.
    for item in audit.get("matched_target_constructs", []):
        row = int(item["row"])
        actual = parse_construct(labels[row], row)
        if actual is None or any(actual[k] != item.get(k) for k in actual):
            raise ValueError("Legacy audited construct identity changed")
    return labels, genes, symbols, controls, counts, eligible, parsed, excluded


def _verify_file(path, obj):
    if obj.get("is_singlecell") is not False or obj["source_scale"] not in (RAW_SCALE, Z_SCALE):
        raise ValueError("Require declared external population source scale")
    if path.stat().st_size != obj["size"] or file_hash(path, "md5") != obj["md5"]:
        raise ValueError("Full publisher bytes/MD5 differ")


def audit_full_source(path, obj, *, h1_expression_symbols=(), chunk_rows=256):
    """Read all external X rows, never H1 expression; return durable audit data."""
    import h5py
    path = Path(path)
    _verify_file(path, obj)
    with h5py.File(path, "r") as handle:
        labels, genes, symbols, controls, counts, eligible, parsed, excluded = _header(handle, obj)
        per_gene_missing = np.zeros(len(genes), dtype=np.int64)
        control_finite = np.zeros(len(genes), dtype=np.int64)
        construct_finite = np.zeros(len(genes), dtype=np.int64)
        control_missing = construct_missing = invalid_or_unparsed_missing = negative = finite_zero = 0
        affected_rows = []
        minimum, maximum = np.inf, -np.inf
        for start in range(0, len(labels), chunk_rows):
            values = np.asarray(handle["X"][start:start+chunk_rows], dtype=float)
            finite = np.isfinite(values)
            per_gene_missing += (~finite).sum(axis=0)
            measured = values[finite]
            if len(measured):
                minimum, maximum = min(minimum, float(measured.min())), max(maximum, float(measured.max()))
            negative += int(np.sum(measured < 0)); finite_zero += int(np.sum(measured == 0))
            for offset, missing in enumerate((~finite).sum(axis=1)):
                row = start+offset
                if controls[row] and eligible[row]:
                    control_finite += finite[offset]
                elif row in parsed:
                    construct_finite += finite[offset]
                if not missing:
                    continue
                if controls[row] and eligible[row]:
                    control_missing += int(missing)
                elif row in parsed:
                    construct_missing += int(missing)
                else:
                    invalid_or_unparsed_missing += int(missing)
                affected_rows.append(dict(row=row, nonfinite=int(missing), eligible_construct=row in parsed,
                                          eligible_control=bool(controls[row] and eligible[row])))
        by_symbol, by_gene = defaultdict(set), defaultdict(set)
        for item in parsed.values():
            by_symbol[item["symbol"]].add(item["ensembl_id"])
            by_gene[item["ensembl_id"]].add(item["symbol"])
        gene_target_counts = Counter(item["ensembl_id"] for item in parsed.values())
        mapping, overlap = gene_mapping(genes, symbols, h1_expression_symbols)
        if obj["source_scale"] == RAW_SCALE and negative:
            raise ValueError("RAW population source has negative finite expression")
        return dict(source_id=obj["id"], source_scale=obj["source_scale"], source_file_sha256=file_hash(path),
                    publisher_md5=obj["md5"], bytes=path.stat().st_size, shape=list(handle["X"].shape),
                    target_axis_sha256=axis_hash(labels), expression_ensembl_axis_sha256=axis_hash(genes),
                    expression_symbol_axis_sha256=axis_hash(symbols), obs_columns=list(handle["obs"].keys()),
                    original_core_controls=int(controls.sum()), eligible_core_controls=int((controls & eligible).sum()),
                    eligible_constructs=len(parsed), eligible_canonical_target_genes=len(gene_target_counts),
                    canonical_target_axis_sha256=axis_hash(sorted(gene_target_counts)),
                    canonical_target_ids=sorted(gene_target_counts), multiple_construct_targets=sum(v > 1 for v in gene_target_counts.values()),
                    symbol_to_multiple_target_ensembl={key:sorted(value) for key,value in by_symbol.items() if len(value)>1},
                    target_ensembl_to_multiple_symbols={key:sorted(value) for key,value in by_gene.items() if len(value)>1},
                    excluded_rows=excluded, excluded_reason_counts=dict(Counter(item["reason"] for item in excluded)),
                    invalid_core_control_rows=np.flatnonzero(controls & ~eligible).tolist(),
                    invalid_population_count_rows=np.flatnonzero(~eligible).tolist(),
                    invalid_population_count_nonfinite=int((~np.isfinite(counts)).sum()),
                    invalid_population_count_nonpositive=int(np.sum(np.isfinite(counts)&(counts<=0))),
                    eligible_refseq_constructs=sum(item["target_unit_type"]=="refseq_transcript" for item in parsed.values()),
                    full_matrix_nonfinite=int(per_gene_missing.sum()), eligible_core_control_nonfinite=control_missing,
                    eligible_construct_nonfinite=construct_missing, excluded_row_nonfinite=invalid_or_unparsed_missing,
                    source_gene_nonfinite_counts={genes[i]:int(v) for i,v in enumerate(per_gene_missing) if v},
                    source_genes_without_finite_eligible_control=[g for g,n in zip(genes,control_finite) if not n],
                    source_genes_without_finite_eligible_construct=[g for g,n in zip(genes,construct_finite) if not n],
                    genes_with_legal_control_and_construct=int(np.sum((control_finite>0)&(construct_finite>0))),
                    affected_rows=affected_rows, finite_negative_values=negative, finite_zero_values=finite_zero,
                    finite_value_min=minimum, finite_value_max=maximum, H1_expression_overlap=overlap,
                    raw_and_normalized_profiles_not_singlecell=True, missingness_causes_inferred=False,
                    H1_expression_opened=False, validation_test_outcomes_accessed=False,
                    policy=POLICY)


def load_public_source(path, obj, *, chunk_rows=256):
    """Aggregate every eligible canonical ENSG target on all source genes.

    Memory is bounded by two target×gene accumulators, not by the single-cell
    dataset. This function must run remotely for real publisher matrices.
    """
    import h5py
    path = Path(path)
    _verify_file(path, obj)
    with h5py.File(path, "r") as handle:
        labels, genes, symbols, controls, counts, eligible, parsed, excluded = _header(handle, obj)
        targets = tuple(sorted({item["ensembl_id"] for item in parsed.values()}))
        positions = {target:i for i,target in enumerate(targets)}
        numerator, denominator = np.zeros(len(genes)), np.zeros(len(genes))
        for row in np.flatnonzero(controls & eligible):
            value = np.asarray(handle["X"][row], dtype=float)
            finite = np.isfinite(value)
            if obj["source_scale"] == RAW_SCALE:
                if np.any(value[finite] < 0):
                    raise ValueError("RAW source contains negative expression")
                value = np.log1p(value)
            numerator += np.where(finite, value, 0) * counts[row]
            denominator += finite * counts[row]
        reference = np.divide(numerator, denominator, out=np.full(len(genes),np.nan), where=denominator>0)
        numerator = np.zeros((len(targets),len(genes)))
        denominator = np.zeros_like(numerator)
        target_symbols = defaultdict(set)
        for start in range(0,len(labels),chunk_rows):
            values = np.asarray(handle["X"][start:start+chunk_rows],dtype=float)
            for offset,value in enumerate(values):
                row=start+offset
                if row not in parsed:
                    continue
                finite=np.isfinite(value) & np.isfinite(reference)
                if obj["source_scale"] == RAW_SCALE:
                    if np.any(value[np.isfinite(value)]<0):
                        raise ValueError("RAW source contains negative expression")
                    value=np.log1p(value)
                delta=np.zeros(len(genes))
                np.subtract(value,reference,out=delta,where=finite)
                item=parsed[row]; position=positions[item["ensembl_id"]]
                numerator[position] += delta * counts[row]
                denominator[position] += finite * counts[row]
                target_symbols[item["ensembl_id"]].add(item["symbol"])
        mask=denominator>0
        delta=np.divide(numerator,denominator,out=np.full_like(numerator,np.nan),where=mask)
    metadata=dict(policy=POLICY, file_sha256=file_hash(path), file_md5=obj["md5"], file_id=obj["file_id"],
                  transform="log1p" if obj["source_scale"]==RAW_SCALE else "identity_gem_z",
                  target_symbols={key:sorted(value) for key,value in target_symbols.items()},
                  original_core_control_rows=np.flatnonzero(controls).tolist(), eligible_core_control_rows=np.flatnonzero(controls & eligible).tolist(),
                  excluded_rows=excluded, source_expression_symbols=list(symbols),
                  source_context=dict(cell_line=obj["cell_line"],time_days=obj["time_days"]),
                  source_clipping_applied=False, integerization_applied=False, H1_expression_opened=False,
                  h1_perturbed_outcomes_accessed=False, validation_test_outcomes_accessed=False)
    return ExternalSource(obj["id"],targets,genes,delta,mask,obj["source_scale"],metadata)


@dataclass(frozen=True)
class BasisConfig:
    rank: int = 64
    seed: int = 20251013
    oversampling: int = 8
    power_iterations: int = 2
    completion_iterations: int = 2
    singular_shrink_fraction: float = 0.02
    encode_ridge: float = 1e-6
    min_gene_observations: int = 2

    def __post_init__(self):
        if self.rank<1 or self.oversampling<0 or self.power_iterations<0 or self.completion_iterations<0 or self.min_gene_observations<2:
            raise ValueError("Invalid basis dimensions/iterations")
        if not np.isfinite(self.singular_shrink_fraction) or not 0<=self.singular_shrink_fraction<1 or not np.isfinite(self.encode_ridge) or self.encode_ridge<=0:
            raise ValueError("Invalid denoising/encoding regularization")


def _svd(matrix, config):
    rank=min(config.rank,min(matrix.shape))
    if not rank or not np.any(matrix):
        return np.zeros((len(matrix),0)),np.zeros(0),np.zeros((0,matrix.shape[1]))
    width=min(rank+config.oversampling,min(matrix.shape))
    omega=np.random.default_rng(config.seed).normal(size=(matrix.shape[1],width))
    q=np.linalg.qr(matrix@omega,mode="reduced")[0]
    for _ in range(config.power_iterations):
        z=np.linalg.qr(matrix.T@q,mode="reduced")[0]
        q=np.linalg.qr(matrix@z,mode="reduced")[0]
    u,s,v=np.linalg.svd(q.T@matrix,full_matrices=False)
    keep=np.flatnonzero(s>max(1e-12,float(s[0])*1e-10))[:rank]
    u,s,v=(q@u[:,keep]),s[keep],v[keep]
    # Sorted source gene coordinates make the tie convention reproducible.
    for index,row in enumerate(v):
        if row[np.argmax(np.abs(row))]<0:
            v[index]*=-1; u[:,index]*=-1
    return u,s,v


def _matrix_hash(ids,genes,values,mask):
    digest=hashlib.sha256((axis_hash(ids)+axis_hash(genes)).encode())
    for start in range(0,len(values),64):
        m=mask[start:start+64]
        digest.update(np.asarray(np.where(m,values[start:start+64],0),dtype="<f8").tobytes())
        digest.update(m.astype(np.uint8).tobytes())
    return digest.hexdigest()


class SourceBasis:
    """Frozen source-only basis; missing rows encoded by observed-feature LS."""
    def __init__(self, config=None):
        self.config=config if isinstance(config,BasisConfig) else BasisConfig(**(config or {}))

    def fit(self, source, *, basis_target_ids=None, excluded_target_ids=(), scope="all_public"):
        if scope not in ("all_public","h1_fit_only"):
            raise ValueError("Explicit basis scope required")
        if scope=="h1_fit_only" and basis_target_ids is None:
            raise ValueError("H1-fit basis requires its exact source target list")
        excluded=set(_ids(excluded_target_ids,"excluded source targets"))
        requested=_ids(source.target_ids if basis_target_ids is None else basis_target_ids,"basis source targets")
        if not set(requested)<=set(source.target_ids):
            raise ValueError("Unknown basis source target")
        ids=tuple(sorted(set(requested)-excluded)); genes=tuple(sorted(source.genes))
        x,mask=source.aligned(ids,genes)
        used_rows=mask.any(axis=1)
        x,mask=x[used_rows],mask[used_rows]
        self.basis_target_ids=tuple(target for target,used in zip(ids,used_rows) if used)
        count=mask.sum(axis=0)
        center=np.divide(np.where(mask,x,0).sum(axis=0),count,out=np.zeros(len(genes)),where=count>0)
        centered=np.where(mask,x-center,0)
        var=np.divide((centered**2).sum(axis=0),count,out=np.zeros(len(genes)),where=count>0)
        supported=(count>=self.config.min_gene_observations)&(var>1e-24)
        self.genes=tuple(g for g,ok in zip(genes,supported) if ok)
        self.center=center[supported]; self.scale=np.sqrt(var[supported])
        design=centered[:,supported]/self.scale
        observed=mask[:,supported]
        completed=design.copy()
        for _ in range(self.config.completion_iterations):
            u,s,v=_svd(completed,self.config)
            shrunk=np.maximum(s-self.config.singular_shrink_fraction*(s[0] if len(s) else 0),0)
            reconstructed=(u*shrunk)@v
            completed=np.where(observed,design,reconstructed)
        _,s,self.components=_svd(completed,self.config)
        self.gains=np.maximum(s-self.config.singular_shrink_fraction*(s[0] if len(s) else 0),0)/s if len(s) else s
        self.source_id,self.source_scale=source.source_id,source.scale
        self.metadata=dict(scope=scope, basis_target_ids=list(self.basis_target_ids), excluded_source_target_ids=sorted(excluded),
                           source_id=source.source_id, source_scale=source.scale, source_file_sha256=source.metadata.get("file_sha256"),
                           input_source_gene_axis_sha256=axis_hash(source.genes), basis_gene_axis_sha256=axis_hash(self.genes),
                           available_basis_targets=len(self.basis_target_ids), supported_genes=len(self.genes),
                           actual_rank=len(s), H1_response_used=False, missing_initialization="centered mean only for latent completion; not observed zero",
                           algorithm="seeded randomized source SVD; missing-only soft singular completion; observed-feature ridge encoding")
        self.metadata["basis_input_sha256"]=_matrix_hash(self.basis_target_ids,genes,x,mask)
        self.metadata["config"]=asdict(self.config)
        return self

    def save(self,path):
        path=Path(path)
        if path.exists():
            raise FileExistsError("Frozen source basis cannot be overwritten")
        path.parent.mkdir(parents=True,exist_ok=True)
        np.savez_compressed(path,genes=np.asarray(self.genes),center=self.center,scale=self.scale,
                            components=self.components,gains=self.gains,metadata=np.asarray(json.dumps(self.metadata,allow_nan=False)))

    @classmethod
    def read(cls,path):
        with np.load(path,allow_pickle=False) as data:
            metadata=json.loads(str(data["metadata"]))
            basis=cls(metadata["config"])
            basis.genes=tuple(map(str,data["genes"])); basis.center=data["center"].copy(); basis.scale=data["scale"].copy()
            basis.components=data["components"].copy(); basis.gains=data["gains"].copy()
        basis.metadata=metadata; basis.source_id=metadata["source_id"]; basis.source_scale=metadata["source_scale"]
        basis.basis_target_ids=tuple(metadata["basis_target_ids"])
        if basis.components.shape!=(len(basis.gains),len(basis.genes)) or basis.center.shape!=(len(basis.genes),) or basis.scale.shape!=basis.center.shape:
            raise ValueError("Frozen basis array axes differ")
        if not all(np.isfinite(value).all() for value in (basis.center,basis.scale,basis.components,basis.gains)) or np.any(basis.scale<=0):
            raise ValueError("Frozen basis has invalid numerical parameters")
        return basis

    def encode(self, source, *, target_ids):
        if source.source_id!=self.source_id or source.scale!=self.source_scale:
            raise ValueError("Source identity/scale differs from frozen basis")
        ids=_ids(target_ids,"encoded source target IDs")
        x,mask=source.aligned(ids,self.genes)
        available=mask.any(axis=1) & (len(self.gains)>0)
        scores=np.zeros((len(ids),len(self.gains)))
        for row in np.flatnonzero(available):
            selected=mask[row]
            components=self.components[:,selected]
            values=(x[row,selected]-self.center[selected])/self.scale[selected]
            scores[row]=np.linalg.solve(components@components.T+self.config.encode_ridge*np.eye(len(self.gains)),components@values)*self.gains
        return scores,available


def resolve_targets(source, targets, authoritative=None):
    """Unique exact source symbol or declared canonical ENSG; never alias TAZ."""
    by_symbol=defaultdict(set)
    for ens,symbols in source.metadata.get("target_symbols",{}).items():
        for symbol in symbols:
            by_symbol[symbol].add(ens)
    output={}
    for target in _ids(targets,"H1 target IDs"):
        explicit=(authoritative or {}).get(target)
        if explicit is not None:
            if not re.fullmatch(r"ENSG\d+",str(explicit)):
                raise ValueError("Authoritative target identity must be canonical ENSG")
            ens=explicit
        elif target=="TAZ":
            ens=None
        elif target in source.target_ids:
            ens=target
        elif len(by_symbol.get(target,()))==1:
            ens=next(iter(by_symbol[target]))
        else:
            ens=None
        output[target]=ens if ens in source.target_ids else None
    return output


class PublicBasisTransferExpert:
    """Same explicit fit/predict contract as E1, with full source-gene features.

    fit receives physically filtered H1 responses when possible; if a wider
    response array is provided, its other values are not inspected. There is no
    truth argument to predict. Public basis and fit-only basis share this API.
    """
    def __init__(self, *, basis_config=None, scope="all_public", track="main_external", alpha=1., shrinkage=.5):
        if scope not in ("all_public","h1_fit_only") or track not in ("main_external","strict_all_context_unseen"):
            raise ValueError("Declare source scope and target-availability track")
        if not np.isfinite(alpha) or alpha<=0 or not np.isfinite(shrinkage) or not 0<=shrinkage<=1:
            raise ValueError("Finite positive ridge and legal shrinkage required")
        self.basis_config=basis_config; self.scope=scope; self.track=track; self.alpha=alpha; self.shrinkage=shrinkage

    def fit(self,sources,response_target_ids,response_genes,response_deltas,*,fit_target_ids,
            authoritative_target_ids=None,excluded_target_ids=(),bases=None):
        ids=_ids(response_target_ids,"response targets"); self.genes=_ids(response_genes,"response genes")
        self.fit_target_ids=_ids(fit_target_ids,"fit targets"); self.excluded=_ids(excluded_target_ids,"excluded H1 targets")
        if set(self.fit_target_ids)&set(self.excluded):
            raise ValueError("Fit and excluded targets overlap")
        if self.track=="strict_all_context_unseen" and not self.excluded:
            raise ValueError("Strict track needs frozen heldout target identities before pretraining")
        if self.track=="strict_all_context_unseen" and "TAZ" in self.excluded and not (authoritative_target_ids or {}).get("TAZ"):
            raise ValueError("Strict TAZ exclusion needs authoritative H1 Ensembl identity")
        values=np.asarray(response_deltas)
        if values.shape!=(len(ids),len(self.genes)) or not set(self.fit_target_ids)<=set(ids) or len(self.fit_target_ids)<2:
            raise ValueError("Explicit response/fit axes differ")
        lookup={target:i for i,target in enumerate(ids)}
        y=np.asarray(values[[lookup[t] for t in self.fit_target_ids]],dtype=float).copy()
        if not np.isfinite(y).all():
            raise ValueError("Fit responses must be finite")
        self.common_response=y.mean(axis=0); self.models={}; self.authoritative=authoritative_target_ids or {}
        for source in sources:
            if source.source_id in self.models:
                raise ValueError("Separate source blocks need unique IDs")
            mapped=resolve_targets(source,self.fit_target_ids+self.excluded,self.authoritative)
            if self.track=="strict_all_context_unseen":
                aliases=defaultdict(set)
                for ens,symbols in source.metadata.get("target_symbols",{}).items():
                    for symbol in symbols: aliases[symbol].add(ens)
                if any(mapped[t] is None and len(aliases[t])>1 for t in self.excluded):
                    raise ValueError("Strict excluded target has unresolved ambiguous source Ensembl identity")
            basis_ids=tuple(sorted({mapped[t] for t in self.fit_target_ids if mapped[t] is not None})) if self.scope=="h1_fit_only" else None
            excluded_source=tuple(sorted({mapped[t] for t in self.excluded if mapped[t] is not None})) if self.track=="strict_all_context_unseen" else ()
            mapped_fit=[mapped[t] for t in self.fit_target_ids if mapped[t] is not None]
            if len(mapped_fit)!=len(set(mapped_fit)):
                raise ValueError("Several fit targets resolve to the same source gene identity")
            if bases is None:
                basis=SourceBasis(self.basis_config).fit(source,basis_target_ids=basis_ids,excluded_target_ids=excluded_source,scope=self.scope)
            else:
                basis=bases[source.source_id]
                expected_ids=tuple(sorted(set(source.target_ids if basis_ids is None else basis_ids)-set(excluded_source)))
                # A prepared public source keeps every eligible target; missing
                # target rows, if supplied synthetically, must be excluded too.
                expected_genes=tuple(sorted(source.genes))
                x,m=source.aligned(expected_ids,expected_genes)
                used=m.any(axis=1); expected_ids=tuple(t for t,valid in zip(expected_ids,used) if valid)
                content_hash=_matrix_hash(expected_ids,expected_genes,x[used],m[used])
                if (basis.source_id!=source.source_id or basis.source_scale!=source.scale or basis.metadata["scope"]!=self.scope or
                    basis.basis_target_ids!=expected_ids or basis.metadata["excluded_source_target_ids"]!=sorted(excluded_source) or
                    basis.metadata["source_file_sha256"]!=source.metadata.get("file_sha256") or
                    basis.metadata["basis_input_sha256"]!=content_hash or
                    basis.metadata["input_source_gene_axis_sha256"]!=axis_hash(source.genes) or
                    basis.metadata["config"]!=asdict(BasisConfig(**(self.basis_config or {})) if not isinstance(self.basis_config,BasisConfig) else self.basis_config)):
                    raise ValueError("Frozen public basis scope/input/config differs")
            encoded_ids=tuple(mapped[t] or "__missing__"+t for t in self.fit_target_ids)
            scores,available=basis.encode(source,target_ids=encoded_ids)
            model=dict(basis=basis,enabled=False,scale=source.scale,mapping=mapped)
            self.models[source.source_id]=model
            if available.sum()<3:
                continue
            x,ys=scores[available],y[available]
            center=x.mean(axis=0); spread=x.std(axis=0); spread=np.where(spread>1e-12,spread,1)
            design=(x-center)/spread; ycenter=ys.mean(axis=0)
            coefficients=np.linalg.solve(design.T@design+self.alpha*len(x)*np.eye(design.shape[1]),design.T@(ys-ycenter))
            model.update(enabled=True,center=center,spread=spread,coefficients=coefficients,offset=ycenter-self.common_response)
        return self

    def predict(self,sources,*,query_ids,genes=None,return_diagnostics=False):
        query_ids=_ids(query_ids,"query targets")
        if set(query_ids)&set(self.fit_target_ids):
            raise ValueError("Query and fit targets overlap")
        if self.track=="strict_all_context_unseen" and not set(query_ids)<=set(self.excluded):
            raise ValueError("Strict query was not excluded before public pretraining")
        by_id={source.source_id:source for source in sources}
        if len(by_id)!=len(sources):
            raise ValueError("Duplicate source identity")
        total=np.zeros((len(query_ids),len(self.genes))); contributors=np.zeros(len(query_ids),dtype=int)
        for source_id,model in self.models.items():
            source=by_id.get(source_id)
            if source is None:
                continue
            if source.scale!=model["scale"]:
                raise ValueError("Source scale changed after calibration")
            if not model["enabled"] or self.track=="strict_all_context_unseen":
                continue
            mapped=resolve_targets(source,query_ids,self.authoritative)
            fit_source_ids={model["mapping"][t] for t in self.fit_target_ids if model["mapping"][t] is not None}
            if any(mapped[t] is not None and mapped[t] in fit_source_ids for t in query_ids):
                raise ValueError("Query resolves to a calibrated fit gene identity")
            scores,available=model["basis"].encode(source,target_ids=tuple(mapped[t] or "__missing__"+t for t in query_ids))
            delta=((scores-model["center"])/model["spread"])@model["coefficients"]+model["offset"]
            total+=np.where(available[:,None],delta,0); contributors+=available
        prediction=self.common_response+self.shrinkage*np.divide(total,contributors[:,None],out=np.zeros_like(total),where=contributors[:,None]>0)
        if genes is not None:
            genes=_ids(genes,"prediction genes"); lookup={g:i for i,g in enumerate(self.genes)}
            if not set(genes)<=set(lookup):
                raise ValueError("Prediction expression gene not fitted")
            prediction=prediction[:,[lookup[g] for g in genes]]
        diagnostic=dict(scope=self.scope,track=self.track,fit_target_ids=list(self.fit_target_ids),query_ids=list(query_ids),
                        source_contributors=contributors.tolist(),fallback="fit-target mean H1 delta",
                        basis_scopes={key:model["basis"].metadata for key,model in self.models.items()},
                        H1_query_truth_accepted=False,agent_used=False,official_score=False)
        return (prediction,diagnostic) if return_diagnostics else prediction
```
