## Supplementary reproducibility code and frozen predictions for "Centered cross-context perturbation transfer on the public 2025 Virtual Cell Challenge benchmark": residual_transfer_diagnostic.py

```
"""Scoped residual memory diagnostic. No outcome-file reader or agent call."""
from __future__ import annotations
import numpy as np
from .stage2_fit_dataset import ids,array_hash,logical_hash,crossfit_splits,source_prediction,validate_expert

CAP=float(np.nextafter(np.float32(15),np.float32(0)))
SOURCE_IDS=("K562_gwps_normalized_bulk_01","rpe1_normalized_bulk_01","K562_essential_normalized_bulk_01")

def gemz_baseline(*,views,fit_ids,fit_deltas,query_ids,genes,control):
    if tuple(v.expert_id for v in views)!=SOURCE_IDS or len({v.background for v in views})!=3:
        raise ValueError("Exactly three independent registered GEMZ backgrounds required")
    total=np.zeros((len(query_ids),len(genes)));count=np.zeros_like(total,dtype=np.int64);records=[]
    scope=logical_hash(dict(fit_ids=list(fit_ids),genes=list(genes),fit_delta_sha256=array_hash(fit_deltas),control_sha256=array_hash(control)))
    for view in views:
        r=source_prediction(view,fit_ids=fit_ids,fit_deltas=fit_deltas,query_ids=query_ids,genes=genes,control=control,fit_scope_hash=scope)
        d,m,_,_,meta=validate_expert(r,fit_ids=fit_ids,fit_deltas=fit_deltas,query_ids=query_ids,genes=genes,control=control,fit_scope_hash=scope)
        total+=np.where(m,d,0);count+=m;records.append(meta)
    common=np.broadcast_to(fit_deltas.mean(0),total.shape).copy()
    delta=np.divide(total,count,out=common,where=count>0)
    return np.clip(control+delta,0,CAP)-control,records

def fit_programs(residuals,donors,genes,*,rank=8,completion_iterations=2):
    """Finite donor-centered residuals; missing-only SVD completion, explicit mask."""
    residuals=np.asarray(residuals,float)
    if residuals.shape!=(len(donors),len(genes)) or not np.isfinite(residuals).all():raise ValueError("Finite current-fit residual coordinates required")
    mask=np.ones(residuals.shape,bool);lookup={g:i for i,g in enumerate(genes)}
    for i,t in enumerate(donors):
        if t!="TAZ" and t in lookup:mask[i,lookup[t]]=False
    values=np.where(mask,residuals,np.nan);count=mask.sum(0)
    center=np.divide(np.where(mask,values,0).sum(0),count,out=np.zeros(len(genes)),where=count>0)
    observed_centered=np.where(mask,values-center,0);completed=observed_centered.copy()
    maxrank=min(rank,len(donors)-1,len(genes))
    def basis(x):
        _,s,v=np.linalg.svd(x,full_matrices=False);tol=np.finfo(float).eps*max(x.shape)*(s[0] if len(s) else 0)
        k=min(maxrank,int((s>max(tol,1e-12)).sum()));v=v[:k].copy()
        for row in v:
            if row[np.argmax(np.abs(row))]<0:row*=-1
        return v,s[:k]
    for _ in range(completion_iterations):
        components,_=basis(completed);reconstruction=(completed@components.T)@components
        completed=np.where(mask,observed_centered,reconstruction)
    components,singular=basis(completed);coefficients=completed@components.T
    return dict(components=components,coefficients=coefficients,center=center,mask=mask,masked_residuals=values,
        singular_values=singular,completion=completed,policy="own target output coordinate NaN/maskFalse except unresolvedTAZ; finite donor-gene mean centering; initialize missing centered0 then two missing-only rank<=8 unshrunk SVD reconstructions; stable max-loading sign; centered residual mean is not added back",actual_rank=len(components))

def centered_cosine(fit,query,fit_available,query_available,*,standardize=False):
    fit=np.asarray(fit,float);query=np.asarray(query,float);fa=np.asarray(fit_available,bool);qa=np.asarray(query_available,bool)
    result=np.full((len(query),len(fit)),np.nan)
    if fa.sum()<2:return result
    center=fit[fa].mean(0);spread=fit[fa].std(0) if standardize else np.ones(fit.shape[1]);spread=np.where(spread>1e-24,spread,1.)
    x=(fit-center)/spread;y=(query-center)/spread;xn=np.linalg.norm(x,axis=1);yn=np.linalg.norm(y,axis=1)
    valid=qa[:,None]&fa[None,:]&(yn[:,None]>1e-12)&(xn[None,:]>1e-12)
    return np.divide(y@x.T,yn[:,None]*xn[None,:],out=result,where=valid)

def candidate_pool(source_similarity,ada_similarity,donor_ids):
    donor_ids=tuple(donor_ids)
    def ordered(values):return sorted((i for i,v in enumerate(values) if np.isfinite(v)),key=lambda i:(-values[i],donor_ids[i]))
    source=ordered(source_similarity);ada=ordered(ada_similarity);selected=source[:4];origin=["source"]*len(selected)
    for i in ada:
        if i not in selected:
            selected.append(i);origin.append("GenePT_ADA")
            if origin.count("GenePT_ADA")==4:break
    if len(selected)>8:raise AssertionError("Candidate capacity changed")
    similarity=[]
    for i in selected:
        v=[a[i] for a in (source_similarity,ada_similarity) if np.isfinite(a[i])];similarity.append(float(np.mean(v)))
    if similarity:
        logits=np.asarray(similarity)/.2;weights=np.exp(logits-logits.max());weights/=weights.sum()
    else:weights=np.asarray([])
    return selected,origin,weights,similarity

class ResidualMemory:
    def fit(self,*,fit_target_ids,fit_deltas,genes,control_mean,baseline_views,retrieval_views,ada_features,seed):
        self.fit_ids=ids(fit_target_ids,"current-fit donor identities");self.genes=ids(genes,"response genes")
        y=np.array(fit_deltas,float,copy=True);self.control=np.array(control_mean,float,copy=True)
        if y.shape!=(len(self.fit_ids),len(self.genes)) or len(self.fit_ids)<6 or self.control.shape!=(len(self.genes),) or not np.isfinite(y).all() or not np.isfinite(self.control).all():raise ValueError("Physical current-fit response matrix required")
        if tuple(v.expert_id for v in retrieval_views)!=SOURCE_IDS or len({v.background for v in retrieval_views})!=3:raise ValueError("Retrieval uses three equal, distinct GEMZ backgrounds")
        self.baseline_views=tuple(baseline_views);self.retrieval_views=tuple(retrieval_views);self.ada=ada_features;self.y=y
        lookup={t:i for i,t in enumerate(self.fit_ids)};self.donor_oof=np.empty_like(y);seen=np.zeros(len(y),int);splits=[]
        for split in crossfit_splits(self.fit_ids,seed=seed):
            f=tuple(split["fit_target_ids"]);q=tuple(split["query_target_ids"]);fi=[lookup[t] for t in f];qi=[lookup[t] for t in q]
            pred,records=gemz_baseline(views=self.baseline_views,fit_ids=f,fit_deltas=y[fi],query_ids=q,genes=self.genes,control=self.control)
            self.donor_oof[qi]=pred;seen[qi]+=1
            splits.append(dict(split,source_fits=records,fit_response_sha256=array_hash(y[fi])))
        if not np.all(seen==1):raise ValueError("Every donor must be predicted exactly once by a model excluding its label")
        self.raw_residuals=y-self.donor_oof
        self.programs=fit_programs(self.raw_residuals,self.fit_ids,self.genes)
        self.metadata=dict(fit_target_ids=list(self.fit_ids),genes=list(self.genes),inner_splits=splits,fit_response_sha256=array_hash(y),
            donor_oof_sha256=array_hash(self.donor_oof),residual_sha256=array_hash(self.raw_residuals),program_basis_sha256=array_hash(self.programs["components"]),
            fit_scope_hash=logical_hash(dict(fit_target_ids=list(self.fit_ids),fit_response_sha256=array_hash(y),seed=seed)),actual_program_rank=self.programs["actual_rank"],
            query_truth_accepted=False,global_fit66_OOF_catalogue_used=False,program_policy=self.programs["policy"],reliability_used=False,
            inner_OOF_fit_ids_exclude_donor=True,residual_contains_own_truth=True)
        return self

    def predict(self,*,query_ids):
        queries=ids(query_ids,"query identities")
        if set(queries)&set(self.fit_ids):raise ValueError("Fit donors and query identities must be disjoint")
        baseline,base_metadata=gemz_baseline(views=self.baseline_views,fit_ids=self.fit_ids,fit_deltas=self.y,query_ids=queries,genes=self.genes,control=self.control)
        per_context=[]
        for view in self.retrieval_views:
            f,fa=view.slice(self.fit_ids);q,qa=view.slice(queries);per_context.append(centered_cosine(f,q,fa,qa))
        stack=np.stack(per_context);count=np.isfinite(stack).sum(0)
        source=np.divide(np.where(np.isfinite(stack),stack,0).sum(0),count,out=np.full(count.shape,np.nan),where=count>0)
        f,fa=self.ada.slice(self.fit_ids);q,qa=self.ada.slice(queries);ada=centered_cosine(f,q,fa,qa,standardize=True)
        means=np.clip(self.control+baseline,0,CAP);correction=np.zeros_like(means);oracle=np.broadcast_to(means[:,None,:],(len(queries),17,len(self.genes))).copy();oracle_valid=np.zeros((len(queries),17),bool);oracle_valid[:,0]=True
        pool=[]
        for j,t in enumerate(queries):
            donors,origin,weights,scores=candidate_pool(source[j],ada[j],self.fit_ids)
            if donors:
                residual_programs=self.programs["coefficients"][donors]@self.programs["components"]
                correction[j]=weights@residual_programs
                for d,r in enumerate(residual_programs):
                    for k,mix in enumerate((.25,.5)):
                        a=1+2*d+k;oracle[j,a]=np.clip(means[j]+mix*r,0,CAP);oracle_valid[j,a]=True
            pool.append(dict(query_id=t,donor_ids=[self.fit_ids[i] for i in donors],selected_from=origin,weights=weights.tolist(),similarity=scores,
                available_source_contexts=[int(count[j,i]) for i in donors],source_similarity=[None if not np.isfinite(source[j,i]) else float(source[j,i]) for i in donors],
                ADA_similarity=[None if not np.isfinite(ada[j,i]) else float(ada[j,i]) for i in donors],temperature=.2,retention=1.,fallback=not bool(donors)))
        predictions={"mix0":means,"mix25":np.clip(means+.25*correction,0,CAP),"mix50":np.clip(means+.5*correction,0,CAP)}
        return dict(predictions=predictions,oracle_candidate_means=oracle,oracle_candidate_valid=oracle_valid,correction=correction,
            metadata=dict(self.metadata,query_ids=list(queries),candidate_pools=pool,baseline_fits=base_metadata,
                source_similarity="equal mean over available pairwise contexts only; selfmissing features, center currentF44 known rows; no RAW view votes",
                ADA_similarity_policy="currentF44 available-feature mean/std then cosine; no PCA; fixed literal dictionary mask"))

def development_crossfit(*,fit_target_ids,fit_deltas,genes,control_mean,baseline_views,retrieval_views,ada_features,seed):
    targets=ids(fit_target_ids,"entire allowed development fit scope");y=np.asarray(fit_deltas,float)
    if y.shape!=(len(targets),len(genes)):raise ValueError("Allowed fit response axis differs")
    lookup={t:i for i,t in enumerate(targets)};arms={a:np.empty_like(y) for a in ("mix0","mix25","mix50")}
    oracle=np.empty((len(targets),17,len(genes)));valid=np.zeros((len(targets),17),bool);folds=[];seen=np.zeros(len(targets),int)
    for split in crossfit_splits(targets,seed=seed):
        f=tuple(split["fit_target_ids"]);q=tuple(split["query_target_ids"]);fi=[lookup[t] for t in f];qi=[lookup[t] for t in q]
        model=ResidualMemory().fit(fit_target_ids=f,fit_deltas=y[fi],genes=genes,control_mean=control_mean,baseline_views=baseline_views,retrieval_views=retrieval_views,ada_features=ada_features,seed=seed+split["fold"]+1)
        result=model.predict(query_ids=q)
        for arm in arms:arms[arm][qi]=result["predictions"][arm]
        oracle[qi]=result["oracle_candidate_means"];valid[qi]=result["oracle_candidate_valid"];seen[qi]+=1
        folds.append(dict(split,model=result["metadata"]))
    if not np.all(seen==1):raise ValueError("Development queries must be predicted exactly once")
    return dict(predictions=arms,oracle_candidate_means=oracle,oracle_candidate_valid=valid,folds=folds,
        metadata=dict(allowed_fit_target_ids=list(targets),all_predictions_generated_once=True,each_pseudoquery_outcome_excluded_from_its_model=True,
            overall_outcomes_globally_unseen=False,interpretation="development crossfit: another pseudoquery model can train on this target; all artifacts commit before diagnostic evaluation",
            genuine_agent_called=False,query34_truth_used=False,outer50_truth_used=False,official_score=False))
```
