## Supplementary reproducibility code and frozen predictions for "Centered cross-context perturbation transfer on the public 2025 Virtual Cell Challenge benchmark": stage2_fit_dataset.py

```
"""Current-fit-only cross-fitted expert arrays; no trainer or outcome reader.

Public source scalers are reusable, supervised heads and row priors are not.
The sole response argument is a physical current-fit array. A caller must pin
this module, provider code, source artifacts and its scope before real use.
"""
from __future__ import annotations

from dataclasses import dataclass
import hashlib
import json
from pathlib import Path
import re
from typing import Callable, Mapping

import numpy as np

from .transfer import Z_SCALE, RAW_SCALE, axis_hash, gene_mapping
from .dual_source_ridge import SourceScaler
from .public_source_basis import resolve_targets, _matrix_hash

VERSION = "stage2_current_fit_crossfit_v1"
STAT_COLUMNS = ("control_mean", "subfit_delta_mean", "subfit_delta_std_ddof0", "subfit_mean_abs_delta")
SOURCE_ALPHA = .001
SOURCE_SHRINKAGE = 1.


def array_hash(value):
    value = np.ascontiguousarray(value)
    h = hashlib.sha256(str(value.dtype).encode() + str(value.shape).encode())
    h.update(value.tobytes())
    return h.hexdigest()


def logical_hash(value):
    return hashlib.sha256(json.dumps(value, sort_keys=True, separators=(",", ":"), allow_nan=False).encode()).hexdigest()


def raw_pvalues_hash(value):
    """Match the separate audited raw-DE reader, retaining native p dtype."""
    value = np.ascontiguousarray(value)
    header = json.dumps({"dtype":value.dtype.str,"shape":list(value.shape)},sort_keys=True,separators=(",", ":"),ensure_ascii=False).encode()
    return hashlib.sha256(header+b"\n"+value.tobytes(order="C")).hexdigest()


def ids(value, name, *, empty=False):
    value = tuple(value)
    if ((not value and not empty) or any(not isinstance(v, str) or not v for v in value)
            or len(set(value)) != len(value) or "non-targeting" in value):
        raise ValueError("Require unique literal " + name + " without the control label")
    return value


def copy_metadata(value):
    return json.loads(json.dumps(value, allow_nan=False))


def readonly(value, dtype=None):
    value = np.array(value, dtype=dtype, copy=True)
    value.setflags(write=False)
    return value


@dataclass(frozen=True)
class FrozenSourceView:
    expert_id: str
    background: tuple[str, str]
    view_kind: str
    targets: tuple[str, ...]
    features: np.ndarray
    available: np.ndarray
    metadata: dict
    evidence_genes: tuple[str, ...] = ()
    evidence: np.ndarray | None = None
    evidence_available: np.ndarray | None = None

    def __post_init__(self):
        targets = ids(self.targets, "source feature targets")
        x = np.asarray(self.features, dtype=np.float64); mask = np.asarray(self.available)
        if (not self.expert_id or self.view_kind not in ("GEMZ", "RAW") or len(self.background) != 2
                or x.ndim != 2 or x.shape[0] != len(targets) or x.shape[1] < 1
                or mask.dtype.kind != "b" or mask.shape != (len(targets),) or not np.isfinite(x[mask]).all()):
            raise ValueError("Finite frozen source features and explicit availability required")
        if self.metadata.get("scope") != "frozen_public" or self.metadata.get("H1_response_used") is not False:
            raise ValueError("Source features must have an explicit external-only public scope")
        object.__setattr__(self, "targets", targets); object.__setattr__(self, "features", readonly(x))
        object.__setattr__(self, "available", readonly(mask)); object.__setattr__(self, "background", tuple(map(str, self.background)))
        object.__setattr__(self, "metadata", copy_metadata(self.metadata))
        if self.evidence is None:
            if self.evidence_available is not None or self.evidence_genes:
                raise ValueError("Evidence axes/mask require evidence values")
        else:
            genes = ids(self.evidence_genes, "source evidence genes"); ev = np.asarray(self.evidence, float)
            em = np.asarray(self.evidence_available)
            if em.dtype.kind != "b" or ev.shape != (len(targets), len(genes)) or em.shape != ev.shape or not np.isfinite(ev[em]).all():
                raise ValueError("Source evidence must have an explicit finite-measurement mask")
            object.__setattr__(self, "evidence_genes", genes); object.__setattr__(self, "evidence", readonly(np.where(em, ev, np.nan)))
            object.__setattr__(self, "evidence_available", readonly(em))

    @property
    def declaration(self):
        return dict(expert_id=self.expert_id, background=list(self.background), view_kind=self.view_kind,
                    feature_targets=list(self.targets), features_sha256=array_hash(self.features),
                    available_sha256=array_hash(self.available), evidence_genes=list(self.evidence_genes),
                    evidence_sha256=None if self.evidence is None else array_hash(self.evidence),
                    evidence_available_sha256=None if self.evidence_available is None else array_hash(self.evidence_available),
                    metadata=self.metadata, source_alpha=SOURCE_ALPHA, source_shrinkage=SOURCE_SHRINKAGE)

    def slice(self, target_ids):
        lookup = {t:i for i,t in enumerate(self.targets)}
        if any(t not in lookup for t in target_ids):
            raise ValueError("Target absent from source feature whitelist")
        rows = [lookup[t] for t in target_ids]
        # Unknown placeholders cannot influence a fit center or kernel.
        return np.where(self.available[rows,None], self.features[rows], 0), self.available[rows].copy()

    def slice_evidence(self, target_ids, genes):
        if self.evidence is None:
            return np.full((len(target_ids),len(genes)), np.nan), np.zeros((len(target_ids),len(genes)),bool)
        if tuple(genes) != self.evidence_genes:
            raise ValueError("Evidence gene coordinate order differs")
        lookup = {t:i for i,t in enumerate(self.targets)}; rows = [lookup[t] for t in target_ids]
        return self.evidence[rows].copy(), self.evidence_available[rows].copy()


def source_view_from_public(source, basis, *, target_ids, gene_names, output_ensembl_ids=None):
    """Encode frozen external inputs once, outside response cross-fitting.

    Full numeric public-source identity is verified. No H1 response is accepted.
    The optional evidence is the original measured source control-relative delta,
    not its standardized features or an H1-calibrated response.
    """
    targets = ids(target_ids,"public encoding targets"); genes = ids(gene_names,"H1 expression genes")
    if basis.metadata["config"]["rank"] != 128 or source.scale not in (Z_SCALE,RAW_SCALE):
        raise ValueError("Use the registered rank128 public centers/spreads and legal source scale")
    scaler = SourceScaler.from_public_basis(basis)
    if scaler.source_id != source.source_id or scaler.metadata["source_file_sha256"] != source.metadata.get("file_sha256"):
        raise ValueError("Public scaler/source identity mismatch")
    full_genes = tuple(sorted(source.genes)); full_targets=tuple(sorted(source.target_ids))
    sx,sm=source.aligned(full_targets,full_genes);eligible=sm.any(axis=1)
    eligible_targets=tuple(t for t,valid in zip(full_targets,eligible) if valid)
    sx,sm=sx[eligible],sm[eligible]
    if tuple(scaler.metadata["public_basis_target_ids"])!=eligible_targets or basis.metadata["config"]["min_gene_observations"]!=2:
        raise ValueError("All-public scaler must cover the full eligible public target axis")
    if _matrix_hash(eligible_targets,full_genes,sx,sm) != scaler.metadata["public_basis_input_sha256"]:
        raise ValueError("Public scaler numeric source input changed")
    count=sm.sum(axis=0);center=np.divide(np.where(sm,sx,0).sum(axis=0),count,out=np.zeros(len(full_genes)),where=count>0)
    centered=np.where(sm,sx-center,0);variance=np.divide((centered**2).sum(axis=0),count,out=np.zeros(len(full_genes)),where=count>0)
    keep=(count>=2)&(variance>1e-24)
    if (scaler.genes!=tuple(g for g,valid in zip(full_genes,keep) if valid) or
            not np.array_equal(scaler.center,center[keep]) or not np.array_equal(scaler.spread,np.sqrt(variance[keep]))):
        raise ValueError("Frozen public scaler genes/centers/spreads differ from exact full-source statistics")
    mapping = resolve_targets(source,targets)
    known = [mapping[t] for t in targets if mapping[t] is not None]
    if len(known) != len(set(known)):
        raise ValueError("Targets duplicate a canonical source identity")
    encoded = tuple(mapping[t] or "__missing__"+t for t in targets)
    features, available = scaler.encode(source, encoded)
    original, observed = source.aligned(encoded,source.genes)
    symbols = source.metadata["source_expression_symbols"]
    coordinates, alignment = gene_mapping(source.genes,symbols,genes,output_ensembl_ids)
    # No unverified interpretation of the ambiguous H1 TAZ coordinate.
    if "TAZ" in genes and not (output_ensembl_ids or {}).get("TAZ"):
        coordinates[genes.index("TAZ")] = -1
    evidence = np.full((len(targets),len(genes)),np.nan); mask = np.zeros(evidence.shape,bool)
    matched = coordinates >= 0
    evidence[:,matched] = original[:,coordinates[matched]]; mask[:,matched] = observed[:,coordinates[matched]]
    context = source.metadata["source_context"]
    return FrozenSourceView(source.source_id,(str(context["cell_line"]),str(context["time_days"])),
        "GEMZ" if source.scale==Z_SCALE else "RAW",targets,features,available,
        dict(scope="frozen_public",H1_response_used=False,public_scaler=scaler.metadata,
             source_scale=source.scale,source_gene_axis_sha256=axis_hash(source.genes),
             evidence_gene_alignment=alignment,evidence_units="signed GEM-Z" if source.scale==Z_SCALE else "log1p population delta",
             TAZ_target_unknown=True,public_components_ignored=True),genes,evidence,mask)


@dataclass(frozen=True)
class ExpertPrediction:
    deltas: np.ndarray
    available: np.ndarray
    metadata: dict
    evidence: np.ndarray | None = None
    evidence_available: np.ndarray | None = None


class GenePTFoldAdapter:
    """Fresh optional numerical expert; it never receives query responses."""
    def __init__(self, *, features, alpha, shrinkage=1., pca_rank=None, standardize=True):
        self.features,self.params = features,dict(alpha=alpha,shrinkage=shrinkage,pca_rank=pca_rank,standardize=standardize)

    @property
    def declaration(self):
        from . import genept_target_kernel as kernel
        return dict(kind="frozen_target_feature_kernel",parameters=self.params,representation_id=self.features.representation_id,
                    frozen_feature_metadata=self.features.metadata,feature_targets=list(self.features.targets),
                    feature_values_sha256=array_hash(self.features.values),feature_available_sha256=array_hash(self.features.available),
                    numerical_code_sha256=hashlib.sha256(Path(kernel.__file__).read_bytes()).hexdigest())

    def fit(self, *, fit_target_ids, fit_deltas, genes, control_mean, fit_scope_hash):
        from .genept_target_kernel import GenePTTargetKernelExpert
        self.scope_hash=fit_scope_hash; self.control=control_mean.copy()
        self.model=GenePTTargetKernelExpert(**self.params).fit(fit_target_ids=fit_target_ids,fit_deltas=fit_deltas,genes=genes,features=self.features)
        return self

    def predict(self, *, query_target_ids):
        result=self.model.predict(query_target_ids=query_target_ids)
        return ExpertPrediction(result.deltas,result.available,dict(result.metadata,fit_scope_hash=self.scope_hash,
            fit_target_ids=list(self.model.fit_ids),query_target_ids=list(query_target_ids),genes=list(self.model.genes),query_truth_accepted=False))


def target_kernel_registration(features):
    """New fixed settings for GenePT or the existing frozen Sol30 arrays.

    The input availability mask is respected verbatim; this function does not
    change TAZ annotations or infer masks from literal feature values.
    """
    factory=lambda:GenePTFoldAdapter(features=features,alpha=.001,shrinkage=1.,pca_rank=None,standardize=True)
    return factory,factory().declaration


def crossfit_splits(fit_targets, *, seed):
    ordered=tuple(sorted(fit_targets)); permutation=np.random.default_rng(seed).permutation(len(ordered))
    result=[]
    for fold,part in enumerate(np.array_split(permutation,3)):
        q=tuple(ordered[i] for i in sorted(part)); qset=set(q); f=tuple(t for t in ordered if t not in qset)
        result.append(dict(fold=fold,fit_target_ids=list(f),query_target_ids=list(q),split_id=logical_hash(dict(seed=seed,fold=fold,fit_targets=f,query_targets=q))))
    return result


def gene_statistics(deltas,control):
    return np.stack((control,deltas.mean(axis=0),deltas.std(axis=0),np.abs(deltas).mean(axis=0)),axis=1)


def bh_labels(raw_pvalues, *, threshold=.05):
    from scipy.stats import false_discovery_control
    p=np.asarray(raw_pvalues)
    if p.ndim!=2 or p.dtype.kind!="f" or p.dtype.itemsize not in (4,8) or not np.isfinite(p).all() or np.any((p<0)|(p>1)):
        raise ValueError("Raw finite float32/float64 p-values after official NaN postprocessing required")
    adjusted=false_discovery_control(p.reshape(-1),method="bh").reshape(p.shape)
    return adjusted<threshold, adjusted


def source_prediction(view,*,fit_ids,fit_deltas,query_ids,genes,control,fit_scope_hash):
    x,known=view.slice(fit_ids); xq,query_known=view.slice(query_ids); common=fit_deltas.mean(axis=0)
    prediction=np.broadcast_to(common,(len(query_ids),len(genes))).copy(); available=np.zeros(len(query_ids),bool)
    meta=dict(fit_scope_hash=fit_scope_hash,fit_target_ids=list(fit_ids),query_target_ids=list(query_ids),genes=list(genes),
              fit_response_sha256=array_hash(fit_deltas),query_truth_accepted=False,source=view.expert_id,
              n_available_fit=int(known.sum()),alpha=SOURCE_ALPHA,shrinkage=SOURCE_SHRINKAGE,
              scaler_scope="frozen_public spreads plus current-subfit feature center",PCA_used=False,enabled=False)
    if known.sum()>=3:
        design=x[known]; center=design.mean(axis=0); design=design-center; gram=design@design.T
        trace=float(np.trace(gram)/len(design))
        if np.isfinite(trace) and trace>1e-24:
            y=fit_deltas[known]; ym=y.mean(axis=0)
            coefficients=np.linalg.solve(gram/trace+SOURCE_ALPHA*len(design)*np.eye(len(design)),y-ym)
            prediction[query_known]=ym+((xq[query_known]-center)@design.T/trace)@coefficients
            available=query_known.copy()
            meta.update(enabled=True,feature_center_sha256=array_hash(center),kernel_trace_scale=trace,
                normalized_regularizer=SOURCE_ALPHA*len(design),raw_materialized_regularizer=SOURCE_ALPHA*len(design)*trace,
                normalized_kernel_mean_diagonal=float(np.trace(gram/trace)/len(design)))
    evidence,em=view.slice_evidence(query_ids,genes)
    return ExpertPrediction(prediction,available,meta,evidence,em)


def validate_expert(result,*,fit_ids,query_ids,genes,fit_deltas,control,fit_scope_hash):
    if not isinstance(result,ExpertPrediction):
        raise ValueError("Provider must return an explicit ExpertPrediction")
    meta=result.metadata
    checks={"fit_scope_hash":fit_scope_hash,"fit_target_ids":list(fit_ids),"query_target_ids":list(query_ids),
            "genes":list(genes),"fit_response_sha256":array_hash(fit_deltas),"query_truth_accepted":False}
    if any(meta.get(k)!=v for k,v in checks.items()):
        raise ValueError("Provider fitting scope/response/axes differ; global OOF replay is forbidden")
    d=np.asarray(result.deltas,float); mask=np.asarray(result.available)
    shape=(len(query_ids),len(genes))
    if mask.dtype.kind!="b" or d.shape!=shape or mask.shape not in ((len(query_ids),),shape):
        raise ValueError("Expert prediction/mask shapes differ")
    mask=np.broadcast_to(mask[:,None],shape).copy() if mask.ndim==1 else mask.copy()
    if not np.isfinite(d[mask]).all():
        raise ValueError("Available expert predictions must be finite")
    fallback=fit_deltas.mean(axis=0)
    d=np.where(mask,d,fallback)
    # Match Stage1 mean projection, preserving delta units after projection.
    d=np.maximum(control+d,0)-control
    if result.evidence is None:
        if result.evidence_available is not None: raise ValueError("Evidence mask without values")
        ev=np.full(shape,np.nan); em=np.zeros(shape,bool)
    else:
        ev=np.asarray(result.evidence,float); em=np.asarray(result.evidence_available)
        if ev.shape!=shape or em.shape!=shape or em.dtype.kind!="b" or not np.isfinite(ev[em]).all():
            raise ValueError("Evidence coordinates/availability invalid")
        ev=np.where(em,ev,np.nan)
    return d,mask,ev,em,dict(meta,fallback="current subfit common delta, unavailable mask remains false",mean_projection="max(control+delta,0)-control")


@dataclass(frozen=True)
class ScopeDataset:
    arrays: dict
    metadata: dict

    @property
    def scope_hash(self): return self.metadata["scope_hash"]

    @property
    def ready_to_train(self): return self.metadata["ready_to_train"]

    @property
    def fit_targets(self): return tuple(self.metadata["fit_targets"])

    @property
    def query_targets(self): return tuple(self.metadata["query_targets"])

    @property
    def gene_names(self): return tuple(self.metadata["gene_names"])

    @property
    def expert_ids(self): return tuple(self.metadata["expert_ids"])

    def validate_scope(self,*,fit_targets,query_targets,gene_names,scope_hash):
        if logical_hash({k:v for k,v in self.metadata.items() if k!="metadata_sha256"})!=self.metadata["metadata_sha256"]:
            raise ValueError("Dataset metadata changed")
        expected=dict(fit_targets=list(fit_targets),query_targets=list(query_targets),gene_names=list(gene_names),scope_hash=scope_hash)
        if any(self.metadata.get(k)!=v for k,v in expected.items()):
            raise ValueError("Dataset belongs to a different current fitting scope")
        declared=self.metadata["scope_declaration"]
        if logical_hash(declared)!=self.scope_hash or any(self.metadata[k]!=v for k,v in declared.items()):
            raise ValueError("Dataset scope declaration changed")
        if {k:array_hash(v) for k,v in self.arrays.items()}!=self.metadata["array_hashes"]:
            raise ValueError("Dataset numerical contents changed")
        return self

    def save(self,path):
        path=Path(path)
        if path.exists(): raise FileExistsError("Scope artifacts are immutable")
        self.validate_scope(fit_targets=self.metadata["fit_targets"],query_targets=self.metadata["query_targets"],gene_names=self.metadata["gene_names"],scope_hash=self.scope_hash)
        path.parent.mkdir(parents=True,exist_ok=True)
        with path.open("xb") as stream:
            np.savez_compressed(stream,metadata=np.asarray(json.dumps(self.metadata,allow_nan=False)),**self.arrays)
        return hashlib.sha256(path.read_bytes()).hexdigest()

    @classmethod
    def load(cls,path,*,fit_targets,query_targets,gene_names,scope_hash):
        with np.load(path,allow_pickle=False) as saved:
            meta=json.loads(str(saved["metadata"]))
            if meta.get("version")!=VERSION or set(saved.files)!={"metadata",*meta["array_hashes"]}:
                raise ValueError("Unexpected scope artifact schema")
            data=cls({k:readonly(saved[k]) for k in meta["array_hashes"]},meta)
        return data.validate_scope(fit_targets=fit_targets,query_targets=query_targets,gene_names=gene_names,scope_hash=scope_hash)


def build_current_fit_dataset(*,fit_targets,query_targets,gene_names,fit_deltas,control_mean,source_views,
        parent_scope_id,frozen_input_hashes,seed=20251013,optional_expert_factories=None,expert_declarations=None,
        required_experts=(),fit_raw_pvalues=None,raw_pvalue_provenance=None,fit_de_labels=None):
    """Build OOF training experts and refitted query experts; accepts no query Y.

    Optional fresh factories return adapters with fit(physical fit arrays,
    fit_scope_hash) and predict(query_target_ids only). Their declared provenance
    is checked separately from prediction scope. This is an API boundary, not
    operating-system isolation against malicious provider implementations.
    """
    fit=ids(fit_targets,"current fit targets"); query=ids(query_targets,"query targets",empty=True); genes=ids(gene_names,"expression genes")
    if len(fit)<6 or set(fit)&set(query) or not parent_scope_id:
        raise ValueError("Three-fold cross-fitting needs at least six disjoint current-fit targets")
    y=np.array(fit_deltas,dtype=np.float64,copy=True); control=np.asarray(control_mean,dtype=np.float64)
    if y.shape!=(len(fit),len(genes)) or control.shape!=(len(genes),) or not np.isfinite(y).all() or not np.isfinite(control).all() or np.any(control<0) or np.any(control+y<0):
        raise ValueError("Physical finite fit response rows and nonnegative shared log1p means required")
    if not frozen_input_hashes or any(not isinstance(v,str) or not re.fullmatch(r"[0-9a-f]{64}",v) for v in frozen_input_hashes.values()):
        raise ValueError("Explicit frozen input hashes required")
    views=tuple(source_views); primary=[v for v in views if v.view_kind=="GEMZ"]; raw=[v for v in views if v.view_kind=="RAW"]
    if len(primary)!=3 or len({v.background for v in primary})!=3 or any(v.background not in {p.background for p in primary} for v in raw):
        raise ValueError("Require three independent GEMZ backgrounds; RAW views must be paired controls")
    if len({(v.background,v.view_kind) for v in views})!=len(views) or len({v.expert_id for v in views})!=len(views):
        raise ValueError("Duplicate source context/view")
    factories=dict(optional_expert_factories or {}); declarations=copy_metadata(expert_declarations or {})
    if set(factories)!=set(declarations): raise ValueError("Each optional expert needs a frozen numerical declaration")
    expert_ids=("control","common",*(v.expert_id for v in views),*factories)
    if len(set(expert_ids))!=len(expert_ids): raise ValueError("Duplicate expert IDs")
    required_experts=tuple(required_experts)
    if len(required_experts)!=len(set(required_experts)) or any(not isinstance(v,str) or not v for v in required_experts):
        raise ValueError("Require a unique literal required-expert whitelist")
    unavailable_required=sorted(set(required_experts)-set(expert_ids))
    rawp=None; labels=None; full_frequency=None
    if fit_raw_pvalues is not None:
        rawp=np.array(fit_raw_pvalues,copy=True)
        if rawp.shape!=y.shape: raise ValueError("Raw p-value matrix must be physical current-fit rows")
        provenance=raw_pvalue_provenance or {}
        required=dict(scope="raw_per_target",fit_targets=list(fit),gene_names=list(genes),adjusted_pvalues=False,NaN_p_postprocessed=True,full_gene_axis=True,normalization_target_sum=54611.)
        if any(provenance.get(k)!=v for k,v in required.items()) or provenance.get("raw_pvalues_sha256")!=raw_pvalues_hash(rawp):
            raise ValueError("Raw p-value provenance must match current fit scope, full genes and L54611")
        labels,adjusted=bh_labels(rawp)
        full_frequency=(labels.sum(axis=0)+1)/(len(fit)+2)
    elif raw_pvalue_provenance is not None:
        raise ValueError("Raw provenance without raw measurements")
    if fit_de_labels is not None:
        provided=np.asarray(fit_de_labels)
        if provided.shape!=y.shape or not np.isfinite(provided).all() or np.any((provided!=0)&(provided!=1)):
            raise ValueError("DE labels must be current-fit aligned binary labels")
        provided=provided.astype(bool)
        if labels is not None and not np.array_equal(provided,labels):
            raise ValueError("DE labels are not this current panel's recomputed BH labels")
        if labels is None: labels=provided
    declaration=dict(version=VERSION,parent_scope_id=parent_scope_id,fit_targets=list(fit),query_targets=list(query),gene_names=list(genes),
        fit_deltas_sha256=array_hash(y),control_mean_sha256=array_hash(control),raw_pvalues_sha256=None if rawp is None else raw_pvalues_hash(rawp),
        fit_de_labels_sha256=None if labels is None else array_hash(labels),raw_pvalue_provenance_sha256=logical_hash(raw_pvalue_provenance),
        external_source_declarations=[v.declaration for v in views],optional_expert_declarations=declarations,
        expert_ids=list(expert_ids),required_experts=list(required_experts),frozen_input_hashes=dict(frozen_input_hashes),
        seed=int(seed),folds=3,source_alpha=SOURCE_ALPHA,source_shrinkage=SOURCE_SHRINKAGE)
    scope_hash=logical_hash(declaration); n,q,g,e=len(fit),len(query),len(genes),len(expert_ids)
    # Provider-facing hashes exclude full-scope H1 response/DE hashes. A
    # provider may use this hash as an RNG seed, so even opaque audit hashes
    # must not convey a heldout OOF row's own outcome into its prediction.
    provider_context={key:declaration[key] for key in ("version","parent_scope_id","fit_targets","query_targets","gene_names",
        "control_mean_sha256","external_source_declarations","optional_expert_declarations","expert_ids","seed","folds","source_alpha","source_shrinkage")}
    arrays=dict(fit_expert_deltas=np.empty((n,e,g)),query_expert_deltas=np.empty((q,e,g)),
        fit_expert_available=np.zeros((n,e,g),bool),query_expert_available=np.zeros((q,e,g),bool),
        fit_source_evidence=np.full((n,e,g),np.nan),query_source_evidence=np.full((q,e,g),np.nan),
        fit_evidence_available=np.zeros((n,e,g),bool),query_evidence_available=np.zeros((q,e,g),bool),
        fit_row_common_deltas=np.empty((n,g)),fit_row_gene_statistics=np.empty((n,g,4)),
        query_gene_statistics=gene_statistics(y,control),common_delta=y.mean(axis=0),control_mean=control.copy(),
        fit_truth_means=control+y,fit_reference_deltas=y.copy(),matched_fit_indices=np.arange(n,dtype=np.int64),
        target_gene_indices=np.asarray([genes.index(t) if t in genes else -1 for t in fit],dtype=np.int64),fit_row_subfold=np.full(n,-1,dtype=np.int64))
    if rawp is not None:
        arrays.update(fit_de_labels=labels,fit_full_panel_fdr=adjusted,fit_row_de_frequency=np.empty((n,g)),query_de_frequency=full_frequency)
    elif labels is not None: arrays["fit_de_labels"]=labels
    lookup={t:i for i,t in enumerate(fit)}; splits=crossfit_splits(fit,seed=seed); receipts=[]; instances=[]
    def predict_panel(subfit,subquery,suby,subscope):
        rows=[]; masks=[]; evidences=[]; evidence_masks=[]; reports={}
        for anchor in ("control","common"):
            delta=np.zeros(g) if anchor=="control" else suby.mean(axis=0)
            rows.append(np.broadcast_to(np.maximum(control+delta,0)-control,(len(subquery),g)).copy());masks.append(np.ones((len(subquery),g),bool))
            evidences.append(np.full((len(subquery),g),np.nan));evidence_masks.append(np.zeros((len(subquery),g),bool))
        for view in views:
            result=source_prediction(view,fit_ids=subfit,fit_deltas=suby,query_ids=subquery,genes=genes,control=control,fit_scope_hash=subscope)
            d,m,ev,em,report=validate_expert(result,fit_ids=subfit,query_ids=subquery,genes=genes,fit_deltas=suby,control=control,fit_scope_hash=subscope)
            rows.append(d);masks.append(m);evidences.append(ev);evidence_masks.append(em);reports[view.expert_id]=report
        for name,factory in factories.items():
            adapter=factory()
            if any(adapter is old for old in instances): raise ValueError("Optional providers must create a fresh model per subfit")
            instances.append(adapter)
            if copy_metadata(adapter.declaration)!=declarations[name]: raise ValueError("Optional expert declaration changed")
            adapter.fit(fit_target_ids=subfit,fit_deltas=suby.copy(),genes=genes,control_mean=control.copy(),fit_scope_hash=subscope)
            result=adapter.predict(query_target_ids=subquery)
            if copy_metadata(adapter.declaration)!=declarations[name]: raise ValueError("Optional expert declaration changed after fit")
            d,m,ev,em,report=validate_expert(result,fit_ids=subfit,query_ids=subquery,genes=genes,fit_deltas=suby,control=control,fit_scope_hash=subscope)
            rows.append(d);masks.append(m);evidences.append(ev);evidence_masks.append(em);reports[name]=report
        return np.stack(rows,axis=1),np.stack(masks,axis=1),np.stack(evidences,axis=1),np.stack(evidence_masks,axis=1),reports
    for split in splits:
        f=tuple(split["fit_target_ids"]); held=tuple(split["query_target_ids"])
        fi=[lookup[t] for t in f]; qi=[lookup[t] for t in held]; suby=y[fi].copy()
        subscope=logical_hash(dict(provider_context=provider_context,split_id=split["split_id"],fit_targets=f,query_targets=held,fit_response_sha256=array_hash(suby)))
        d,m,ev,em,reports=predict_panel(f,held,suby,subscope)
        for key,value in (("fit_expert_deltas",d),("fit_expert_available",m),("fit_source_evidence",ev),("fit_evidence_available",em)):
            arrays[key][qi]=value
        arrays["fit_row_common_deltas"][qi]=suby.mean(axis=0);arrays["fit_row_gene_statistics"][qi]=gene_statistics(suby,control)
        arrays["fit_row_subfold"][qi]=split["fold"]
        receipt=dict(**split,fit_scope_hash=subscope,fit_response_sha256=array_hash(suby),expert_reports=reports)
        if rawp is not None:
            sublabels,_=bh_labels(rawp[fi]);arrays["fit_row_de_frequency"][qi]=(sublabels.sum(axis=0)+1)/(len(fi)+2)
            receipt.update(BH_family=len(fi)*g,raw_pvalues_sha256=raw_pvalues_hash(rawp[fi]),DE_targets=list(f))
        receipts.append(receipt)
    if np.any(arrays["fit_row_subfold"]<0): raise AssertionError("Crossfit missed a current-fit target")
    query_scope=logical_hash(dict(provider_context=provider_context,fit_targets=fit,query_targets=query,fit_response_sha256=array_hash(y)))
    if q:
        d,m,ev,em,query_reports=predict_panel(fit,query,y.copy(),query_scope)
        arrays.update(query_expert_deltas=d,query_expert_available=m,query_source_evidence=ev,query_evidence_available=em)
    else: query_reports={}
    metadata=dict(declaration,scope_hash=scope_hash,scope_declaration=copy_metadata(declaration),expert_ids=list(expert_ids),source_backgrounds=3,RAW_GEMZ_views_independent=False,
        expert_backgrounds={v.expert_id:list(v.background) for v in views},crossfit_splits=receipts,query_fit_scope_hash=query_scope,query_expert_reports=query_reports,
        ready_to_train=rawp is not None and not unavailable_required,missing_required_experts=unavailable_required,
        DE_ready=rawp is not None,DE_unready_reason=None if rawp is not None else "Raw current-fit p-values absent: subfit BH and row prior unavailable",
        gene_statistics_columns=list(STAT_COLUMNS),gene_statistics_standardized=False,statistics_ddof=0,frequency_smoothing="Beta(1,1): (subfit DE count + 1)/(subfit target count + 2)",
        crossfit_provenance=dict(fit_targets=list(fit),query_truth_used=False,all_fit_rows_predicted_once=True,fit_row_priors_exclude_own_outcome=True,
                                folds=3,seed=int(seed),current_scope_hash=scope_hash,split_ids=[s["split_id"] for s in splits]),
        BH_policy="Separate entire-current-fit and each entire-subfit full target-by-gene BH, threshold strictly < .05",raw_pvalue_provenance=copy_metadata(raw_pvalue_provenance),
        response_dtype="float64",availability_dtype="bool",raw_pvalue_dtype=None if rawp is None else str(rawp.dtype),
        expert_missing_representation="finite current-subfit common fallback with explicit false availability; never inferred observed zero",
        raw_pvalue_hash_convention="SHA256(canonical UTF8 {dtype:dtype.str,shape:list} + LF + C-order bytes)",
        internal_array_hash_convention="SHA256(str(dtype) + str(shape) + C-order bytes)",
        evidence_missing_representation="NaN plus explicit false mask",fit_row_priors="subfit outcomes only; exclude every heldout crossfit row",query_priors="all and only current-fit outcomes",
        pds_references="fit_reference_deltas only, positive matching indices and literal target gene coordinates; no query references",
        query_truth_accepted=False,validation_test_expression_opened=False,training_launched=False,agent_called=False,official_score=False,
        scope_isolation="explicit array/API/hash protocol, not an operating-system sandbox",allocated_array_bytes=sum(v.nbytes for v in arrays.values()),
        array_hashes={k:array_hash(v) for k,v in arrays.items()})
    metadata["metadata_sha256"]=logical_hash(metadata)
    return ScopeDataset({k:readonly(v) for k,v in arrays.items()},metadata)
```
