## Supplementary reproducibility code and frozen predictions for "Centered cross-context perturbation transfer on the public 2025 Virtual Cell Challenge benchmark": transfer.py

```
"""E1: fit-target-only cross-context pseudobulk transfer.

Replogle bulk profiles are floating point population profiles, not integer
single-cell counts. Raw log1p-pseudobulk and GEM-group Z profiles remain
separate domains. Neither is an evaluator submission space. This module never
opens H1 validation/test expression and does not compute an official score.
"""
from __future__ import annotations

from dataclasses import dataclass, field
import hashlib
import json
from pathlib import Path
import re
from typing import Iterable, Mapping, Sequence

import numpy as np


RAW_SCALE = "raw-expression pseudobulk profile"
Z_SCALE = "gemgroup control-relative z-score pseudobulk"
POPULATION_FILTER_POLICY = "exclude non-finite or non-positive num_cells_filtered from core controls and requested constructs; no weight imputation; empty targets remain missing"
_LABEL = re.compile(r"\d+_(.+)_((?:P\d+(?:P\d+)*)|(?:ENST\d+(?:\.\d+)?(?:,ENST\d+(?:\.\d+)?)*))_(ENSG\d+|nan)")


def axis_hash(values: Sequence[str]) -> str:
    return hashlib.sha256("\n".join(map(str, values)).encode("utf-8")).hexdigest()


def file_hash(path: Path, algorithm: str = "sha256") -> str:
    result = hashlib.new(algorithm)
    with Path(path).open("rb") as handle:
        while block := handle.read(8 * 1024 * 1024):
            result.update(block)
    return result.hexdigest()


def _ids(values: Iterable[str], name: str) -> tuple[str, ...]:
    result = tuple(map(str, values))
    if len(set(result)) != len(result) or any(not value for value in result):
        raise ValueError(f"{name} must be unique, nonempty identities")
    return result


def _ensembl(value: str | None) -> str | None:
    if value is None or not str(value).startswith("ENSG"):
        return None
    return str(value).split(".")[0]


def parse_construct(label: str, row: int) -> dict | None:
    match = _LABEL.fullmatch(label)
    if not match:
        return None
    return dict(row=int(row), symbol=match[1], target_unit=match[2],
                target_unit_type="promoter" if match[2].startswith("P") else "transcript",
                promoter=match[2] if match[2].startswith("P") else None,
                ensembl_id=match[3] if match[3] != "nan" else None)


def _column(frame, key: str) -> np.ndarray:
    """Read current and pre-0.7 AnnData categorical/string columns with h5py."""
    import h5py

    node = frame[key]
    if isinstance(node, h5py.Group):
        if not {"codes", "categories"} <= set(node):
            raise ValueError(f"Unsupported AnnData column encoding for {key}")
        codes, categories = node["codes"][...], _dataset(node["categories"])
    elif "__categories" in frame and key in frame["__categories"]:
        codes, categories = node[...], _dataset(frame["__categories"][key])
    else:
        return _dataset(node)
    result = np.empty(len(codes), dtype=object)
    if np.any(codes >= len(categories)) or np.any(codes < -1):
        raise ValueError(f"Invalid categorical codes for {key}")
    result[:] = None
    valid = codes >= 0
    result[valid] = categories[codes[valid].astype(int)]
    return result


def _dataset(node) -> np.ndarray:
    import h5py

    return node.asstr()[...] if h5py.check_string_dtype(node.dtype) else node[...]


def _index(frame) -> tuple[str, ...]:
    name = frame.attrs.get("_index", "_index")
    if isinstance(name, bytes):
        name = name.decode("utf-8")
    return _ids(_column(frame, str(name)), "AnnData axis")


def _booleans(values: np.ndarray) -> np.ndarray:
    result = []
    for value in values:
        if isinstance(value, (bool, np.bool_)) or value in (0, 1):
            result.append(bool(value))
        elif str(value).lower() in ("true", "false"):
            result.append(str(value).lower() == "true")
        else:
            raise ValueError("core_control has a missing or invalid boolean")
    return np.asarray(result, dtype=bool)


def gene_mapping(source_ids: Sequence[str], source_symbols: Sequence[str],
                 output_genes: Sequence[str],
                 output_ensembl_ids: Mapping[str, str] | None = None) -> tuple[np.ndarray, dict]:
    """Prefer declared Ensembl identity, otherwise require a unique exact symbol.

    Duplicate symbols never get silently averaged or assigned to the first gene.
    Explicit Ensembl metadata overrides a conflicting symbol match.
    """
    source_ids = _ids(source_ids, "source gene axis")
    output_genes = _ids(output_genes, "output gene axis")
    if len(source_symbols) != len(source_ids):
        raise ValueError("Source symbol axis length differs from Ensembl axis")
    by_ens: dict[str, list[int]] = {}
    by_symbol: dict[str, list[int]] = {}
    for i, (ens, symbol) in enumerate(zip(source_ids, source_symbols)):
        by_ens.setdefault(_ensembl(ens) or ens, []).append(i)
        if symbol is not None and str(symbol):
            by_symbol.setdefault(str(symbol), []).append(i)
    mapping = np.full(len(output_genes), -1, dtype=int)
    ambiguous, missing = [], []
    for i, gene in enumerate(output_genes):
        ens = _ensembl((output_ensembl_ids or {}).get(gene)) or _ensembl(gene)
        matches = by_ens.get(ens, []) if ens else by_symbol.get(gene, [])
        if len(matches) == 1:
            mapping[i] = matches[0]
        elif matches:
            ambiguous.append(gene)
        else:
            missing.append(gene)
    return mapping, dict(matched=int(np.sum(mapping >= 0)), missing=missing,
                         ambiguous=ambiguous, alias_substitution=False)


@dataclass
class ExternalSource:
    """One context/time/scale block. NaN plus mask denotes absent measurement."""
    source_id: str
    target_ids: tuple[str, ...]
    genes: tuple[str, ...]
    deltas: np.ndarray
    mask: np.ndarray
    scale: str
    metadata: dict = field(default_factory=dict)

    def __post_init__(self):
        self.target_ids = _ids(self.target_ids, "source targets")
        self.genes = _ids(self.genes, "source genes")
        self.deltas = np.asarray(self.deltas, dtype=np.float64)
        self.mask = np.asarray(self.mask, dtype=bool)
        shape = (len(self.target_ids), len(self.genes))
        if self.deltas.shape != shape or self.mask.shape != shape:
            raise ValueError("Source delta and availability mask have incorrect shape")
        if not np.isfinite(self.deltas[self.mask]).all():
            raise ValueError("Observed source deltas must be finite")
        if self.scale not in (RAW_SCALE, Z_SCALE):
            raise ValueError("Declare raw-pseudobulk or GEM-Z scale explicitly")

    def aligned(self, target_ids: Sequence[str], genes: Sequence[str]) -> tuple[np.ndarray, np.ndarray]:
        targets, genes = _ids(target_ids, "requested targets"), _ids(genes, "requested genes")
        output = np.full((len(targets), len(genes)), np.nan)
        mask = np.zeros(output.shape, dtype=bool)
        row_map = {target: i for i, target in enumerate(self.target_ids)}
        gene_map = {gene: i for i, gene in enumerate(self.genes)}
        rows = [(i, row_map[target]) for i, target in enumerate(targets) if target in row_map]
        columns = [(i, gene_map[gene]) for i, gene in enumerate(genes) if gene in gene_map]
        if rows and columns:
            out_row, src_row = np.asarray(rows).T
            out_col, src_col = np.asarray(columns).T
            output[np.ix_(out_row, out_col)] = self.deltas[np.ix_(src_row, src_col)]
            mask[np.ix_(out_row, out_col)] = self.mask[np.ix_(src_row, src_col)]
        return output, mask

    def without_targets(self, excluded: Iterable[str]) -> "ExternalSource":
        mask = self.mask.copy()
        mask[np.isin(self.target_ids, list(excluded))] = False
        return ExternalSource(self.source_id, self.target_ids, self.genes,
                              self.deltas.copy(), mask, self.scale, self.metadata.copy())


def load_replogle(path: Path, audit_object: dict, output_genes: Sequence[str],
                   target_ids: Sequence[str], *,
                   output_ensembl_ids: Mapping[str, str] | None = None,
                   target_gene_ids: Mapping[str, str] | None = None,
                   raw_transform: str = "log1p", construct_aggregation: str = "cell_weighted",
                   verify_file: bool = True) -> ExternalSource:
    """Subtract legal core controls per construct before target aggregation.

    Only explicitly audited construct rows are consumed. All declared HDF5 axis
    fingerprints and row identities are verified against the real file. Raw
    log1p is applied to population profiles, without integer-count checks,
    single-cell normalization, or reconstructing cell-level Wilcoxon tests.
    """
    import h5py

    path = Path(path)
    output_genes, target_ids = _ids(output_genes, "output genes"), _ids(target_ids, "target ids")
    if raw_transform not in ("log1p", "identity"):
        raise ValueError("Supported raw transforms are log1p and identity")
    if construct_aggregation not in ("cell_weighted", "equal"):
        raise ValueError("Declare equal or cell_weighted construct aggregation")
    scale = audit_object["source_scale"]
    if scale not in (RAW_SCALE, Z_SCALE) or audit_object.get("is_singlecell") is not False:
        raise ValueError("An explicitly identified raw or GEM-Z pseudobulk source is required")
    integrity = {"file_bytes": path.stat().st_size, "file_sha256": file_hash(path)}
    if verify_file:
        if path.stat().st_size != audit_object["size"]:
            raise ValueError("Source file byte count differs from publisher manifest")
        integrity["file_md5"] = file_hash(path, "md5")
        if integrity["file_md5"] != audit_object["md5"]:
            raise ValueError("Source file MD5 differs from publisher manifest")
    audit = audit_object["metadata"]
    with h5py.File(path, "r") as data:
        labels, genes = _index(data["obs"]), _index(data["var"])
        symbols = tuple(map(str, _column(data["var"], "gene_name")))
        for key, values in (("target_axis_sha256", labels), ("gene_axis_sha256", genes),
                            ("gene_symbol_axis_sha256", symbols)):
            if axis_hash(values) != audit[key]:
                raise ValueError(f"Source {key} differs from inspected axis")
        if tuple(data["X"].shape) != tuple(audit["shape"]) or data["X"].shape != (len(labels), len(genes)):
            raise ValueError("Source matrix shape differs from manifest")
        if not isinstance(data["X"], h5py.Dataset) or data["X"].dtype.kind != "f":
            raise ValueError("Expected a dense floating-point population profile")
        original_controls = np.flatnonzero(_booleans(_column(data["obs"], "core_control")))
        if not len(original_controls) or len(original_controls) != audit["core_control_rows"]:
            raise ValueError("Core-control reference is absent or differs from audit")
        counts = np.asarray(_column(data["obs"], "num_cells_filtered"), dtype=np.float64)
        if len(counts) != len(labels):
            raise ValueError("Source population cell-count axis has incorrect length")
        eligible_population = np.isfinite(counts) & (counts > 0)
        controls = original_controls[eligible_population[original_controls]]
        excluded_controls = [dict(row=int(row), target="non-targeting", reason="invalid_population_cell_count",
                                  observed_num_cells_filtered=float(counts[row]) if np.isfinite(counts[row]) else None)
                             for row in original_controls if not eligible_population[row]]
        if not len(controls):
            raise ValueError("No core-control population has a finite positive cell count")
        selected: dict[str, list[dict]] = {target: [] for target in target_ids}
        excluded, declared_rows, requested_construct_rows = [], set(), set()
        for declared in audit["matched_target_constructs"]:
            row = int(declared["row"])
            if row in declared_rows or not 0 <= row < len(labels):
                raise ValueError("Construct mapping repeats a row or is out of range")
            declared_rows.add(row)
            parsed = parse_construct(labels[row], row)
            if parsed is None or any(parsed.get(key) != declared.get(key) for key in parsed):
                raise ValueError("Audited construct identity differs from the real observation label")
            target = declared["symbol"]
            if target not in selected:
                continue
            requested_construct_rows.add(row)
            expected_id = _ensembl((target_gene_ids or {}).get(target))
            if (target == "TAZ" and not expected_id) or (expected_id and expected_id != _ensembl(declared["ensembl_id"])):
                excluded.append(dict(target=target, row=row, reason="unresolved_or_conflicting_target_ensembl"))
                continue
            if row in original_controls:
                raise ValueError("A declared perturbation construct is an original core-control row")
            if not eligible_population[row]:
                excluded.append(dict(target=target, row=row, reason="invalid_population_cell_count",
                                     observed_num_cells_filtered=float(counts[row]) if np.isfinite(counts[row]) else None))
                continue
            selected[target].append(declared)
        selected_rows = sorted(set(controls.tolist()) | {item["row"] for group in selected.values() for item in group})
        profiles = np.asarray(data["X"][selected_rows, :], dtype=np.float64)
        if not np.isfinite(profiles).all():
            raise ValueError("Selected bulk profiles contain nonfinite values")
        transform = raw_transform if scale == RAW_SCALE else "identity_gem_z"
        if scale == RAW_SCALE:
            if np.any(profiles < 0):
                raise ValueError("Raw bulk profiles contain negative expression")
            if raw_transform == "log1p":
                profiles = np.log1p(profiles)
        row_lookup = {row: i for i, row in enumerate(selected_rows)}
        control_weights = counts[controls]
        if np.sum(control_weights) <= 0:
            raise ValueError("Core control populations have no cells")
        # Common reference uses only author-declared core-control populations.
        reference = np.average(profiles[[row_lookup[row] for row in controls]],
                               axis=0, weights=control_weights)
        gene_indices, gene_audit = gene_mapping(genes, symbols, output_genes, output_ensembl_ids)
        valid_genes = gene_indices >= 0
        deltas = np.full((len(target_ids), len(output_genes)), np.nan)
        mask = np.zeros(deltas.shape, dtype=bool)
        construct_log = {}
        for i, target in enumerate(target_ids):
            constructs = selected[target]
            if not constructs:
                continue
            rows = [item["row"] for item in constructs]
            weights = counts[rows] if construct_aggregation == "cell_weighted" else np.ones(len(rows))
            construct_deltas = profiles[[row_lookup[row] for row in rows]] - reference
            delta = np.average(construct_deltas, axis=0, weights=weights)
            deltas[i, valid_genes] = delta[gene_indices[valid_genes]]
            mask[i, valid_genes] = True
            construct_log[target] = dict(constructs=constructs, cell_counts=counts[rows].tolist(),
                                         normalized_weights=(weights / weights.sum()).tolist())
    metadata = dict(**integrity, file_id=audit_object.get("file_id"),
                    source_context=dict(cell_line=audit_object["cell_line"], time_days=audit_object["time_days"]),
                    transform=transform, profile_unit="population expression; not single-cell integer counts",
                    raw_cell_normalization_applied=False, core_control_rows=original_controls.tolist(),
                    eligible_control_rows=controls.tolist(), excluded_control_rows=excluded_controls,
                    eligible_control_normalized_weights=(control_weights / control_weights.sum()).tolist(),
                    population_filtering_policy=POPULATION_FILTER_POLICY,
                    invalid_unused_population_rows=[int(row) for row in np.flatnonzero(~eligible_population)
                                                    if row not in set(original_controls.tolist()) | requested_construct_rows],
                    core_control_weighting="num_cells_filtered", construct_aggregation=construct_aggregation,
                    construct_rule="transform each profile; subtract core-control mean per construct; aggregate deltas",
                    constructs=construct_log, excluded_constructs=excluded, gene_mapping=gene_audit,
                    source_target_axis_sha256=axis_hash(labels), source_gene_axis_sha256=axis_hash(genes),
                    output_gene_axis_sha256=axis_hash(output_genes), target_axis_sha256=axis_hash(target_ids),
                    h1_perturbed_outcomes_accessed=False, official_score=False)
    return ExternalSource(audit_object["id"], target_ids, output_genes, deltas, mask, scale, metadata)


def save_source(source: ExternalSource, path: Path) -> None:
    path = Path(path)
    path.parent.mkdir(parents=True, exist_ok=True)
    np.savez_compressed(path, source_id=np.array(source.source_id), target_ids=np.asarray(source.target_ids),
                        genes=np.asarray(source.genes), deltas=source.deltas.astype(np.float32),
                        mask=source.mask, scale=np.array(source.scale),
                        metadata_json=np.array(json.dumps(source.metadata, sort_keys=True)))


def read_source(path: Path) -> ExternalSource:
    with np.load(path, allow_pickle=False) as values:
        return ExternalSource(str(values["source_id"]), tuple(values["target_ids"]), tuple(values["genes"]),
                              values["deltas"], values["mask"], str(values["scale"]),
                              json.loads(str(values["metadata_json"])))


@dataclass(frozen=True)
class TransferConfig:
    kind: str = "low_rank_ridge"
    rank: int = 16
    alpha: float = 1.0
    shrinkage: float = 1.0
    min_source_targets: int = 3

    def __post_init__(self):
        if self.kind not in ("low_rank_ridge", "scalar", "diagonal"):
            raise ValueError("Unknown transfer model kind")
        if self.rank < 1 or self.alpha <= 0 or not 0 <= self.shrinkage <= 1 or self.min_source_targets < 2:
            raise ValueError("Invalid rank, regularization, shrinkage or minimum source count")


class TransferExpert:
    """Train source-specific mappings and blend only available source blocks.

    Fit accepts an outcome matrix but selects explicit fit_target_ids before
    inspecting values. Pass a physically filtered outcome artifact in production.
    Prediction has no truth argument. Missing source is the fit-target mean
    H1 response, not a zero-response claim. Source masks select observations;
    centered imputation is used only inside a fitted latent representation.
    """
    def __init__(self, config: TransferConfig | Mapping | None = None):
        self.config = config if isinstance(config, TransferConfig) else TransferConfig(**(config or {}))
        self.models: dict[str, dict] = {}

    def fit(self, sources: Sequence[ExternalSource], response_target_ids: Sequence[str],
            response_genes: Sequence[str], response_deltas: np.ndarray, *,
            fit_target_ids: Sequence[str]) -> "TransferExpert":
        response_ids = _ids(response_target_ids, "response targets")
        self.genes = _ids(response_genes, "response genes")
        self.fit_target_ids = _ids(fit_target_ids, "fit targets")
        if len(self.fit_target_ids) < 2:
            raise ValueError("At least two explicit fit targets are required")
        response_deltas = np.asarray(response_deltas)
        if response_deltas.shape != (len(response_ids), len(self.genes)):
            raise ValueError("Response matrix does not match its declared axes")
        response_lookup = {target: i for i, target in enumerate(response_ids)}
        if not set(self.fit_target_ids) <= set(response_lookup):
            raise ValueError("Fit identity absent from response axis")
        # No heldout value, target-wide response statistic or validation row is inspected.
        y = np.asarray(response_deltas[[response_lookup[target] for target in self.fit_target_ids]], dtype=np.float64).copy()
        if not np.isfinite(y).all():
            raise ValueError("Fit target responses contain nonfinite values")
        self.common_response = y.mean(axis=0)
        self.models = {}
        for source in sources:
            if source.source_id in self.models:
                raise ValueError("Source blocks must have unique IDs, including context/time/scale")
            x, mask = source.aligned(self.fit_target_ids, self.genes)
            rows = mask.any(axis=1)
            record = dict(scale=source.scale, available_fit_targets=int(rows.sum()),
                          fit_source_target_ids=[target for target, valid in zip(self.fit_target_ids, rows) if valid],
                          enabled=False)
            self.models[source.source_id] = record
            if rows.sum() < self.config.min_source_targets:
                continue
            x, mask, ys = x[rows], mask[rows], y[rows]
            count = mask.sum(axis=0)
            center = np.divide(np.where(mask, x, 0).sum(axis=0), count,
                               out=np.zeros(len(self.genes)), where=count > 0)
            centered = np.where(mask, x - center, 0.0)
            variance = np.divide(np.square(centered).sum(axis=0), count,
                                 out=np.zeros(len(self.genes)), where=count > 0)
            scale = np.where(variance > 1e-24, np.sqrt(variance), 1.0)
            supported_genes = count >= 2
            offset = ys.mean(axis=0) - self.common_response
            yc = ys - ys.mean(axis=0)
            record.update(enabled=True, center=center, spread=scale, supported_genes=supported_genes,
                          offset=offset, fit_observation_counts=count, kind=self.config.kind)
            if self.config.kind == "low_rank_ridge":
                design = centered / scale
                design[:, ~supported_genes] = 0
                # Thin source PCA via the n_fit x n_fit Gram, never H1 heldout outcomes.
                eigenvalues, u = np.linalg.eigh(design @ design.T)
                order = np.argsort(eigenvalues)[::-1]
                eigenvalues, u = eigenvalues[order], u[:, order]
                keep = np.flatnonzero(eigenvalues > max(1e-12, float(eigenvalues[0]) * 1e-12))[:self.config.rank]
                if len(keep):
                    components = (design.T @ u[:, keep] / np.sqrt(eigenvalues[keep])).T
                    latent = design @ components.T
                    coefficients = np.linalg.solve(latent.T @ latent + self.config.alpha * len(x) * np.eye(len(keep)),
                                                   latent.T @ yc)
                else:
                    components = np.zeros((0, len(self.genes)))
                    coefficients = np.zeros((0, len(self.genes)))
                record.update(components=components, coefficients=coefficients)
            elif self.config.kind == "diagonal":
                weights = np.sum(centered * yc, axis=0) / (np.sum(centered**2, axis=0) + self.config.alpha * len(x))
                weights[~supported_genes] = 0
                record["coefficients"] = weights
                record["offset"] = np.where(supported_genes, offset, 0)
            else:
                eligible = centered[:, supported_genes]
                record["coefficients"] = float(np.sum(eligible * yc[:, supported_genes]) /
                                                (np.sum(eligible**2) + self.config.alpha * len(x)))
                record["offset"] = np.where(supported_genes, offset, 0)
        return self

    def predict(self, sources: Sequence[ExternalSource], *, query_ids: Sequence[str],
                genes: Sequence[str] | None = None, strict_unseen: bool = False,
                return_diagnostics: bool = False):
        if not hasattr(self, "common_response"):
            raise ValueError("Fit the model before prediction")
        query_ids = _ids(query_ids, "query targets")
        sources_by_id = {source.source_id: source for source in sources}
        if len(sources_by_id) != len(sources):
            raise ValueError("Duplicate source ID")
        total = np.zeros((len(query_ids), len(self.genes)))
        contributors = np.zeros(total.shape, dtype=int)
        source_available = {}
        for source_id, record in self.models.items():
            source = sources_by_id.get(source_id)
            if source is None or not record["enabled"]:
                source_available[source_id] = [False] * len(query_ids)
                continue
            if source.scale != record["scale"]:
                raise ValueError("Source scale changed after fitting")
            x, mask = source.aligned(query_ids, self.genes)
            if strict_unseen:
                mask[:] = False
            mask &= record["supported_genes"]
            available = mask.any(axis=1)
            source_available[source_id] = available.tolist()
            centered = np.where(mask, x - record["center"], 0.0)
            if self.config.kind == "low_rank_ridge":
                delta = (centered / record["spread"]) @ record["components"].T @ record["coefficients"] + record["offset"]
                contribution_mask = np.broadcast_to(available[:, None], total.shape)
            else:
                delta = centered * record["coefficients"] + record["offset"]
                contribution_mask = mask
            total += np.where(contribution_mask, delta, 0)
            contributors += contribution_mask
        deviation = np.divide(total, contributors, out=np.zeros_like(total), where=contributors > 0)
        prediction = self.common_response + self.config.shrinkage * deviation
        diagnostics = dict(query_ids=list(query_ids), source_available=source_available,
                           contributing_sources_per_target=[
                               {str(count): int(np.sum(row == count)) for count in np.unique(row)}
                               for row in contributors],
                           fallback_gene_counts=np.sum(contributors == 0, axis=1).tolist(),
                           n_output_genes=len(self.genes),
                           fallback="training-fold target mean response", strict_unseen=bool(strict_unseen),
                           fit_target_ids=list(self.fit_target_ids), outcome_accessed=False)
        if genes is not None:
            requested = _ids(genes, "prediction gene axis")
            lookup = {gene: i for i, gene in enumerate(self.genes)}
            if not set(requested) <= set(lookup):
                raise ValueError("Prediction requested an unfitted expression coordinate")
            prediction = prediction[:, [lookup[gene] for gene in requested]]
        return (prediction, diagnostics) if return_diagnostics else prediction


def target_folds(target_ids: Sequence[str], n_splits: int = 3, seed: int = 20251013):
    """Split whole perturbation targets, never individual constructs/cells."""
    target_ids = _ids(target_ids, "CV target axis")
    if not 2 <= n_splits <= len(target_ids):
        raise ValueError("CV requires at least one target per heldout fold")
    parts = np.array_split(np.random.default_rng(seed).permutation(len(target_ids)), n_splits)
    return [(tuple(target_ids[i] for i in np.sort(np.concatenate([part for j, part in enumerate(parts) if j != fold]))),
             tuple(target_ids[i] for i in np.sort(query))) for fold, query in enumerate(parts)]


def nested_target_cv(sources: Sequence[ExternalSource], target_ids: Sequence[str],
                     genes: Sequence[str], deltas: np.ndarray, configs: Sequence[Mapping], *,
                     control: np.ndarray | None = None, outer_folds: int = 3, inner_folds: int = 3,
                     seed: int = 20251013, strict_unseen: bool = False) -> tuple[np.ndarray, dict]:
    """Inner fitting/selection uses only outer-fit outcomes; outer evaluation follows commit.

    Reports target MAE diagnostics in the declared modeling space. These are not
    the historical competition DES/PDS/MAE or normalized leaderboard aggregate.
    """
    target_ids, genes = _ids(target_ids, "CV targets"), _ids(genes, "CV genes")
    deltas = np.asarray(deltas, dtype=np.float64)
    if deltas.shape != (len(target_ids), len(genes)) or not np.isfinite(deltas).all():
        raise ValueError("CV requires a finite training-only outcome matrix with declared axes")
    configurations = [TransferConfig(**configuration) for configuration in configs]
    if not configurations:
        raise ValueError("Register a nonempty candidate grid before CV")
    lookup = {target: i for i, target in enumerate(target_ids)}
    baseline = np.empty_like(deltas)
    predictions = np.empty_like(deltas)
    records = []

    def clip(delta):
        return np.maximum(delta + control, 0) - control if control is not None else delta

    for fold, (fit, query) in enumerate(target_folds(target_ids, outer_folds, seed)):
        fit_rows = [lookup[target] for target in fit]
        # Physically select the outer fit labels before calling inner selectors.
        fit_y = deltas[fit_rows].copy()
        fit_lookup = {target: i for i, target in enumerate(fit)}
        inner = target_folds(fit, inner_folds, seed + fold + 1)
        losses = []
        for config in configurations:
            errors = []
            for inner_fit, inner_query in inner:
                inner_sources = [source.without_targets(inner_query) for source in sources] if strict_unseen else sources
                model = TransferExpert(config).fit(inner_sources, fit, genes, fit_y, fit_target_ids=inner_fit)
                predicted = clip(model.predict(inner_sources, query_ids=inner_query, strict_unseen=strict_unseen))
                inner_truth = fit_y[[fit_lookup[target] for target in inner_query]]
                errors.extend(np.mean(np.abs(predicted - inner_truth), axis=1).tolist())
            losses.append(float(np.mean(errors)))
        chosen = int(np.argmin(losses))
        outer_sources = [source.without_targets(query) for source in sources] if strict_unseen else sources
        model = TransferExpert(configurations[chosen]).fit(outer_sources, fit, genes, fit_y, fit_target_ids=fit)
        predicted, diagnostics = model.predict(outer_sources, query_ids=query, strict_unseen=strict_unseen,
                                               return_diagnostics=True)
        rows = [lookup[target] for target in query]
        predictions[rows] = clip(predicted)
        baseline[rows] = clip(np.broadcast_to(model.common_response, predicted.shape))
        records.append(dict(fold=fold, fit_target_ids=list(fit), query_ids=list(query),
                            inner_mean_target_mae=losses, chosen_candidate=chosen,
                            chosen_config=vars(configurations[chosen]), prediction_diagnostics=diagnostics,
                            diagnostic_mae=float(np.mean(np.abs(predictions[rows] - deltas[rows])))))
    report = dict(expert="E1_external_pseudobulk_transfer", outer_folds=outer_folds, inner_folds=inner_folds,
                  seed=seed, track="strict_all_context_target_unseen" if strict_unseen else "non_H1_same_target_transfer",
                  official_score=False, benchmark_comparable=False, validation_test_outcomes_accessed=False,
                  diagnostic_target_mae=np.mean(np.abs(predictions - deltas), axis=1).tolist(),
                  diagnostic_mae=float(np.mean(np.abs(predictions - deltas))),
                  common_response_diagnostic_mae=float(np.mean(np.abs(baseline - deltas))),
                  candidate_grid=[vars(config) for config in configurations], folds=records)
    return predictions, report
```
