## Supplementary figures and images for "Centered cross-context perturbation transfer on the public 2025 Virtual Cell Challenge benchmark"

### final_v5_center_ablation_20261021.png

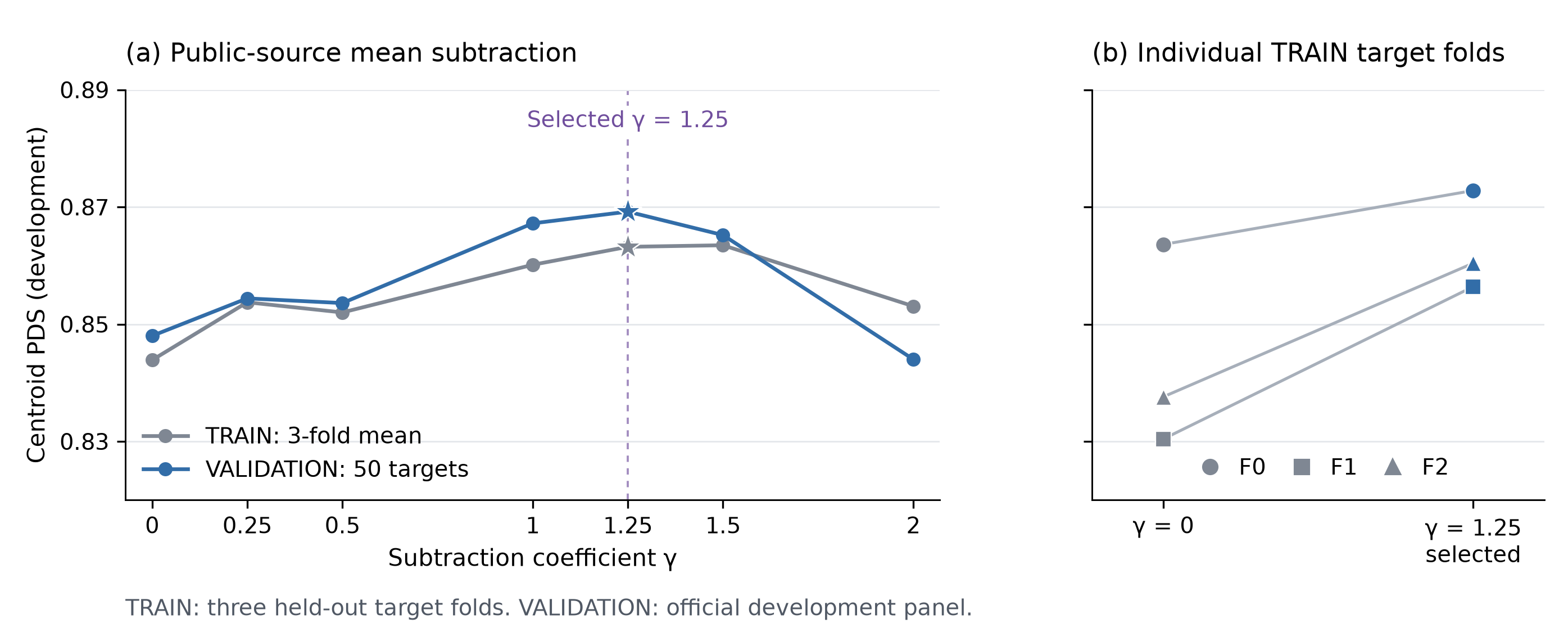

### final_v5_comparison_20261021.png

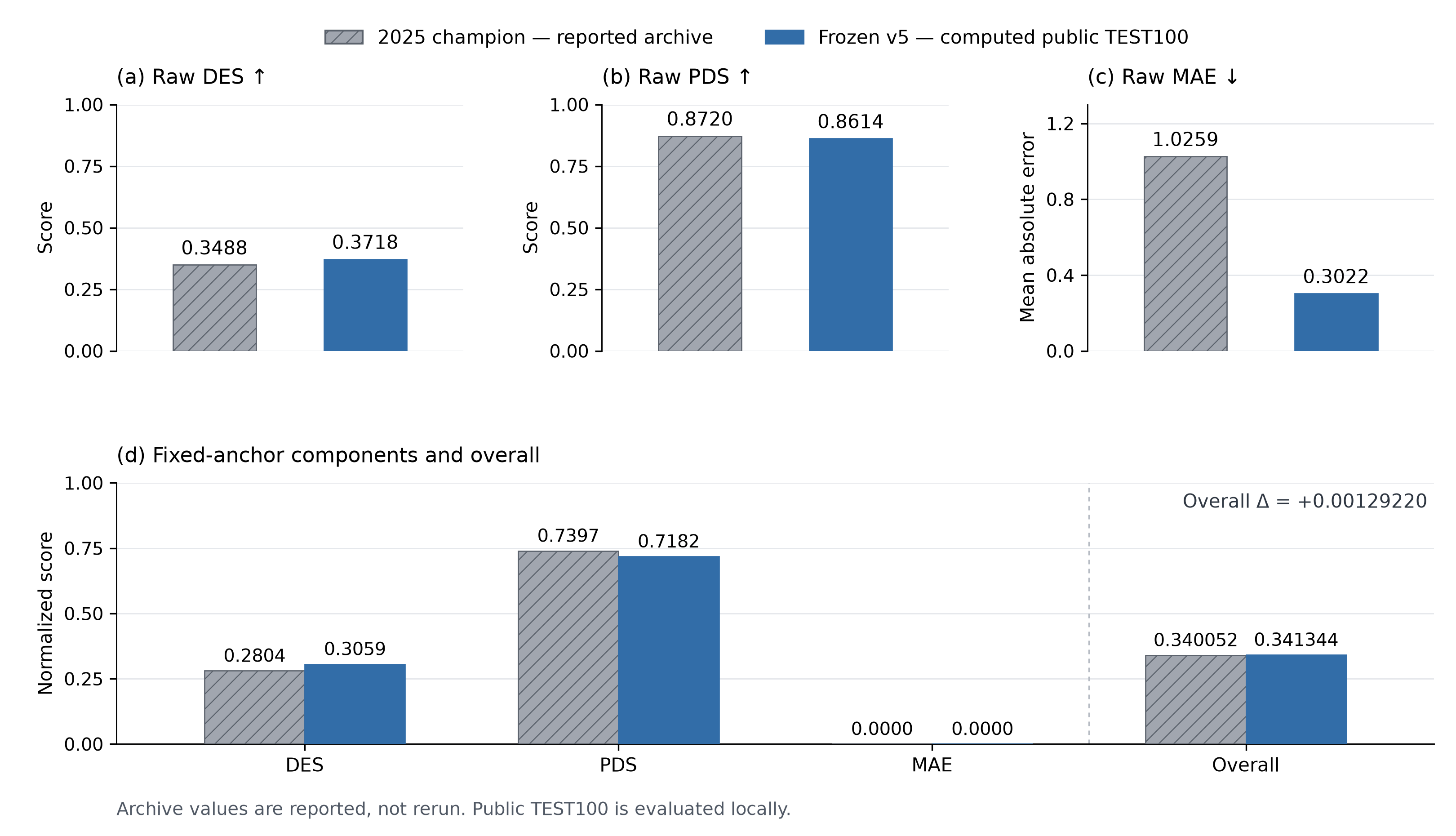
